# Bacterial growth in multicellular aggregates leads to the emergence of complex lifecycles

**DOI:** 10.1101/2021.11.01.466752

**Authors:** Julia A Schwartzman, Ali Ebrahimi, Grayson Chadwick, Yuya Sato, Victoria Orphan, Otto X Cordero

## Abstract

In response to environmental stresses such as starvation, many bacteria facultatively aggregate into multicellular structures that can attain new metabolic functions and behaviors. Despite the ubiquity and relevance of this form of collective behavior, we lack an understanding of how the spatiotemporal dynamics of aggregate development emerge from cellular physiology. Here, we show that the coupling between growth and spatial gradient formation leads to the emergence of a complex lifecycle, akin to those known for multicellular bacteria. Under otherwise carbon-limited growth conditions, the marine bacterium *Vibrio splendidus* 12B01 forms multicellular groups to collectively harvest carbon from the brown-algal polysaccharide alginate. This is achieved during growth on dissolved alginate polymer through formation of spherical, clonal clusters of cells that grow up to 40 µm in radius. Clusters develop striking spatial patterning as they grow due to phenotypic differentiation of sub-populations into a ‘shell’ of static cells surrounding a motile ‘core’. Combining *in situ* measurements of cell physiology with transcriptomics, we show that shell cells express adhesive type IV pili, while motile core cells express carbon storage granules. The emergence of shell and core phenotypes is cued by opposing gradients of carbon and nitrogen that form within cell clusters due to local metabolic activity. Eventually, the shell ruptures, releasing the carbon-storing core, and we show that carbon-storing cells more readily propagate on alginate than non-carbon storing cells. We propose that phenotypic differentiation promotes the resilience of 12B01 groups by enabling clonal groups to grow larger and propagate more effectively. Phenotypic differentiation may be a widespread, but overlooked, strategy among bacteria to enhance resilience in the context of resource limitation.

## INTRODUCTION

Clonal groups of microbes often form transient multicellular structures such as biofilms and fruiting bodies (Claessen et al., 2014; Shapiro, 1998). The emergent behaviors of these structures underly ecologically important functions such as antimicrobial resistance (Høiby et al., 2010), immune evasion (Arciola et al., 2018), and the mobilization of complex nutrients (Koschwanez et al., 2013; Ratzke and Gore, 2016). We typically consider only the benefits of life in a group. However, life at high cell density is also an inevitable source of cellular conflict. Even where cells cooperate by sharing metabolic tasks such as resource mobilization or detoxification, they compete for access to other growth-essential nutrients. Thus, cells in transient multicellular states must balance cooperation and conflict for the population to derive and ecological benefit. One possible mechanism through which this is achieved within clonal populations is the emergence of phenotypic heterogeneity. There is a growing understanding that phenotypic heterogeneity is a ubiquitous aspect of microbial populations (Dar et al., 2021; Vlamakis et al., 2013). However, we have little understanding of when heterogeneity provides an ecological benefit (van Gestel et al., 2015). Here, we explore how the emergence of phenotypic heterogeneity in transient multicellular structures formed by a marine bacterium enhances resource partitioning and gives rise to complex reproductive cycles.

The focus of our study is transient multicellular behaviors that emerge in the context of decomposition. Decomposers have an ecological advantage in resource poor environments because they can access otherwise recalcitrant nutrients (Sichert and Cordero, 2021). Because decomposition takes place outside of cells, the ability to pool enzymes accelerates the rate of decomposition, and in turn, enhances the recovery of breakdown products. It is common for this behavior to be positively dependent on cell density; strong positive-density dependence results in critical population thresholds where cells that cannot create local density die off (Ratzke and Gore, 2016). In this context, the ecological benefit of local cell density can exert strong evolutionary selection for multicellular behavior (Koschwanez et al., 2011), suggesting that multicellular behaviors are likely widespread among degraders in nature (D’Souza et al., 2021).

Here, we demonstrate how phenotypic heterogeneity within multicellular groups shapes resource allocation, growth, and reproduction in marine polysaccharide decomposer. The Gammaproteobacterium *Vibrio splendidus* 12B01 grows on the brown-algae derived polysaccharide alginate through the formation of cooperative multicellular clusters. Cells in the clusters exist in a mix of phenotypic states and form reproducible spherical structures that grow up to 40 µm in radius and develop mixing in their cores. In previous work, we have shown that growth of 12B01 is strongly dependent on local population density (Ebrahimi et al., 2019a; Hehemann et al., 2016). Using a combination of transcriptomics, quantitative light microscopy and single cell stable isotope probing we characterized the phenotypic differentiation that occurs within 12B01 groups. We show that phenotypic differentiation allows groups to form self-organized structures that promote sharing of polysaccharide-derived carbon and mitigate competition for other growth essential nutrients. Our results highlight the role of phenotypic differentiation in the context of polysaccharide decomposition: a common microbial group behavior.

## RESULTS

Alginate decomposition is a common trait among marine *Vibrio* (Hehemann et al., 2016), and the cellular mechanisms by which 12B01 decomposes alginate are well characterized (Badur et al., 2017, 2015; Jagtap et al., 2014; Wargacki et al., 2012). A striking feature of 12B01 growth on dissolved alginate is the emergence of motility within cell clusters (Ebrahimi et al., 2019a). To better understand the process leading to the emergence of motility, we monitored the development of clusters during growth of 12B01 on alginate in shaking flasks. The population (composed initially of solitary cells) underwent a long lag phase before tightly packed clusters of cells started to form and population growth was detectable. Clusters underwent three distinct stages as they grew: i) initial growth as structurally homogenous clusters (Figure 1A), ii) self-organization of clusters into layered structures composed of an outer sub-population of cells with clear cell-cell adhesion, and an inner motile sub-population of cells (Figure 1B, Movie S1), and iii) rupture of layered structures, releasing motile internal contents (Figure 1C). This behavior is reminiscent of the hollowing and seeding dispersal observed in diverse biofilm-forming bacteria (Kaplan, 2010). Stages corresponded to changes in cluster growth rate: clusters in stage i grew slowly compared to stage ii (Figure 1D), suggesting that the amount of cellular cooperation increases from stage i to stage ii, and then ceases upon rupture.

**Figure 1.**
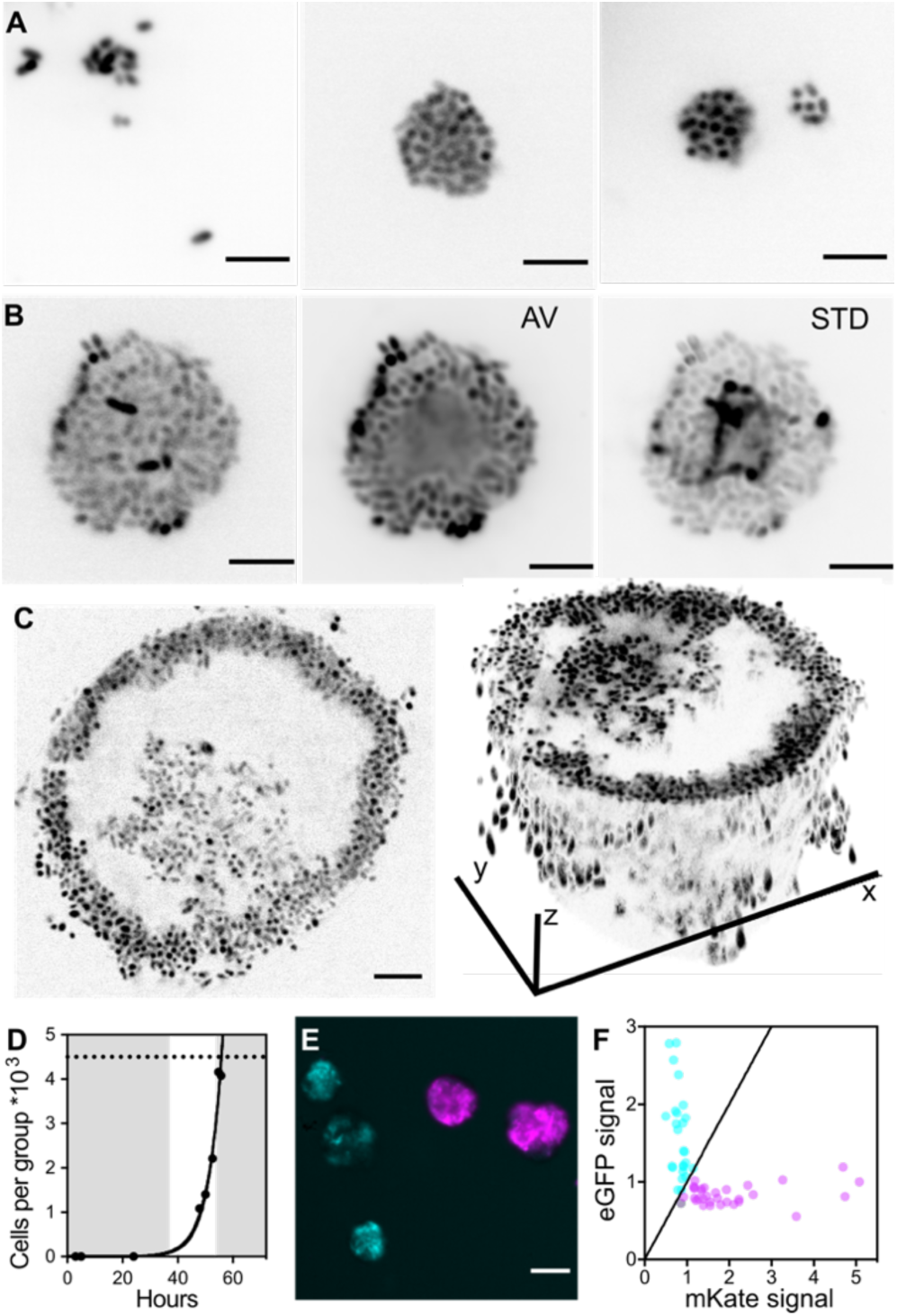
12B01 growth on alginate involves formation of phenotypically distinct sub-populations by clonal groups. **A-C**) Stages of 12B01 growth on alginate. Scale bars are 5 µm. **A**) In stage i, single cells are suspended in medium containing soluble alginate polysaccharides, and slowly form undifferentiated groups. Panels show three examples of stage i groups. **B**) In stage ii, groups grow and differentiate into layered structures, with a solid shell and a mobile core. Panels show mixing in the center of groups. AV; average intensity projection of time-lapse, showing populations of cells that are motile and not motile (Movie S1); STD, standard deviation projection, showing differences in pixel intensity between frames. **C**) Rupture of clusters leads to hollowing in stage iii. Left: optical section through a cluster, showing absence of cells in center; right: 3D projection, showing structure of hollowed cluster. **D**) Growth of 12B01 in groups, expressed as cells per group. Grey shaded regions delineate growth stages. Exponential growth rate of cells in stage ii (fit line): µmax=0.19 +/−0.03 h^−1^. Dashed line indicates size at which growth of clusters ceased due to rupture. **E**) Formation of 12B01 groups tracked using isogenic strains that express fluorescent proteins (Figure S2A). Representative images show distribution of fluorescent protein among eGFP (cyan) or mKate (magenta)-labeled groups at stage ii. Scale bar= 20 µm. **F**) Quantification of fluorescence intensity within 12B01 groups. Each point represents the total intensity per group, normalized by the mean intensity of groups in each channel. The black diagonal line represents identify (equal proportion of both strains).

Self-organization emerges from local cellular interactions. Both physical interactions with alginate polysaccharide, and physiological changes resulting from the availability of alginate-derived carbon, could contribute to self-organization of 12B01. We investigated these ideas by growing cells on alginate polysaccharide, uronic acids mannuronate and guluronate (metabolizable break-down products of alginate), or a mix of polysaccharide and breakdown products. Although the presence of alginate polysaccharide was sufficient to induce cellular aggregation, aggregates forming in the presence of the breakdown products differed greatly in their morphology from the compact clusters formed when only polysaccharide was available (Figure S1A). Staining with a cell-impermeant fluorescent dye specific for DNA revealed that the loose biofilm-like aggregates contained dead cells and extracellular DNA, whereas the clusters formed during growth on alginate polymer only did not (Figure S1B). These results indicated that, while cellular interactions with alginate polysaccharide are likely to take place, alginate polysaccharide must be the only available carbon source for 12B01 to develop morphological complexity.

Biofilms, swarms, and other bacterial social structures often form by the coalescence of many individual cells. Unlike clonal structures, conflict can emerge within aggregative social structures due to the presence of multiple independent genotypes (Grosberg and Strathmann, 2007; Márquez-Zacarías et al., 2021; Nadell et al., 2016; Pentz et al., 2020). This led us to investigate whether 12B01 clusters were aggregative or clonal. We mixed isogenic populations of 12B01 each expressing a fluorescent protein and monitored the proportion of clusters that developed with each fluorescent protein, or with a mix of both (Figure 1E, Figure S2A). Strikingly, clusters expressed either one or the other fluorescent protein rather than forming mixed assemblages (Figure 1F, Figure S2B), demonstrating that the emergent cooperation of 12B01 clusters on alginate polymer is achieved by division of a progenitor cell into a clonal collective. The clonality of 12B01 cooperation indicates that individual progenitor cells can support initial divisions to create local cell density, and that cell clusters do not merge during development. One consequence of the clonality of 12B01 clusters is that cooperative resource sharing and phenotypic heterogeneity emerge from interactions among individuals that share high genetic relatedness: on average, the population undergoes fewer than 10 rounds of division prior to rupture.

To define the physiological differences between sub-populations, we profiled global transcription at different stages of 12B01 growth. The synchronous timing of cluster development in our experimental setup enabled collection of RNA from cells in early stage ii, cells released from late-stage iii clusters, and cells remaining in the shells of late-stage iii clusters (Figure 2A). These samples were chosen to maximize the homogeneity within the sampled population, and to capture populations from distinct stages of growth. Only 3% of the 12B01 transcriptome was differentially expressed between stage ii and stage iii free-living populations (Figure 2B, Table S1). About 10% of the transcriptome was differentially expressed more than 2-fold between samples and those collected from stage iii clusters and either of the other sample types (Figure 2B). Although the transcriptional state of free-living stage iii populations is likely to differ from that of cells belonging to the motile core of 12B01 clusters, samples were collected no more than 30 minutes after the first signs of rupture, as assessed by microscopy. Thus, the free-living population is likely to reflect some aspects of the transcriptional state of the ‘core’ subpopulation directly prior to rupture. We compared all pairs of samples to define sets of genes differentially more than 2-fold between conditions (Table S1).

**Figure 2.**
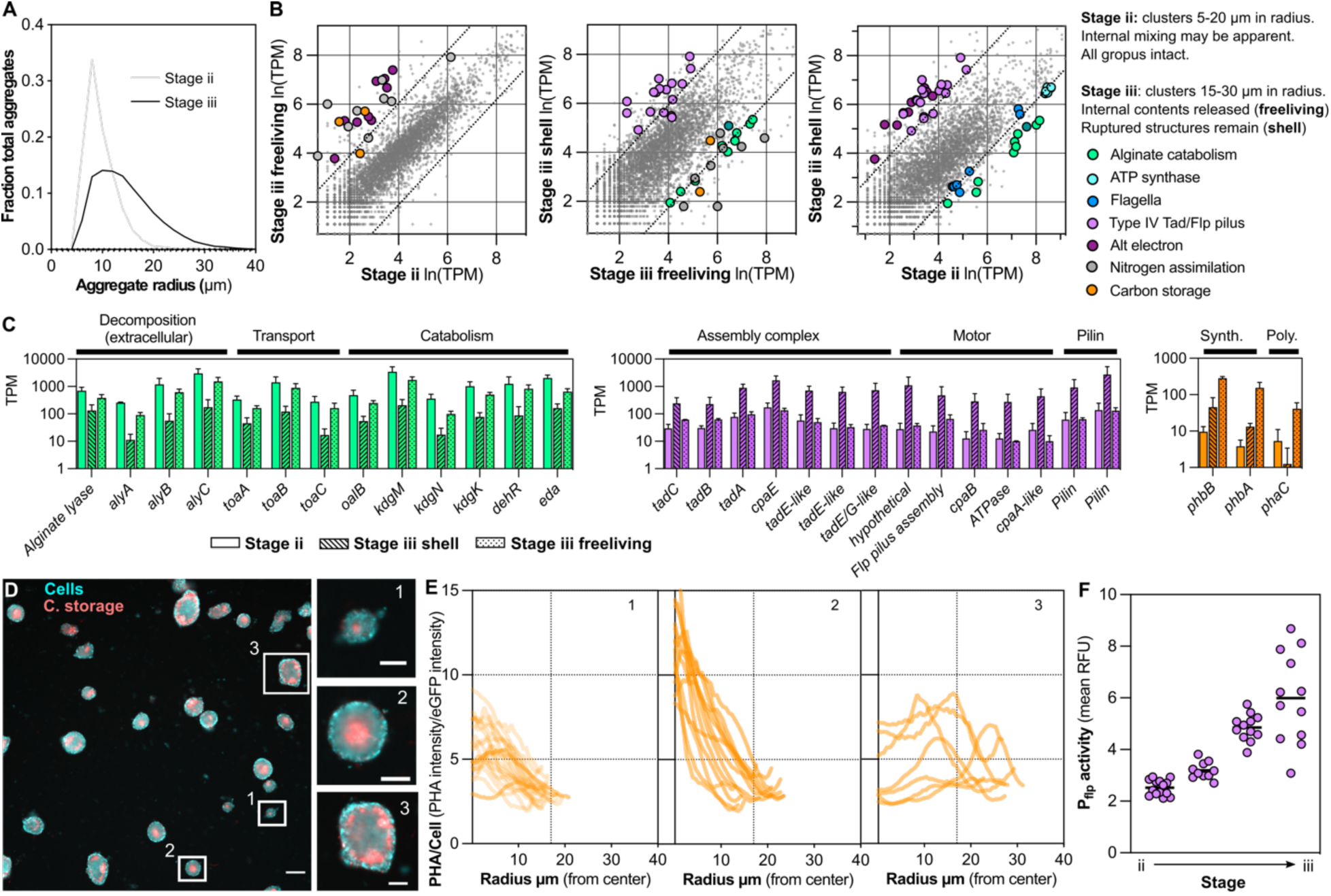
Transcriptional and physiological differences that define cluster sub-populations. **A**) Histograms showing the change in the distribution of cluster radii at different timepoints of 12B01 growth on alginate. ‘Stage ii’ represents a timepoint when most clusters are in the first stage of development, and ‘Stage ii’ represents a timepoint when most clusters are in stage ii of development. Clusters measured per timepoint: Stage ii=1085, Stage iii=6105. **B**) Pair-wise comparisons of normalized gene expression, showing patterns of differentially expressed genes that characterize each sample type. Points represent individual genes, plotted as the mean ln(TPM) for all replicates; stage iii free-living samples, n=2, stage iii shell and stage ii samples, n=3. Diagonal lines indicate threshold 2-fold differential expression. Highlighted genes are discussed in the text, and differentially expressed in the comparison. See Table S1 for detailed information on all differentially expressed genes, including P-values for fold change expression, and TPM (Transcripts per Kilobase Million) values for differentially expressed genes. **C**) Normalized transcript abundance (TPM) for gene clusters encoding key metabolic functions: alginate catabolism (left), type IV *tad/flp* pili (middle), carbon storage (right). Error bars are the standard deviation of 3 replicates (stage ii and stage iii shell samples), or 2 replicates (stage iii free-living samples). **D**) Accumulation of polyhydroxyalkanoate (PHA) in a subset of cells making up 12B01 clusters. Composite image showing localization of PHA (C. storage) in eGFP-labeled 12B01 (cells). Three distinct patterns of carbon storage granule formation are shown inset, right. Scale bar main image= 50 µm, scale bar inset=20 µm. **E)** Quantification of image in 2D, showing radial profiles of carbon storage intensity, normalized by cellular expression of GFP. Each trace represents a measurement from an individual cluster. Measurements are made from the cluster center (zero) to the periphery. Vertical dashed line notes 17 µm radial distance. Traces are grouped by their pattern of PHA accumulation: numbers indicate the patterns represented by inset images in Figure 2D. First panel, n=22, second panel n=13, third panel n=7. **F**) Transcriptional activity of *tad/flp* pili within clusters at timepoints during growth of 12B01 on alginate. Each data point represents the mean activity measured within an individual cluster. Timepoints were taken in the transition from most clusters being in stage ii to most clusters being in stage iii.

Genes differentially expressed in stage ii pointed to cellular processes underlying the fast growth and mixing observed in this stage. Relative to stage iii shell samples, stage ii samples more highly expressed genes encoding flagella (V12B01_05395-V12B01_05425), F1F0 ATP synthase (V12B01_01872-V12B01_01902), and the metabolic enzyme Eda (V12B01_24199): a key aldolase in the conversion of alginate-derived uronic acids into glycolytic metabolites via the Entner-Doudoroff pathway (Figure 2B, Figure 2C). Relative to either stage iii sample type, stage ii samples also transcribed lower levels of genes involved in alternative electron acceptor utilization, such as fumarate reductase (*frd* V12B01_23345-V12B01_23360), and nitrate reductase (*nrf*, V12B01_23499-V12B01_23529) (Figure 2B). In *Escherichia coli*, ATP synthase genes are subject to growth-rate dependent transcriptional regulation (Kasimoglu et al., 1996), and the *eda* gene is transcribed in response to uronic acids (Murray and Conway, 2005). By analogy to *E. coli*, the transcriptional activation of F1F0 ATP synthase and *eda* in stage ii relative to the stage iii shell could reflect more active growth on alginate oligosaccharides. In addition, the lower transcription of alternative electron acceptor genes in stage ii suggests that cells in this stage have more access to oxygen than cells in stage iii. Culturing 12B01 on alginate under anoxic conditions revealed that a consequence of oxygen limitation was diminished yield after 24 h of growth (Figure S1C). Together, the comparison of growth phenotypes and transcriptional signatures in stage ii suggest that the rapid growth and visible motility within stage ii clusters may be due to availability of both alginate and oxygen.

Genes differentially expressed at least 2-fold in the stage iii shell pointed to differences in alginate catabolism and cellular adhesion. Cells remaining in clusters during stage iii transcribed lower levels of alginate catabolism genes (V12B01_24194-V12B01_24214 and V12B01_24234-V12B01_24274) relative to other samples (Figure 2B, Figure 2C). Additionally, stage iii shell samples highly expressed a cluster of genes encoding a putative type IV Tad/Flp pilus (V12B01_22446-V12B01_22511, Figure 2C). Type IV pili are known to mediate adhesion in a variety of bacteria (McCallum et al., 2019), including social behaviors such as twitching motility in *Pseudomonas aeruginosa* (Burrows, 2012) and gliding in *Myxococcus xanthus* (Wu and Kaiser, 1995). To validate the type iii shell-specific transcription of *tad/flp* genes, we localized cellular transcription within 12B01 clusters using a fluorescent reporter construct (Figure S3A,B). Consistent with the transcriptional profiles, we found that transcription of tad/*flp* pilus genes were elevated in cells associated with late-stage iii clusters, relative to stage ii (Figure 2F). Together, the decreased transcription of alginate catabolism genes and increased transcription of *tad/flp* pilus prompted us to ask whether transcriptional regulation of this type IV pilus was growth-rate dependent. Transcriptional activity of *tad/flp* increased in cells as growth in batch culture entered stationary phase, a pattern not shared by the synthetic *tac* promoter (Figure S3C). The specific activation of the *tad/flp* promoter during stationary phase growth suggests that the transcriptional regulation of this gene cluster is responsive to cellular growth state. Together, these results suggest that the increase in transcription of the *tad/flp* pilus genes during development of 12B01 clusters may be linked to decreased availability of alginate-derived carbon and slower growth in the shell sub-population of stage iii clusters.

Differential gene expression specific to stage iii free-living samples highlighted genes related to nitrogen assimilation and carbon storage. These genes included ammonium transport (V12B01_25139), urease (V12B01_09226, V12B01_09231) glutamine synthase *glnA* (V12B01_02370), and nitrogen responsive regulator *glnGL* (V12B01_02360, V12B01_02355) (Table S1). In *E. coli*, transcription of *glnG* and *glnL* is activated by nitrogen limitation, and this regulator activates transcription of nitrogen assimilation genes (Pahel et al., 1982), suggesting that stage iii free-living cells may be nitrogen limited. A polyhydroxyalkanoate (PHA) polymerase (*phaC*, V12B01_15111), and associated PHA biosynthetic genes (*phbA* V12B01_15121, *phbB* V12B01_15126), were also highly transcribed by stage iii free-living samples (Figure 2B, Figure 2C). Together, *phbA, phbB*, and *phaC* encode machinery to synthesize carbon storage organelles called carbonosomes (Greening and Lithgow, 2020; Jendrossek, 2009). While *phbA* and *phbB* expression were highest in stage iii free-living samples, expression of these genes was also elevated in stage iii shell samples (Figure 2C). In contrast, *phaC* expression was found to be significantly elevated only in stage iii free-living samples (Figure 2C). Together, these results suggest that the ability to polymerize carbonosomes is a specific attribute of a sub-population of stage iii cells. We experimentally validated the prediction that carbon storage granules differentiate subpopulations within the cluster by staining clusters with the lipophilic fluorescent dye Nile Red (Figure 2D). We found that cells stained brightly by Nile Red, the presumptive PHA-storing sub-population, were indeed localized within the cluster core. Carbon storage was detectable only when clusters grew to a size larger than 17 µm in radius (Figure 2E), corresponding to late stage ii, as predicted from the global transcriptional profiles (Figure 2C). Together, these observations suggest that the microenvironment in the core of cell clusters, in which alginate-derived carbon is available in excess but ammonia-derived nitrogen is limiting, induces accumulation of PHA as clusters grow larger.

The accumulation of PHA within cells in the core of clusters, combined with transcriptional signatures hinting at nitrogen limitation in stage iii free-living cells, led us to hypothesize that nitrogen availability cues cells to accumulate carbon storage polymer. We tested this idea by measuring the yield of 12B01 during growth on alginate oligomers under differing concentrations of ammonium to define nitrogen-limited growth conditions (Figure 3A). Cells grown under nitrogen limitation and carbon excess accumulated more PHA, measured by intensity of Nile Red staining (Figure 3B). Numerical estimates and direct measurement of ammonium assimilation using stable isotope probing combined with nanoscale secondary ion mass spectrometry (NanoSIMS) showed that N gradients emerged within 12B01 cell clusters, and that the emergence of gradients occurred at a size predicted by cellular consumption rate. Importantly, our numerical estimate assumed that cells in clusters were phenotypically undifferentiated and did not move, a simplification that allowed us to approximate the length scale over which cellular consumption depleted ammonium supplied by diffusion from outside the cluster (Figure 3C). For clusters growing at the maximum observed rate of 0.2 h^−1^, radial cluster size reached 15-20 µm before cellular consumption depleted ammonium from the center of clusters (Figure 3C). Using NanoSIMS, we quantified the incorporation of heavy-labeled nitrogen into cellular biomass following four hours of incubation with ^15^N-ammonium (see methods, Figure S4). The NanoSIMS measurements revealed that cellular uptake of ammonium decreased towards the center of cell clusters (where PHA accumulated). The outer layers of cells assimilated more than twice the amount of ammonium than cells in the core of clusters (Figures 3D-E). The qualitative agreement between the gradient estimates and direct measurements of ammonium assimilation into biomass support the idea that cellular consumption creates local ammonium limitation within the center of growing clusters, favoring PHA accumulation.

**Figure 3.**
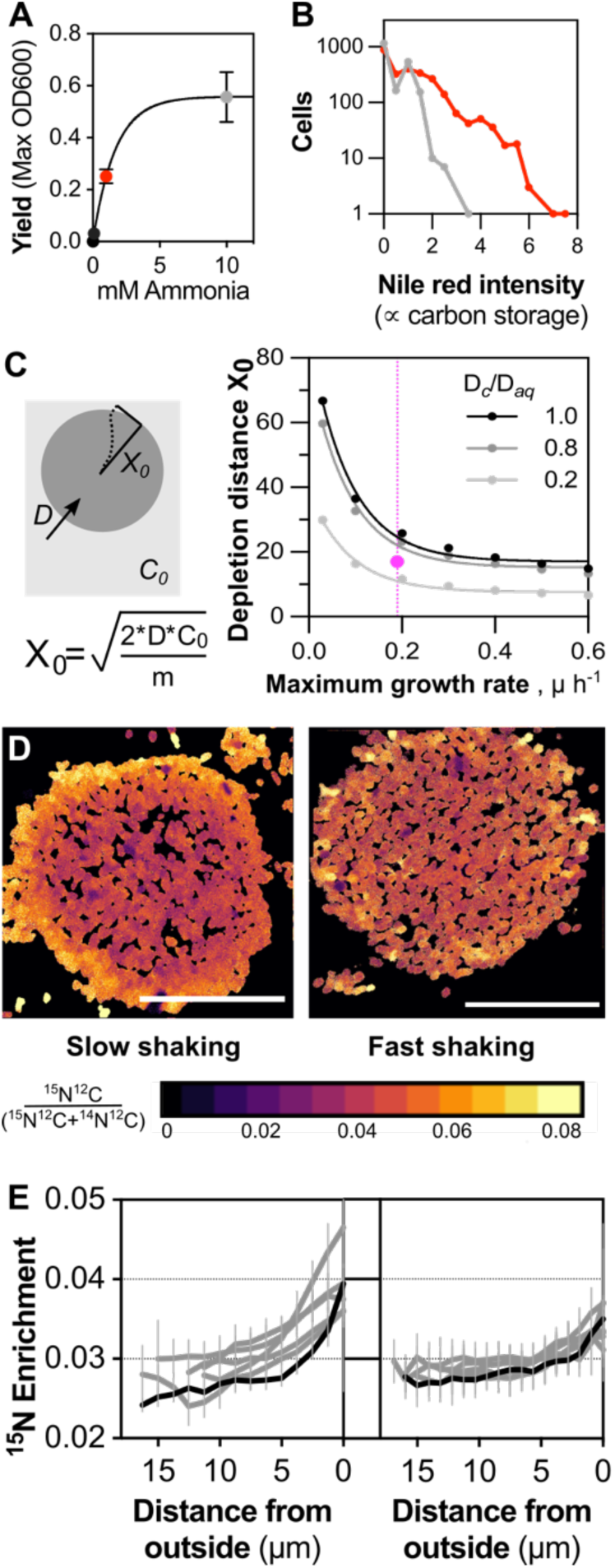
Carbon-storing inner sub-populations are nitrogen limited. **A**) Yield of 12B01 grown on alginate oligosaccharides and varying amounts of ammonium provided as the sole source of nitrogen. Grey dot indicates ammonia-replete growth, red dot indicates ammonia-limited growth. Error bars: standard deviation, n=3. **B**) PHA content under ammonium limitation. Measurements are of individual cells stained by Nile Red staining, made on populations analyzed in panel A. Grey: population grown under ammonia replete conditions; red: population growth under ammonia limited conditions. **C**) Estimates of gradients emerging within clusters due to cellular consumption of ammonium as it diffuses in from the bulk medium. Steady state approximation derived as in (Crank, 1975). *X*_*0*_= distance from outer surface where no ammonium remains, *D*_*aq*_= diffusion coefficient of ammonium in water, D_*c*_/D_*aq*_= ratiometric difference between diffusion coefficient in water and in the extracellular space of cell clusters estimated from (Stewart, 2003), *C*_*0*_=concentration of ammonium ion in bulk medium, *m*=per cell consumption of ammonium, µ= cellular growth rate. Pink dot indicates the 17 µm radius size threshold from Figure 2E where cells inside clusters begin to express carbon storage granules. Vertical pink dashed line indicates the growth rate of clusters with slow shaking (pink). Cellular consumption was estimated from the yield of 12B01 grown on alginate oligomers under ammonium limitation (Figure 1A), and from measurements of cell size in clusters. Length scale estimates were calculated for cellular growth rates ranging from the maximum rate of exponential growth on alginate oligomers (0.6 h^−1^), to the maximum rate of 12B01 cluster growth (0.2 h^−1^, Figure 1D). **D**) ^15^N-ammonium assimilation within stage ii clusters formed under slow shaking (left), or fast shaking (right) measured by nanoscale secondary ion mass spectrometry. Representative secondary ion images show spatial patterns of ^15^N enrichment in an aggregate (fractional abundance ^15^N), where warmer colors represent higher levels of enrichment. Cellular N assimilation is defined from heavy ammonium uptake measured as fractional abundance (^15^N^12^C /(^14^N^12^C+^15^N^12^C)). Scale bars= 20 µm. **E**) Radial profiles of ^15^N-ammonium enrichment within 12B01 clusters grown under slow (left, n=6) or fast (right, n=7) shaking. Bars indicate standard deviation of pixel intensity. Black traces indicate profiles for images shown in 3D. Samples were incubated with ^15^N-lableled ammonium for 4 h.

To test the idea that cooperation for carbon and competition for nitrogen create nutrient gradients and cue self-organized formation of shell and core structures, we sought to perturb the environmental context in which competition and cooperation take place. As we had already established that the addition of alginate oligomers to the growth environment profoundly changes cellular interactions, we instead sought to shift gradients more subtly. It follows from our prediction of gradients emerging from diffusive supply and local cellular uptake (Figure 3C), that decreasing cellular uptake will flatten gradients of ammonia within clusters. We found that increasing the shaking speed at which we grew 12B01 clusters decreased the cellular growth rate and reasoned that this decrease might also decrease cellular demand for ammonium. Consistent with this expectation, NanoSIMS profiling of clusters formed under fast shaking revealed decreased ^15^N-ammonium assimilation across their radial profile of clusters and no detectable gradient in assimilation (Figure 3E, Figure S5). Notably, clusters formed under fast shaking not only grew slower, but they also failed to differentiate into motile ‘core’ and immobile ‘shell’ sub-populations. Instead, only a much smaller sub-population of cells developed motility. Thus, shifting cellular physiology by changing environmental context perturbed both gradient formation and the emergence of complex self-organized structures during growth of 12B01 on alginate.

What does the ability to phenotypically differentiate in response to nitrogen limitation contribute to 12B01 clusters? Previous work has suggested that the emergence of mixing in the core of clusters can collapse resource gradients(Ebrahimi et al., 2019a). Mixing would thus homogenize allocation of carbon and nitrogen among cells in the core sub-population. But this explanation does not account for the accumulation of PHA. To determine whether the ability to store carbon enhanced the ability of 12B01 cells to form new clusters on alginate, we derived populations with high or low proportions of PHA-rich cells (Figure 3B) and compared the growth of these two populations on alginate polysaccharide (Figure 4A). The high PHA population resumed growth on alginate more readily than the population with fewer PHA-rich cells. Moreover, high PHA populations grew at a faster rate than low PHA populations. Together, the faster resumption of growth on alginate and faster population-level growth rate suggest that carbon storage accelerates reproductive cycles during growth of 12B01 on alginate. Thus, phenotypic differentiation of sub-populations may balance cellular conflict and cooperation within aggregates through mixing and may enhance reproduction of clonal lineages by promoting carbon storage.

**Figure 4.**
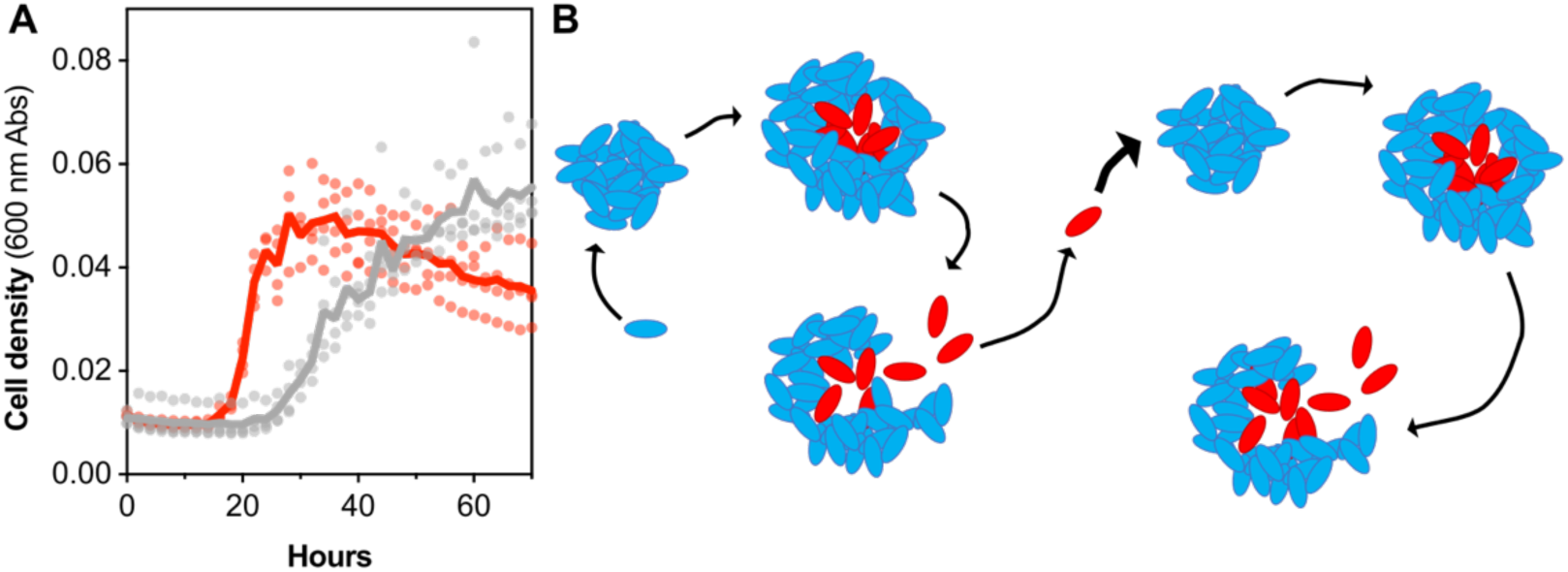
Carbon storage enhances propagation of 12B01 on alginate polysaccharide. A) Growth on alginate polymer by 12B01 populations with high or low proportions of PHA-rich cells. Populations were derived by growing 12B01 on alginate oligosaccharides with 1 mM or 10 mM ammonium chloride as the sole nitrogen source. The two populations were then used to inoculate polymeric alginate cultures containing 10 mM ammonium. Blue, high PHA population formed under ammonia-limited growth on alginate oligosaccharides; red, low PHA population formed under ammonia-replete growth on alginate oligosaccharides. Proportion of PHA in each population quantified in Figure 3B. All replicates are shown, solid line represents mean of replicates. B) Proposed ‘reproductive cycles’ supporting cooperative growth of 12B01 growth on alginate. Cells form clonal clusters. Local density promotes growth on alginate polysaccharide in a carbon-limited environment. As clusters grow, they phenotypically differentiate into ‘shell’ (grey) and ‘core’(blue) sub-populations in response to resource gradients. Clusters rupture, releasing carbon-storing ‘core’ sub-population. Cells carrying high levels of PHA exit stage i faster and initiate a new cycle of self-organization.

## DISCUSSION

Phenotypic heterogeneity is a property of most clonal populations of bacteria (Ackermann, 2015; Dar et al., 2021; Nadell et al., 2016). Where this heterogeneity allows cells to perform specific and synergistic tasks, it can give rise to a phenomenon called division of labor. We propose that the phenotypic differentiation of 12B01 cells during cooperative growth on alginate falls into the framework of a division of labor. We show that gradients of resource limitation arising from cellular conflict and cooperation cue the expression cell-surface adhesins and the accumulation of carbon-storage granules in distinct sub-populations within clusters (Figure 4B). These sub-populations form a structural shell, and a motile, carbon-storing inner core. The ability of the inner sub-population to store carbon enhances propagation of clonal lineages upon dispersal (Figure 4B). It is well known that bacteria integrate environmental cues and gradients into the regulation of biofilm formation (Gerstel and Römling, 2001; Glick et al., 2010; Zacharia et al., 2021). However, it remains challenging to demonstrate that heterogenous cell sub-populations interact synergistically; outside of elegant experiments that have engineered such interactions in the lab (Dragoš et al., 2018), multicellular behaviors (Geerlings et al., 2020; van Gestel et al., 2015) and infection (Ackermann et al., 2008), few natural examples of bacterial division of labor are known. Our work demonstrates that this barrier in understanding can be overcome by studying bacteria in micro-scale ecological contexts that favor growth of collectives over individuals; such environments are typically resource limited, in contrast to the nutrient rich growth environments favored by laboratory studies.

The formation and rupture of the multicellular structures described in this work are reminiscent of the formation and dispersal of surface-attached biofilms (Hunt et al., 2004; Kaplan et al., 2003; Mai-Prochnow et al., 2004; Purevdorj-Gage et al., 2005; Rumbaugh and Sauer, 2020; Sauer et al., 2002; Stewart et al., 2007). This similarity extends beyond the formation of convergent multicellular structures: our work demonstrates that type IV pili are involved in the formation of 12B01 clonal groups, like the role that these dynamic cellular adhesins play in microcolony formation. Interestingly, many type IV pilins have been shown to transfer cargo, such as DNA or bacteriophage (McCallum et al., 2019). Further work will be needed to determine if the Tad/flp type IV pilus expressed by 12B01 shell cells performs cellular functions other than adhesion. In addition, the emergence of a motile core in 12B01 clonal groups recalls a well-known dispersal phenotype of proteobacterial biofilms sometimes called hollowing (Rumbaugh and Sauer, 2020). We show that hollowing emerges in 12B01 clonal groups in the context of a resource imbalance, which promotes formation of carbon storage granules. Although this physiological adaptation is likely to provide an ecological benefit only if cells disperse into carbon-limited environments, hollowing may more generally promote the re-allocation of cellular resources towards a sub-population of cells that will disperse as propagules. More work will be needed to determine whether phenotypic differentiation is a conserved strategy of resource allocation in natural bacterial populations.

The ability to form large, self-organized clusters and undergo reproductive cycles with coordinated bursts of dispersal may shape the ecological dynamics of 12B01. We do not know whether this strain can form similar structures in coastal seawater, however, the concentrations of alginate and ammonium chloride used in our experimental model fall within what can accumulate in coastal seawater during algal blooms (Martin-Platero et al., 2018). Blooms are also characterized by waves of bacteriophage infection. Biofilms and other bacterial structures have been proposed as barriers to phage infection (McDougald et al., 2011; Rumbaugh and Sauer, 2020; Vidakovic et al., 2018). Thus, the highly structured clusters formed by 12B01 may provide a barrier to bacteriophage infection, particularly of the cells in the interior of clusters. Blooms are also accompanied by increases in bacterivores such as protists. The grazing efficiency of protists has been shown to depend on the size and structural integrity of their prey, thus, the formation of dense 40 µm groups may enhance resistance to predation (Pernthaler, 2005). Future work will be needed to determine whether, in addition to accelerating cycles or reproduction, formation of phenotypically differentiated multicellular groups also enhances resistance to bacteriophage or protist predators.

The division of labor, emergence of complex self-organized structures, and clonality that characterize 12B01 growth on alginate are attributes shared by simple multicellular organisms (Szathmáry and Smith, 1995). In this context, the ability of 12B01 to grow in resource-limited environments as clonal structures is particularly intriguing: simulations have suggested that microbial decomposition is constrained both by resource availability and by the evolution of cheats (Allison, 2005). We speculate that 12B01’s ability to form clonal, rather than aggregative, structures that phenotypically specialize in response to resource limitation stabilizes this strain’s strong cooperation, much like clonality stabilizes the evolution of complex multicellularity by minimizing cheating and evolutionary conflict (Márquez-Zacarías et al., 2021; Pentz et al., 2020). This parallel suggests that 12B01 may be a useful experimental model in which to study the evolution of multicellular behaviors such as reproductive specialization.

## METHODS

### Strain information and culture conditions

*Vibrio splendidus* strain 12B01(Le Roux et al., 2009) was maintained in Marine Broth (Difco 2216), as a liquid culture, or on medium solidified with 1.5% agarose. Unless otherwise specified, incubation temperature was 25°C. For experimental measurements, precultures were established in a defined minimal medium (Ebrahimi et al., 2019b) containing 10 mM ammonium as sole source nitrogen, and an indicated carbon source, using a previously described protocol (Amarnath et al., 2021). Precultures were grown with 10 mM glucose to an optical density of 0.4-0.6, as measured at 600 nm with a 1 cm path cuvette on a Genesys 20 Spectrophotometer (Thermo Scientific, Waltham, MA), rinsed by pelleting cells at 5000 rcf for 1 min in a tabletop microcentrifuge (Eppendorf, Hamburg, Germany), and resuspended in carbon-free defined minimal medium such that the final was 1.0. Experimental cultures were initiated by inoculating cells to an initial OD of 0.01 in 250 mL borosilicate glass Erelenmeyer flasks containing 70 mL of alginate minimal medium. Low viscosity alginate (Sigma-Aldrich A1112, St. Louis, MO) was added as the medium carbon source at 700 mg/L (0.07% w/v). ‘Fast’ shaking was established by orbital rotation of flasks at setting 4 on an orbital shaker (Model 3500, VWR, Radnor, PA), which corresponds to ∼200 rpm, ‘Slow’ shaking was established at a setting of 2, corresponding to ∼100 rpm. To pre-digest alginate polysaccharide, 0.01 µg/mL commercial alginate lyase (Sigma-Aldrich A1603, 0.75” rotational orbit) was added to culture medium and incubated overnight with stirring at 37°C. Alginate lyase was removed from culture medium prior to assay by filtering the medium through 3 kDa Amicon Ultra molecular weight cutoff filters (MilliPore Sigma, Burlington, MA). To maintain plasmids in 12B01,12.5 µg/mL chloramphenicol was added to the medium.

### Constructs and validation

Multicopy plasmids expressing eGFP or mKate2 from the synthetic Taq promoter were constructed in previous work (Pollak et al., 2020). These constructs encoded P_*tac*_-eGFP/mKate, the chloramphenicol resistance gene *cat* and the segment of plasmid pVSV208 released by restriction digest by SpeI and SphI, which contains an origin of transfer for RP4 conjugation (Dunn et al., 2006), as well as an origin of replication from a *Vibrio fischeri* plasmid that is stably maintained in several species of *Vibrio* (Le Roux et al., 2011).

To construct a transcriptional reporter for the *tad/flp* locus, we mapped mRNA-Seq reads onto the region of the 12B01 chromosome encoding the 17 open reading frames of the locus, revealing coverage above background levels of read mapping across the cluster. We observed elevated mapping of genes encoding the pilin and inner-membrane assembly complex (Figure S2A). Tad/Flp are a family of type IV pili with broad distribution across bacteria and archaea (Giltner et al., 2012). In the genus *Vibrio*, the genetics and biochemistry of Tad/Flp pili are best characterized in *V. vulnificus* (Ellison et al., 2019; Pu and Rowe-Magnus, 2018). 12B01 encodes a single *tad/flp* locus, which contains homologs of Flp pilin, the inner membrane assembly complex Tad, and the outer membrane motor Rcp all encoded on the minus strand. Accordingly, we mapped the ribosome binding site associated with the putative pilin ORF V12B01_22511 and used 142 bp segment of DNA falling upstream of this site to promote transcription in our reporter construct. The reporter was engineered to express the red fluorescent protein mKate2 from the *tad/flp* promoter by cloning this segment of DNA into plasmid pLL103 through digest by KpnI and SphI restriction enzymes (NEB, Ipswich, MA) to generate pJS2020.1.

The activity of the promoter was confirmed by measuring mKate fluorescence as a function of growth on a Tecan Spark plate reader (Tecan). We performed titration experiments to ensure that the amount of chloramphenicol used to maintain positive selection (12.5 µg/mL) was sufficient to prevent plasmid loss and did not alter growth kinetics or self-organization relative to a no-antibiotic control (Figure S2B). For these experiments, strains were grown in 10 mM glucose, 10 mM ammonium minimal medium with shaking at 25°C.

### Experimental manipulation of PHA expression in 12B01 populations

Nitrogen limitation was used as a cue to induce the expression of carbon storage granules in 12B01. Cells were cultured as described above, except that precultures were established in 0.07% alginate digest minimal medium containing 1 mM or 10 mM ammonium chloride. The eGFP-expressing version of 12B01 (carrying plasmid pLL104) was cultivated under these conditions. The fraction of cell populations expressing carbonosomes was quantified by Nile Red staining.

### Light microscopy-staining conditions

Self-organization of 12B01 was visualized using a ImageXpress high content microscope equipped with Metamorph Software (Molecular devices, San Jose, CA), operating in widefield mode. Images were acquired in widefield mode at 40x with a Ph2 ELWD objective (0.6 NA, Nikon), and filter sets: Ex 482/35 nm, Em: 536/40 nm, dichroic 506 nm to detect eGFP; Ex: 562/40 nm, Em:624/40 nm, dichroic 593 nm to detect mKate, Nile Red, and Priopidium Iodide (Invitrogen); Ex:377/50 nm Em: 447/60 nm dichroic 409 nm to detect TOTO-3 (Invitrogen). The excitation light source was LED lines from a Lumencore light engine set to 100% of their maximum intensity. Images were acquired with a sCMOS detector (Andor Zyla, Andor, Belfast, North Ireland). Unless otherwise noted, raw images were the average of 8 frames, and were collected with exposure times of 100 ms.

Higher resolution images of cells in 12B01 structures were acquired using a Yokogawa CSU-22 spinning disk confocal light path, installed on a Ziess AxioVert 200M inverted microscope with DIC optics (Ziess, Jena, Germany). Images were acquired using Metamorph software, with 63x or 100x 1.4 NA Plan Apochromat oil objectives and illumination from a 488 nm 150 mM OPSL excitation laser with a 525/50 nm bandpass emission filter and 488 nm dichroic. Unless otherwise noted, images were collected with exposure times of 100 ms, and processed with ImageJ, (Version 2.1.0/1.53h).

To visualize DNA, samples of live culture were stained with 5 µM of the DNA-binding dye SYTO9 (Invitrogen), which emits green fluorescence when bound to DNA. Extracellular DNA was stained as described previously (Jemielita et al., 2018) except that 1 µM TOTO-3 iodide was used in place of TOTO-1 iodide (fluorescent molecules are both cell-impermeant DNA stains, but with different excitation/emission properties). Dead cells were detected by staining with 20 µM Propidium iodide. PHA accumulation within cells was stained with 0.5 µg/mL lipophilic fluorescent dye Nile Red (Spiekermann et al., 1999). Dye was resuspended in DMSO at 1 mg/mL, and a fresh 10 µg/mL working solution was created for each experiment to enhance solubility in experimental medium.

### Analysis of light microscopy images

Custom analysis scripts were written to segment 12B01 aggregates to measure aggregate growth, cell composition, expression of transcriptional reporters, and accumulation of PHA. Scripts and raw data are available on the Github repository for this project (github.com/jaschwartzman/12B01.git).

Aggregate growth (plotted in Figure 1D) was measured by segmenting images of aggregates obtained by brightfield microscopy. An intensity-based threshold was used to define the 2-dimensional area of aggregates, and this area was quantified. To convert from segmented 2D area to an estimate of cells per aggregate, cells were assumed to be cylinders of radius 0.5 µm and length 3 µm (volume 3 µm^3^), aggregates were assumed to be spherical with no space between cells. A radius was derived from the 2D area and used to calculate aggregate volume then convert to number of cells per aggregate.

Composition of aggregates (plotted in Figure 1F) was measured by segmenting images of eGFP and mKate expressing cells. A brightfield image was used to create a mask that defined aggregates, and the intensity of eGFP or mKate expression was quantified within each area.

The mean radial intensity profile of PHA accumulation (Figure 2E) was quantified measuring mean intensity of Nile Red fluorescence, or cell-associated eGFP signal within concentric bands spaced from the center of aggregates to the periphery. Accumulation of PHA was defined as an increase in the ratio of Nile Red fluorescence to GFP fluorescence.

### RNA isolation and RNA-Seq

Cultures were grown an indicated stage of self-organization, as defined by microscopy. 30 mL RNA Protect Bacterial Reagent (Qiagen, Hilden, Germany) was added to 15 mL cultures and incubated at 4 °C overnight. Cells were collected by centrifugation at 5000 rcf and 4°C in a Sorvall Legend XIR centrifuge (Thermo Fisher, Waltham, MA), equipped with a Fiberlite F15-6×100g rotor for 30 min. Supernatant was decanted, leaving pelleted cell material which was frozen at −20°C. A Qiagen RNeasy kit was used to isolate total RNA, following manufacturer’s protocol, except for the following modifications: cells were resuspended in 15 mg/mL lysozyme in TE buffer, and incubated for room temperature for 30 min, prior to addition of buffer RLT. Samples were lysed by mechanical disruption using lysing matrix B (MPBio, Santa Ana, CA). Samples in lysing matrix shaken in a homogenizer (MPBio) for 10X 30 second intervals, taking care to keep samples from heating up. Samples were treated with DNase digestion using a Turbo DNA free DNAse kit (Ambion, Austin, TX), following manufacturer’s instructions. The integrity and purity of total RNA was assessed running a sample on an Agilent 4200 Tapestation in a HS RNA screen tape (Agilent, Santa Clara, CA). Depletion of rRNA was accomplished using the Ribominus Yeast and Bacteria transcriptome isolation kit (Invitrogen, Waltham, MA) and included ethanol precipitation with glycogen to concentrate remaining mRNA. Ethanol precipitated samples were resuspended in FPF sample buffer from the TruSeq stranded mRNA-Seq kit reagents (Illumina, San Diego, CA). mRNA abundance was quantified using a Quant-iT RNA Assay kit (Invitrogen) on a Tecan Spark plate reader equipped with a monochrometer, following manufacturer’s instructions (Tecan, Männendorf, Switzeland). The quality of the mRNA was further assessed using an Agilent HS tape. We analyzed resulting samples on a Bioanalyzer to confirm mRNA depletion, and concentrated samples using ethanol precipitation with glycogen. Samples were screened for quality by requiring an RNA integrity number (RIN) or more than 8. mRNA remaining from rRNA depletion of 2-10 µg total RNA was used as template to create sequencing libraries. Sequencing libraries were prepared using an Illumina TruSeq Stranded mRNA library preparation kit (Illumina). Libraries were assessed for quality using screen tapes and fluorescence-based plate reader assays prior to pooling. Libraries were sequenced at the Whitehead sequencing core (Whitehead Institute, MIT) on an Illumina HiSeq in 60×60 paired-end mode.

### RNASeq data analysis

Paired-end Illumina reads were trimmed using Trimmomatic (v0.30) to remove sequencing adapters and low quality reads (phred >30)(Bolger et al., 2014). The remaining paired reads were checked for quality using FastQC (version 0.11.9) (Brown et al., 2017) and mapped to the predicted coding regions of the *V. splendidus* 12B01 genome (assembly ASM15276v1) using Bowtie2 (v 2.3.4.1)(Langmead and Salzberg, 2012), outputting SAM files. SAM files were sorted by position using SAMTools (V 1.9) (Danecek et al., 2021), and mapping was parsed using HTSeq (Anders et al., 2015) to obtain count tables. Differential gene expression was assessed from count tables DeSeq2 (Love et al., 2014) (run in R Studio Version 1.2.5042). Briefly, tables were processed using a variance stabilizing transformation to assess clustering of replicates, after which the statistical significance of log fold change between pairs of samples was assessed using a Wald test. P values were adjusted using the Benjamin-Hochberg method, to account for multiple tests in followed by a Wald test to determine differential gene expression in log_2_ normalized abundance. Normalized transcript abundance was calculated from count tables with custom analysis scripts to implement the transcripts per kilobase million (TPM) method (github.com/jaschwartzman/12B01.git for code). Predicted 12B01 coding regions were further annotated using Eggnogg mapper 2.1 to make functional inferences (Cantalapiedra et al., 2021; Huerta-Cepas et al., 2019).

### Stable isotope amendment experiments and nanoSIMS analysis

To trace the assimilation of ammonium into cellular biomass, 10 mM of ^15^N-labeled ammonium (99% 15N, Cambridge Isotope Laboratories, Tewksbury, MA, USA) was added to cultures of 12B01 growing on 0.07% alginate. The cultures initially contained 10 mM of unlabeled ammonium as the sole nitrogen source which was depleted upon labelled ammonium amendment. Sample collection and processing was conducted following previously established protocols (McGlynn et al., 2018), with minor modifications: Samples were collected after a 4 h incubation with the heavy-labeled ammonium. Incubation time includes a 1 h step in which aggregates settle out. After removal of supernatant, an equal volume of 4% paraformaldehyde mPBS (50 mM sodium phosphate pH 7.4, 0.4 M NaCl) was added to the concentrated aggregate mixture for 45 min. 30 min of this time was static, to allow aggregates to settle out. Three 1-hour rinses with 1xmPBS were performed, in which aggregates were incubated with rotation, then allowed to settle for 30 min. Following the final wash step, concentrated aggregates were combined with 1 part molten 9% noble agar in phosphate buffered saline. The agar-aggregate mixture was allowed to solidify in a 1.5 mL centrifuge tube. Samples were stored at 4°C prior to processing. Immobilized samples were dehydrated in ethanol and stored at −20 °C prior to embedding in Technovit 8100 resin (Heraeus Kulzer GmbH) and continuing with the manufacturer’s standard protocol. Embedded samples were sectioned into 1 µm thick slices with a glass knife using a Lecia microtome and mounted onto poly-l-lysine-coated glass slides (Tekdon, Myakka City, FL), sputter coated with 40nm of gold using a Cressington sputter coater. Samples were analyzed on a CAMECA nanoSIMS 50L in the Center for Microanalysis at Caltech Data acquisition began with a pre-sputtering step using a 90-pA primary Cs+ ion beam with aperture diaphragm setting of D1 = 1 until the ^14^N^12^C− ion counts stabilized. ^14^N^12^C− and ^15^N^12^C− ions were imaged along with secondary electron images. Data were collected using a 0.5-pA primary Cs+ beam with aperture diaphragm setting of D1 = 3 and the entrance slit set at (ES) = 2. Dwell times ranged from 1-10ms/pixel with acquisition area between 10 and 40 microns.

Raw images were processed with Look@nanosims (Polerecky et al., 2012) analysis scripts, to convert secondary ion count data output from the CAMECA into a format readable by MATLAB. Converted ^14^N^12^C and ^15^N^12^C secondary ion count data were further processed with custom analysis scripts to calculate the ^15^N fractional abundance and to calculate the average radial ^15^N fractional abundance. Where aggregates were imaged with multiple fields of view, images were stitched together in ImageJ (Version 2.1.0/1.53h) using the Grid collection/stitching plugin (internal version 1.2) (Preibisch et al., 2009). Image analysis code and raw data are available in the Github repository for this manuscript (github.com/jaschwartzman/12B01.git).

## Supporting information

Move S1

## ACKNOWLEDGEMENTS

We thank the Polz lab for kindly providing *Vibrio splendidus* 12B01. Glen D’Souza, Jan Hendrick Hehemann and Andreas Sichert provided critical advice about working with alginate. Steven Biller, Allison Coe, and members of the Chisholm lab for advice regarding RNA extraction and RNASeq data analysis. Yunbin Guan for assistance with nanoSIMS operations. We thank Terrence Hwa, Kapil Amernath, Ghita Ghessous and members of the Hwa lab, Martin Ackermann, and Ben Roller for their advice and discussions. A.E. acknowledges funding from Swiss National Science Foundation: Grants P2EZP2 175128 and P400PB_186751. Y.S. was funded through the Japan Society for the Promotion of Science KAKENHI (Grant Number 20H02291). This work was supported by Simons Foundation: Principles of Microbial Ecosystems (PriME) award number 542395. O.X.C and J.S acknowledge support from the Kavli Institute of Theoretical Physics National Science Foundation Grant No. NSF PHY-1748958, NIH Grant No. R25GM067110, the Gordon and Betty Moore Foundation Grant No. 2919.02, National Science Foundation under Grant No. NSF PHY-1748958.

## AUTHOR CONTRIBUTIONS

J.S., A.E, and O.X. designed the study, with help from V.O. and G.C, about the design of stable isotope experiments. J.S, A.E., Y.S. and G.C. performed experiments. All authors contributed to data analysis and writing the paper.

## REFERENCES

Ackermann M. 2015. A functional perspective on phenotypic heterogeneity in microorganisms. Nat Rev Microbiol 13:497–508.

Ackermann M, Stecher B, Freed NE, Songhet P, Hardt W-D, Doebeli M. 2008. Self-destructive cooperation mediated by phenotypic noise. Nature 454:987–990.

Allison SD. 2005. Cheaters, diffusion and nutrients constrain decomposition by microbial enzymes in spatially structured environments. Ecol Lett 8:626–635.

Amarnath K, Narla AV, Pontrelli S, Dong J, Caglar T, Taylor BR, Schwartzman J, Sauer U, Cordero OX, Hwa T. 2021. Stress-induced cross-feeding of internal metabolites provides a dynamic mechanism of microbial cooperation. bioRxiv. doi:10.1101/2021.06.24.449802

Anders S, Pyl PT, Huber W. 2015. HTSeq--a Python framework to work with high-throughput sequencing data. Bioinformatics 31:166–169.

Arciola CR, Campoccia D, Montanaro L. 2018. Implant infections: adhesion, biofilm formation and immune evasion. Nat Rev Microbiol 16:397–409.

Badur AH, Jagtap SS, Yalamanchili G, Lee J-K, Zhao H, Rao CV. 2015. Alginate lyases from alginate-degrading Vibrio splendidus 12B01 are endolytic. Appl Environ Microbiol 81:1865–1873.

Badur AH, Plutz MJ, Yalamanchili G, Jagtap SS, Schweder T, Unfried F, Markert S, Polz MF, Hehemann J-H, Rao CV. 2017. Exploiting fine-scale genetic and physiological variation of closely related microbes to reveal unknown enzyme functions. J Biol Chem 292:13056– 13067.

Bolger AM, Lohse M, Usadel B. 2014. Trimmomatic: a flexible trimmer for Illumina sequence data. Bioinformatics 30:2114–2120.

Brown J, Pirrung M, McCue LA. 2017. FQC Dashboard: integrates FastQC results into a web-based, interactive, and extensible FASTQ quality control tool. Bioinformatics 33:3137– 3139.

Burrows LL. 2012. Pseudomonas aeruginosa twitching motility: type IV pili in action. Annu Rev Microbiol 66:493–520.

Cantalapiedra CP, Hernández-Plaza A, Letunic I, Bork P, Huerta-Cepas J. 2021. eggNOG-mapper v2: Functional Annotation, Orthology Assignments, and Domain Prediction at the Metagenomic Scale. Mol Biol Evol. doi:10.1093/molbev/msab293

Claessen D, Rozen DE, Kuipers OP, Søgaard-Andersen L, van Wezel GP. 2014. Bacterial solutions to multicellularity: a tale of biofilms, filaments and fruiting bodies. Nat Rev Microbiol 12:115–124.

Crank J. 1975. xThe Mathematics of Diffusion. Oxford University Press.

Danecek P, Bonfield JK, Liddle J, Marshall J, Ohan V, Pollard MO, Whitwham A, Keane T, McCarthy SA, Davies RM, Li H. 2021. Twelve years of SAMtools and BCFtools. Gigascience 10. doi:10.1093/gigascience/giab008

Dar D, Dar N, Cai L, Newman DK. 2021. Spatial transcriptomics of planktonic and sessile bacterial populations at single-cell resolution. Science 373. doi:10.1126/science.abi4882

Dragoš A, Kiesewalter H, Martin M, Hsu C-Y, Hartmann R, Wechsler T, Eriksen C, Brix S, Drescher K, Stanley-Wall N, Kümmerli R, Kovács ÁT. 2018. Division of Labor during Biofilm Matrix Production. Curr Biol 28:1903-1913.e5.

D’Souza GG, Povolo VR, Keegstra JM, Stocker R, Ackermann M. 2021. Nutrient complexity triggers transitions between solitary and colonial growth in bacterial populations. ISME J. doi:10.1038/s41396-021-00953-7

Dunn AK, Millikan DS, Adin DM, Bose JL, Stabb EV. 2006. New rfp-and pES213-derived tools for analyzing symbiotic Vibrio fischeri reveal patterns of infection and lux expression in situ. Appl Environ Microbiol 72:802–810.

Ebrahimi A, Schwartzman J, Cordero OX. 2019a. Multicellular behaviour enables cooperation in microbial cell aggregates. Philos Trans R Soc Lond B Biol Sci 374:20190077.

Ebrahimi A, Schwartzman J, Cordero OX. 2019b. Cooperation and spatial self-organization determine rate and efficiency of particulate organic matter degradation in marine bacteria. Proc Natl Acad Sci U S A 116:23309–23316.

Ellison CK, Kan J, Chlebek JL, Hummels KR, Panis G, Viollier PH, Biais N, Dalia AB, Brun YV. 2019. A bifunctional ATPase drives tad pilus extension and retraction. Sci Adv 5:eaay2591.

Geerlings NMJ, Karman C, Trashin S, As KS, Kienhuis MVM, Hidalgo-Martinez S, Vasquez-Cardenas D, Boschker HTS, De Wael K, Middelburg JJ, Polerecky L, Meysman FJR. 2020. Division of labor and growth during electrical cooperation in multicellular cable bacteria. Proc Natl Acad Sci U S A 117:5478–5485.

Gerstel U, Römling U. 2001. Oxygen tension and nutrient starvation are major signals that regulate agfD promoter activity and expression of the multicellular morphotype in Salmonella typhimurium. Environ Microbiol 3:638–648.

Giltner CL, Nguyen Y, Burrows LL. 2012. Type IV pilin proteins: versatile molecular modules. Microbiol Mol Biol Rev 76:740–772.

Glick R, Gilmour C, Tremblay J, Satanower S, Avidan O, Déziel E, Greenberg EP, Poole K, Banin E. 2010. Increase in rhamnolipid synthesis under iron-limiting conditions influences surface motility and biofilm formation in Pseudomonas aeruginosa. J Bacteriol 192:2973– 2980.

Greening C, Lithgow T. 2020. Formation and function of bacterial organelles. Nat Rev Microbiol. doi:10.1038/s41579-020-0413-0

Grosberg RK, Strathmann RR. 2007. The Evolution of Multicellularity: A Minor Major Transition? Annu Rev Ecol Evol Syst 38:621–654.

Hehemann J-H, Arevalo P, Datta MS, Yu X, Corzett CH, Henschel A, Preheim SP, Timberlake S, Alm EJ, Polz MF. 2016. Adaptive radiation by waves of gene transfer leads to fine-scale resource partitioning in marine microbes. Nat Commun 7:12860.

Høiby N, Bjarnsholt T, Givskov M, Molin S, Ciofu O. 2010. Antibiotic resistance of bacterial biofilms. Int J Antimicrob Agents 35:322–332.

Huerta-Cepas J, Szklarczyk D, Heller D, Hernández-Plaza A, Forslund SK, Cook H, Mende DR, Letunic I, Rattei T, Jensen LJ, von Mering C, Bork P. 2019. eggNOG 5.0: a hierarchical, functionally and phylogenetically annotated orthology resource based on 5090 organisms and 2502 viruses. Nucleic Acids Res 47:D309–D314.

Hunt SM, Werner EM, Huang B, Hamilton MA, Stewart PS. 2004. Hypothesis for the role of nutrient starvation in biofilm detachment. Appl Environ Microbiol 70:7418–7425.

Jagtap SS, Hehemann J-H, Polz MF, Lee J-K, Zhao H. 2014. Comparative biochemical characterization of three exolytic oligoalginate lyases from Vibrio splendidus reveals complementary substrate scope, temperature, and pH adaptations. Appl Environ Microbiol 80:4207–4214.

Jemielita M, Wingreen NS, Bassler BL. 2018. Quorum sensing controls Vibrio cholerae multicellular aggregate formation. Elife 7. doi:10.7554/eLife.42057

Jendrossek D. 2009. Polyhydroxyalkanoate granules are complex subcellular organelles (carbonosomes). J Bacteriol 191:3195–3202.

Kaplan JB. 2010. Biofilm dispersal: mechanisms, clinical implications, and potential therapeutic uses. J Dent Res 89:205–218.

Kaplan JB, Meyenhofer MF, Fine DH. 2003. Biofilm growth and detachment of Actinobacillus actinomycetemcomitans. J Bacteriol 185:1399–1404.

Kasimoglu E, Park SJ, Malek J, Tseng CP, Gunsalus RP. 1996. Transcriptional regulation of the proton-translocating ATPase (atpIBEFHAGDC) operon of Escherichia coli: control by cell growth rate. J Bacteriol 178:5563–5567.

Koschwanez JH, Foster KR, Murray AW. 2013. Improved use of a public good selects for the evolution of undifferentiated multicellularity. Elife 2:e00367.

Koschwanez JH, Foster KR, Murray AW. 2011. Sucrose utilization in budding yeast as a model for the origin of undifferentiated multicellularity. PLoS Biol 9:e1001122.

Langmead B, Salzberg SL. 2012. Fast gapped-read alignment with Bowtie 2. Nat Methods 9:357– 359.

Le Roux F, Davis BM, Waldor MK. 2011. Conserved small RNAs govern replication and incompatibility of a diverse new plasmid family from marine bacteria. Nucleic Acids Res 39:1004–1013.

Le Roux F, Zouine M, Chakroun N, Binesse J, Saulnier D, Bouchier C, Zidane N, Ma L, Rusniok C, Lajus A, Buchrieser C, Médigue C, Polz MF, Mazel D. 2009. Genome sequence of Vibrio splendidus: an abundant planctonic marine species with a large genotypic diversity. Environ Microbiol 11:1959–1970.

Love MI, Huber W, Anders S. 2014. Moderated estimation of fold change and dispersion for RNA-seq data with DESeq2. Genome Biol 15:550.

Mai-Prochnow A, Evans F, Dalisay-Saludes D, Stelzer S, Egan S, James S, Webb JS, Kjelleberg S. 2004. Biofilm development and cell death in the marine bacterium Pseudoalteromonas tunicata. Appl Environ Microbiol 70:3232–3238.

Márquez-Zacarías P, Conlin PL, Tong K, Pentz JT, Ratcliff WC. 2021. Why have aggregative multicellular organisms stayed simple? Curr Genet. doi:10.1007/s00294-021-01193-0

Martin-Platero AM, Cleary B, Kauffman K, Preheim SP, McGillicuddy DJ, Alm EJ, Polz MF. 2018. High resolution time series reveals cohesive but short-lived communities in coastal plankton. Nat Commun 9:1–11.

McCallum M, Burrows LL, Howell PL. 2019. The Dynamic Structures of the Type IV Pilus. Microbiol Spectr 7. doi:10.1128/microbiolspec.PSIB-0006-2018

McDougald D, Rice SA, Barraud N, Steinberg PD, Kjelleberg S. 2011. Should we stay or should we go: mechanisms and ecological consequences for biofilm dispersal. Nat Rev Microbiol 10:39–50.

McGlynn SE, Chadwick GL, O’Neill A, Mackey M, Thor A, Deerinck TJ, Ellisman MH, Orphan VJ. 2018. Subgroup Characteristics of Marine Methane-Oxidizing ANME-2 Archaea and Their Syntrophic Partners as Revealed by Integrated Multimodal Analytical Microscopy. Appl Environ Microbiol 84. doi:10.1128/AEM.00399-18

Murray EL, Conway T. 2005. Multiple Regulators Control Expression of the Entner-Doudoroff Aldolase (Eda) of Escherichia coli. J Bacteriol 187:991–1000.

Nadell CD, Drescher K, Foster KR. 2016. Spatial structure, cooperation and competition in biofilms. Nat Rev Microbiol 14:589–600.

Pahel G, Rothstein DM, Magasanik B. 1982. Complex glnA-glnL-glnG operon of Escherichia coli. J Bacteriol 150:202–213.

Pentz JT, Márquez-Zacarías P, Bozdag GO, Burnetti A, Yunker PJ, Libby E, Ratcliff WC. 2020. Ecological Advantages and Evolutionary Limitations of Aggregative Multicellular Development. Curr Biol 30:4155-4164.e6.

Pernthaler J. 2005. Predation on prokaryotes in the water column and its ecological implications. Nat Rev Microbiol 3:537–546.

Polerecky L, Adam B, Milucka J, Musat N, Vagner T, Kuypers MMM. 2012. Look@NanoSIMS--a tool for the analysis of nanoSIMS data in environmental microbiology. Environ Microbiol 14:1009–1023.

Pollak S, Gralka M, Sato Y, Schwartzman J, Lu L, Cordero OX. 2020. Public good exploitation in natural bacterioplankton communities. bioRxiv. doi:10.1101/2020.12.13.422583

Preibisch S, Saalfeld S, Tomancak P. 2009. Globally optimal stitching of tiled 3D microscopic image acquisitions. Bioinformatics 25:1463–1465.

Pu M, Rowe-Magnus DA. 2018. A Tad pilus promotes the establishment and resistance of Vibrio vulnificus biofilms to mechanical clearance. NPJ Biofilms Microbiomes 4:10.

Purevdorj-Gage B, Costerton WJ, Stoodley P. 2005. Phenotypic differentiation and seeding dispersal in non-mucoid and mucoid Pseudomonas aeruginosa biofilms. Microbiology 151:1569–1576.

Ratzke C, Gore J. 2016. Self-organized patchiness facilitates survival in a cooperatively growing Bacillus subtilis population. Nat Microbiol 1:16022.

Rumbaugh KP, Sauer K. 2020. Biofilm dispersion. Nat Rev Microbiol 18:571–586.

Sauer K, Camper AK, Ehrlich GD, Costerton JW, Davies DG. 2002. Pseudomonas aeruginosa displays multiple phenotypes during development as a biofilm. J Bacteriol 184:1140–1154.

Shapiro JA. 1998. Thinking about bacterial populations as multicellular organisms. Annu Rev Microbiol 52:81–104.

Sichert A, Cordero OX. 2021. Polysaccharide-Bacteria Interactions From the Lens of Evolutionary Ecology. Front Microbiol 12:2903.

Spiekermann P, Rehm BH, Kalscheuer R, Baumeister D, Steinbüchel A. 1999. A sensitive, viable-colony staining method using Nile red for direct screening of bacteria that accumulate polyhydroxyalkanoic acids and other lipid storage compounds. Arch Microbiol 171:73–80.

Stewart PS. 2003. Diffusion in biofilms. J Bacteriol 185:1485–1491.

Stewart PS, Rani SA, Gjersing E, Codd SL, Zheng Z, Pitts B. 2007. Observations of cell cluster hollowing in Staphylococcus epidermidis biofilms. Lett Appl Microbiol 44:454–457.

Szathmáry E, Smith JM. 1995. The major evolutionary transitions. Nature 374:227–232.

van Gestel J, Vlamakis H, Kolter R. 2015. Division of Labor in Biofilms: the Ecology of Cell Differentiation. Microbiol Spectr 3:MB-0002-2014.

Vidakovic L, Singh PK, Hartmann R, Nadell CD, Drescher K. 2018. Dynamic biofilm architecture confers individual and collective mechanisms of viral protection. Nat Microbiol 3:26–31.

Vlamakis H, Chai Y, Beauregard P, Losick R, Kolter R. 2013. Sticking together: building a biofilm the Bacillus subtilis way. Nat Rev Microbiol 11:157–168.

Wargacki AJ, Leonard E, Win MN, Regitsky DD, Santos CNS, Kim PB, Cooper SR, Raisner RM, Herman A, Sivitz AB, Lakshmanaswamy A, Kashiyama Y, Baker D, Yoshikuni Y. 2012. An engineered microbial platform for direct biofuel production from brown macroalgae. Science 335:308–313.

Wu SS, Kaiser D. 1995. Genetic and functional evidence that Type IV pili are required for social gliding motility in Myxococcus xanthus. Mol Microbiol 18:547–558.

Zacharia VM, Ra Y, Sue C, Alcala E, Reaso JN, Ruzin SE, Traxler MF. 2021. Genetic Network Architecture and Environmental Cues Drive Spatial Organization of Phenotypic Division of Labor in Streptomyces coelicolor. MBio 12. doi:10.1128/mBio.00794-21

