## Supplemental figures and tables for "Bacterial growth in multicellular aggregates leads to the emergence of complex lifecycles"

File contains:
Figure S1-S5
Table S1

[see additional file]
**Movie S1**. Time series showing motility in the core of stage ii aggregates. Cells constitutively express eGFP from plasmid pLL104. Scale bar= 10μm.

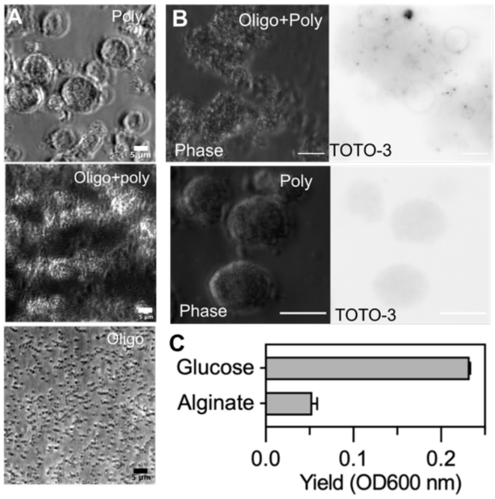


**Figure S1. Properties of 12B01 clonal groups formed during growth on alginate polysaccharide, in comparison to growth on oligosaccharide.** Scale bar =20 μm A) Morphology of 12B01 grown on alginate oligosaccharides (Oligo), a combination of alginate oligosaccharides and low viscosity polysaccharide (Oligo+Poly), or polysaccharide only (Poly). B) Staining of 12B01 during growth on alginate polysaccharide or oligosaccharide with TOTO-3 iodide, a cell-impermeant fluorophore that detects extracellular DNA and dead cells. Left-hand companion images were acquired in phase-contrast and show the area occupied by 12B01. For TOTO-3 staining, darker signal indicates increased staining of DNA. C) Growth of 12B01 cultures on alginate or glucose under anoxic conditions.

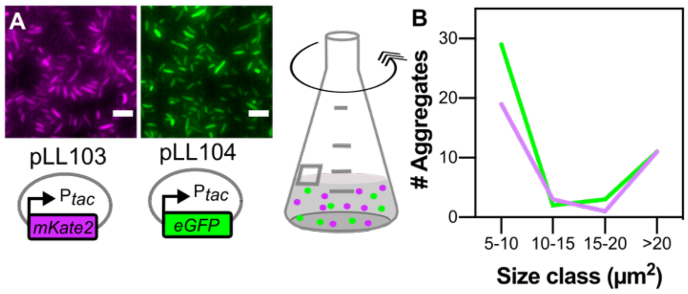


**Figure S2. Growth of labeled 12B01 populations is not affected by expression of fluorescent proteins.** A) Construction of fluorescent-protein expressing 12B01. Plasmids pLL103 and pLL104 express mKate2 of eGFP from the synthetic $P_{tac}$ promoter. The plasmid backbone encoded a chloramphenicol acetyltransferase, and a *Vibrio*-specific origin of replication. Scale bars= 2 μm. Populations of the two plasmid-expressing isogenic strains were mixed to initiate the experiment. B) Distribution of group sizes for each fluorescent marker, measured for clusters collected from a single time-point during stage ii self-organization. Raw dataset analyzed is the same as shown in figure 2F.

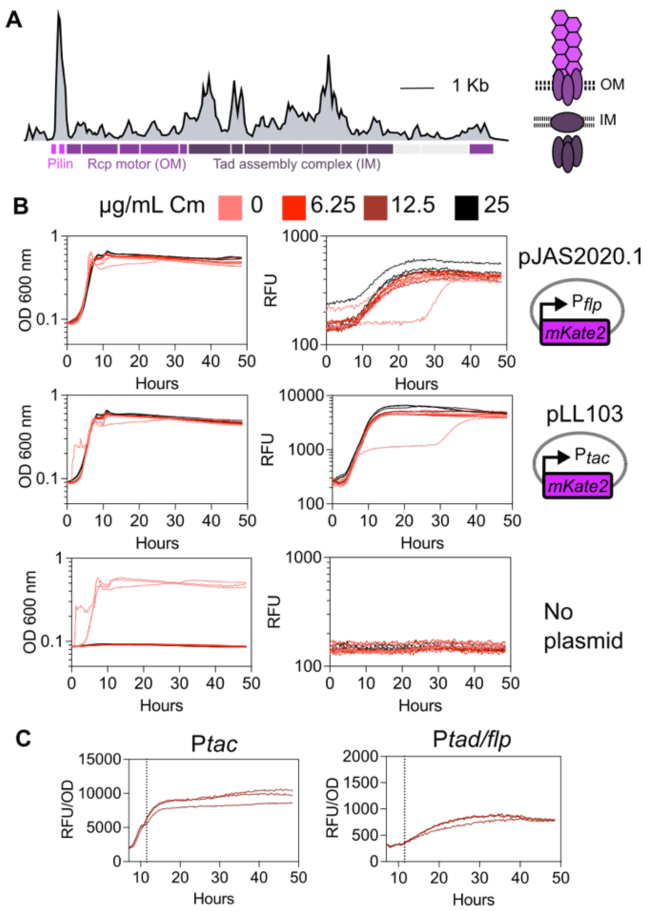


**Figure S3**. **Construction of transcriptional reporter constructs and characterization of growth rate and growth stage dependent gene expression in *Vibrio splendidus* 12B01.** A) Construction of $P_{flp}$::mKate transcriptional reporter. The promoter region for the *tad/flp* locus on the 12B01 chromosome was cloned upstream of fluorescent protein mKate2 to create a transcriptional reporter construct pJAS2020.1. B) Antibiotic selection was sufficient to maintain expression of the multicopy plasmids used in this study. The growth and fluorescence of 12B01 expressing mKate from either pJAS2020.1 or pLL103 was monitored during growth on glucose minimal medium containing different concentrations of chloramphenicol. A no-plasmid control was included to assess intrinsic resistance of 12B01 to chloramphenicol. Three biological replicates are shown. C) Activity of the *tac* or *tad/flp* promoter in 10 mM glucose batch culture. Vertical dashed line indicates maximum OD of culture. Three biological replicates are shown.

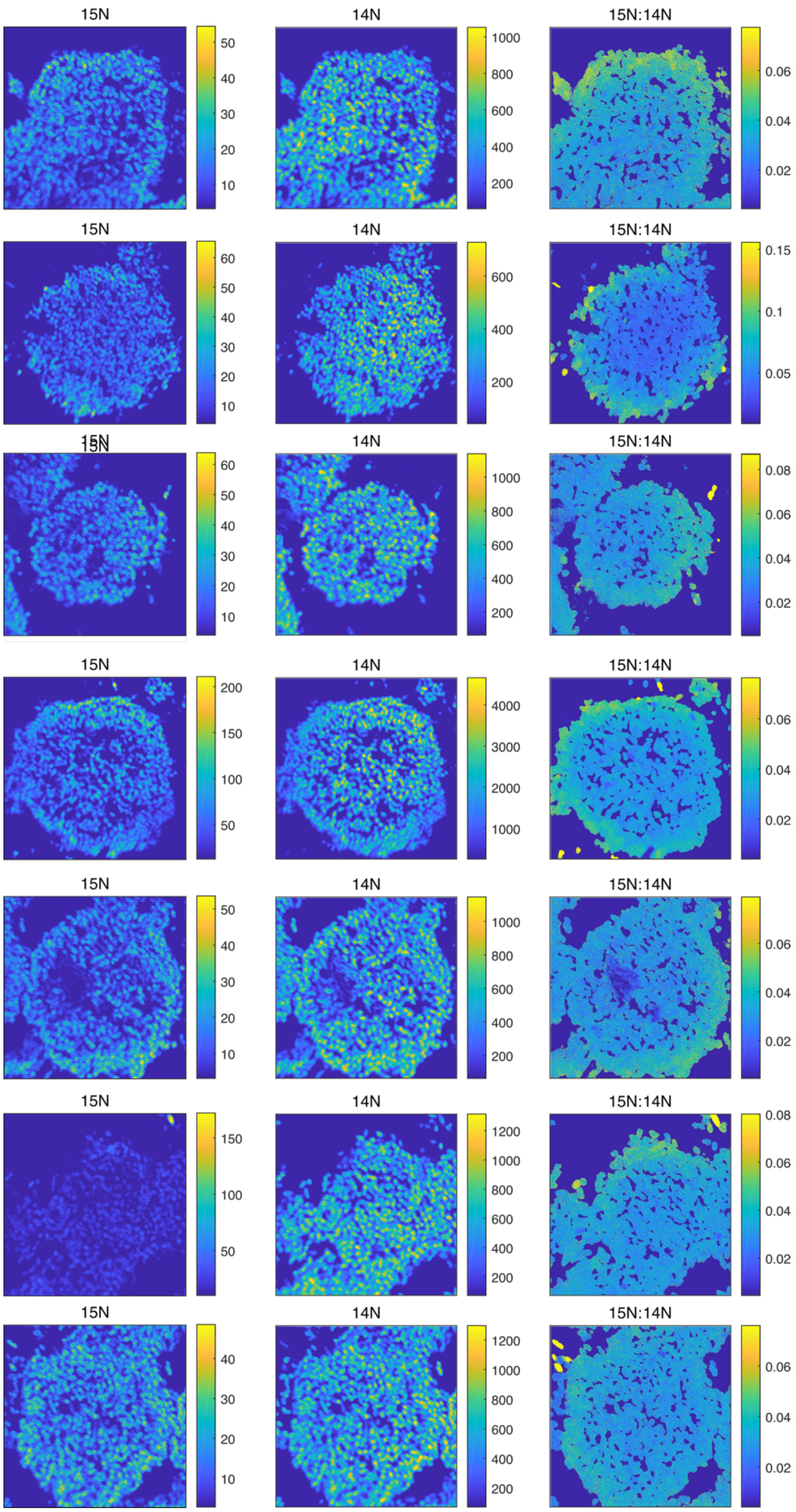


**Figure S4. Quantification of ammonia assimilation by stage ii 12B01 clusters cultivated under slow shaking.** Clusters were incubated for 4 h with $^{15}N$ ammonia. Incorporation of $^{15}N$ and $^{14}N$ into carbon containing biomass was quantified by secondary ion mass spectrometry. Left: $^{15}N^{12}C$ signal, middle, $^{14}N^{12}C$ signal, right enrichment of $^{15}N^{12}C$ ($^{15}N^{12}C/(^{15}N^{12}C+^{14}N^{12}C)$). All measurements taken are shown.

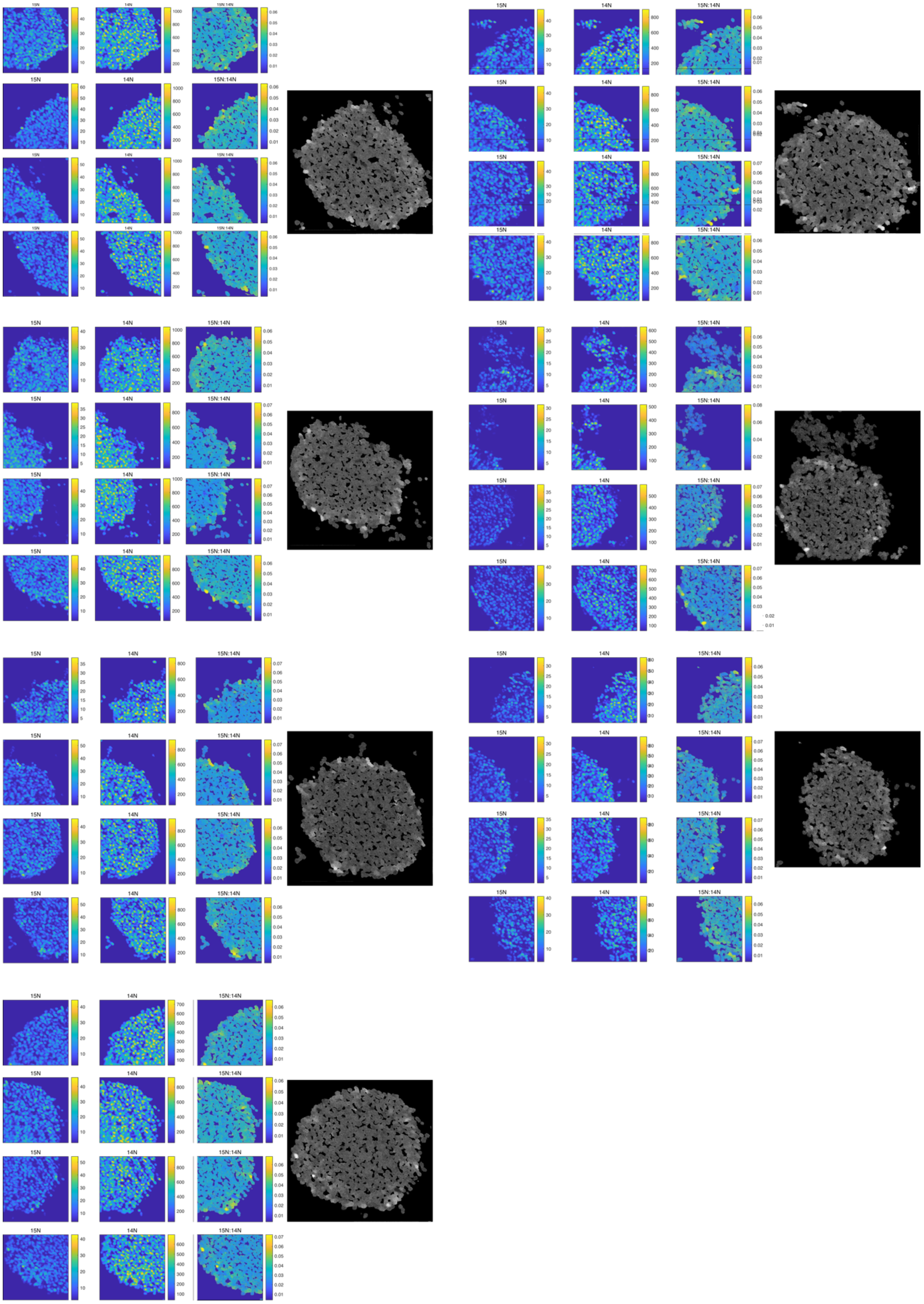
15N
14N
15N:14N

**Figure S5. Quantification of ammonia assimilation by stage ii 12B01 clusters cultivated under faster shaking.** As in Figure S4, clusters in stage ii of development were incubated for 4 h with $^{15}N$ ammonia. Incorporation of $^{15}N$ and $^{14}N$ into carbon containing biomass was quantified by secondary ion mass spectrometry. Left: $^{15}N^{12}C$ signal, middle, $^{14}N^{12}C$ signal, right enrichment of $^{15}N^{12}C$ ($^{15}N^{12}C/(^{15}N^{12}C+^{14}N^{12}C)$). Stitched image is shown to the far right. All measurements taken are shown.

**Table S1. Differential gene expression in 12B01 samples collected during stage ii and iii of morphogenesis on alginate.**

| Samples compared | | | | | | Annotations | | | | | | DeSeq Fold Change Statistics | | | TPM values | | | | | |
|---|---|---|---|---|---|---|---|---|---|---|---|---|---|---|---|---|---|---|---|---|
| Condition 1 | Condition 2 | log2FoldChange (condition 1 vs. 2) | Adjusted P value | Gene_ID | Protein ID | Gene annotation (NCBI) | Biological process (Biocyc Pathway) | KEGG | COG | Description | Gene name | Base Mean | Standard Error | P value | Comparitor 1 rep 1 | Comparitor 1 rep 2 | Comparitor 1 rep 3 | Comparitor 2 rep 1 | Comparitor 2 rep 2 | Comparitor 2 rep 3 |
| stage3_free | stage3_shell | 2.50 | 4.46E-04 | V12B01_00010 | EAP91702.1 | putative phosphoheptose isomerase | Carbohydrate Biosynthesis | K12961 | G | Catalyzes the isomerization of sedoheptulose 7-phosphate in D-glycero-D-manno-heptose 7-phosphate | diaA | 26.3 | 0.69 | 4.49E-05 | 87.4 | 72.3 | NA | 10.8 | 2.6 | 14.5 |
| stage3_free | stage3_shell | 2.15 | 3.60E-04 | V12B01_00035 | EAP91699.1 | Ubiquinol-cytochrome c reductase, cytochrome B | Aerobic Respiration | K00412 | C | Component of the ubiquinol-cytochrome c reductase complex (complex III or cytochrome b-c1 complex), which is a respiratory chain that generates an electrochemical potential coupled to ATP synthesis | petB | 481.5 | 0.58 | 3.48E-05 | 1316.8 | 1196.0 | NA | 170.4 | 68.1 | 267.6 |
| stage2 | stage3_shell | 2.23 | 3.76E-05 | V12B01_00240 | EAP93508.1 | translation initiation inhibitor | Translation | | J | translation initiation inhibitor, yjgF family | - | 48.5 | 0.52 | 3.41E-06 | 217.2 | 337.9 | 408.9 | 82.6 | 17.4 | 32.7 |
| stage3_free | stage3_shell | 3.40 | 7.39E-14 | V12B01_00250 | EAP93510.1 | argininosuccinate lyase | Amino Acid Biosynthesis | K14681 | E | belongs to the lyase 1 family. Argininosuccinate lyase subfamily | argH | 639.6 | 0.43 | 3.58E-16 | 1248.9 | 755.5 | NA | 74.1 | 35.5 | 82.0 |
| stage3_free | stage3_shell | 3.15 | 3.70E-04 | V12B01_00255 | EAP93511.1 | argininosuccinate synthase | Amino Acid Biosynthesis | K01940 | F | Belongs to the argininosuccinate synthase family. Type 1 subfamily | argG | 876.9 | 0.88 | 3.59E-05 | 2534.7 | 728.5 | NA | 23.7 | 25.0 | 223.2 |
| **stage2** | **stage3_free** | **-2.37** | **2.67E-03** | **V12B01_00260** | **EAP93512.1** | **acetylglutamate kinase** | **Amino Acid Biosynthesis** | **K00930** | **F** | **Belongs to the acetylglutamate kinase family. ArgB subfamily** | **argB** | **499.8** | **0.77** | **1.30E-04** | **245.2** | **243.3** | **352.0** | **2730.8** | **1181.7** | **NA** |
| stage3_free | stage3_shell | 3.27 | 2.72E-05 | V12B01_00260 | EAP93512.1 | acetylglutamate kinase | Amino Acid Biosynthesis | K00930 | F | Belongs to the acetylglutamate kinase family. ArgB subfamily | argB | 499.8 | 0.76 | 1.54E-06 | 2730.8 | 1181.7 | NA | 54.7 | 23.0 | 252.6 |
| stage3_free | stage3_shell | 3.26 | 2.32E-05 | V12B01_00265 | EAP93513.1 | N-acetyl-gamma-glutamyl-phosphate reductase | Amino Acid Biosynthesis | K00145 | E | Catalyzes the NADPH-dependent reduction of N-acetyl-5- glutamyl phosphate to yield N-acetyl-L-glutamate 5-semialdehyde | argC | 768.5 | 0.74 | 1.26E-06 | 2324.1 | 1914.8 | NA | 154.2 | 30.1 | 192.6 |
| **stage2** | **stage3_free** | **-2.17** | **4.26E-05** | **V12B01_00270** | **EAP93514.1** | **acetylornithine deacetylase** | **Amino Acid Biosynthesis** | **K01438** | **E** | **Belongs to the peptidase M20A family. ArgE subfamily** | **argE** | **85.7** | **0.49** | **8.64E-07** | **58.7** | **48.6** | **36.8** | **339.5** | **159.7** | **NA** |
| stage3_free | stage3_shell | 3.09 | 6.79E-08 | V12B01_00270 | EAP93514.1 | acetylornithine deacetylase | Amino Acid Biosynthesis | K01438 | E | Belongs to the peptidase M20A family. ArgE subfamily | argE | 85.7 | 0.55 | 1.39E-09 | 339.5 | 159.7 | NA | 12.6 | 16.0 | 30.0 |
| stage2 | stage3_shell | 2.41 | 2.89E-03 | V12B01_00275 | EAP93515.1 | phosphoenolpyruvate carboxylase | TCA cycle | K01595 | H | Forms oxaloacetate, a four-carbon dicarboxylic acid source for the tricarboxylic acid cycle | ppc | 280.4 | 0.87 | 5.35E-04 | 248.8 | 63.4 | 55.5 | 1.2 | 21.9 | 14.1 |
| stage2 | stage3_shell | 2.66 | 1.33E-03 | V12B01_00285 | EAP93517.1 | 5,10-methylenetetrahydrofolate reductase | Cofactor, Carrier, and Vitamin Biosynthesis | K00297 | E | reductase | metF | 24.2 | 0.89 | 2.11E-04 | 47.4 | 27.2 | 87.0 | 0.0 | 3.4 | 12.7 |
| **stage2** | **stage3_shell** | **-3.78** | **5.38E-06** | **V12B01_00365** | **EAP93533.1** | **hypothetical protein** | | | **S** | **Protein of unknown function (DUF3135)** | **VV3022** | **32.3** | **0.83** | **3.52E-07** | **2.4** | **5.7** | **3.4** | **25.8** | **56.9** | **61.2** |
| **stage3_free** | **stage3_shell** | **-2.56** | **2.66E-03** | **V12B01_00365** | **EAP93533.1** | **hypothetical protein** | | | **S** | **Protein of unknown function (DUF3135)** | **VV3022** | **32.3** | **0.91** | **3.79E-04** | **12.3** | **3.9** | **NA** | **25.8** | **56.9** | **61.2** |
| stage2 | stage3_shell | 2.21 | 1.04E-03 | V12B01_00420 | EAP91672.1 | serine hydroxymethyltransferase | Amino Acid Biosynthesis | K00600 | E | Catalyzes the reversible interconversion of serine and glycine with tetrahydrofolate (THF) serving as the one-carbon carrier. This reaction serves as the major source of one-carbon groups required for the biosynthesis of purines, thymidylate, methionine, and other important biomolecules. Also exhibits THF-independent aldolase activity toward beta-hydroxyamino acids, producing glycine and aldehydes, via a retro-aldol mechanism | glyA | 175.4 | 0.69 | 1.56E-04 | 518.8 | 148.4 | 126.4 | 34.6 | 5.7 | 59.0 |
| stage3_free | stage3_shell | 3.23 | 2.97E-05 | V12B01_00420 | EAP91672.1 | serine hydroxymethyltransferase | Amino Acid Biosynthesis | K00600 | E | Catalyzes the reversible interconversion of serine and glycine with tetrahydrofolate (THF) serving as the one-carbon carrier. This reaction serves as the major source of one-carbon groups required for the biosynthesis of purines, thymidylate, methionine, and other important biomolecules. Also exhibits THF-independent aldolase activity toward beta-hydroxyamino acids, producing glycine and aldehydes, via a retro-aldol mechanism | glyA | 175.4 | 0.76 | 1.72E-06 | 643.4 | 435.6 | NA | 34.6 | 5.7 | 59.0 |
| stage2 | stage3_shell | 2.06 | 8.44E-06 | V12B01_00475 | EAP91721.1 | ribosomal protein L17 | Translation | K02879 | J | Ribosomal protein L17 | rplQ | 557.8 | 0.45 | 6.19E-07 | 3833.9 | 2045.1 | 1982.1 | 213.9 | 516.2 | 508.2 |
| **stage2** | **stage3_shell** | **-4.48** | **3.03E-15** | **V12B01_00602** | **EAP92072.1** | **putative stomatin-like protein** | **Membrane** | | **O** | **COG0330 Membrane protease subunits, stomatin prohibitin homologs** | **-** | **321.7** | **0.55** | **2.17E-17** | **3.3** | **11.1** | **6.3** | **162.2** | **147.9** | **67.8** |
| **stage3_free** | **stage3_shell** | **-3.83** | **2.31E-09** | **V12B01_00602** | **EAP92072.1** | **putative stomatin-like protein** | **Membrane** | | **O** | **COG0330 Membrane protease subunits, stomatin prohibitin homologs** | **-** | **321.7** | **0.62** | **3.31E-11** | **5.7** | **15.7** | **NA** | **162.2** | **147.9** | **67.8** |
| **stage2** | **stage3_shell** | **-4.61** | **8.16E-17** | **V12B01_00607** | **EAP92073.1** | **hypothetical membrane protein** | **Membrane** | **K07403** | **O** | **COG1030 Membrane-bound serine protease (ClpP class)** | **-** | **247.6** | **0.53** | **4.65E-19** | **1.9** | **3.8** | **3.5** | **56.8** | **68.8** | **54.6** |
| **stage3_free** | **stage3_shell** | **-4.16** | **5.19E-11** | **V12B01_00607** | **EAP92073.1** | **hypothetical membrane protein** | **Membrane** | **K07403** | **O** | **COG1030 Membrane-bound serine protease (ClpP class)** | **-** | **247.6** | **0.61** | **4.93E-13** | **4.2** | **4.1** | **NA** | **56.8** | **68.8** | **54.6** |
| **stage2** | **stage3_shell** | **-2.41** | **1.03E-12** | **V12B01_00747** | **EAP95276.1** | **inosine monophosphate dehydrogenase-related protein** | **Nucleoside and Nucleotide Degradation** | | **S** | **Domain in cystathionine beta-synthase and other proteins.** | **-** | **243.1** | **0.32** | **1.26E-14** | **77.8** | **123.2** | **114.5** | **344.8** | **722.0** | **232.1** |
| **stage3_free** | **stage3_shell** | **-2.82** | **3.87E-13** | **V12B01_00747** | **EAP95276.1** | **inosine monophosphate dehydrogenase-related protein** | **Nucleoside and Nucleotide Degradation** | | **S** | **Domain in cystathionine beta-synthase and other proteins.** | **-** | **243.1** | **0.37** | **2.12E-15** | **72.4** | **85.2** | **NA** | **344.8** | **722.0** | **232.1** |

| | | | | | | | | | | | | | | | | | | | | |
|---|---|---|---|---|---|---|---|---|---|---|---|---|---|---|---|---|---|---|---|---|
| stage2 | stage3_shell | 3.28 | 4.07E-03 | V12B01_00777 | EAP95282.1 | Thiol-disulfide isomerase | | K03673 | O | Thiol disulfide interchange protein | - | 9.0 | 1.17 | 7.95E-04 | 42.2 | 57.8 | 25.5 | 5.3 | 0.0 | 0.0 |
| stage2 | stage3_free | 5.22 | 1.92E-03 | V12B01_00857 | EAP95298.1 | cytochrome o ubiquinol oxidase, subunit III | Aerobic Respiration | K02299 | C | oxidase, subunit | cyoC | 26.2 | 2.19 | 9.18E-05 | 11.5 | 12.6 | 5.6 | 0.0 | 0.0 | NA |
| **stage3_free** | **stage3_shell** | **-6.02** | **6.13E-04** | **V12B01_00857** | **EAP95298.1** | **cytochrome o ubiquinol oxidase, subunit III** | **Aerobic Respiration** | **K02299** | **C** | **oxidase, subunit** | **cyoC** | **26.2** | **2.61** | **6.39E-05** | **0.0** | **0.0** | **NA** | **5.2** | **17.1** | **4.6** |
| **stage3_free** | **stage3_shell** | **-2.13** | **1.37E-04** | **V12B01_00892** | **EAP95305.1** | **putative transcriptional regulator** | **Regulation** | | **K** | **Transcriptional regulator** | **VPA1607** | **29.3** | **0.55** | **1.09E-05** | **14.0** | **13.7** | **NA** | **38.5** | **63.0** | **50.0** |
| **stage3_free** | **stage3_shell** | **-3.33** | **1.16E-02** | **V12B01_00902** | **EAP95307.1** | **hypothetical protein** | | | **-** | **-** | **VPA1605** | **8.2** | **1.53** | **2.39E-03** | **0.0** | **2.6** | **NA** | **0.0** | **40.3** | **15.2** |
| stage2 | stage3_free | 2.14 | 1.86E-02 | V12B01_00952 | EAP95317.1 | ferric aerobactin receptor precursor | Regulation | K02014 | M | COG1629 Outer membrane receptor proteins, mostly Fe transport | iutA | 97.3 | 1.09 | 1.47E-03 | 34.8 | 3.9 | 3.0 | 1.7 | 2.2 | NA |
| stage2 | stage3_free | 2.12 | 1.54E-04 | V12B01_00992 | EAP95325.1 | putative membrane protein, suppressor for copper-sensitivity A | Membrane | | - | - | - | 46.0 | 0.53 | 4.08E-06 | 190.3 | 329.1 | 239.0 | 61.6 | 39.7 | NA |
| **stage3_free** | **stage3_shell** | **-2.59** | **1.50E-02** | **V12B01_01007** | **EAP95328.1** | **hypothetical protein** | | | **S** | **Protein of unknown function (DUF2860)** | **-** | **5.4** | **1.29** | **3.30E-03** | **0.9** | **1.0** | **NA** | **3.2** | **10.6** | **8.6** |
| **stage3_free** | **stage3_shell** | **-3.59** | **1.10E-03** | **V12B01_01022** | **EAP95331.1** | **hypothetical protein** | | **K01053** | **G** | **SMP-30/Gluconolaconase/LRE-like region** | **-** | **8.3** | **1.13** | **1.30E-04** | **1.0** | **1.1** | **NA** | **3.6** | **30.8** | **6.4** |
| **stage2** | **stage3_shell** | **-2.47** | **2.59E-03** | **V12B01_01047** | **EAP95336.1** | **hypothetical protein** | | | **-** | **#N/A** | **#N/A** | **10.6** | **0.87** | **4.64E-04** | **3.9** | **2.3** | **9.0** | **45.9** | **13.4** | **25.1** |
| **stage2** | **stage3_shell** | **-2.53** | **6.38E-06** | **V12B01_01092** | **EAP95345.1** | **Rare lipoprotein A** | **Membrane** | **K03642** | **M** | **Lytic transglycosylase with a strong preference for naked glycan strands that lack stem peptides** | **-** | **56.0** | **0.55** | **4.36E-07** | **8.8** | **26.6** | **37.9** | **56.4** | **116.9** | **176.0** |
| **stage3_free** | **stage3_shell** | **-3.50** | **2.99E-07** | **V12B01_01092** | **EAP95345.1** | **Rare lipoprotein A** | **Membrane** | **K03642** | **M** | **Lytic transglycosylase with a strong preference for naked glycan strands that lack stem peptides** | **-** | **56.0** | **0.67** | **7.70E-09** | **8.1** | **17.0** | **NA** | **56.4** | **116.9** | **176.0** |
| **stage2** | **stage3_shell** | **-2.56** | **1.50E-04** | **V12B01_01097** | **EAP95346.1** | **hypothetical protein** | | | **-** | **-** | **-** | **89.2** | **0.67** | **1.65E-05** | **4.8** | **10.5** | **13.1** | **65.9** | **77.4** | **9.4** |
| **stage3_free** | **stage3_shell** | **-2.17** | **3.44E-03** | **V12B01_01097** | **EAP95346.1** | **hypothetical protein** | | | **-** | **-** | **-** | **89.2** | **0.78** | **5.13E-04** | **3.0** | **20.2** | **NA** | **65.9** | **77.4** | **9.4** |
| **stage2** | **stage3_shell** | **-3.22** | **6.82E-06** | **V12B01_01102** | **EAP95347.1** | **hypothetical protein** | | | **S** | **nuclease activity** | **-** | **137.6** | **0.71** | **4.78E-07** | **8.9** | **27.4** | **39.5** | **364.3** | **266.3** | **22.8** |
| **stage3_free** | **stage3_shell** | **-3.70** | **7.32E-06** | **V12B01_01102** | **EAP95347.1** | **hypothetical protein** | | | **S** | **nuclease activity** | **-** | **137.6** | **0.83** | **3.24E-07** | **21.3** | **14.1** | **NA** | **364.3** | **266.3** | **22.8** |
| stage2 | stage3_shell | 2.67 | 3.50E-02 | V12B01_01162 | EAP95359.1 | sugar transporter family protein | Transport Proteins | | EGP | MFS_1 like family | - | 4.2 | 2.38 | 1.08E-02 | 6.2 | 4.9 | 4.6 | 0.0 | 0.0 | 0.0 |
| stage3_free | stage3_shell | 3.28 | 2.57E-02 | V12B01_01162 | EAP95359.1 | sugar transporter family protein | Transport Proteins | | EGP | MFS_1 like family | - | 4.2 | 2.52 | 6.47E-03 | 7.4 | 7.0 | NA | 0.0 | 0.0 | 0.0 |
| stage2 | stage3_shell | 3.74 | 9.43E-03 | V12B01_01167 | EAP95360.1 | hypothetical protein | | | S | SnoaL-like domain | - | 5.3 | 2.28 | 2.17E-03 | 15.7 | 6.6 | 6.4 | 0.0 | 0.0 | 0.0 |
| stage3_free | stage3_shell | 3.48 | 1.74E-02 | V12B01_01167 | EAP95360.1 | hypothetical protein | | | S | SnoaL-like domain | - | 5.3 | 2.35 | 3.95E-03 | 13.5 | 4.7 | NA | 0.0 | 0.0 | 0.0 |
| **stage2** | **stage3_shell** | **-2.01** | **1.01E-06** | **V12B01_01182** | **EAP95363.1** | **sulfatase family protein** | | | **P** | **C-terminal region of aryl-sulfatase** | **-** | **1464.3** | **0.40** | **5.49E-08** | **273.7** | **359.6** | **332.6** | **591.1** | **1949.7** | **628.9** |
| **stage2** | **stage3_shell** | **-2.42** | **1.81E-13** | **V12B01_01187** | **EAP95364.1** | **hypothetical protein** | | | **S** | **Domain of unknown function (DUF4345)** | **-** | **200.2** | **0.31** | **1.83E-15** | **126.6** | **93.5** | **127.2** | **356.4** | **709.9** | **357.7** |
| **stage3_free** | **stage3_shell** | **-2.45** | **5.99E-07** | **V12B01_01192** | **EAP95365.1** | **hypothetical protein** | | | **I** | **long-chain fatty acid transport protein** | **-** | **526.2** | **0.47** | **1.72E-08** | **156.6** | **205.6** | **NA** | **580.2** | **1444.2** | **413.6** |
| **stage3_free** | **stage3_shell** | **-4.55** | **8.68E-03** | **V12B01_01197** | **EAP95366.1** | **hypothetical protein** | | | **-** | **-** | **-** | **3.9** | **2.75** | **1.61E-03** | **0.0** | **0.0** | **NA** | **5.7** | **10.8** | **0.0** |
| **stage2** | **stage3_shell** | **-2.13** | **1.03E-05** | **V12B01_01237** | **EAP95374.1** | **glutamate synthase domain protein** | | **K22083** | **E** | **Belongs to the glutamate synthase family** | **-** | **196.8** | **0.47** | **7.90E-07** | **11.9** | **19.0** | **20.4** | **36.8** | **120.0** | **29.1** |
| **stage3_free** | **stage3_shell** | **-2.38** | **6.16E-06** | **V12B01_01242** | **EAP95375.1** | **hypothetical protein** | | | **S** | **Protein of unknown function (DUF1439)** | **-** | **66.9** | **0.50** | **2.59E-07** | **14.8** | **31.0** | **NA** | **68.7** | **160.0** | **61.2** |
| **stage2** | **stage3_shell** | **-2.67** | **3.52E-04** | **V12B01_01247** | **EAP95376.1** | **sigma-54 dependent transcriptional regulator** | **Regulation** | | **K** | **Bacterial regulatory protein, Fis family** | **-** | **22.3** | **0.76** | **4.39E-05** | **0.7** | **2.6** | **1.2** | **2.4** | **15.8** | **8.5** |
| **stage3_free** | **stage3_shell** | **-2.09** | **9.10E-03** | **V12B01_01247** | **EAP95376.1** | **sigma-54 dependent transcriptional regulator** | | | **K** | **Bacterial regulatory protein, Fis family** | **-** | **22.3** | **0.86** | **1.71E-03** | **3.4** | **0.7** | **NA** | **2.4** | **15.8** | **8.5** |
| **stage2** | **stage3_shell** | **-2.12** | **3.31E-03** | **V12B01_01277** | **EAP95382.1** | **hypothetical protein** | | | **S** | **Protein of unknown function (DUF3429)** | **VPA1252** | **52.6** | **0.74** | **6.25E-04** | **7.6** | **9.1** | **11.3** | **13.7** | **100.2** | **0.0** |
| **stage3_free** | **stage3_shell** | **-2.17** | **7.97E-03** | **V12B01_01277** | **EAP95382.1** | **hypothetical protein** | | | **S** | **Protein of unknown function (DUF3429)** | **VPA1252** | **52.6** | **0.88** | **1.45E-03** | **10.7** | **6.2** | **NA** | **13.7** | **100.2** | **0.0** |
| **stage2** | **stage3_free** | **-2.25** | **3.28E-02** | **V12B01_01297** | **EAP95386.1** | **hypothetical protein** | | | **-** | **-** | **-** | **5.7** | **1.21** | **3.09E-03** | **1.3** | **8.5** | **1.9** | **49.4** | **18.8** | **NA** |
| stage2 | stage3_shell | 2.03 | 4.68E-03 | V12B01_01392 | EAP95405.1 | hypothetical protein | | K09948 | S | protein conserved in bacteria | - | 22.3 | 0.72 | 9.34E-04 | 142.7 | 97.1 | 129.3 | 19.9 | 28.3 | 0.0 |
| **stage2** | **stage3_shell** | **-2.75** | **NA** | **V12B01_01562** | **EAP93176.1** | **aspartate ammonia-lyase** | **Amino Acid Degradation** | **K01744** | **E** | **Aspartate ammonia-lyase** | **aspA** | **366.5** | **0.89** | **NA** | **53.6** | **58.5** | **107.0** | **188.2** | **167.8** | **953.0** |
| **stage2** | **stage3_free** | **-2.01** | **1.60E-02** | **V12B01_01567** | **EAP93177.1** | **anaerobic C4-dicarboxylate transporter** | **Transport Proteins** | **K07791** | **S** | **Responsible for the transport of C4-dicarboxylates from the periplasm across the inner membrane** | **dcuA** | **54.6** | **0.85** | **1.20E-03** | **18.0** | **12.7** | **19.6** | **98.3** | **97.7** | **NA** |
| **stage2** | **stage3_shell** | **-5.74** | **3.09E-06** | **V12B01_01607** | **EAP93185.1** | **hypothetical arginine repressor** | **Regulation** | **K03402** | **K** | **Regulates arginine biosynthesis genes** | **ARGR** | **10.8** | **1.23** | **1.91E-07** | **1.8** | **0.0** | **1.3** | **32.7** | **18.6** | **104.8** |
| **stage2** | **stage3_shell** | **-2.06** | **NA** | **V12B01_01627** | **EAP93189.1** | **Predicted membrane protein** | **Membrane** | | **S** | **membrane** | **-** | **367.6** | **1.26** | **NA** | **316.4** | **281.2** | **481.5** | **5864.4** | **221.0** | **277.3** |
| **stage2** | **stage3_free** | **-2.41** | **6.45E-04** | **V12B01_01632** | **EAP93190.1** | **Predicted symporter** | **Transport Proteins** | **K14393** | **S** | **Belongs to the sodium solute symporter (SSF) (TC 2.A.21) family** | **actP** | **702.5** | **0.67** | **2.27E-05** | **220.5** | **36.4** | **194.5** | **991.5** | **1015.1** | **NA** |
| stage3_free | stage3_shell | 2.05 | 2.37E-03 | V12B01_01632 | EAP93190.1 | Predicted symporter | Transport Proteins | K14393 | S | Belongs to the sodium solute symporter (SSF) (TC 2.A.21) family | actP | 702.5 | 0.66 | 3.29E-04 | 991.5 | 1015.1 | NA | 289.7 | 78.1 | 76.9 |
| stage2 | stage3_shell | 2.03 | 9.53E-05 | V12B01_01647 | EAP93193.1 | sensor histidine kinase | Regulation | | T | COG0591 Na proline symporter | - | 56.0 | 0.50 | 9.87E-06 | 34.0 | 29.6 | 34.1 | 3.7 | 7.0 | 5.0 |
| **stage2** | **stage3_free** | **-5.67** | **1.53E-07** | **V12B01_01657** | **EAP93195.1** | **predicted signal-transduction protein** | **Regulation** | **K07182** | **T** | **containing cAMP-binding and CBS domains** | **-** | **85.1** | **1.01** | **1.71E-09** | **1.9** | **1.5** | **4.9** | **211.3** | **160.6** | **NA** |
| **stage2** | **stage3_shell** | **-2.61** | **3.60E-03** | **V12B01_01657** | **EAP93195.1** | **predicted signal-transduction protein** | **Regulation** | **K07182** | **T** | **containing cAMP-binding and CBS domains** | **-** | **85.1** | **1.00** | **6.82E-04** | **1.9** | **1.5** | **4.9** | **34.9** | **0.0** | **18.6** |
| stage3_free | stage3_shell | 2.27 | 1.67E-02 | V12B01_01657 | EAP93195.1 | predicted signal-transduction protein | Regulation | K07182 | T | containing cAMP-binding and CBS domains | - | 85.1 | 1.05 | 3.74E-03 | 211.3 | 160.6 | NA | 34.9 | 0.0 | 18.6 |
| **stage2** | **stage3_free** | **-3.55** | **3.46E-03** | **V12B01_01662** | **EAP93196.1** | **DNA polymerase III subunit epsilon** | **Replication** | **K02342** | **L** | **DNA polymerase III, epsilon subunit** | **-** | **8.9** | **1.21** | **1.81E-04** | **2.8** | **0.0** | **5.9** | **68.6** | **43.3** | **NA** |
| stage3_free | stage3_shell | 3.51 | 1.13E-02 | V12B01_01662 | EAP93196.1 | DNA polymerase III subunit epsilon | Replication | K02342 | L | DNA polymerase III, epsilon subunit | - | 8.9 | 1.54 | 2.30E-03 | 68.6 | 43.3 | NA | 0.0 | 4.8 | 0.0 |

| | | | | | | | | | | | | | | | | | | | | |
|---|---|---|---|---|---|---|---|---|---|---|---|---|---|---|---|---|---|---|---|---|
| **stage2** | **stage3_free** | **-5.03** | **1.60E-13** | **V12B01_01667** | **EAP93197.1** | **acetyl-coenzyme A synthetase** | **Carbohydrates and Carboxylates Degradation** | **K01895** | **F** | **Catalyzes the conversion of acetate into acetyl-CoA (AcCoA), an essential intermediate at the junction of anabolic and catabolic pathways. AcsA undergoes a two-step reaction. In the first half reaction, AcsA combines acetate with ATP to form acetyl-adenylate (AcAMP) intermediate. In the second half reaction, it can then transfer the acetyl group from AcAMP to the sulfhydryl group of CoA, forming the product AcCoA** | **acsA** | **2273.1** | **0.65** | **3.64E-16** | **156.2** | **78.3** | **182.9** | **6350.4** | **4142.2** | **NA** |
| **stage2** | **stage3_shell** | **-2.13** | **3.39E-04** | **V12B01_01667** | **EAP93197.1** | **acetyl-coenzyme A synthetase** | **Carbohydrates and Carboxylates Degradation** | **K01895** | **F** | **Catalyzes the conversion of acetate into acetyl-CoA (AcCoA), an essential intermediate at the junction of anabolic and catabolic pathways. AcsA undergoes a two-step reaction. In the first half reaction, AcsA combines acetate with ATP to form acetyl-adenylate (AcAMP) intermediate. In the second half reaction, it can then transfer the acetyl group from AcAMP to the sulfhydryl group of CoA, forming the product AcCoA** | **acsA** | **2273.1** | **0.59** | **4.17E-05** | **156.2** | **78.3** | **182.9** | **866.1** | **92.3** | **559.2** |
| stage3_free | stage3_shell | 2.63 | 1.03E-04 | V12B01_01667 | EAP93197.1 | acetyl-coenzyme A synthetase | Carbohydrates and Carboxylates Degradation | K01895 | F | Catalyzes the conversion of acetate into acetyl-CoA (AcCoA), an essential intermediate at the junction of anabolic and catabolic pathways. AcsA undergoes a two-step reaction. In the first half reaction, AcsA combines acetate with ATP to form acetyl-adenylate (AcAMP) intermediate. In the second half reaction, it can then transfer the acetyl group from AcAMP to the sulfhydryl group of CoA, forming the product AcCoA | acsA | 2273.1 | 0.65 | 7.69E-06 | 6350.4 | 4142.2 | NA | 866.1 | 92.3 | 559.2 |
| stage2 | stage3_shell | 2.61 | 5.78E-07 | V12B01_01672 | EAP93198.1 | 3-dehydroquinate dehydratase | Other Biosynthesis | K03786 | E | Catalyzes a trans-dehydration via an enolate intermediate | aroQ | 72.8 | 0.50 | 2.86E-08 | 445.7 | 308.5 | 305.1 | 56.8 | 47.1 | 6.3 |
| stage3_free | stage3_shell | 2.12 | 2.22E-04 | V12B01_01672 | EAP93198.1 | 3-dehydroquinate dehydratase | Other Biosynthesis | K03786 | E | Catalyzes a trans-dehydration via an enolate intermediate | aroQ | 72.8 | 0.54 | 1.91E-05 | 264.2 | 258.3 | NA | 56.8 | 47.1 | 6.3 |
| stage2 | stage3_shell | 2.50 | 9.56E-06 | V12B01_01677 | EAP93199.1 | acetyl-CoA carboxylase | | K02160 | I | first, biotin carboxylase catalyzes the carboxylation of the carrier protein and then the transcarboxylase transfers the carboxyl group to form malonyl-CoA | accB | 117.7 | 0.55 | 7.28E-07 | 884.4 | 381.5 | 380.3 | 91.7 | 46.8 | 50.3 |
| stage3_free | stage3_shell | 2.20 | 4.18E-04 | V12B01_01677 | EAP93199.1 | acetyl-CoA carboxylase | | K02160 | I | first, biotin carboxylase catalyzes the carboxylation of the carrier protein and then the transcarboxylase transfers the carboxyl group to form malonyl-CoA | accB | 117.7 | 0.60 | 4.14E-05 | 468.5 | 474.0 | NA | 91.7 | 46.8 | 50.3 |
| stage2 | stage3_shell | 2.66 | 1.90E-11 | V12B01_01682 | EAP93200.1 | acetyl-CoA carboxylase | | K01961 | I | Biotin carboxylase | accC | 444.6 | 0.38 | 2.96E-13 | 895.6 | 559.2 | 665.9 | 44.0 | 81.0 | 103.8 |
| stage3_free | stage3_shell | 2.22 | 6.61E-07 | V12B01_01682 | EAP93200.1 | acetyl-CoA carboxylase | | K01961 | I | Biotin carboxylase | accC | 444.6 | 0.42 | 1.95E-08 | 542.1 | 539.0 | NA | 44.0 | 81.0 | 103.8 |
| stage3_free | stage3_shell | 2.73 | 5.85E-07 | V12B01_01707 | EAP93205.1 | bifunctional phosphoribosylaminoimidazolecarboxamide formyltransferase/IMP cyclohydrolase | Nucleoside and Nucleotide Biosynthesis | K00602 | F | bifunctional purine biosynthesis protein purh | purH | 214.1 | 0.52 | 1.67E-08 | 532.3 | 505.7 | NA | 61.5 | 29.1 | 60.1 |
| stage3_free | stage3_shell | 3.00 | 2.58E-07 | V12B01_01712 | EAP93206.1 | phosphoribosylamine--glycine ligase | Nucleoside and Nucleotide Biosynthesis | K01945 | F | Belongs to the GARS family | purD | 151.3 | 0.56 | 6.31E-09 | 514.8 | 290.5 | NA | 19.8 | 32.8 | 44.1 |
| stage2 | stage3_shell | 4.09 | 7.32E-06 | V12B01_01792 | EAP91796.1 | amino acid ABC transporter, periplasmic amino acid-binding portion | Transport Proteins | K16961 | ET | belongs to the bacterial solute-binding protein 3 family | patH | 639.8 | 0.94 | 5.25E-07 | 486.3 | 393.3 | 1514.6 | 13.6 | 5.1 | 58.1 |
| stage3_free | stage3_shell | 4.19 | 3.90E-05 | V12B01_01792 | EAP91796.1 | amino acid ABC transporter, periplasmic amino acid-binding portion | Transport Proteins | K16961 | ET | belongs to the bacterial solute-binding protein 3 family | patH | 639.8 | 1.02 | 2.49E-06 | 1158.2 | 639.7 | NA | 13.6 | 5.1 | 58.1 |
| stage3_free | stage3_shell | 2.60 | 1.24E-08 | V12B01_01822 | EAP91802.1 | putative inner membrane protein translocase component YidC | Membrane | K03217 | U | Required for the insertion and or proper folding and or complex formation of integral membrane proteins into the membrane. Involved in integration of membrane proteins that insert both dependently and independently of the Sec translocase complex, as well as at least some lipoproteins. Aids folding of multispanning membrane proteins | yidC | 173.9 | 0.43 | 2.04E-10 | 298.7 | 319.4 | NA | 51.2 | 23.3 | 28.1 |
| stage2 | stage3_shell | 2.01 | 1.64E-12 | V12B01_01872 | EAP92816.1 | ATP synthase subunit A | Transport Proteins | K02108 | C | it plays a direct role in the translocation of protons across the membrane | atpB | 1518.9 | 0.27 | 2.11E-14 | 5953.4 | 3640.3 | 4594.8 | 792.9 | 773.5 | 888.4 |
| stage2 | stage3_shell | 2.23 | 6.32E-10 | V12B01_01882 | EAP92818.1 | ATP synthase subunit B | Transport Proteins | K02109 | C | Component of the F(0) channel, it forms part of the peripheral stalk, linking F(1) to F(0) | atpF | 1019.8 | 0.34 | 1.31E-11 | 6760.9 | 4127.5 | 5516.3 | 491.4 | 927.2 | 961.3 |

| | | | | | | | | | | | | | | | | | | | | |
|---|---|---|---|---|---|---|---|---|---|---|---|---|---|---|---|---|---|---|---|---|
| stage2 | stage3_shell | 2.30 | 1.05E-09 | V12B01_01887 | EAP92819.1 | ATP synthase subunit D | Transport Proteins | K02113 | C | F(1)F(0) ATP synthase produces ATP from ADP in the presence of a proton or sodium gradient. F-type ATPases consist of two structural domains, F(1) containing the extramembraneous catalytic core and F(0) containing the membrane proton channel, linked together by a central stalk and a peripheral stalk. During catalysis, ATP synthesis in the catalytic domain of F(1) is coupled via a rotary mechanism of the central stalk subunits to proton translocation | atpH | 937.3 | 0.36 | 2.31E-11 | 6202.0 | 3620.0 | 4525.4 | 368.0 | 821.6 | 773.1 |
| stage2 | stage3_shell | 2.18 | 3.33E-07 | V12B01_01892 | EAP92820.1 | ATP synthase subunit A | Transport Proteins | K02111 | C | Produces ATP from ADP in the presence of a proton gradient across the membrane. The alpha chain is a regulatory subunit | atpA | 2626.5 | 0.41 | 1.56E-08 | 6617.2 | 3092.5 | 3345.6 | 299.4 | 821.2 | 784.7 |
| stage2 | stage3_free | 2.64 | 3.53E-09 | V12B01_01897 | EAP92821.1 | ATP synthase subunit C | Transport Proteins | K02115 | C | Produces ATP from ADP in the presence of a proton gradient across the membrane. The gamma chain is believed to be important in regulating ATPase activity and the flow of protons through the CF(0) complex | atpG | 1437.9 | 0.43 | 2.57E-11 | 6154.1 | 3333.4 | 3832.8 | 690.5 | 634.4 | NA |
| stage2 | stage3_shell | 2.12 | 7.46E-08 | V12B01_01897 | EAP92821.1 | ATP synthase subunit C | Transport Proteins | K02115 | C | Produces ATP from ADP in the presence of a proton gradient across the membrane. The gamma chain is believed to be important in regulating ATPase activity and the flow of protons through the CF(0) complex | atpG | 1437.9 | 0.38 | 2.85E-09 | 6154.1 | 3333.4 | 3832.8 | 359.4 | 947.5 | 761.9 |
| stage2 | stage3_free | 2.73 | 3.45E-08 | V12B01_01902 | EAP92822.1 | ATP synthase subunit B | Transport Proteins | K02112 | C | Produces ATP from ADP in the presence of a proton gradient across the membrane. The catalytic sites are hosted primarily by the beta subunits | atpD | 2989.3 | 0.47 | 3.19E-10 | 7599.9 | 4143.4 | 4799.7 | 752.2 | 769.8 | NA |
| stage2 | stage3_shell | 2.15 | 5.93E-07 | V12B01_01902 | EAP92822.1 | ATP synthase subunit B | Transport Proteins | K02112 | C | Produces ATP from ADP in the presence of a proton gradient across the membrane. The catalytic sites are hosted primarily by the beta subunits | atpD | 2989.3 | 0.41 | 2.98E-08 | 7599.9 | 4143.4 | 4799.7 | 335.7 | 1141.1 | 971.4 |
| stage2 | stage3_free | 2.53 | 2.60E-07 | V12B01_01907 | EAP92823.1 | ATP synthase subunit epsilon | Transport Proteins | K02114 | C | Produces ATP from ADP in the presence of a proton gradient across the membrane | atpC | 703.8 | 0.47 | 3.03E-09 | 4798.8 | 3053.6 | 3380.2 | 594.7 | 590.5 | NA |
| stage2 | stage3_free | 3.07 | 1.23E-09 | V12B01_01927 | EAP92827.1 | threonine dehydratase | Amino Acid Degradation | K01754 | E | Catalyzes the anaerobic formation of alpha-ketobutyrate and ammonia from threonine in a two-step reaction. The first step involved a dehydration of threonine and a production of enamine intermediates (aminocrotonate), which tautomerizes to its imine form (iminobutyrate). Both intermediates are unstable and short- lived. The second step is the nonenzymatic hydrolysis of the enamine imine intermediates to form 2-ketobutyrate and free ammonia. In the low water environment of the cell, the second step is accelerated by RidA | ilvA | 463.9 | 0.48 | 7.07E-12 | 477.2 | 427.3 | 508.8 | 56.2 | 47.3 | NA |
| stage2 | stage3_shell | 2.44 | 5.94E-08 | V12B01_01927 | EAP92827.1 | threonine dehydratase | Amino Acid Degradation | K01754 | E | Catalyzes the anaerobic formation of alpha-ketobutyrate and ammonia from threonine in a two-step reaction. The first step involved a dehydration of threonine and a production of enamine intermediates (aminocrotonate), which tautomerizes to its imine form (iminobutyrate). Both intermediates are unstable and short- lived. The second step is the nonenzymatic hydrolysis of the enamine imine intermediates to form 2-ketobutyrate and free ammonia. In the low water environment of the cell, the second step is accelerated by RidA | ilvA | 463.9 | 0.44 | 2.22E-09 | 477.2 | 427.3 | 508.8 | 31.0 | 91.1 | 51.6 |
| stage2 | stage3_shell | 2.26 | 2.78E-05 | V12B01_01932 | EAP92828.1 | dihydroxy-acid dehydratase | Amino Acid Biosynthesis | K01687 | H | Belongs to the IlvD Edd family | ilvD | 326.8 | 0.53 | 2.44E-06 | 392.0 | 217.7 | 221.6 | 14.7 | 73.1 | 26.3 |
| stage2 | stage3_shell | 3.00 | 1.25E-06 | V12B01_01937 | EAP92829.1 | branched-chain amino acid aminotransferase | Amino Acid Degradation | K00826 | E | Belongs to the class-IV pyridoxal-phosphate-dependent aminotransferase family | ilvE | 87.2 | 0.61 | 6.90E-08 | 174.4 | 92.8 | 134.0 | 8.2 | 12.5 | 11.8 |
| stage3_free | stage3_shell | 2.12 | 2.28E-03 | V12B01_01957 | EAP92833.1 | hypothetical protein | | | S | Protein of unknown function (DUF2860) | - | 39.5 | 0.70 | 3.15E-04 | 87.3 | 75.5 | NA | 6.7 | 9.5 | 17.9 |
| stage2 | stage3_shell | 2.70 | 2.15E-09 | V12B01_01962 | EAP92834.1 | thiol:disulfide interchange protein | | K03673 | O | Thiol disulfide interchange protein | dsbA | 72.6 | 0.44 | 4.94E-11 | 397.2 | 246.7 | 247.0 | 21.3 | 25.2 | 52.2 |

| | | | | | | | | | | | | | | | | | | | | |
|---|---|---|---|---|---|---|---|---|---|---|---|---|---|---|---|---|---|---|---|---|
| stage3_free | stage3_shell | 3.00 | 4.89E-08 | V12B01_02052 | EAP92852.1 | phosphoribosylaminoimidazole carboxylase | Nucleoside and Nucleotide Biosynthesis | K01589 | F | Catalyzes the ATP-dependent conversion of 5- aminoimidazole ribonucleotide (AIR) and HCO(3)(-) to N5-carboxyaminoimidazole ribonucleotide (N5-CAIR) | purK | 90.3 | 0.52 | 9.28E-10 | 331.2 | 319.4 | NA | 25.4 | 29.4 | 25.2 |
| stage3_free | stage3_shell | 4.14 | 4.38E-05 | V12B01_02057 | EAP92853.1 | phosphoribosylaminoimidazole carboxylase catalytic subunit | Nucleoside and Nucleotide Biosynthesis | K01588 | F | Catalyzes the conversion of N5-carboxyaminoimidazole ribonucleotide (N5-CAIR) to 4-carboxy-5-aminoimidazole ribonucleotide (CAIR) | purE | 18.0 | 0.99 | 2.87E-06 | 190.1 | 147.3 | NA | 6.6 | 0.0 | 11.7 |
| stage2 | stage3_shell | 3.38 | 2.00E-07 | V12B01_02082 | EAP92858.1 | carbonic anhydrase, family 3 | | | S | COG0663 Carbonic anhydrases acetyltransferases, isoleucine patch superfamily | yrdA | 41.5 | 0.64 | 8.65E-09 | 191.4 | 195.3 | 233.8 | 8.8 | 11.1 | 20.9 |
| stage3_free | stage3_shell | 2.16 | 1.69E-03 | V12B01_02082 | EAP92858.1 | carbonic anhydrase, family 3 | | | S | COG0663 Carbonic anhydrases acetyltransferases, isoleucine patch superfamily | yrdA | 41.5 | 0.68 | 2.14E-04 | 93.8 | 98.5 | NA | 8.8 | 11.1 | 20.9 |
| stage2 | stage3_shell | 3.29 | 3.09E-07 | V12B01_02099 | EAP91810.1 | ketol-acid reductoisomerase | Amino Acid Biosynthesis | K00053 | EH | Involved in the biosynthesis of branched-chain amino acids (BCAA). Catalyzes an alkyl-migration followed by a ketol- acid reduction of (S)-2-acetolactate (S2AL) to yield (R)-2,3- dihydroxy-isovalerate. In the isomerase reaction, S2AL is rearranged via a Mg-dependent methyl migration to produce 3- hydroxy-3-methyl-2-ketobutyrate (HMKB). In the reductase reaction, this 2-ketoacid undergoes a metal-dependent reduction by NADPH to yield (R)-2,3-dihydroxy-isovalerate | ilvC | 614.0 | 0.63 | 1.43E-08 | 1711.6 | 800.9 | 753.3 | 52.7 | 56.0 | 97.8 |
| **stage2** | **stage3_shell** | **-2.44** | **1.68E-03** | **V12B01_02185** | **EAP95559.1** | **Integrase, catalytic region** | | **K07497** | **L** | **Evidence 2b Function of strongly homologous gene** | **-** | **11.0** | **0.80** | **2.75E-04** | **0.6** | **5.3** | **5.4** | **27.7** | **20.6** | **10.6** |
| **stage3_free** | **stage3_shell** | **-2.44** | **5.55E-03** | **V12B01_02185** | **EAP95559.1** | **Integrase, catalytic region** | | **K07497** | **L** | **Evidence 2b Function of strongly homologous gene** | **-** | **11.0** | **0.96** | **9.22E-04** | **2.8** | **4.2** | **NA** | **27.7** | **20.6** | **10.6** |
| stage3_free | stage3_shell | 2.26 | 3.74E-04 | V12B01_02190 | EAP95560.1 | glutathione reductase | Cofactor, Carrier, and Vitamin Biosynthesis | K00383 | C | Belongs to the class-I pyridine nucleotide-disulfide oxidoreductase family | gor | 110.2 | 0.62 | 3.65E-05 | 142.3 | 142.4 | NA | 18.9 | 11.2 | 25.2 |
| **stage2** | **stage3_shell** | **-2.66** | **6.33E-05** | **V12B01_02230** | **EAP95568.1** | **universal stress protein A** | **Regulation** | **K06149** | **T** | **COG0589 Universal stress protein UspA and related nucleotide-binding proteins** | **uspA** | **567.4** | **0.66** | **6.28E-06** | **276.0** | **340.3** | **528.7** | **576.7** | **1443.3** | **3717.7** |
| **stage3_free** | **stage3_shell** | **-2.13** | **3.39E-03** | **V12B01_02230** | **EAP95568.1** | **universal stress protein A** | **Regulation** | **K06149** | **T** | **COG0589 Universal stress protein UspA and related nucleotide-binding proteins** | **uspA** | **567.4** | **0.78** | **5.04E-04** | **462.6** | **554.3** | **NA** | **576.7** | **1443.3** | **3717.7** |
| **stage2** | **stage3_shell** | **-2.47** | **9.45E-11** | **V12B01_02235** | **EAP95569.1** | **ferritin** | | **K02217** | **P** | **Iron-storage protein** | **ftnA** | **2346.6** | **0.36** | **1.58E-12** | **563.4** | **1489.8** | **1442.7** | **3952.0** | **7991.3** | **3284.0** |
| **stage3_free** | **stage3_shell** | **-2.85** | **3.34E-11** | **V12B01_02235** | **EAP95569.1** | **ferritin** | | **K02217** | **P** | **Iron-storage protein** | **ftnA** | **2346.6** | **0.41** | **3.10E-13** | **865.3** | **921.6** | **NA** | **3952.0** | **7991.3** | **3284.0** |
| **stage2** | **stage3_shell** | **-2.56** | **1.75E-10** | **V12B01_02240** | **EAP95570.1** | **universal stress protein UspB** | **Regulation** | **K06144** | **S** | **Universal stress protein B homolog** | **uspB** | **1310.0** | **0.39** | **3.07E-12** | **576.4** | **1272.4** | **1072.0** | **3570.3** | **8334.8** | **2117.8** |
| **stage3_free** | **stage3_shell** | **-3.16** | **4.06E-12** | **V12B01_02240** | **EAP95570.1** | **universal stress protein UspB** | **Regulation** | **K06144** | **S** | **Universal stress protein B homolog** | **uspB** | **1310.0** | **0.43** | **3.00E-14** | **612.6** | **692.4** | **NA** | **3570.3** | **8334.8** | **2117.8** |
| stage2 | stage3_shell | 2.10 | 3.22E-08 | V12B01_02325 | EAP95587.1 | Cytochrome c4 | Respiration | | C | COG2863 Cytochrome c553 | cc4 | 375.1 | 0.36 | 1.05E-09 | 1410.3 | 1005.8 | 1207.3 | 294.7 | 120.0 | 165.9 |
| stage3_free | stage3_shell | 2.11 | 7.42E-07 | V12B01_02325 | EAP95587.1 | Cytochrome c4 | Respiration | | C | COG2863 Cytochrome c553 | cc4 | 375.1 | 0.40 | 2.27E-08 | 1247.7 | 1250.7 | NA | 294.7 | 120.0 | 165.9 |
| **stage3_free** | **stage3_shell** | **-2.01** | **5.41E-06** | **V12B01_02330** | **EAP95588.1** | **methyltransferase-related protein** | | | **S** | **2-polyprenyl-3-methyl-5-hydroxy-6-metoxy-1, 4-benzoquinol methylase** | **-** | **126.2** | **0.42** | **2.20E-07** | **58.0** | **108.3** | **NA** | **290.0** | **322.0** | **187.1** |
| **stage2** | **stage3_free** | **-2.16** | **3.28E-04** | **V12B01_02355** | **EAP95593.1** | **Response regulator for NtrB** | **Regulation** | **K07712** | **T** | **COG2204 Response regulator containing CheY-like receiver, AAA-type ATPase, and DNA-binding domains** | **glnG** | **100.1** | **0.56** | **9.91E-06** | **39.9** | **48.9** | **42.5** | **333.6** | **135.5** | **NA** |
| stage3_free | stage3_shell | 2.05 | 5.78E-04 | V12B01_02355 | EAP95593.1 | Response regulator for NtrB | Regulation | K07712 | T | COG2204 Response regulator containing CheY-like receiver, AAA-type ATPase, and DNA-binding domains | glnG | 100.1 | 0.57 | 5.97E-05 | 333.6 | 135.5 | NA | 15.9 | 61.4 | 28.4 |
| **stage2** | **stage3_free** | **-2.48** | **3.20E-05** | **V12B01_02360** | **EAP95594.1** | **nitrogen regulation protein** | **Regulation** | **K07708** | **T** | **nitrogen regulation protein** | **glnL** | **27.8** | **0.55** | **6.19E-07** | **16.7** | **16.5** | **13.6** | **124.9** | **79.8** | **NA** |
| stage3_free | stage3_shell | 3.41 | 4.93E-06 | V12B01_02360 | EAP95594.1 | nitrogen regulation protein | Regulation | K07708 | T | nitrogen regulation protein | glnL | 27.8 | 0.71 | 1.96E-07 | 124.9 | 79.8 | NA | 1.6 | 10.5 | 5.6 |
| **stage2** | **stage3_free** | **-2.30** | **2.13E-04** | **V12B01_02370** | **EAP95596.1** | **glutamine synthetase** | **Amino Acid Biosynthesis** | **K01915** | **E** | **glutamine synthetase** | **glnA** | **822.3** | **0.58** | **6.15E-06** | **778.6** | **385.3** | **229.1** | **3790.5** | **1781.8** | **NA** |
| stage3_free | stage3_shell | 4.15 | 9.55E-12 | V12B01_02370 | EAP95596.1 | glutamine synthetase | Amino Acid Biosynthesis | K01915 | E | glutamine synthetase | glnA | 822.3 | 0.58 | 8.06E-14 | 3790.5 | 1781.8 | NA | 95.2 | 52.6 | 147.4 |
| stage2 | stage3_free | 2.43 | 4.58E-10 | V12B01_02445 | EAP95611.1 | type II secretory pathway, pseudopilin EpsG | Secretion | K02456 | U | general secretion pathway protein | epsG | 136.3 | 0.37 | 2.32E-12 | 465.7 | 557.6 | 526.4 | 84.5 | 96.3 | NA |
| stage2 | stage3_free | 2.64 | 1.09E-05 | V12B01_02475 | EAP95617.1 | general secretion pathway protein M | Secretion | K02462 | U | Involved in a type II secretion system (T2SS, formerly general secretion pathway, GSP) for the export of proteins | epsM | 35.4 | 0.57 | 1.81E-07 | 116.4 | 150.7 | 134.0 | 15.0 | 22.6 | NA |
| **stage2** | **stage3_shell** | **-3.52** | **4.17E-09** | **V12B01_02525** | **EAP95627.1** | **hypothetical protein** | | | **-** | **#N/A** | **#N/A** | **92.8** | **0.58** | **1.06E-10** | **14.3** | **87.6** | **78.9** | **257.7** | **561.2** | **826.6** |
| **stage3_free** | **stage3_shell** | **-3.57** | **2.06E-07** | **V12B01_02525** | **EAP95627.1** | **hypothetical protein** | | | **-** | **#N/A** | **#N/A** | **92.8** | **0.67** | **4.91E-09** | **59.0** | **54.2** | **NA** | **257.7** | **561.2** | **826.6** |
| **stage2** | **stage3_shell** | **-2.93** | **1.51E-12** | **V12B01_02530** | **EAP95628.1** | **site-specific recombinase, phage integrase family** | | | **L** | **Phage integrase family** | **-** | **262.5** | **0.40** | **1.91E-14** | **13.7** | **38.5** | **38.8** | **127.0** | **262.7** | **151.6** |
| **stage3_free** | **stage3_shell** | **-4.04** | **1.39E-16** | **V12B01_02530** | **EAP95628.1** | **site-specific recombinase, phage integrase family** | | | **L** | **Phage integrase family** | **-** | **262.5** | **0.47** | **2.06E-19** | **14.5** | **13.7** | **NA** | **127.0** | **262.7** | **151.6** |
| **stage3_free** | **stage3_shell** | **-2.71** | **1.51E-04** | **V12B01_02570** | **EAP95636.1** | **hypothetical protein** | | | **-** | **-** | **-** | **55.2** | **0.72** | **1.21E-05** | **32.4** | **55.3** | **NA** | **513.7** | **165.1** | **100.8** |
| stage2 | stage3_shell | 2.03 | 8.97E-03 | V12B01_02575 | EAP95637.1 | hypothetical protein | | | - | #N/A | #N/A | 57.0 | 0.86 | 2.04E-03 | 120.6 | 177.7 | 105.1 | 28.6 | 0.0 | 23.8 |
| stage2 | stage3_shell | 3.69 | 3.20E-14 | V12B01_02580 | EAP95638.1 | hypothetical protein | | | - | #N/A | #N/A | 884.0 | 0.47 | 2.90E-16 | 10340.6 | 13278.3 | 9036.8 | 442.8 | 481.6 | 746.5 |

| | | | | | | | | | | | | | | | | | | | | |
|---|---|---|---|---|---|---|---|---|---|---|---|---|---|---|---|---|---|---|---|---|
| stage2 | stage3_shell | 2.27 | 5.35E-05 | V12B01_02585 | EAP95639.1 | toxin secretion, membrane fusion protein | | K13408 | M | HlyD family secretion protein | - | 72.7 | 0.55 | 5.11E-06 | 92.0 | 152.0 | 118.8 | 30.9 | 3.7 | 13.8 |
| stage2 | stage3_shell | 2.08 | 9.28E-03 | V12B01_02595 | EAP95641.1 | transposase | | | L | User locus_tag | - | 15.2 | 0.81 | 2.12E-03 | 35.5 | 55.4 | 35.8 | 5.1 | 9.7 | 0.0 |
| stage3_free | stage3_shell | -2.07 | 3.18E-09 | V12B01_02615 | EAP95645.1 | transcription elongation factor GreB | | | - | #N/A | #N/A | 177.6 | 0.33 | 4.63E-11 | 95.1 | 103.8 | NA | 276.5 | 388.3 | 287.7 |
| stage3_free | stage3_shell | 2.16 | 4.22E-08 | V12B01_02650 | EAP95652.1 | Guanosine-3',5'-bis(diphosphate) 3'-pyrophosphohydrolase | Metabolic Regulator Biosynthesis | K01139 | KT | In eubacteria ppGpp (guanosine 3'-diphosphate 5-' diphosphate) is a mediator of the stringent response that coordinates a variety of cellular activities in response to changes in nutritional abundance | spoT | 212.1 | 0.37 | 7.74E-10 | 266.8 | 291.8 | NA | 55.6 | 34.9 | 37.5 |
| stage3_free | stage3_shell | -3.00 | 1.02E-03 | V12B01_02690 | EAP95660.1 | hypothetical protein | | | S | Inovirus Gp2 | - | 13.3 | 0.94 | 1.18E-04 | 4.3 | 1.5 | NA | 29.7 | 21.1 | 17.7 |
| stage3_free | stage3_shell | -5.07 | 6.05E-03 | V12B01_02695 | EAP95661.1 | hypothetical protein | | | S | Inovirus Gp2 | - | 2.8 | 2.90 | 1.02E-03 | 0.0 | 0.0 | NA | 4.7 | 11.2 | 4.2 |
| stage3_free | stage3_shell | -2.60 | 3.67E-02 | V12B01_02840 | EAP95690.1 | putative lipoprotein | | | - | - | - | 1.8 | 3.70 | 1.01E-02 | 0.0 | 0.0 | NA | 7.5 | 7.1 | 0.0 |
| stage3_free | stage3_shell | -2.14 | 5.77E-03 | V12B01_02850 | EAP95692.1 | putative lipoprotein | | | - | - | - | 8.1 | 0.81 | 9.66E-04 | 9.1 | 7.9 | NA | 42.2 | 44.9 | 18.8 |
| stage2 | stage3_shell | -2.35 | 6.32E-10 | V12B01_02880 | EAP95698.1 | methyl-accepting chemotaxis protein, putative | Chemotaxis | K03406 | NT | Methyl-accepting chemotaxis protein | - | 1757.9 | 0.36 | 1.30E-11 | 143.6 | 302.8 | 281.7 | 1159.7 | 1412.2 | 412.8 |
| stage3_free | stage3_shell | -2.53 | 4.20E-09 | V12B01_02880 | EAP95698.1 | methyl-accepting chemotaxis protein, putative | Chemotaxis | K03406 | NT | Methyl-accepting chemotaxis protein | - | 1757.9 | 0.41 | 6.38E-11 | 188.5 | 238.8 | NA | 1159.7 | 1412.2 | 412.8 |
| stage2 | stage3_free | -2.21 | 4.92E-03 | V12B01_03000 | EAP95722.1 | orotate phosphoribosyltransferase | Nucleoside and Nucleotide Biosynthesis | K00762 | F | Catalyzes the transfer of a ribosyl phosphate group from 5-phosphoribose 1-diphosphate to orotate, leading to the formation of orotidine monophosphate (OMP) | pyrE | 35.0 | 0.77 | 2.75E-04 | 36.0 | 52.4 | 44.2 | 223.0 | 332.2 | NA |
| stage3_free | stage3_shell | 3.56 | 3.45E-05 | V12B01_03000 | EAP95722.1 | orotate phosphoribosyltransferase | Nucleoside and Nucleotide Biosynthesis | K00762 | F | Catalyzes the transfer of a ribosyl phosphate group from 5-phosphoribose 1-diphosphate to orotate, leading to the formation of orotidine monophosphate (OMP) | pyrE | 35.0 | 0.82 | 2.16E-06 | 223.0 | 332.2 | NA | 39.8 | 0.0 | 0.0 |
| stage3_free | stage3_shell | 2.35 | 1.24E-03 | V12B01_03033 | EAP91665.1 | orotidine 5'-phosphate decarboxylase | Nucleoside and Nucleotide Biosynthesis | K01591 | F | Catalyzes the decarboxylation of orotidine 5'- monophosphate (OMP) to uridine 5'-monophosphate (UMP) | pyrF | 27.6 | 0.74 | 1.48E-04 | 147.9 | 118.4 | NA | 5.6 | 5.3 | 39.8 |
| stage2 | stage3_free | -5.86 | 7.63E-36 | V12B01_03058 | EAP92109.1 | Alanine dehydrogenase | Amino Acid Degradation | K00259 | C | Belongs to the AlaDH PNT family | ald | 1340.6 | 0.45 | 1.68E-39 | 62.1 | 60.4 | 46.6 | 4197.5 | 2763.5 | NA |
| stage2 | stage3_shell | -4.19 | 6.02E-23 | V12B01_03058 | EAP92109.1 | Alanine dehydrogenase | Amino Acid Degradation | K00259 | C | Belongs to the AlaDH PNT family | ald | 1340.6 | 0.41 | 1.02E-25 | 62.1 | 60.4 | 46.6 | 1058.1 | 309.3 | 928.8 |
| stage2 | stage3_shell | -2.71 | 6.32E-04 | V12B01_03108 | EAP92119.1 | biotin synthesis protein BioC | Cofactor, Carrier, and Vitamin Biosynthesis | K02169 | H | Converts the free carboxyl group of a malonyl-thioester to its methyl ester by transfer of a methyl group from S-adenosyl- L-methionine (SAM). It allows to synthesize pimeloyl-ACP via the fatty acid synthetic pathway | bioC | 23.5 | 0.84 | 8.59E-05 | 4.4 | 8.8 | 7.4 | 19.9 | 26.3 | 77.9 |
| stage2 | stage3_shell | -2.67 | 2.58E-03 | V12B01_03113 | EAP92120.1 | dithiobiotin synthetase | Cofactor, Carrier, and Vitamin Biosynthesis | K01935 | H | Catalyzes a mechanistically unusual reaction, the ATP- dependent insertion of CO2 between the N7 and N8 nitrogen atoms of 7,8-diaminopelargonic acid (DAPA) to form an ureido ring | bioD | 53.7 | 0.98 | 4.61E-04 | 7.2 | 17.6 | 16.4 | 28.0 | 64.1 | 162.3 |
| stage3_free | stage3_shell | -2.72 | 5.56E-03 | V12B01_03113 | EAP92120.1 | dithiobiotin synthetase | Cofactor, Carrier, and Vitamin Biosynthesis | K01935 | H | Catalyzes a mechanistically unusual reaction, the ATP- dependent insertion of CO2 between the N7 and N8 nitrogen atoms of 7,8-diaminopelargonic acid (DAPA) to form an ureido ring | bioD | 53.7 | 1.19 | 9.25E-04 | 13.4 | 10.5 | NA | 28.0 | 64.1 | 162.3 |
| stage3_free | stage3_shell | 3.18 | 1.39E-05 | V12B01_03128 | EAP92123.1 | adenylosuccinate lyase | Nucleoside and Nucleotide Biosynthesis | K01756 | F | Belongs to the lyase 1 family. Adenylosuccinate lyase subfamily | purB | 136.6 | 0.72 | 6.95E-07 | 743.8 | 509.2 | NA | 9.8 | 24.1 | 87.2 |
| stage3_free | stage3_shell | 2.20 | 2.95E-04 | V12B01_03133 | EAP93326.1 | adenylosuccinate lyase | Nucleoside and Nucleotide Biosynthesis | K01756 | F | Belongs to the lyase 1 family. Adenylosuccinate lyase subfamily | purB | 44.6 | 0.59 | 2.73E-05 | 469.6 | 293.6 | NA | 94.2 | 8.1 | 53.4 |
| stage2 | stage3_free | -2.44 | 7.25E-05 | V12B01_03163 | EAP93332.1 | hypothetical protein | | K18218 | C | COG1757 Na H antiporter | - | 112.0 | 0.57 | 1.66E-06 | 25.3 | 22.8 | 23.6 | 155.6 | 153.2 | NA |
| stage2 | stage3_shell | 2.01 | 1.26E-03 | V12B01_03183 | EAP93336.1 | imidazole glycerol-phosphate dehydratase/histidinol phosphatase | Amino Acid Biosynthesis | K01089 | E | Histidine biosynthesis bifunctional protein HisB | hisB | 236.2 | 0.64 | 1.97E-04 | 481.0 | 138.2 | 160.5 | 11.7 | 32.0 | 70.6 |
| stage2 | stage3_shell | 2.54 | 1.50E-07 | V12B01_03188 | EAP93337.1 | imidazole glycerol phosphate synthase subunit HisH | Amino Acid Degradation | K02501 | E | IGPS catalyzes the conversion of PRFAR and glutamine to IGP, AICAR and glutamate. The HisH subunit provides the glutamine amidotransferase activity that produces the ammonia necessary to HisF for the synthesis of IGP and AICAR | hisH | 137.5 | 0.46 | 6.29E-09 | 435.3 | 172.8 | 213.6 | 28.6 | 41.8 | 23.1 |
| stage2 | stage3_shell | 2.01 | 2.21E-06 | V12B01_03193 | EAP93338.1 | 1-(5-phosphoribosyl)-5-[(5-phosphoribosylamino)methylideneamino] imidazole-4-carboxamide isomerase | Amino Acid Biosynthesis | K01814 | E | 1-(5-phosphoribosyl)-5- (5-phosphoribosylamino)methylideneamino imidazole-4-carboxamide isomerase | hisA | 157.5 | 0.41 | 1.32E-07 | 412.7 | 242.8 | 264.3 | 30.3 | 67.6 | 57.9 |
| stage3_free | stage3_shell | -2.58 | 3.98E-05 | V12B01_03268 | EAP93353.1 | NapE protein | | K02571 | C | COG4459 Periplasmic nitrate reductase system, NapE component | torE | 108.7 | 0.62 | 2.56E-06 | 87.5 | 102.4 | NA | 602.4 | 810.2 | 77.8 |
| stage3_free | stage3_shell | -2.50 | 1.57E-03 | V12B01_03283 | EAP93356.1 | hypothetical protein | | | S | Putative, 10TM heavy-metal exporter | VP1164 | 152.6 | 0.83 | 1.96E-04 | 2.3 | 6.3 | NA | 32.9 | 10.0 | 25.8 |

| | | | | | | | | | | | | | | | | | | | | |
|---|---|---|---|---|---|---|---|---|---|---|---|---|---|---|---|---|---|---|---|---|
| stage2 | stage3_shell | -2.11 | 1.14E-05 | V12B01_03328 | EAP93365.1 | psp operon transcriptional activator | Regulation | K03974 | K | COG1221 Transcriptional regulators containing an AAA-type ATPase domain and a DNA-binding domain | pspF | 65.9 | 0.46 | 8.93E-07 | 10.4 | 36.7 | 29.9 | 70.3 | 140.5 | 61.2 |
| stage3_free | stage3_shell | -2.48 | 7.63E-06 | V12B01_03328 | EAP93365.1 | psp operon transcriptional activator | Regulation | K03974 | K | COG1221 Transcriptional regulators containing an AAA-type ATPase domain and a DNA-binding domain | pspF | 65.9 | 0.54 | 3.46E-07 | 19.2 | 20.2 | NA | 70.3 | 140.5 | 61.2 |
| stage3_free | stage3_shell | 2.38 | 3.03E-05 | V12B01_03393 | EAP93378.1 | putative sodium-dependent transporter | Transport Proteins | K03308 | S | COG0733 Na -dependent transporters of the SNF family | - | 368.3 | 0.54 | 1.79E-06 | 653.2 | 542.4 | NA | 86.2 | 61.8 | 70.6 |
| stage2 | stage3_shell | 2.83 | 2.87E-02 | V12B01_03423 | EAP93384.1 | CDP-diacylglycerol--glycerol-3-phosphate 3-phosphatidyltransferase | Fatty Acid and Lipid Biosynthesis | | I | Belongs to the CDP-alcohol phosphatidyltransferase class-I family | - | 3.9 | 2.29 | 8.52E-03 | 7.2 | 13.1 | 12.6 | 0.0 | 0.0 | 0.0 |
| stage2 | stage3_shell | -3.21 | 3.01E-04 | V12B01_03458 | EAP92251.1 | hypothetical protein | | | - | #N/A | #N/A | 7.7 | 0.91 | 3.67E-05 | 9.9 | 13.1 | 9.2 | 153.9 | 134.5 | 0.0 |
| stage3_free | stage3_shell | -4.61 | 6.74E-04 | V12B01_03458 | EAP92251.1 | hypothetical protein | | | - | #N/A | #N/A | 7.7 | 1.32 | 7.14E-05 | 6.8 | 0.0 | NA | 153.9 | 134.5 | 0.0 |
| stage2 | stage3_shell | 2.15 | 5.09E-02 | V12B01_03468 | EAP92253.1 | DNA and RNA helicase | Replication | K11927 | JKL | Belongs to the DEAD box helicase family | - | 4.9 | 1.38 | 1.72E-02 | 11.5 | 5.7 | 4.0 | 1.3 | 0.0 | 0.0 |
| stage2 | stage3_free | -3.66 | 1.37E-04 | V12B01_03503 | EAP92260.1 | glycine betaine/L-proline transport ATP-binding protein ProV | Transport Proteins | K02000 | E | COG4175 ABC-type proline glycine betaine transport system, ATPase component | proV | 42.5 | 0.91 | 3.46E-06 | 14.2 | 4.2 | 1.6 | 108.2 | 116.9 | NA |
| stage2 | stage3_shell | -3.43 | 2.71E-05 | V12B01_03503 | EAP92260.1 | glycine betaine/L-proline transport ATP-binding protein ProV | Transport Proteins | K02000 | E | COG4175 ABC-type proline glycine betaine transport system, ATPase component | proV | 42.5 | 0.83 | 2.37E-06 | 14.2 | 4.2 | 1.6 | 158.7 | 15.3 | 28.8 |
| stage2 | stage3_free | -3.13 | 2.01E-06 | V12B01_03513 | EAP92262.1 | glycine betaine/L-proline transport system permease protein ProW | Transport Proteins | K02001 | P | COG4176 ABC-type proline glycine betaine transport system, permease component | proW | 26.9 | 0.62 | 2.71E-08 | 21.9 | 5.8 | 6.6 | 112.4 | 121.5 | NA |
| stage3_free | stage3_shell | 2.51 | 3.09E-04 | V12B01_03513 | EAP92262.1 | glycine betaine/L-proline transport system permease protein ProW | Transport Proteins | K02001 | P | COG4176 ABC-type proline glycine betaine transport system, permease component | proW | 26.9 | 0.68 | 2.87E-05 | 112.4 | 121.5 | NA | 16.1 | 8.3 | 13.0 |
| stage2 | stage3_free | -3.03 | 5.31E-09 | V12B01_03518 | EAP92263.1 | glycine betaine/L-proline-binding protein | Transport Proteins | K02002 | E | Substrate binding domain of ABC-type glycine betaine transport system | proX | 90.5 | 0.49 | 4.21E-11 | 45.0 | 21.8 | 24.1 | 325.8 | 228.7 | NA |
| stage3_free | stage3_shell | -2.19 | 1.54E-03 | V12B01_03553 | EAP92270.1 | BLUF | | | S | Sensors of blue-light using FAD | - | 128.7 | 0.71 | 1.92E-04 | 51.5 | 68.8 | NA | 435.1 | 316.1 | 26.7 |
| stage2 | stage3_shell | -2.61 | 9.74E-05 | V12B01_03558 | EAP92271.1 | hypothetical protein | | | - | - | - | 19.9 | 0.67 | 1.02E-05 | 6.9 | 38.7 | 32.5 | 166.4 | 110.3 | 148.3 |
| stage2 | stage3_shell | 2.16 | 2.25E-08 | V12B01_03563 | EAP92272.1 | hypothetical protein | | | - | - | - | 129.1 | 0.37 | 7.31E-10 | 560.5 | 574.5 | 364.8 | 67.5 | 63.9 | 106.9 |
| stage2 | stage3_shell | -2.60 | 3.65E-06 | V12B01_03603 | EAP92280.1 | 50S ribosomal protein L25 | Translation | | - | #N/A | #N/A | 77.5 | 0.55 | 2.28E-07 | 23.2 | 129.4 | 130.3 | 449.2 | 650.1 | 343.0 |
| stage3_free | stage3_shell | -3.60 | 5.28E-08 | V12B01_03603 | EAP92280.1 | 50S ribosomal protein L25 | Translation | | - | #N/A | #N/A | 77.5 | 0.64 | 1.01E-09 | 36.7 | 57.9 | NA | 449.2 | 650.1 | 343.0 |
| stage2 | stage3_shell | 2.49 | 2.07E-08 | V12B01_03608 | EAP92281.1 | ribosomal protein L25 | Translation | K02897 | J | This is one of the proteins that binds to the 5S RNA in the ribosome where it forms part of the central protuberance | rplY | 163.8 | 0.42 | 6.65E-10 | 1160.3 | 842.9 | 1079.3 | 211.9 | 86.8 | 71.4 |
| stage3_free | stage3_shell | 2.89 | 4.08E-09 | V12B01_03608 | EAP92281.1 | ribosomal protein L25 | Translation | K02897 | J | This is one of the proteins that binds to the 5S RNA in the ribosome where it forms part of the central protuberance | rplY | 163.8 | 0.46 | 6.11E-11 | 1477.0 | 1282.7 | NA | 211.9 | 86.8 | 71.4 |
| stage2 | stage3_shell | -2.31 | 4.46E-03 | V12B01_03623 | EAP91697.1 | Predicted enzyme related to lactoylglutathione lyase | | K06996 | S | enzyme related to lactoylglutathione lyase | - | 30.8 | 0.88 | 8.86E-04 | 12.8 | 30.5 | 79.1 | 321.1 | 318.5 | 0.0 |
| stage3_free | stage3_shell | -2.16 | 1.35E-02 | V12B01_03623 | EAP91697.1 | Predicted enzyme related to lactoylglutathione lyase | | K06996 | S | enzyme related to lactoylglutathione lyase | - | 30.8 | 1.06 | 2.89E-03 | 32.8 | 48.3 | NA | 321.1 | 318.5 | 0.0 |
| stage2 | stage3_shell | -2.98 | 2.22E-07 | V12B01_03693 | EAP92440.1 | putative outer membrane protein | Outer membrane | K03286 | M | COG2885 Outer membrane protein and related peptidoglycan-associated (lipo)proteins | - | 824.7 | 0.56 | 9.78E-09 | 40.7 | 181.7 | 162.7 | 624.0 | 1651.3 | 335.0 |
| stage3_free | stage3_shell | -3.12 | 1.77E-06 | V12B01_03693 | EAP92440.1 | putative outer membrane protein | Outer membrane | K03286 | M | COG2885 Outer membrane protein and related peptidoglycan-associated (lipo)proteins | - | 824.7 | 0.64 | 6.07E-08 | 68.8 | 158.6 | NA | 624.0 | 1651.3 | 335.0 |
| stage2 | stage3_shell | -2.11 | 1.42E-04 | V12B01_03698 | EAP92441.1 | Outer membrane protein | Outer membrane | K12543 | MU | COG1538 Outer membrane protein | - | 887.9 | 0.54 | 1.54E-05 | 18.4 | 45.8 | 34.8 | 128.2 | 219.2 | 26.6 |
| stage2 | stage3_shell | -2.91 | 1.21E-06 | V12B01_03703 | EAP92442.1 | putative biofilm-associated surface protein | Membrane | | M | Conserved repeat domain | - | 5727.7 | 0.59 | 6.66E-08 | 13.2 | 43.2 | 38.6 | 302.4 | 260.1 | 61.3 |
| stage3_free | stage3_shell | -2.89 | 1.96E-05 | V12B01_03703 | EAP92442.1 | putative biofilm-associated surface protein | Outer membrane | | M | Conserved repeat domain | - | 5727.7 | 0.67 | 1.02E-06 | 31.9 | 30.1 | NA | 302.4 | 260.1 | 61.3 |
| stage2 | stage3_shell | -2.13 | 1.73E-03 | V12B01_03723 | EAP92446.1 | ISPsy11, transposase OrfA | | K07483 | L | Evidence 2b Function of strongly homologous gene | - | 11.2 | 0.69 | 2.85E-04 | 3.3 | 24.7 | 13.2 | 91.4 | 45.2 | 28.3 |
| stage3_free | stage3_shell | -2.47 | 1.83E-03 | V12B01_03723 | EAP92446.1 | ISPsy11, transposase OrfA | | K07483 | L | Evidence 2b Function of strongly homologous gene | - | 11.2 | 0.82 | 2.38E-04 | 13.7 | 7.2 | NA | 91.4 | 45.2 | 28.3 |
| stage3_free | stage3_shell | 2.10 | 3.20E-03 | V12B01_03758 | EAP93225.1 | hypothetical protein | | | S | membrane | ybjE | 16.9 | 0.68 | 4.69E-04 | 36.9 | 46.1 | NA | 3.5 | 11.7 | 0.0 |
| stage2 | stage3_shell | -2.05 | 1.95E-04 | V12B01_03768 | EAP93227.1 | hypothetical protein | | K07343 | K | COG3070 Regulator of competence-specific genes | VV1446 | 537.7 | 0.54 | 2.23E-05 | 257.5 | 478.3 | 548.0 | 1120.1 | 1156.8 | 1971.5 |
| stage2 | stage3_shell | -2.39 | 2.56E-02 | V12B01_03778 | EAP93229.1 | response regulator | Regulation | | T | response regulator | - | 25.4 | 2.02 | 7.40E-03 | 10.7 | 9.2 | 13.8 | 28.9 | 18.2 | 268.7 |
| stage2 | stage3_shell | -3.27 | 2.62E-04 | V12B01_03783 | EAP93230.1 | signal transduction histidine kinase | Regulation | | T | COG0642 Signal transduction histidine kinase | VP1245 | 39.8 | 0.95 | 3.12E-05 | 4.3 | 5.6 | 5.0 | 9.0 | 18.7 | 101.1 |
| stage3_free | stage3_shell | -2.20 | 1.47E-02 | V12B01_03783 | EAP93230.1 | signal transduction histidine kinase | Regulation | | T | COG0642 Signal transduction histidine kinase | VP1245 | 39.8 | 1.17 | 3.21E-03 | 8.0 | 9.4 | NA | 9.0 | 18.7 | 101.1 |
| stage2 | stage3_shell | 2.27 | 2.05E-08 | V12B01_03793 | EAP93232.1 | phosphoserine aminotransferase | Amino Acid Biosynthesis | K00831 | E | Catalyzes the reversible conversion of 3-phosphohydroxypyruvate to phosphoserine and of 3-hydroxy-2-oxo-4-phosphonooxybutanoate to phosphohydroxythreonine | serC | 320.4 | 0.39 | 6.54E-10 | 429.6 | 271.1 | 335.4 | 70.0 | 30.4 | 46.8 |

| | | | | | | | | | | | | | | | | | | | | |
|---|---|---|---|---|---|---|---|---|---|---|---|---|---|---|---|---|---|---|---|---|
| stage3_free | stage3_shell | 2.07 | 5.02E-06 | V12B01_03793 | EAP93232.1 | phosphoserine aminotransferase | Amino Acid Biosynthesis | K00831 | E | Catalyzes the reversible conversion of 3-phosphohydroxypyruvate to phosphoserine and of 3-hydroxy-2-oxo-4-phosphonooxybutanoate to phosphohydroxythreonine | serC | 320.4 | 0.43 | 2.01E-07 | 347.9 | 270.2 | NA | 70.0 | 30.4 | 46.8 |
| stage2 | stage3_shell | 2.16 | 3.50E-06 | V12B01_03828 | EAP93239.1 | aspartate aminotransferase | TCA cycle | K00832 | E | COG1448 Aspartate tyrosine aromatic aminotransferase | aspC | 266.5 | 0.45 | 2.18E-07 | 448.8 | 226.9 | 228.5 | 29.5 | 33.0 | 71.7 |
| stage3_free | stage3_shell | 2.50 | 1.60E-06 | V12B01_03828 | EAP93239.1 | aspartate aminotransferase | TCA cycle | K00832 | E | COG1448 Aspartate tyrosine aromatic aminotransferase | aspC | 266.5 | 0.50 | 5.42E-08 | 416.6 | 367.0 | NA | 29.5 | 33.0 | 71.7 |
| stage2 | stage3_shell | 2.34 | 4.39E-06 | V12B01_03863 | EAP93246.1 | asparaginyl-tRNA synthetase | Other Biosynthesis | K01893 | J | COG0017 Aspartyl asparaginyl-tRNA synthetases | asnS | 405.3 | 0.50 | 2.81E-07 | 696.0 | 296.5 | 313.6 | 82.1 | 18.4 | 67.1 |
| stage3_free | stage3_shell | 2.60 | 5.87E-06 | V12B01_03863 | EAP93246.1 | asparaginyl-tRNA synthetase | Other Biosynthesis | K01893 | J | COG0017 Aspartyl asparaginyl-tRNA synthetases | asnS | 405.3 | 0.55 | 2.43E-07 | 557.2 | 519.8 | NA | 82.1 | 18.4 | 67.1 |
| stage2 | stage3_shell | -3.48 | 7.59E-06 | V12B01_03868 | EAP93247.1 | sulfate transporter | Transport Proteins | K03321 | P | COG0659 Sulfate permease and related transporters (MFS superfamily) | - | 12.3 | 0.78 | 5.51E-07 | 0.6 | 1.4 | 1.3 | 10.7 | 4.1 | 19.1 |
| stage3_free | stage3_shell | -3.19 | 3.50E-04 | V12B01_03868 | EAP93247.1 | sulfate transporter | Transport Proteins | K03321 | P | COG0659 Sulfate permease and related transporters (MFS superfamily) | - | 12.3 | 0.92 | 3.36E-05 | 1.2 | 1.3 | NA | 10.7 | 4.1 | 19.1 |
| stage3_free | stage3_shell | -2.48 | 9.17E-05 | V12B01_03908 | EAP93255.1 | MutT/nudix family protein | | | L | COG0494 NTP pyrophosphohydrolases including oxidative damage repair enzymes | nudL | 63.5 | 0.62 | 6.64E-06 | 6.1 | 21.6 | NA | 74.6 | 75.6 | 47.5 |
| stage3_free | stage3_shell | -2.52 | 3.63E-06 | V12B01_03913 | EAP93256.1 | putative guanylate cyclase-related protein | | | S | Haem-NO-binding | - | 248.2 | 0.53 | 1.35E-07 | 114.5 | 148.6 | NA | 541.4 | 1075.5 | 288.4 |
| stage2 | stage3_shell | -5.60 | 4.94E-27 | V12B01_03968 | EAP93267.1 | hypothetical protein | | | - | - | - | 5964.9 | 0.50 | 4.17E-30 | 430.3 | 1629.9 | 2849.0 | 56483.5 | 54239.5 | 68847.3 |
| stage3_free | stage3_shell | -4.71 | 1.64E-15 | V12B01_03968 | EAP93267.1 | hypothetical protein | | | - | - | - | 5964.9 | 0.57 | 4.09E-18 | 2543.0 | 3375.9 | NA | 56483.5 | 54239.5 | 68847.3 |
| stage3_free | stage3_shell | -5.00 | 9.78E-03 | V12B01_04068 | EAP94821.1 | hypothetical protein | | | - | #N/A | #N/A | 1.9 | 3.37 | 1.88E-03 | 0.0 | 0.0 | NA | 72.6 | 57.3 | 0.0 |
| stage3_free | stage3_shell | -2.08 | 3.05E-10 | V12B01_04073 | EAP94822.1 | Transcriptional regulator | Regulation | | K | COG1309 Transcriptional regulator | - | 489.9 | 0.31 | 3.42E-12 | 199.0 | 231.6 | NA | 575.3 | 1004.7 | 510.8 |
| stage2 | stage3_shell | 4.48 | 1.99E-03 | V12B01_04118 | EAP94831.1 | hypothetical protein | | | - | #N/A | #N/A | 6.5 | 2.21 | 3.36E-04 | 18.1 | 27.5 | 27.8 | 0.0 | 0.0 | 0.0 |
| stage3_free | stage3_shell | 4.90 | 1.40E-03 | V12B01_04118 | EAP94831.1 | hypothetical protein | | | - | #N/A | #N/A | 6.5 | 2.27 | 1.71E-04 | 38.1 | 25.1 | NA | 0.0 | 0.0 | 0.0 |
| stage2 | stage3_shell | -3.15 | 3.90E-03 | V12B01_04138 | EAP94835.1 | esterase | | | S | Alpha/beta hydrolase family | - | 5.6 | 1.20 | 7.52E-04 | 1.0 | 2.3 | 0.7 | 10.3 | 29.3 | 0.0 |
| stage2 | stage3_free | -3.94 | 1.24E-04 | V12B01_04178 | EAP94843.1 | putative membrane protein | Membrane | | S | Protein of unknown function (DUF3360) | - | 127.2 | 0.97 | 3.07E-06 | 13.3 | 8.2 | 20.7 | 354.9 | 236.5 | NA |
| stage3_free | stage3_shell | 2.25 | 1.40E-02 | V12B01_04178 | EAP94843.1 | putative membrane protein | Membrane | | S | Protein of unknown function (DUF3360) | - | 127.2 | 0.99 | 3.01E-03 | 354.9 | 236.5 | NA | 86.8 | 4.0 | 3.8 |
| stage2 | stage3_shell | -2.16 | 1.57E-02 | V12B01_04198 | EAP94847.1 | oxidoreductase, short chain dehydrogenase/reductase family | | K13938 | IQ | Enoyl-(Acyl carrier protein) reductase | - | 23.7 | 1.03 | 4.13E-03 | 1.9 | 3.0 | 1.3 | 8.9 | 21.1 | 0.0 |
| stage3_free | stage3_shell | -2.22 | 2.74E-02 | V12B01_04198 | EAP94847.1 | oxidoreductase, short chain dehydrogenase/reductase family | | K13938 | IQ | Enoyl-(Acyl carrier protein) reductase | - | 23.7 | 1.29 | 7.02E-03 | 0.6 | 2.7 | NA | 8.9 | 21.1 | 0.0 |
| stage3_free | stage3_shell | -3.56 | 2.02E-02 | V12B01_04203 | EAP94848.1 | GTP cyclohydrolase I | Cofactor, Carrier, and Vitamin Biosynthesis | K01495 | H | GTP cyclohydrolase I | - | 49.9 | 2.82 | 4.79E-03 | 0.0 | 0.0 | NA | 5.0 | 4.7 | 0.0 |
| stage2 | stage3_shell | -3.99 | 4.21E-20 | V12B01_04238 | EAP94855.1 | Cys/Met metabolism pyridoxal-phosphate-dependent enzyme | | K01740 | E | Cys Met metabolism pyridoxal-phosphate-dependent enzyme | - | 268.0 | 0.42 | 1.30E-22 | 4.9 | 19.7 | 17.9 | 198.4 | 162.0 | 163.4 |
| stage3_free | stage3_shell | -3.81 | 1.90E-14 | V12B01_04238 | EAP94855.1 | Cys/Met metabolism pyridoxal-phosphate-dependent enzyme | | K01740 | E | Cys Met metabolism pyridoxal-phosphate-dependent enzyme | - | 268.0 | 0.48 | 7.22E-17 | 13.4 | 18.8 | NA | 198.4 | 162.0 | 163.4 |
| stage2 | stage3_shell | 2.80 | 2.72E-02 | V12B01_04348 | EAP94877.1 | hypothetical protein | | K06933 | S | Belongs to the UPF0251 family | - | 4.8 | 2.19 | 7.98E-03 | 19.1 | 21.6 | 28.0 | 0.0 | 0.0 | 0.0 |
| stage2 | stage3_shell | -3.67 | 2.28E-14 | V12B01_04358 | EAP94879.1 | glutamate-gated potassium channel | Transport Proteins | K02030 | ET | COG0834 ABC-type amino acid transport signal transduction systems, periplasmic component domain | - | 127.3 | 0.46 | 2.02E-16 | 2.4 | 9.4 | 6.9 | 48.3 | 76.2 | 65.2 |
| stage3_free | stage3_shell | -3.48 | 4.34E-10 | V12B01_04358 | EAP94879.1 | glutamate-gated potassium channel | Transport Proteins | K02030 | ET | COG0834 ABC-type amino acid transport signal transduction systems, periplasmic component domain | - | 127.3 | 0.54 | 4.94E-12 | 6.3 | 7.9 | NA | 48.3 | 76.2 | 65.2 |
| stage2 | stage3_free | -3.09 | 1.49E-06 | V12B01_04413 | EAP94890.1 | major facilitator family transporter | Transport Proteins | | EGP | Uncharacterised MFS-type transporter YbfB | - | 14.7 | 0.60 | 1.90E-08 | 2.6 | 4.2 | 2.1 | 31.5 | 27.4 | NA |
| stage2 | stage3_shell | -2.22 | 4.06E-04 | V12B01_04413 | EAP94890.1 | major facilitator family transporter | Transport Proteins | | EGP | Uncharacterised MFS-type transporter YbfB | - | 14.7 | 0.63 | 5.14E-05 | 2.6 | 4.2 | 2.1 | 18.8 | 7.6 | 9.6 |
| stage2 | stage3_shell | -2.33 | 1.36E-08 | V12B01_04458 | EAP94899.1 | transcriptional regulators, LysR family | Regulation | | K | Transcriptional regulator | - | 412.7 | 0.39 | 4.10E-10 | 84.1 | 225.2 | 254.9 | 736.6 | 1099.8 | 437.5 |
| stage3_free | stage3_shell | -3.11 | 2.25E-11 | V12B01_04458 | EAP94899.1 | transcriptional regulators, LysR family | Regulation | | K | Transcriptional regulator | - | 412.7 | 0.45 | 2.04E-13 | 123.8 | 96.0 | NA | 736.6 | 1099.8 | 437.5 |
| stage2 | stage3_shell | 3.22 | 1.86E-02 | V12B01_04503 | EAP94908.1 | hypothetical protein | | K10915 | E | Aminotransferase class I and II | cqsA | 4.0 | 2.29 | 5.07E-03 | 9.0 | 6.9 | 3.2 | 0.0 | 0.0 | 0.0 |
| stage3_free | stage3_shell | 2.35 | 5.25E-02 | V12B01_04503 | EAP94908.1 | hypothetical protein | | K10915 | E | Aminotransferase class I and II | cqsA | 4.0 | 2.29 | 1.60E-02 | 8.1 | 1.6 | NA | 0.0 | 0.0 | 0.0 |
| stage3_free | stage3_shell | -2.04 | 1.05E-05 | V12B01_04543 | EAP94916.1 | integration host factor alpha subunit | | K04764 | K | This protein is one of the two subunits of integration host factor, a specific DNA-binding protein that functions in genetic recombination as well as in transcriptional and translational control | himA | 2146.0 | 0.44 | 5.00E-07 | 1170.3 | 2012.0 | NA | 4432.8 | 9357.4 | 2406.2 |
| stage3_free | stage3_shell | 4.22 | 1.53E-03 | V12B01_04593 | EAP91790.1 | 5'-phosphoribosylglycinamide transformylase | Nucleoside and Nucleotide Biosynthesis | K08289 | F | Involved in the de novo purine biosynthesis. Catalyzes the transfer of formate to 5-phospho-ribosyl-glycinamide (GAR), producing 5-phospho-ribosyl-N-formylglycinamide (FGAR). Formate is provided by PurU via hydrolysis of 10-formyl-tetrahydrofolate | purT | 10.6 | 1.33 | 1.89E-04 | 43.9 | 39.2 | NA | 2.7 | 0.0 | 0.0 |
| stage2 | stage3_shell | -2.00 | 1.66E-04 | V12B01_04598 | EAP91791.1 | cytidine deaminase | Nucleoside and Nucleotide Degradation | K01489 | F | This enzyme scavenges exogenous and endogenous cytidine and 2'-deoxycytidine for UMP synthesis | cdd | 28.9 | 0.53 | 1.85E-05 | 8.0 | 6.8 | 6.7 | 10.8 | 27.3 | 35.3 |
| stage3_free | stage3_shell | 2.25 | 1.07E-02 | V12B01_04618 | EAP94609.1 | exonuclease I | | K01141 | L | COG2925 Exonuclease I | sbcB | 13.7 | 0.91 | 2.13E-03 | 34.7 | 26.4 | NA | 9.1 | 0.0 | 0.0 |

| | | | | | | | | | | | | | | | | | | | | |
|---|---|---|---|---|---|---|---|---|---|---|---|---|---|---|---|---|---|---|---|---|
| stage2 | stage3_shell | 3.72 | 1.66E-02 | V12B01_04623 | EAP94610.1 | hypothetical protein | | | S | Domain of unknown function (DUF4402) | - | 3.1 | 2.64 | 4.43E-03 | 34.8 | 13.8 | 26.5 | 0.0 | 0.0 | 0.0 |
| stage2 | stage3_shell | 3.84 | 8.86E-10 | V12B01_04698 | EAP94625.1 | sodium/dicarboxylate symporter | Transport Proteins | K06956 | U | Belongs to the dicarboxylate amino acid cation symporter (DAACS) (TC 2.A.23) family | - | 420.6 | 0.61 | 1.91E-11 | 958.6 | 239.5 | 511.2 | 17.4 | 26.4 | 33.2 |
| stage3_free | stage3_shell | 3.41 | 1.14E-06 | V12B01_04698 | EAP94625.1 | sodium/dicarboxylate symporter | Transport Proteins | K06956 | U | Belongs to the dicarboxylate amino acid cation symporter (DAACS) (TC 2.A.23) family | - | 420.6 | 0.67 | 3.79E-08 | 632.8 | 256.6 | NA | 17.4 | 26.4 | 33.2 |
| stage3_free | stage3_shell | 2.19 | 1.17E-02 | V12B01_04748 | EAP94635.1 | vtamin B12-transporter permease | Transport Proteins | K06073 | H | Part of the ABC transporter complex BtuCDF involved in vitamin B12 import. Involved in the translocation of the substrate across the membrane | btuC | 10.6 | 0.88 | 2.43E-03 | 23.9 | 27.0 | NA | 3.2 | 4.6 | 0.0 |
| stage2 | stage3_shell | 3.27 | 9.14E-07 | V12B01_04783 | EAP94642.1 | uncharacterized protein conserved in bacteria | | | S | Lytic polysaccharide mono-oxygenase, cellulose-degrading | - | 73.5 | 0.67 | 4.86E-08 | 200.7 | 184.7 | 154.6 | 2.2 | 12.4 | 21.3 |
| **stage3_free** | **stage3_shell** | **-2.34** | **1.35E-02** | **V12B01_04808** | **EAP94647.1** | **hypothetical protein** | | | **-** | **-** | **-** | **14.2** | **1.19** | **2.88E-03** | **0.0** | **45.4** | **NA** | **257.4** | **117.7** | **47.5** |
| stage2 | stage3_free | 2.04 | 1.53E-07 | V12B01_04838 | EAP94653.1 | hypothetical protein | | | LO | Belongs to the peptidase S16 family | - | 168.3 | 0.37 | 1.70E-09 | 165.4 | 127.0 | 156.2 | 38.0 | 30.1 | NA |
| stage3_free | stage3_shell | 2.02 | 1.08E-02 | V12B01_04873 | EAP94660.1 | hypothetical protein | | | S | PFAM KAP P-loop domain protein | - | 17.9 | 0.80 | 2.17E-03 | 17.1 | 17.6 | NA | 2.4 | 1.1 | 4.2 |
| **stage2** | **stage3_shell** | **-2.09** | **4.56E-09** | **V12B01_04893** | **EAP94664.1** | **site-specific recombinase, phage integrase family** | | | **L** | **Phage Integrase Family** | **-** | **131.2** | **0.34** | **1.18E-10** | **32.8** | **42.4** | **30.9** | **101.9** | **178.9** | **71.4** |
| **stage3_free** | **stage3_shell** | **-2.15** | **1.21E-07** | **V12B01_04893** | **EAP94664.1** | **site-specific recombinase, phage integrase family** | | | **L** | **Phage Integrase Family** | **-** | **131.2** | **0.38** | **2.63E-09** | **29.8** | **37.5** | **NA** | **101.9** | **178.9** | **71.4** |
| **stage3_free** | **stage3_shell** | **-2.85** | **1.04E-04** | **V12B01_04898** | **EAP94665.1** | **hypothetical protein** | | | **-** | **#N/A** | **#N/A** | **43.6** | **0.74** | **7.82E-06** | **16.0** | **23.4** | **NA** | **133.9** | **141.6** | **105.5** |
| **stage3_free** | **stage3_shell** | **-2.25** | **6.02E-05** | **V12B01_04903** | **EAP94666.1** | **site-specific recombinase, phage integrase family domain protein** | | | **-** | **#N/A** | **#N/A** | **120.8** | **0.54** | **4.08E-06** | **9.0** | **20.7** | **NA** | **60.8** | **98.3** | **20.7** |
| **stage2** | **stage3_shell** | **-3.65** | **NA** | **V12B01_04973** | **EAP94680.1** | **peptidase, M20A family** | | **K01258** | **E** | **COG2195 Di- and tripeptidases** | **-** | **43.1** | **1.18** | **NA** | **2.8** | **5.1** | **3.4** | **11.5** | **16.4** | **113.2** |
| **stage2** | **stage3_free** | **-2.15** | **3.46E-02** | **V12B01_04983** | **EAP94682.1** | **hypothetical protein** | | | **-** | **-** | **-** | **81.5** | **1.17** | **3.30E-03** | **47.8** | **169.5** | **97.3** | **885.2** | **779.9** | **NA** |
| stage2 | stage3_shell | 2.78 | 2.29E-04 | V12B01_05068 | EAP94699.1 | hypothetical protein | | K07454 | L | restriction endonuclease | - | 26.2 | 0.75 | 2.70E-05 | 51.2 | 44.9 | 57.0 | 8.5 | 0.0 | 5.0 |
| **stage2** | **stage3_shell** | **-2.48** | **6.58E-07** | **V12B01_05083** | **EAP94702.1** | **hypothetical protein** | | | **-** | **-** | **-** | **240.0** | **0.48** | **3.34E-08** | **36.3** | **151.0** | **164.7** | **365.5** | **917.1** | **335.0** |
| **stage3_free** | **stage3_shell** | **-2.67** | **2.61E-06** | **V12B01_05083** | **EAP94702.1** | **hypothetical protein** | | | **-** | **-** | **-** | **240.0** | **0.55** | **9.44E-08** | **104.6** | **98.1** | **NA** | **365.5** | **917.1** | **335.0** |
| stage3_free | stage3_shell | 2.31 | 1.26E-05 | V12B01_05115 | EAP91836.1 | kinase, GHMP family | | K00919 | F | Catalyzes the phosphorylation of the position 2 hydroxy group of 4-diphosphocytidyl-2C-methyl-D-erythritol | ispE | 57.0 | 0.50 | 6.14E-07 | 160.3 | 238.0 | NA | 45.6 | 15.5 | 16.3 |
| stage3_free | stage3_shell | 2.22 | 2.45E-03 | V12B01_05135 | EAP91840.1 | HemK protein | | K02493 | J | Methylates the class 1 translation termination release factors RF1 PrfA and RF2 PrfB on the glutamine residue of the universally conserved GGQ motif | prmC | 27.6 | 0.76 | 3.44E-04 | 91.2 | 72.6 | NA | 1.8 | 5.2 | 26.1 |
| stage3_free | stage3_shell | 2.39 | 2.88E-02 | V12B01_05160 | EAP91845.1 | hypothetical protein | | K03976 | S | Belongs to the prolyl-tRNA editing family. YbaK EbsC subfamily | ybaK | 7.6 | 1.21 | 7.44E-03 | 35.7 | 30.4 | NA | 6.7 | 0.0 | 0.0 |
| **stage2** | **stage3_shell** | **-5.47** | **3.60E-04** | **V12B01_05175** | **EAP91848.1** | **hypothetical protein** | | | **S** | **Protein of unknown function (DUF3149)** | **-** | **2468.4** | **1.76** | **4.49E-05** | **198.7** | **1147.9** | **768.0** | **4917.9** | **13347.7** | **95210.2** |
| stage2 | stage3_shell | 2.22 | 2.65E-02 | V12B01_05245 | EAP93999.1 | 3-oxoacyl-(acyl carrier protein) synthase | Fatty Acid and Lipid Biosynthesis | K00647 | IQ | Belongs to the beta-ketoacyl-ACP synthases family | - | 7.5 | 1.05 | 7.73E-03 | 11.1 | 9.7 | 10.4 | 0.0 | 2.5 | 0.0 |
| **stage2** | **stage3_shell** | **-3.33** | **1.73E-18** | **V12B01_05305** | **EAP94011.1** | **putative phospholipid biosynthesis acyltransferase** | | | **I** | **Phosphate acyltransferases** | **-** | **58.4** | **0.36** | **6.95E-21** | **12.7** | **17.9** | **17.4** | **107.8** | **170.7** | **96.0** |
| **stage3_free** | **stage3_shell** | **-2.62** | **7.47E-10** | **V12B01_05305** | **EAP94011.1** | **putative phospholipid biosynthesis acyltransferase** | **Membrane** | | **I** | **Phosphate acyltransferases** | **-** | **58.4** | **0.41** | **9.61E-12** | **30.8** | **21.2** | **NA** | **107.8** | **170.7** | **96.0** |
| **stage3_free** | **stage3_shell** | **-2.18** | **7.16E-05** | **V12B01_05330** | **EAP94016.1** | **hypothetical protein** | | | **-** | **-** | **-** | **129.8** | **0.53** | **4.95E-06** | **28.0** | **37.9** | **NA** | **192.2** | **122.3** | **56.2** |
| stage2 | stage3_shell | 2.73 | 2.03E-06 | V12B01_05340 | EAP94018.1 | Predicted integral membrane protein | Membrane | | S | Integral membrane protein | VC2207 | 69.2 | 0.56 | 1.18E-07 | 220.9 | 119.2 | 133.5 | 20.1 | 9.5 | 17.9 |
| stage2 | stage3_shell | 2.19 | 1.86E-04 | V12B01_05345 | EAP94019.1 | hypothetical protein | | K09860 | S | LPP20 lipoprotein | rim | 58.2 | 0.58 | 2.11E-05 | 239.8 | 132.0 | 174.1 | 29.6 | 10.5 | 39.5 |
| stage2 | stage3_free | 1.99 | 2.89E-09 | V12B01_05390 | EAP94028.1 | flagellar hook protein | Motility | K02390 | N | Flagellar hook protein FlgE | flgE | 1460.1 | 0.32 | 2.02E-11 | 1349.7 | 1402.9 | 1568.6 | 362.0 | 327.2 | NA |
| stage2 | stage3_shell | 2.49 | 5.64E-06 | V12B01_05395 | EAP94029.1 | polar flagellar FlgF | Motility | K02391 | N | Belongs to the flagella basal body rod proteins family | flgF | 53.9 | 0.55 | 3.75E-07 | 142.3 | 107.2 | 99.0 | 4.3 | 8.1 | 34.2 |
| stage2 | stage3_shell | 3.50 | 2.39E-07 | V12B01_05400 | EAP94030.1 | Flagellar basal body rod protein FlgG | Motility | K02392 | N | Belongs to the flagella basal body rod proteins family | flgG | 47.2 | 0.64 | 1.06E-08 | 102.0 | 70.9 | 81.1 | 4.1 | 0.0 | 14.4 |
| stage3_free | stage3_shell | 2.52 | 3.24E-04 | V12B01_05400 | EAP94030.1 | Flagellar basal body rod protein FlgG | Motility | K02392 | N | Belongs to the flagella basal body rod proteins family | flgG | 47.2 | 0.67 | 3.04E-05 | 48.1 | 42.6 | NA | 4.1 | 0.0 | 14.4 |
| stage2 | stage3_shell | 2.45 | 2.96E-07 | V12B01_05405 | EAP94031.1 | flagellar L-ring protein precursor | Motility | K02393 | N | Assembles around the rod to form the L-ring and probably protects the motor basal body from shearing forces during rotation | flgH | 65.8 | 0.46 | 1.37E-08 | 116.7 | 83.4 | 111.6 | 12.3 | 7.8 | 22.0 |
| stage2 | stage3_shell | 4.08 | 7.21E-09 | V12B01_05410 | EAP94032.1 | flagellar P-ring protein precursor | Motility | K02394 | N | Assembles around the rod to form the L-ring and probably protects the motor basal body from shearing forces during rotation | flgI | 73.0 | 0.65 | 1.99E-10 | 68.8 | 91.5 | 96.6 | 5.9 | 2.8 | 0.0 |
| stage3_free | stage3_shell | 3.13 | 2.02E-05 | V12B01_05410 | EAP94032.1 | flagellar P-ring protein precursor | Motility | K02394 | N | Assembles around the rod to form the L-ring and probably protects the motor basal body from shearing forces during rotation | flgI | 73.0 | 0.68 | 1.06E-06 | 51.1 | 42.2 | NA | 5.9 | 2.8 | 0.0 |
| stage2 | stage3_shell | 2.36 | 6.23E-06 | V12B01_05415 | EAP94033.1 | peptidoglycan hydrolase | Motility | K02395 | N | Flagellar rod assembly protein muramidase FlgJ | flgJ | 71.0 | 0.51 | 4.19E-07 | 113.1 | 94.6 | 84.7 | 10.2 | 6.5 | 24.3 |
| stage2 | stage3_shell | 2.32 | 8.21E-08 | V12B01_05420 | EAP94034.1 | flagellar hook-associated protein | Motility | K02396 | N | flagellar hook-associated protein | flgK | 252.9 | 0.42 | 3.19E-09 | 171.6 | 193.8 | 216.5 | 27.2 | 15.3 | 36.3 |
| stage2 | stage3_shell | 3.12 | 5.13E-12 | V12B01_05425 | EAP94035.1 | flagellar hook-associated protein | Motility | K02397 | N | COG1344 Flagellin and related hook-associated proteins | flgL | 103.6 | 0.43 | 7.03E-14 | 134.8 | 116.3 | 139.9 | 8.0 | 10.2 | 14.3 |
| stage2 | stage3_free | 2.09 | 1.02E-03 | V12B01_05430 | EAP94036.1 | flagellin | Motility | K02406 | N | Flagellin is the subunit protein which polymerizes to form the filaments of bacterial flagella | flaA | 1356.3 | 0.63 | 4.09E-05 | 974.6 | 1744.1 | 1910.2 | 302.5 | 295.7 | NA |
| **stage3_free** | **stage3_shell** | **-2.46** | **9.31E-13** | **V12B01_05450** | **EAP94040.1** | **flagellar protein FlaG** | **Motility** | **K06603** | **N** | **flagellar protein FlaG** | **flaG** | **781.4** | **0.33** | **6.08E-15** | **472.4** | **620.2** | **NA** | **1492.7** | **3352.5** | **2005.1** |

| | | | | | | | | | | | | | | | | | | | | |
|---|---|---|---|---|---|---|---|---|---|---|---|---|---|---|---|---|---|---|---|---|
| **stage3_free** | **stage3_shell** | **-2.09** | **5.89E-10** | **V12B01_05455** | **EAP94041.1** | **flagellar hook-associated protein** | **Motility** | **K02407** | **N** | **morphogenesis and for the elongation of the flagellar filament by facilitating polymerization of the flagellin monomers at the tip of growing filament. Forms a capping structure, which prevents flagellin subunits (transported through the central channel of the flagellum) from leaking out without polymerization at the distal end** | **fliD** | **617.1** | **0.32** | **7.21E-12** | **94.0** | **104.2** | **NA** | **190.7** | **443.1** | **320.1** |
| stage3_free | stage3_shell | 2.04 | 1.16E-07 | V12B01_05675 | EAP94252.1 | long-chain fatty acid transport protein | Transport Proteins | K06076 | I | long-chain fatty acid transport protein | fadL | 433.3 | 0.36 | 2.48E-09 | 765.5 | 812.9 | NA | 151.6 | 152.0 | 90.8 |
| **stage3_free** | **stage3_shell** | **-2.21** | **2.23E-04** | **V12B01_05765** | **EAP94270.1** | **serine alkaline protease (subtilisin E)** | | | **M** | **Subtilase family** | **-** | **183.1** | **0.58** | **1.94E-05** | **7.6** | **3.4** | **NA** | **22.5** | **26.6** | **15.0** |
| stage2 | stage3_shell | 2.69 | 9.31E-03 | V12B01_05775 | EAP94272.1 | omega-3 polyunsaturated fatty acid synthase PfaD | Lipid biosynthesis | K15329 | S | 2-nitropropane dioxygenase | - | 14.6 | 1.16 | 2.13E-03 | 48.1 | 21.8 | 13.0 | 0.0 | 5.5 | 0.0 |
| stage2 | stage3_shell | 2.63 | 5.23E-05 | V12B01_05780 | EAP94273.1 | omega-3 polyunsaturated fatty acid synthase PfaC | Lipid biosynthesis | | IQ | FabA-like domain | - | 55.7 | 0.64 | 4.96E-06 | 35.2 | 25.8 | 18.8 | 2.9 | 4.0 | 1.0 |
| stage2 | stage3_free | 2.55 | 9.79E-04 | V12B01_05785 | EAP94274.1 | omega-3 polyunsaturated fatty acid synthase PfaB | Lipid biosynthesis | | Q | synthase | - | 23.2 | 0.77 | 3.91E-05 | 46.8 | 37.2 | 35.1 | 4.0 | 6.7 | NA |
| stage2 | stage3_shell | 4.47 | 1.14E-05 | V12B01_05785 | EAP94274.1 | omega-3 polyunsaturated fatty acid synthase PfaB | Lipid biosynthesis | | Q | synthase | - | 23.2 | 1.02 | 8.91E-07 | 46.8 | 37.2 | 35.1 | 0.0 | 0.0 | 4.1 |
| stage2 | stage3_shell | 2.14 | 6.08E-06 | V12B01_05790 | EAP94275.1 | omega-3 polyunsaturated fatty acid synthase PfaA | Lipid biosynthesis | | IQ | PKS_KR | - | 247.4 | 0.46 | 4.07E-07 | 88.1 | 57.7 | 49.7 | 4.5 | 8.5 | 15.9 |
| stage3_free | stage3_shell | 2.46 | 3.72E-06 | V12B01_05790 | EAP94275.1 | omega-3 polyunsaturated fatty acid synthase PfaA | | | IQ | PKS_KR | - | 247.4 | 0.51 | 1.40E-07 | 100.5 | 67.2 | NA | 4.5 | 8.5 | 15.9 |
| **stage2** | **stage3_free** | **-4.09** | **2.62E-13** | **V12B01_05805** | **EAP94278.1** | **C4-dicarboxylate-binding periplasmic protein** | **Transport Proteins** | **K11688** | **G** | **TRAP-type C4-dicarboxylate transport system, periplasmic component** | **dctP** | **544.6** | **0.53** | **7.87E-16** | **155.4** | **233.3** | **62.7** | **3449.0** | **2268.7** | **NA** |
| stage3_free | stage3_shell | 2.04 | 2.57E-04 | V12B01_05805 | EAP94278.1 | C4-dicarboxylate-binding periplasmic protein | | K11688 | G | TRAP-type C4-dicarboxylate transport system, periplasmic component | dctP | 544.6 | 0.53 | 2.29E-05 | 3449.0 | 2268.7 | NA | 622.4 | 477.0 | 254.4 |
| **stage2** | **stage3_free** | **-4.21** | **5.80E-12** | **V12B01_05810** | **EAP94279.1** | **putative C4-dicarboxylate transport protein DctQ** | **Transport Proteins** | **K11689** | **G** | **TRAP-type C4-dicarboxylate transport system, small permease component** | **-** | **195.8** | **0.58** | **2.17E-14** | **90.6** | **69.3** | **17.3** | **1561.4** | **914.3** | **NA** |
| **stage2** | **stage3_shell** | **-2.54** | **2.82E-06** | **V12B01_05810** | **EAP94279.1** | **putative C4-dicarboxylate transport protein DctQ** | **Transport Proteins** | **K11689** | **G** | **TRAP-type C4-dicarboxylate transport system, small permease component** | **-** | **195.8** | **0.53** | **1.73E-07** | **90.6** | **69.3** | **17.3** | **175.4** | **386.2** | **282.5** |
| **stage2** | **stage3_free** | **-2.97** | **1.09E-04** | **V12B01_05815** | **EAP94280.1** | **C4-dicarboxylate transport protein** | **Transport Proteins** | **K11690** | **G** | **TRAP-type C4-dicarboxylate transport system, large permease component** | **dctM** | **203.7** | **0.72** | **2.65E-06** | **53.9** | **74.9** | **17.0** | **575.2** | **388.3** | **NA** |
| **stage2** | **stage3_shell** | **-2.43** | **1.78E-04** | **V12B01_05815** | **EAP94280.1** | **C4-dicarboxylate transport protein** | **Transport Proteins** | **K11690** | **G** | **TRAP-type C4-dicarboxylate transport system, large permease component** | **dctM** | **203.7** | **0.65** | **2.00E-05** | **53.9** | **74.9** | **17.0** | **85.6** | **447.6** | **167.2** |
| **stage2** | **stage3_shell** | **-5.25** | **7.87E-22** | **V12B01_05820** | **EAP94281.1** | **hypothetical protein** | | | **-** | **-** | **-** | **768.6** | **0.53** | **1.70E-24** | **74.1** | **423.4** | **276.2** | **10238.5** | **7694.8** | **5416.3** |
| **stage3_free** | **stage3_shell** | **-5.39** | **8.44E-18** | **V12B01_05820** | **EAP94281.1** | **hypothetical protein** | | | **-** | **-** | **-** | **768.6** | **0.60** | **7.12E-21** | **210.3** | **251.2** | **NA** | **10238.5** | **7694.8** | **5416.3** |
| stage2 | stage3_shell | 3.57 | 1.32E-02 | V12B01_05925 | EAP94302.1 | hypothetical protein | | | - | - | - | 42.2 | 1.89 | 3.29E-03 | 52.9 | 118.1 | 620.4 | 0.0 | 18.0 | 0.0 |
| stage3_free | stage3_shell | 2.18 | 6.16E-02 | V12B01_05925 | EAP94302.1 | hypothetical protein | | | - | - | - | 42.2 | 2.24 | 1.96E-02 | 239.6 | 85.8 | NA | 0.0 | 18.0 | 0.0 |
| stage3_free | stage3_shell | 2.55 | 1.73E-02 | V12B01_05930 | EAP94303.1 | hypothetical membrane spanning protein | Membrane | | S | membrane | - | 10.5 | 1.23 | 3.93E-03 | 19.4 | 23.5 | NA | 0.0 | 0.0 | 6.2 |
| stage2 | stage3_shell | 2.70 | 2.68E-07 | V12B01_06021 | EAP91973.1 | ATPase of the AAA+ class | | | S | Protein of unknown function (DUF1422) | yijD | 75.9 | 0.52 | 1.21E-08 | 393.3 | 282.8 | 350.6 | 21.5 | 28.5 | 61.2 |
| stage3_free | stage3_shell | 2.58 | 6.67E-06 | V12B01_06021 | EAP91973.1 | ATPase of the AAA+ class | | | S | Protein of unknown function (DUF1422) | yijD | 75.9 | 0.55 | 2.89E-07 | 361.5 | 286.7 | NA | 21.5 | 28.5 | 61.2 |
| stage2 | stage3_shell | 2.07 | 4.40E-02 | V12B01_06041 | EAP91977.1 | DNA-damage-inducible protein F | | K03327 | V | Na driven multidrug efflux pump | dinF | 18.8 | 1.24 | 1.44E-02 | 8.9 | 11.0 | 9.2 | 0.0 | 2.2 | 0.0 |
| stage3_free | stage3_shell | 2.59 | 2.52E-02 | V12B01_06041 | EAP91977.1 | DNA-damage-inducible protein F | | K03327 | V | Na driven multidrug efflux pump | dinF | 18.8 | 1.32 | 6.28E-03 | 12.1 | 15.6 | NA | 0.0 | 2.2 | 0.0 |
| **stage3_free** | **stage3_shell** | **-2.59** | **8.46E-13** | **V12B01_06056** | **EAP91980.1** | **diacylglycerol kinase** | | **K00901** | **M** | **Recycling of diacylglycerol produced during the turnover of membrane phospholipid** | **dgkA** | **68.1** | **0.34** | **5.18E-15** | **52.9** | **50.3** | **NA** | **193.7** | **291.7** | **227.5** |
| **stage3_free** | **stage3_shell** | **-3.47** | **1.27E-07** | **V12B01_06146** | **EAP92712.1** | **hypothetical protein** | | | **-** | **-** | **VP2964** | **196.9** | **0.64** | **2.78E-09** | **101.2** | **90.9** | **NA** | **196.0** | **677.8** | **1579.3** |
| stage2 | stage3_shell | 2.10 | 1.31E-04 | V12B01_06221 | EAP92727.1 | GGDEF family protein | Regulation | K20964 | T | FIST_C | - | 96.6 | 0.53 | 1.40E-05 | 81.7 | 64.1 | 63.8 | 13.4 | 13.3 | 4.6 |
| stage2 | stage3_shell | 2.32 | 9.14E-03 | V12B01_06332 | EAP92609.1 | hypothetical protein | | K09125 | U | Involved in the import of queuosine (Q) precursors, required for Q precursor salvage | yhhQ | 15.6 | 0.95 | 2.08E-03 | 101.2 | 32.2 | 36.2 | 11.9 | 4.5 | 0.0 |
| **stage3_free** | **stage3_shell** | **-3.83** | **2.47E-06** | **V12B01_06387** | **EAP92620.1** | **Predicted integral membrane protein** | **Membrane** | | **S** | **Integral membrane protein** | **-** | **373.8** | **0.82** | **8.79E-08** | **180.5** | **237.4** | **NA** | **1971.2** | **5790.7** | **665.4** |
| **stage3_free** | **stage3_shell** | **-2.24** | **3.18E-05** | **V12B01_06392** | **EAP92621.1** | **hypothetical protein** | | **K09930** | **S** | **protein conserved in bacteria** | **-** | **67.7** | **0.52** | **1.92E-06** | **15.9** | **14.5** | **NA** | **46.4** | **110.7** | **19.8** |
| **stage3_free** | **stage3_shell** | **-2.41** | **5.39E-05** | **V12B01_06402** | **EAP92623.1** | **putative transmembrane protein** | **Membrane** | **K15977** | **S** | **DoxX** | **-** | **34.2** | **0.58** | **3.63E-06** | **6.0** | **22.0** | **NA** | **41.8** | **105.5** | **37.2** |
| stage2 | stage3_free | 2.15 | 1.06E-05 | V12B01_06501 | EAP92169.1 | acetyl-CoA acetyltransferase | Lipid biosynthesis | K00632 | I | Catalyzes the final step of fatty acid oxidation in which acetyl-CoA is released and the CoA ester of a fatty acid two carbons shorter is formed | fadA | 182.0 | 0.46 | 1.71E-07 | 222.7 | 173.7 | 113.7 | 29.9 | 39.6 | NA |
| **stage3_free** | **stage3_shell** | **-2.36** | **9.43E-07** | **V12B01_06501** | **EAP92169.1** | **acetyl-CoA acetyltransferase** | | **K00632** | **I** | **Catalyzes the final step of fatty acid oxidation in which acetyl-CoA is released and the CoA ester of a fatty acid two carbons shorter is formed** | **fadA** | **182.0** | **0.46** | **3.03E-08** | **29.9** | **39.6** | **NA** | **102.9** | **231.4** | **95.4** |
| stage3_free | stage3_shell | 2.01 | 3.23E-03 | V12B01_06536 | EAP92176.1 | glycyl-tRNA synthetase alpha subunit | Translation | K01878 | J | glycyl-trna synthetase alpha subunit | glyQ | 96.5 | 0.67 | 4.73E-04 | 268.5 | 253.1 | NA | 49.8 | 17.9 | 45.9 |
| **stage2** | **stage3_shell** | **-2.70** | **8.78E-17** | **V12B01_06681** | **EAP92400.1** | **hypothetical protein** | | | **S** | **SpoVR family** | **ycgB** | **708.5** | **0.31** | **5.19E-19** | **64.0** | **46.5** | **42.7** | **157.4** | **313.6** | **273.2** |
| **stage2** | **stage3_shell** | **-2.92** | **7.96E-08** | **V12B01_06686** | **EAP92401.1** | **hypothetical protein** | | **K09786** | **S** | **Belongs to the UPF0229 family** | **yeaH** | **441.0** | **0.53** | **3.07E-09** | **60.5** | **39.8** | **23.8** | **91.7** | **249.8** | **385.0** |
| **stage2** | **stage3_shell** | **-2.74** | **8.59E-07** | **V12B01_06691** | **EAP92402.1** | **hypothetical protein** | | **K07180** | **T** | **Serine protein kinase** | **prkA** | **1344.2** | **0.55** | **4.48E-08** | **132.6** | **69.9** | **44.7** | **157.7** | **395.6** | **732.8** |
| **stage2** | **stage3_shell** | **-2.15** | **NA** | **V12B01_06706** | **EAP92506.1** | **hypothetical protein** | | **K09161** | **S** | **Belongs to the UPF0304 family** | **yfbU** | **14.8** | **1.41** | **NA** | **3.6** | **2.8** | **3.7** | **12.8** | **18.1** | **34.1** |
| stage3_free | stage3_shell | 2.10 | 6.67E-03 | V12B01_06741 | EAP92513.1 | amino acid ABC transporter, permease protein | Transport Proteins | K10024 | P | COG4215 ABC-type arginine transport system, permease component | aotQ | 37.4 | 0.77 | 1.17E-03 | 111.7 | 58.3 | NA | 10.8 | 20.4 | 0.0 |

| | | | | | | | | | | | | | | | | | | | | |
|---|---|---|---|---|---|---|---|---|---|---|---|---|---|---|---|---|---|---|---|---|
| stage3_free | stage3_shell | 2.55 | 8.35E-09 | V12B01_06796 | EAP92524.1 | hypothetical protein | | | M | Forms passive diffusion pores that allow small molecular weight hydrophilic materials across the outer membrane | - | 2841.5 | 0.41 | 1.32E-10 | 9814.6 | 8948.7 | NA | 1303.8 | 520.3 | 1311.1 |
| **stage2** | stage3_shell | **-2.79** | **1.22E-14** | **V12B01_06816** | **EAP92528.1** | **ATP-dependent Clp protease adaptor protein ClpS** | **Protein degradation** | **K06891** | **S** | **Involved in the modulation of the specificity of the ClpAP-mediated ATP-dependent protein degradation** | **clpS** | **630.4** | **0.35** | **1.03E-16** | **433.6** | **738.1** | **665.9** | **1796.8** | **3704.4** | **3938.2** |
| **stage3_free** | stage3_shell | **-2.89** | **1.53E-12** | **V12B01_06816** | **EAP92528.1** | **ATP-dependent Clp protease adaptor protein ClpS** | **Protein degradation** | **K06891** | **S** | **Involved in the modulation of the specificity of the ClpAP-mediated ATP-dependent protein degradation** | **clpS** | **630.4** | **0.39** | **1.07E-14** | **541.4** | **592.6** | **NA** | **1796.8** | **3704.4** | **3938.2** |
| **stage2** | stage3_shell | **-2.10** | **6.34E-15** | **V12B01_06821** | **EAP92529.1** | **ATP-dependent Clp protease, ATP-binding subunit ClpA** | **Protein degradation** | **K03694** | **O** | **Belongs to the ClpA ClpB family** | **clpA** | **3092.4** | **0.26** | **5.05E-17** | **346.9** | **388.4** | **378.5** | **819.3** | **1709.1** | **1064.3** |
| **stage3_free** | stage3_shell | **-1.99** | **7.33E-11** | **V12B01_06821** | **EAP92529.1** | **ATP-dependent Clp protease, ATP-binding subunit ClpA** | **Protein degradation** | **K03694** | **O** | **Belongs to the ClpA ClpB family** | **clpA** | **3092.4** | **0.29** | **7.12E-13** | **404.5** | **394.6** | **NA** | **819.3** | **1709.1** | **1064.3** |
| **stage2** | stage3_shell | **-2.51** | **8.52E-11** | **V12B01_06841** | **EAP93911.1** | **putative outer membrane protein** | **Outer membrane** | **K06077** | **M** | **Glycine zipper 2TM domain** | **slyB** | **136.3** | **0.37** | **1.38E-12** | **60.5** | **105.6** | **106.8** | **227.7** | **437.9** | **518.2** |
| **stage3_free** | stage3_shell | **-2.43** | **3.72E-08** | **V12B01_06841** | **EAP93911.1** | **putative outer membrane protein** | **Outer membrane** | **K06077** | **M** | **Glycine zipper 2TM domain** | **slyB** | **136.3** | **0.42** | **6.73E-10** | **93.3** | **98.0** | **NA** | **227.7** | **437.9** | **518.2** |
| stage3_free | stage3_shell | 2.30 | 3.20E-05 | V12B01_06861 | EAP93915.1 | glucose-1-phosphate adenylyltransferase | Carbohydrate Biosynthesis | K00975 | H | Catalyzes the synthesis of ADP-glucose, a sugar donor used in elongation reactions on alpha-glucans | glgC | 296.5 | 0.53 | 1.94E-06 | 893.7 | 682.3 | NA | 77.4 | 49.7 | 166.0 |
| **stage2** | stage3_shell | **-4.28** | **4.63E-13** | **V12B01_06876** | **EAP93918.1** | **hypothetical protein** | | | **S** | **Putative prokaryotic signal transducing protein** | **-** | **107.0** | **0.58** | **5.27E-15** | **26.6** | **49.0** | **33.4** | **180.9** | **404.4** | **1011.7** |
| **stage3_free** | stage3_shell | **-3.80** | **1.66E-08** | **V12B01_06876** | **EAP93918.1** | **hypothetical protein** | | | **S** | **Putative prokaryotic signal transducing protein** | **-** | **107.0** | **0.66** | **2.83E-10** | **46.0** | **52.9** | **NA** | **180.9** | **404.4** | **1011.7** |
| **stage2** | stage3_shell | **-2.75** | **8.65E-09** | **V12B01_06881** | **EAP93919.1** | **hypothetical protein** | | | **S** | **SNARE associated Golgi protein** | **-** | **62.7** | **0.46** | **2.48E-10** | **13.1** | **16.6** | **7.8** | **61.0** | **64.4** | **75.3** |
| **stage2** | stage3_shell | **-2.11** | **1.43E-04** | **V12B01_06901** | **EAP93923.1** | **DNA-binding response regulator TorR** | **Regulation** | **K07772** | **K** | **COG0745 Response regulators consisting of a CheY-like receiver domain and a winged-helix DNA-binding domain** | **torR** | **520.7** | **0.55** | **1.56E-05** | **170.0** | **414.7** | **436.6** | **1324.7** | **1222.6** | **1132.5** |
| **stage3_free** | stage3_shell | **-2.17** | **5.45E-04** | **V12B01_06901** | **EAP93923.1** | **DNA-binding response regulator TorR** | **Regulation** | **K07772** | **K** | **COG0745 Response regulators consisting of a CheY-like receiver domain and a winged-helix DNA-binding domain** | **torR** | **520.7** | **0.63** | **5.58E-05** | **287.6** | **351.9** | **NA** | **1324.7** | **1222.6** | **1132.5** |
| **stage2** | stage3_shell | **-2.02** | **2.89E-04** | **V12B01_06931** | **EAP93929.1** | **anaerobic dehydrogenase** | **Fermentation** | **K00123** | **C** | **Belongs to the prokaryotic molybdopterin-containing oxidoreductase family** | **-** | **216.3** | **0.55** | **3.50E-05** | **12.4** | **30.0** | **21.2** | **24.6** | **65.1** | **115.8** |
| stage2 | stage3_shell | 2.23 | 2.86E-02 | V12B01_06946 | EAP93932.1 | uncharacterized protein conserved in archaea | | | O | protein conserved in archaea | - | 11.8 | 1.18 | 8.48E-03 | 52.8 | 15.3 | 11.8 | 6.5 | 0.0 | 0.0 |
| stage3_free | stage3_shell | 2.07 | 4.68E-02 | V12B01_06946 | EAP93932.1 | uncharacterized protein conserved in archaea | | | O | protein conserved in archaea | - | 11.8 | 1.26 | 1.37E-02 | 18.5 | 34.4 | NA | 6.5 | 0.0 | 0.0 |
| stage3_free | stage3_shell | 2.25 | 3.55E-03 | V12B01_06951 | EAP93933.1 | hypothetical protein | | K15257 | J | Catalyzes carboxymethyl transfer from carboxy-S- adenosyl-L-methionine (Cx-SAM) to 5-hydroxyuridine (ho5U) to form 5-carboxymethoxyuridine (cmo5U) at position 34 in tRNAs | cmoB | 14.3 | 0.77 | 5.40E-04 | 31.1 | 52.9 | NA | 3.3 | 6.2 | 5.9 |
| stage2 | stage3_shell | 2.20 | 5.50E-02 | V12B01_06956 | EAP93934.1 | hypothetical protein | | K15256 | H | Catalyzes the conversion of S-adenosyl-L-methionine (SAM) to carboxy-S-adenosyl-L-methionine (Cx-SAM) | cmoA | 5.7 | 2.62 | 1.89E-02 | 13.4 | 3.9 | 10.3 | 0.0 | 0.0 | 0.0 |
| stage3_free | stage3_shell | 2.04 | 6.80E-02 | V12B01_06956 | EAP93934.1 | hypothetical protein | | K15256 | H | Catalyzes the conversion of S-adenosyl-L-methionine (SAM) to carboxy-S-adenosyl-L-methionine (Cx-SAM) | cmoA | 5.7 | 2.74 | 2.23E-02 | 5.0 | 15.2 | NA | 0.0 | 0.0 | 0.0 |
| **stage3_free** | stage3_shell | **-2.09** | **4.45E-03** | **V12B01_06971** | **EAP93937.1** | **Holliday junction resolvase** | **Replication** | **K01159** | **L** | **Nuclease that resolves Holliday junction intermediates in genetic recombination. Cleaves the cruciform structure in supercoiled DNA by nicking to strands with the same polarity at sites symmetrically opposed at the junction in the homologous arms and leaves a 5'-terminal phosphate and a 3'-terminal hydroxyl group** | **ruvC** | **12.3** | **0.76** | **7.02E-04** | **3.5** | **10.1** | **NA** | **24.5** | **34.8** | **21.8** |
| stage2 | stage3_shell | 2.87 | 3.43E-04 | V12B01_06981 | EAP93939.1 | Holliday junction DNA helicase motor protein | Replication | K03550 | L | The RuvA-RuvB complex in the presence of ATP renatures cruciform structure in supercoiled DNA with palindromic sequence, indicating that it may promote strand exchange reactions in homologous recombination. RuvAB is a helicase that mediates the Holliday junction migration by localized denaturation and reannealing. RuvA stimulates, in the presence of DNA, the weak ATPase activity of RuvB | ruvA | 27.5 | 0.76 | 4.24E-05 | 63.9 | 66.6 | 48.4 | 7.8 | 4.9 | 0.0 |

| | | | | | | | | | | | | | | | | | | | | |
|---|---|---|---|---|---|---|---|---|---|---|---|---|---|---|---|---|---|---|---|---|
| stage3_free | stage3_shell | 3.20 | 1.56E-04 | V12B01_06981 | EAP93939.1 | Holliday junction DNA helicase motor protein | Replication | K03550 | L | The RuvA-RuvB complex in the presence of ATP renatures cruciform structure in supercoiled DNA with palindromic sequence, indicating that it may promote strand exchange reactions in homologous recombination. RuvAB is a helicase that mediates the Holliday junction migration by localized denaturation and reannealing. RuvA stimulates, in the presence of DNA, the weak ATPase activity of RuvB | ruvA | 27.5 | 0.79 | 1.26E-05 | 88.2 | 64.4 | NA | 7.8 | 4.9 | 0.0 |
| stage2 | stage3_free | -3.37 | 5.40E-15 | V12B01_06991 | EAP93941.1 | cytochrome d ubiquinol oxidase, subunit I | Aerobic Respiration | K00425 | C | COG1271 Cytochrome bd-type quinol oxidase, subunit 1 | cydA | 433.5 | 0.40 | 7.14E-18 | 69.8 | 46.0 | 50.1 | 594.5 | 642.3 | NA |
| stage2 | stage3_free | -2.49 | 1.98E-05 | V12B01_06996 | EAP93942.1 | Cytochrome d ubiquinol oxidase, subunit II | Aerobic Respiration | K00426 | C | COG1294 Cytochrome bd-type quinol oxidase, subunit 2 | - | 317.8 | 0.54 | 3.53E-07 | 71.2 | 70.7 | 77.1 | 605.4 | 357.4 | NA |
| stage2 | stage3_shell | 2.58 | 2.90E-09 | V12B01_07036 | EAP93950.1 | Peptidoglycan-associated lipoprotein | Membrane | K03640 | M | Belongs to the ompA family | pal | 553.2 | 0.42 | 6.90E-11 | 3005.4 | 1711.5 | 1963.7 | 126.5 | 312.0 | 309.4 |
| stage2 | stage3_shell | 2.50 | 3.18E-09 | V12B01_07046 | EAP93952.1 | hypothetical protein | | | D | Mediates coordination of peptidoglycan synthesis and outer membrane constriction during cell division | cpoB | 150.8 | 0.40 | 7.79E-11 | 621.1 | 382.9 | 320.4 | 30.4 | 90.1 | 39.7 |
| stage2 | stage3_free | -3.97 | 3.01E-08 | V12B01_07086 | EAP93960.1 | D-amino acid dehydrogenase small subunit | Membrane biosynthesis | K00285 | C | Oxidative deamination of D-amino acids | dadA | 204.2 | 0.67 | 2.72E-10 | 33.3 | 31.4 | 54.7 | 804.2 | 650.8 | NA |
| stage3_free | stage3_shell | 2.56 | 2.58E-04 | V12B01_07086 | EAP93960.1 | D-amino acid dehydrogenase small subunit | | K00285 | C | Oxidative deamination of D-amino acids | dadA | 204.2 | 0.68 | 2.31E-05 | 804.2 | 650.8 | NA | 133.8 | 12.1 | 72.7 |
| stage2 | stage3_shell | -3.14 | 5.07E-12 | V12B01_07116 | EAP93966.1 | hypothetical protein | | | M | 4-amino-4-deoxy-L-arabinose transferase and related glycosyltransferases of PMT family | - | 151.4 | 0.44 | 6.85E-14 | 9.3 | 25.0 | 27.3 | 104.2 | 171.8 | 145.4 |
| stage3_free | stage3_shell | -4.21 | 3.77E-15 | V12B01_07116 | EAP93966.1 | hypothetical protein | | | M | 4-amino-4-deoxy-L-arabinose transferase and related glycosyltransferases of PMT family | - | 151.4 | 0.51 | 1.19E-17 | 7.4 | 12.3 | NA | 104.2 | 171.8 | 145.4 |
| stage2 | stage3_shell | -2.20 | 3.76E-05 | V12B01_07121 | EAP93967.1 | hypothetical protein | | | S | GtrA-like protein | - | 77.7 | 0.52 | 3.41E-06 | 14.3 | 72.4 | 59.1 | 137.4 | 325.3 | 104.1 |
| stage3_free | stage3_shell | -2.73 | 1.05E-05 | V12B01_07121 | EAP93967.1 | hypothetical protein | | | S | GtrA-like protein | - | 77.7 | 0.61 | 5.02E-07 | 24.6 | 42.4 | NA | 137.4 | 325.3 | 104.1 |
| stage2 | stage3_shell | -3.47 | 2.05E-18 | V12B01_07136 | EAP93970.1 | hypothetical protein | | | S | Domain of unknown function (DUF4250) | VC1707 | 840.4 | 0.38 | 8.63E-21 | 265.5 | 588.5 | 641.2 | 3473.7 | 7007.4 | 2563.9 |
| stage3_free | stage3_shell | -2.85 | 2.52E-10 | V12B01_07136 | EAP93970.1 | hypothetical protein | | | S | Domain of unknown function (DUF4250) | VC1707 | 840.4 | 0.43 | 2.76E-12 | 598.4 | 910.5 | NA | 3473.7 | 7007.4 | 2563.9 |
| stage2 | stage3_shell | 3.72 | 1.20E-02 | V12B01_07141 | EAP93971.1 | transcriptional activator MetR | Regulation | K03576 | K | COG0583 Transcriptional regulator | metR | 7.0 | 2.40 | 2.89E-03 | 7.8 | 2.3 | 24.3 | 0.0 | 0.0 | 0.0 |
| stage3_free | stage3_shell | 2.39 | 5.12E-02 | V12B01_07141 | EAP93971.1 | transcriptional activator MetR | Regulation | K03576 | K | COG0583 Transcriptional regulator | metR | 7.0 | 2.45 | 1.54E-02 | 7.5 | 7.4 | NA | 0.0 | 0.0 | 0.0 |
| stage3_free | stage3_shell | 3.76 | 1.54E-02 | V12B01_07176 | EAP93978.1 | hypothetical protein | | K03566 | K | Bacterial regulatory helix-turn-helix protein, lysR family | - | 6.2 | 2.49 | 3.40E-03 | 15.5 | 10.8 | NA | 0.0 | 0.0 | 0.0 |
| stage2 | stage3_shell | 3.79 | 4.79E-08 | V12B01_07191 | EAP93981.1 | putative dicarboxylate-binding periplasmic protein | | | G | TRAP-type C4-dicarboxylate transport system, periplasmic component | - | 432.5 | 0.68 | 1.72E-09 | 3362.1 | 3198.2 | 2246.0 | 33.8 | 154.8 | 200.9 |
| stage3_free | stage3_shell | 2.21 | 3.42E-03 | V12B01_07191 | EAP93981.1 | putative dicarboxylate-binding periplasmic protein | | | G | TRAP-type C4-dicarboxylate transport system, periplasmic component | - | 432.5 | 0.76 | 5.08E-04 | 1355.1 | 856.5 | NA | 33.8 | 154.8 | 200.9 |
| stage2 | stage3_shell | 3.46 | 5.76E-09 | V12B01_07196 | EAP93982.1 | DctP protein | Transport Proteins | | G | TRAP-type C4-dicarboxylate transport system, periplasmic component | - | 247.5 | 0.58 | 1.58E-10 | 3490.7 | 3011.1 | 2483.1 | 58.6 | 194.3 | 269.9 |
| stage2 | stage3_shell | 2.70 | 8.50E-04 | V12B01_07201 | EAP93983.1 | C4-dicarboxylate transport system (permease small protein) | Transport Proteins | | G | TRAP-type C4-dicarboxylate transport system, small permease component | - | 26.6 | 0.85 | 1.22E-04 | 293.7 | 197.0 | 125.5 | 0.0 | 25.2 | 29.7 |
| stage2 | stage3_shell | 2.33 | 1.04E-04 | V12B01_07206 | EAP93984.1 | hypothetical protein | | | G | TRAP-type C4-dicarboxylate transport system, large permease component | - | 28.1 | 0.58 | 1.11E-05 | 77.2 | 75.3 | 37.5 | 12.2 | 6.9 | 4.4 |
| stage2 | stage3_free | -4.48 | 1.80E-09 | V12B01_07211 | EAP93985.1 | proton/glutamate symporter | Transport Proteins | | U | Belongs to the dicarboxylate amino acid cation symporter (DAACS) (TC 2.A.23) family | gltP | 66.1 | 0.69 | 1.07E-11 | 2.5 | 9.0 | 10.4 | 171.0 | 211.2 | NA |
| stage2 | stage3_shell | -2.81 | 2.18E-05 | V12B01_07211 | EAP93985.1 | proton/glutamate symporter | Transport Proteins | | U | Belongs to the dicarboxylate amino acid cation symporter (DAACS) (TC 2.A.23) family | gltP | 66.1 | 0.66 | 1.87E-06 | 2.5 | 9.0 | 10.4 | 60.9 | 15.6 | 54.2 |
| stage2 | stage3_shell | 2.99 | 1.34E-02 | V12B01_07278 | EAP94158.1 | sulfate adenylyltransferase subunit 2 | | K00957 | H | Sulfate adenylyltransferase subunit 2 | cysD | 287.6 | 1.91 | 3.37E-03 | 341.5 | 133.3 | 883.3 | 1.8 | 0.0 | 43.9 |
| stage3_free | stage3_shell | 3.32 | 1.71E-02 | V12B01_07278 | EAP94158.1 | sulfate adenylyltransferase subunit 2 | | K00957 | H | Sulfate adenylyltransferase subunit 2 | cysD | 287.6 | 1.97 | 3.86E-03 | 889.5 | 321.8 | NA | 1.8 | 0.0 | 43.9 |
| stage2 | stage3_shell | 3.93 | 4.53E-06 | V12B01_07283 | EAP94159.1 | sulfate adenylyltransferase subunit 1 | | K00956 | H | Belongs to the TRAFAC class translation factor GTPase superfamily. Classic translation factor GTPase family. CysN NodQ subfamily | cysN | 340.3 | 0.87 | 2.90E-07 | 337.5 | 76.7 | 526.4 | 8.9 | 6.3 | 19.9 |
| stage3_free | stage3_shell | 3.92 | 4.14E-05 | V12B01_07283 | EAP94159.1 | sulfate adenylyltransferase subunit 1 | | K00956 | H | Belongs to the TRAFAC class translation factor GTPase superfamily. Classic translation factor GTPase family. CysN NodQ subfamily | cysN | 340.3 | 0.95 | 2.67E-06 | 528.3 | 123.9 | NA | 8.9 | 6.3 | 19.9 |
| stage2 | stage3_shell | 4.02 | 1.18E-03 | V12B01_07288 | EAP94160.1 | putative sodium/sulfate symporter | Transport Proteins | | P | COG0471 Di- and tricarboxylate transporters | VP0295 | 59.8 | 1.40 | 1.83E-04 | 28.1 | 14.6 | 57.3 | 0.0 | 0.0 | 3.3 |
| stage3_free | stage3_shell | 4.05 | 2.23E-03 | V12B01_07288 | EAP94160.1 | putative sodium/sulfate symporter | Transport Proteins | | P | COG0471 Di- and tricarboxylate transporters | VP0295 | 59.8 | 1.48 | 3.07E-04 | 59.9 | 12.5 | NA | 0.0 | 0.0 | 3.3 |
| stage2 | stage3_shell | 3.63 | 9.40E-03 | V12B01_07293 | EAP94161.1 | adenylylsulfate kinase | | K00860 | F | Catalyzes the synthesis of activated sulfate | cysC | 17.3 | 1.62 | 2.16E-03 | 35.1 | 10.3 | 61.8 | 0.0 | 2.4 | 0.0 |
| stage2 | stage3_shell | 2.33 | 3.14E-03 | V12B01_07308 | EAP94164.1 | hypothetical protein | | K09950 | S | protein conserved in bacteria | - | 21.7 | 0.77 | 5.89E-04 | 69.8 | 48.7 | 32.7 | 6.2 | 8.8 | 0.0 |
| stage3_free | stage3_shell | 2.41 | 3.94E-03 | V12B01_07308 | EAP94164.1 | hypothetical protein | | K09950 | S | protein conserved in bacteria | - | 21.7 | 0.81 | 6.11E-04 | 51.4 | 57.7 | NA | 6.2 | 8.8 | 0.0 |
| stage3_free | stage3_shell | -2.14 | 3.87E-06 | V12B01_07358 | EAP94174.1 | hypothetical protein | | | S | Gamma-glutamyl cyclotransferase, AIG2-like | - | 97.3 | 0.44 | 1.47E-07 | 89.7 | 127.8 | NA | 192.4 | 487.3 | 446.6 |

| | | | | | | | | | | | | | | | | | | | | |
|---|---|---|---|---|---|---|---|---|---|---|---|---|---|---|---|---|---|---|---|---|
| stage2 | stage3_shell | -2.18 | 4.56E-08 | V12B01_07363 | EAP94175.1 | hypothetical protein | | | S | Protein of unknown function (DUF2799) | - | 357.6 | 0.38 | 1.60E-09 | 164.4 | 325.4 | 347.0 | 608.7 | 1466.3 | 893.2 |
| stage2 | stage3_shell | 3.27 | 1.18E-11 | V12B01_07518 | EAP94206.1 | hypothetical protein | | K09022 | J | translation initiation inhibitor, yjgF family | yjgF | 87.7 | 0.45 | 1.76E-13 | 556.4 | 448.6 | 470.5 | 57.4 | 38.8 | 0.0 |
| stage3_free | stage3_shell | 2.65 | 4.65E-07 | V12B01_07518 | EAP94206.1 | hypothetical protein | | K09022 | J | translation initiation inhibitor, yjgF family | yjgF | 87.7 | 0.49 | 1.27E-08 | 324.8 | 337.5 | NA | 57.4 | 38.8 | 0.0 |
| stage2 | stage3_free | -2.78 | 3.61E-04 | V12B01_07523 | EAP94207.1 | aspartate carbamoyltransferase regulatory subunit | Nucleoside and Nucleotide Biosynthesis | K00610 | F | Involved in allosteric regulation of aspartate carbamoyltransferase | pyrI | 15.2 | 0.74 | 1.12E-05 | 30.8 | 10.7 | 24.3 | 225.7 | 162.2 | NA |
| stage2 | stage3_shell | 2.33 | 3.40E-02 | V12B01_07523 | EAP94207.1 | aspartate carbamoyltransferase regulatory subunit | Nucleoside and Nucleotide Biosynthesis | K00610 | F | Involved in allosteric regulation of aspartate carbamoyltransferase | pyrI | 15.2 | 1.38 | 1.05E-02 | 30.8 | 10.7 | 24.3 | 0.0 | 0.0 | 6.2 |
| stage3_free | stage3_shell | 5.99 | 2.79E-05 | V12B01_07523 | EAP94207.1 | aspartate carbamoyltransferase regulatory subunit | Nucleoside and Nucleotide Biosynthesis | K00610 | F | Involved in allosteric regulation of aspartate carbamoyltransferase | pyrI | 15.2 | 1.45 | 1.60E-06 | 225.7 | 162.2 | NA | 0.0 | 0.0 | 6.2 |
| stage2 | stage3_free | -2.74 | 3.94E-04 | V12B01_07528 | EAP94208.1 | aspartate carbamoyltransferase catalytic subunit | Nucleoside and Nucleotide Biosynthesis | K00609 | F | Belongs to the ATCase OTCase family | pyrB | 52.9 | 0.73 | 1.24E-05 | 45.4 | 31.6 | 33.2 | 285.2 | 346.6 | NA |
| stage3_free | stage3_shell | 3.42 | 1.89E-05 | V12B01_07528 | EAP94208.1 | aspartate carbamoyltransferase catalytic subunit | Nucleoside and Nucleotide Biosynthesis | K00609 | F | Belongs to the ATCase OTCase family | pyrB | 52.9 | 0.79 | 9.82E-07 | 285.2 | 346.6 | NA | 6.9 | 3.3 | 42.9 |
| stage3_free | stage3_shell | 3.24 | 5.14E-07 | V12B01_07533 | EAP94209.1 | ornithine carbamoyltransferase | Amino Acid Degradation | K00611 | E | Reversibly catalyzes the transfer of the carbamoyl group from carbamoyl phosphate (CP) to the N(epsilon) atom of ornithine (ORN) to produce L-citrulline | argF | 433.4 | 0.62 | 1.45E-08 | 1166.4 | 604.6 | NA | 41.3 | 28.6 | 102.0 |
| stage2 | stage3_free | -2.49 | 1.85E-09 | V12B01_07538 | EAP94210.1 | arginine deiminase | Amino Acid Degradation | K01478 | E | COG2235 Arginine deiminase | arcA | 92.2 | 0.39 | 1.14E-11 | 13.1 | 13.1 | 9.7 | 71.2 | 74.4 | NA |
| stage2 | stage3_shell | -2.23 | 2.02E-08 | V12B01_07538 | EAP94210.1 | arginine deiminase | Amino Acid Degradation | K01478 | E | COG2235 Arginine deiminase | arcA | 92.2 | 0.38 | 6.39E-10 | 13.1 | 13.1 | 9.7 | 68.0 | 43.4 | 18.7 |
| stage2 | stage3_shell | 2.41 | 4.18E-03 | V12B01_07613 | EAP94225.1 | hypothetical protein | | | S | membrane protein domain | VP2641 | 15.0 | 0.82 | 8.22E-04 | 52.2 | 43.1 | 48.4 | 13.4 | 0.0 | 0.0 |
| stage3_free | stage3_shell | 2.27 | 9.35E-03 | V12B01_07613 | EAP94225.1 | hypothetical protein | | | S | membrane protein domain | VP2641 | 15.0 | 0.85 | 1.78E-03 | 34.5 | 56.4 | NA | 13.4 | 0.0 | 0.0 |
| stage2 | stage3_shell | -3.05 | 2.56E-07 | V12B01_07638 | EAP94230.1 | hypothetical protein | | | S | Protein of unknown function (DUF2787) | - | 16.3 | 0.57 | 1.15E-08 | 6.2 | 5.8 | 12.4 | 41.0 | 88.2 | 39.8 |
| stage3_free | stage3_shell | -2.92 | 2.36E-05 | V12B01_07638 | EAP94230.1 | hypothetical protein | | | S | Protein of unknown function (DUF2787) | - | 16.3 | 0.67 | 1.30E-06 | 10.7 | 6.7 | NA | 41.0 | 88.2 | 39.8 |
| stage2 | stage3_shell | -2.25 | 2.39E-07 | V12B01_07755 | EAP92215.1 | hypothetical protein | | K09131 | S | Belongs to the UPF0235 family | yggU | 215.2 | 0.42 | 1.07E-08 | 116.1 | 354.2 | 328.1 | 746.7 | 1466.1 | 802.3 |
| stage3_free | stage3_shell | -3.08 | 6.08E-10 | V12B01_07755 | EAP92215.1 | hypothetical protein | | K09131 | S | Belongs to the UPF0235 family | yggU | 215.2 | 0.48 | 7.56E-12 | 110.0 | 189.8 | NA | 746.7 | 1466.1 | 802.3 |
| stage3_free | stage3_shell | -2.93 | 1.93E-08 | V12B01_07785 | EAP92221.1 | Holliday junction resolvase-like protein | Replication | K07447 | L | Could be a nuclease involved in processing of the 5'-end of pre-16S rRNA | yqgF | 68.3 | 0.50 | 3.33E-10 | 21.6 | 22.7 | NA | 68.0 | 243.2 | 94.2 |
| stage2 | stage3_shell | -3.59 | 9.53E-12 | V12B01_07815 | EAP92227.1 | putative cytochrome c oxidase, subunit I | Aerobic Respiration | | S | Predicted integral membrane protein (DUF2189) | - | 335.1 | 0.51 | 1.41E-13 | 18.4 | 49.3 | 16.8 | 214.2 | 474.0 | 146.3 |
| stage3_free | stage3_shell | -3.16 | 1.34E-07 | V12B01_07815 | EAP92227.1 | putative cytochrome c oxidase, subunit I | Aerobic Respiration | | S | Predicted integral membrane protein (DUF2189) | - | 335.1 | 0.58 | 2.96E-09 | 32.7 | 41.5 | NA | 214.2 | 474.0 | 146.3 |
| stage3_free | stage3_shell | 2.53 | 9.56E-06 | V12B01_07825 | EAP92229.1 | transketolase | Pentose Phosphate Pathway | K00615 | G | Catalyzes the transfer of a two-carbon ketol group from a ketose donor to an aldose acceptor, via a covalent intermediate with the cofactor thiamine pyrophosphate | tktA | 392.2 | 0.54 | 4.46E-07 | 670.3 | 666.2 | NA | 97.7 | 33.4 | 85.6 |
| stage2 | stage3_shell | 3.39 | 1.39E-06 | V12B01_07865 | EAP93613.1 | Phosphoglycerate dehydrogenase | Amino Acid Biosynthesis | K00058 | C | Belongs to the D-isomer specific 2-hydroxyacid dehydrogenase family | serA | 190.9 | 0.71 | 7.85E-08 | 429.8 | 127.6 | 142.3 | 2.6 | 13.5 | 25.5 |
| stage3_free | stage3_shell | 3.65 | 3.68E-06 | V12B01_07865 | EAP93613.1 | Phosphoglycerate dehydrogenase | Amino Acid Biosynthesis | K00058 | C | Belongs to the D-isomer specific 2-hydroxyacid dehydrogenase family | serA | 190.9 | 0.77 | 1.37E-07 | 381.5 | 194.1 | NA | 2.6 | 13.5 | 25.5 |
| stage2 | stage3_shell | 2.02 | 6.91E-07 | V12B01_07870 | EAP93614.1 | ribose-5-phosphate isomerase A | Pentose Phosphate Pathway | K01807 | G | Catalyzes the reversible conversion of ribose-5- phosphate to ribulose 5-phosphate | rpiA | 69.4 | 0.39 | 3.53E-08 | 215.9 | 184.9 | 145.8 | 36.5 | 23.0 | 34.7 |
| stage3_free | stage3_shell | 2.78 | 2.40E-10 | V12B01_07870 | EAP93614.1 | ribose-5-phosphate isomerase A | Pentose Phosphate Pathway | K01807 | G | Catalyzes the reversible conversion of ribose-5- phosphate to ribulose 5-phosphate | rpiA | 69.4 | 0.42 | 2.58E-12 | 320.2 | 301.2 | NA | 36.5 | 23.0 | 34.7 |
| stage3_free | stage3_shell | -2.27 | 6.15E-15 | V12B01_07970 | EAP93634.1 | GTP-binding protein Era | | K03595 | S | An essential GTPase that binds both GDP and GTP, with rapid nucleotide exchange. Plays a role in 16S rRNA processing and 30S ribosomal subunit biogenesis and possibly also in cell cycle regulation and energy metabolism | era | 471.1 | 0.28 | 2.08E-17 | 184.4 | 147.8 | NA | 555.6 | 765.8 | 486.3 |
| stage3_free | stage3_shell | -2.20 | 5.57E-04 | V12B01_08005 | EAP93641.1 | MazG protein | | K04765 | S | Nucleoside triphosphate | mazG | 165.8 | 0.64 | 5.73E-05 | 65.6 | 102.4 | NA | 573.7 | 422.7 | 59.3 |
| stage3_free | stage3_shell | -2.17 | 2.21E-02 | V12B01_08085 | EAP93657.1 | ribonuclease H | | K03469 | L | RNase H | - | 8.2 | 1.17 | 5.39E-03 | 8.3 | 0.0 | NA | 21.7 | 34.3 | 12.9 |
| stage3_free | stage3_shell | 2.06 | 7.28E-10 | V12B01_08195 | EAP93679.1 | alanyl-tRNA synthetase | Translation | K01872 | J | Catalyzes the attachment of alanine to tRNA(Ala) in a two-step reaction alanine is first activated by ATP to form Ala- AMP and then transferred to the acceptor end of tRNA(Ala). Also edits incorrectly charged Ser-tRNA(Ala) and Gly-tRNA(Ala) via its editing domain | alaS | 509.6 | 0.31 | 9.22E-12 | 435.9 | 483.2 | NA | 85.4 | 56.8 | 83.8 |
| stage2 | stage3_shell | 2.42 | 6.50E-06 | V12B01_08210 | EAP92003.1 | putative oxaloacetate decarboxylase subunit gamma | | K01573 | C | Lyase and sodium transporter | oadG | 104.2 | 0.53 | 4.47E-07 | 368.8 | 394.0 | 370.1 | 49.5 | 11.7 | 88.3 |
| stage3_free | stage3_shell | 3.91 | 2.05E-11 | V12B01_08210 | EAP92003.1 | putative oxaloacetate decarboxylase subunit gamma | | K01573 | C | Lyase and sodium transporter | oadG | 104.2 | 0.56 | 1.81E-13 | 1145.2 | 938.4 | NA | 49.5 | 11.7 | 88.3 |
| stage3_free | stage3_shell | 2.74 | 5.51E-04 | V12B01_08215 | EAP92004.1 | oxaloacetate decarboxylase | Transport Proteins | K01571 | CI | oxaloacetate | oadA | 691.8 | 0.79 | 5.66E-05 | 1158.5 | 841.8 | NA | 16.1 | 43.3 | 172.6 |

| | | | | | | | | | | | | | | | | | | | | |
|---|---|---|---|---|---|---|---|---|---|---|---|---|---|---|---|---|---|---|---|---|
| stage2 | stage3_shell | 2.07 | 1.84E-04 | V12B01_08477 | EAP93013.1 | carbamoyl-phosphate synthase small subunit | Amino Acid Degradation | K01956 | F | carbamoyl-phosphate synthetase glutamine chain | carA | 287.8 | 0.55 | 2.09E-05 | 631.8 | 270.8 | 240.2 | 66.4 | 24.9 | 78.9 |
| stage3_free | stage3_shell | 3.46 | 4.35E-08 | V12B01_08477 | EAP93013.1 | carbamoyl-phosphate synthase small subunit | Amino Acid Degradation | K01956 | F | carbamoyl-phosphate synthetase glutamine chain | carA | 287.8 | 0.60 | 8.08E-10 | 988.0 | 998.2 | NA | 66.4 | 24.9 | 78.9 |
| stage2 | stage3_shell | 2.03 | 7.91E-04 | V12B01_08482 | EAP93014.1 | carbamoyl-phosphate synthase large subunit | Amino Acid Degradation | K01955 | F | Carbamoyl-phosphate synthetase ammonia chain | carB | 1316.9 | 0.60 | 1.12E-04 | 884.1 | 490.7 | 471.3 | 25.2 | 70.7 | 163.9 |
| stage3_free | stage3_shell | 3.23 | 3.79E-06 | V12B01_08482 | EAP93014.1 | carbamoyl-phosphate synthase large subunit | Amino Acid Degradation | K01955 | F | Carbamoyl-phosphate synthetase ammonia chain | carB | 1316.9 | 0.67 | 1.43E-07 | 1978.0 | 849.2 | NA | 25.2 | 70.7 | 163.9 |
| stage2 | stage3_shell | 2.39 | 8.13E-03 | V12B01_08512 | EAP93020.1 | adenosylcobinamide-phosphate synthase | Cofactor, Carrier, and Vitamin Biosynthesis | K02227 | H | COG1270 Cobalamin biosynthesis protein CobD CbiB | - | 11.0 | 0.91 | 1.82E-03 | 25.6 | 18.6 | 16.4 | 0.0 | 1.6 | 6.0 |
| stage3_free | stage3_shell | 2.83 | 3.47E-03 | V12B01_08512 | EAP93020.1 | adenosylcobinamide-phosphate synthase | Cofactor, Carrier, and Vitamin Biosynthesis | K02227 | H | COG1270 Cobalamin biosynthesis protein CobD CbiB | - | 11.0 | 0.95 | 5.19E-04 | 24.4 | 30.2 | NA | 0.0 | 1.6 | 6.0 |
| stage2 | stage3_free | 2.36 | 4.77E-05 | V12B01_08537 | EAP93025.1 | glutamate synthase [NADPH] small chain | TCA cycle | K00266 | C | COG0493 NADPH-dependent glutamate synthase beta chain and related oxidoreductases | gltD | 811.5 | 0.56 | 9.80E-07 | 1406.3 | 814.6 | 724.5 | 179.2 | 154.6 | NA |
| stage2 | stage3_shell | 2.64 | 1.57E-07 | V12B01_08537 | EAP93025.1 | glutamate synthase [NADPH] small chain | TCA cycle | K00266 | C | COG0493 NADPH-dependent glutamate synthase beta chain and related oxidoreductases | gltD | 811.5 | 0.49 | 6.63E-09 | 1406.3 | 814.6 | 724.5 | 37.0 | 169.8 | 106.5 |
| stage2 | stage3_shell | 3.09 | 4.50E-08 | V12B01_08542 | EAP93026.1 | glutamate synthase [NADPH] large chain | TCA cycle | K00265 | E | Glutamate synthase | gltB | 1933.7 | 0.55 | 1.54E-09 | 1374.9 | 500.2 | 434.6 | 23.9 | 63.2 | 83.9 |
| stage3_free | stage3_shell | 2.10 | 8.56E-04 | V12B01_08542 | EAP93026.1 | glutamate synthase [NADPH] large chain | TCA cycle | K00265 | E | Glutamate synthase | gltB | 1933.7 | 0.61 | 9.55E-05 | 557.8 | 276.5 | NA | 23.9 | 63.2 | 83.9 |
| **stage3_free** | **stage3_shell** | **-3.06** | **2.01E-04** | **V12B01_08572** | **EAP93032.1** | **ribonuclease activity regulator protein RraA** | **Regulation** | | **-** | **#N/A** | **#N/A** | **30.0** | **0.84** | **1.70E-05** | **11.5** | **66.5** | **NA** | **462.2** | **352.1** | **107.4** |
| **stage2** | **stage3_shell** | **-2.11** | **7.91E-04** | **V12B01_08582** | **EAP93034.1** | **hypothetical protein** | | | **-** | **-** | **-** | **47.0** | **0.63** | **1.12E-04** | **21.0** | **47.1** | **40.8** | **291.5** | **105.6** | **25.7** |
| **stage3_free** | **stage3_shell** | **-2.86** | **9.94E-05** | **V12B01_08582** | **EAP93034.1** | **hypothetical protein** | | | **-** | **-** | **-** | **47.0** | **0.74** | **7.32E-06** | **14.4** | **28.1** | **NA** | **291.5** | **105.6** | **25.7** |
| stage2 | stage3_shell | 2.49 | 4.64E-09 | V12B01_08597 | EAP93037.1 | threonine synthase | Amino Acid Biosynthesis | K01733 | E | Threonine synthase | thrC | 290.4 | 0.41 | 1.21E-10 | 512.9 | 230.3 | 281.2 | 27.4 | 48.3 | 48.8 |
| **stage2** | **stage3_free** | **-4.61** | **NA** | **V12B01_08602** | **EAP93038.1** | **formate acetyltransferase** | **Fermentation** | **K06866** | **S** | **Acts as a radical domain for damaged PFL and possibly other radical proteins** | **grcA** | **227.1** | **1.07** | **NA** | **42.3** | **46.8** | **52.0** | **1708.4** | **1499.5** | **NA** |
| **stage2** | **stage3_shell** | **-2.21** | **NA** | **V12B01_08602** | **EAP93038.1** | **formate acetyltransferase** | **Fermentation** | **K06866** | **S** | **Acts as a radical domain for damaged PFL and possibly other radical proteins** | **grcA** | **227.1** | **1.03** | **NA** | **42.3** | **46.8** | **52.0** | **384.6** | **28.0** | **286.2** |
| **stage2** | **stage3_free** | **-2.46** | **3.57E-04** | **V12B01_08617** | **EAP93041.1** | **hypothetical protein** | | | **S** | **Hemerythrin HHE cation binding domain** | **-** | **656.6** | **0.64** | **1.09E-05** | **33.8** | **100.7** | **86.5** | **482.2** | **513.4** | **NA** |
| **stage2** | **stage3_shell** | **-5.12** | **1.73E-18** | **V12B01_08617** | **EAP93041.1** | **hypothetical protein** | | | **S** | **Hemerythrin HHE cation binding domain** | **-** | **656.6** | **0.56** | **6.78E-21** | **33.8** | **100.7** | **86.5** | **1047.8** | **1521.1** | **3146.4** |
| **stage3_free** | **stage3_shell** | **-2.22** | **5.45E-04** | **V12B01_08617** | **EAP93041.1** | **hypothetical protein** | | | **S** | **Hemerythrin HHE cation binding domain** | **-** | **656.6** | **0.65** | **5.57E-05** | **482.2** | **513.4** | **NA** | **1047.8** | **1521.1** | **3146.4** |
| **stage2** | **stage3_free** | **-2.82** | **8.88E-05** | **V12B01_08647** | **EAP92308.1** | **Branched-chain amino acid permease** | **Transport Proteins** | **K03311** | **U** | **Component of the transport system for branched-chain amino acids** | **brnQ** | **27.3** | **0.68** | **2.13E-06** | **14.5** | **6.1** | **5.3** | **44.2** | **106.2** | **NA** |
| stage2 | stage3_shell | 2.09 | 5.49E-06 | V12B01_08662 | EAP92311.1 | thiol:disulfide interchange protein DsbC | | K03981 | O | Required for disulfide bond formation in some periplasmic proteins. Acts by transferring its disulfide bond to other proteins and is reduced in the process | dsbC | 59.4 | 0.44 | 3.62E-07 | 128.3 | 92.6 | 86.0 | 22.5 | 11.6 | 14.6 |
| **stage2** | **stage3_shell** | **-2.12** | **4.83E-09** | **V12B01_08692** | **EAP92317.1** | **hypothetical protein** | | | **-** | **-** | **VP0517** | **161.9** | **0.34** | **1.30E-10** | **89.8** | **187.3** | **182.0** | **445.5** | **790.4** | **328.5** |
| **stage3_free** | **stage3_shell** | **-2.88** | **5.60E-12** | **V12B01_08692** | **EAP92317.1** | **hypothetical protein** | | | **-** | **-** | **VP0517** | **161.9** | **0.40** | **4.25E-14** | **66.0** | **115.5** | **NA** | **445.5** | **790.4** | **328.5** |
| **stage2** | **stage3_shell** | **-3.70** | **6.10E-04** | **V12B01_08802** | **EAP92681.1** | **Carbon starvation protein, predicted membrane protein** | **Membrane** | | **T** | **Carbon starvation protein** | **cstA** | **12.0** | **1.17** | **8.24E-05** | **1.8** | **0.5** | **0.8** | **36.4** | **0.0** | **7.6** |
| **stage3_free** | **stage3_shell** | **-2.16** | **2.74E-02** | **V12B01_08802** | **EAP92681.1** | **Carbon starvation protein, predicted membrane protein** | **Membrane** | | **T** | **Carbon starvation protein** | **cstA** | **12.0** | **1.49** | **6.97E-03** | **2.8** | **1.6** | **NA** | **36.4** | **0.0** | **7.6** |
| stage2 | stage3_shell | 2.12 | 2.19E-02 | V12B01_08862 | EAP92693.1 | hypothetical protein | | K01529 | F | Phosphatase that hydrolyzes non-canonical purine nucleotides such as XTP and ITP to their respective diphosphate derivatives. Probably excludes non-canonical purines from DNA precursor pool, thus preventing their incorporation into DNA and avoiding chromosomal lesions | yjjX | 8.9 | 0.94 | 6.17E-03 | 26.4 | 37.3 | 25.1 | 0.0 | 8.7 | 0.0 |
| stage2 | stage3_shell | 2.02 | 2.49E-02 | V12B01_08867 | EAP92694.1 | c-di-GMP phosphodiesterase A-related protein | Regulation | K20966 | T | COG2199 FOG GGDEF domain | mbaA | 8.1 | 0.97 | 7.16E-03 | 13.8 | 6.1 | 4.3 | 1.3 | 1.3 | 0.0 |
| stage2 | stage3_shell | 2.46 | 3.79E-03 | V12B01_08887 | EAP92698.1 | DNA uptake lipoprotein | Transport Proteins | K05807 | M | Part of the outer membrane protein assembly complex, which is involved in assembly and insertion of beta-barrel proteins into the outer membrane | bamD | 16.2 | 0.87 | 7.26E-04 | 94.5 | 34.0 | 15.0 | 8.8 | 4.2 | 0.0 |
| stage3_free | stage3_shell | 2.01 | 2.11E-02 | V12B01_08887 | EAP92698.1 | DNA uptake lipoprotein | Transport Proteins | K05807 | M | Part of the outer membrane protein assembly complex, which is involved in assembly and insertion of beta-barrel proteins into the outer membrane | bamD | 16.2 | 0.92 | 5.07E-03 | 37.6 | 38.9 | NA | 8.8 | 4.2 | 0.0 |

| | | | | | | | | | | | | | | | | | | | |
|---|---|---|---|---|---|---|---|---|---|---|---|---|---|---|---|---|---|---|---|
| **stage2** | stage3_shell | **-3.03** | **3.63E-13** | **V12B01_08909** | **EAP92358.1** | **penicillin-insensitive murein endopeptidase** | **Membrane** | **K07261** | **M** | **Murein endopeptidase that cleaves the D-alanyl-meso-2,6- diamino-pimelyl amide bond that connects peptidoglycan strands. Likely plays a role in the removal of murein from the sacculus** | **mepA** | **82.8** | **0.40** | **3.91E-15** | **11.6** | **26.1** | **23.8** | **91.3** | **138.7** | **159.3** |
| **stage3_free** | stage3_shell | **-3.50** | **9.37E-13** | **V12B01_08909** | **EAP92358.1** | **penicillin-insensitive murein endopeptidase** | **Membrane** | **K07261** | **M** | **Murein endopeptidase that cleaves the D-alanyl-meso-2,6- diamino-pimelyl amide bond that connects peptidoglycan strands. Likely plays a role in the removal of murein from the sacculus** | **mepA** | **82.8** | **0.47** | **6.33E-15** | **13.1** | **16.6** | **NA** | **91.3** | **138.7** | **159.3** |
| **stage2** | stage3_shell | **-3.60** | **7.80E-08** | **V12B01_08914** | **EAP92359.1** | **hypothetical protein** | | | **-** | **-** | **VP0565** | **658.8** | **0.65** | **3.00E-09** | **79.1** | **560.5** | **346.4** | **5382.7** | **4675.7** | **598.9** |
| **stage3_free** | stage3_shell | **-4.87** | **1.98E-10** | **V12B01_08914** | **EAP92359.1** | **hypothetical protein** | | | **-** | **-** | **VP0565** | **658.8** | **0.75** | **2.05E-12** | **116.9** | **157.0** | **NA** | **5382.7** | **4675.7** | **598.9** |
| stage2 | stage3_shell | 2.18 | 6.38E-06 | V12B01_08999 | EAP92376.1 | alkyl hydroperoxide reductase C22 protein | | K03386 | O | COG0450 Peroxiredoxin | - | 849.6 | 0.47 | 4.34E-07 | 2971.0 | 2230.9 | 2604.6 | 514.2 | 159.0 | 439.5 |
| stage3_free | stage3_shell | 2.54 | 3.59E-06 | V12B01_08999 | EAP92376.1 | alkyl hydroperoxide reductase C22 protein | | K03386 | O | COG0450 Peroxiredoxin | - | 849.6 | 0.52 | 1.32E-07 | 3875.1 | 2969.7 | NA | 514.2 | 159.0 | 439.5 |
| **stage2** | stage3_shell | **-2.24** | **3.74E-03** | **V12B01_09046** | **EAP94316.1** | **Putative acetoin utilization protein AcuB** | | **K07168** | **S** | **acetoin utilization protein** | **-** | **28.4** | **0.84** | **7.17E-04** | **38.1** | **3.2** | **21.1** | **43.2** | **64.7** | **166.7** |
| stage2 | stage3_free | 2.25 | 8.14E-04 | V12B01_09106 | EAP94328.1 | co-chaperone HscB | | K04082 | O | Co-chaperone involved in the maturation of iron-sulfur cluster-containing proteins. Seems to help targeting proteins to be folded toward HscA | hscB | 98.4 | 0.66 | 3.03E-05 | 174.9 | 169.5 | 403.4 | 43.4 | 41.9 | NA |
| stage2 | stage3_free | 2.17 | 1.33E-03 | V12B01_09116 | EAP94330.1 | ferredoxin | | K04755 | C | Ferredoxin | fdx | 53.6 | 0.67 | 6.01E-05 | 225.6 | 140.0 | 268.8 | 44.5 | 31.2 | NA |
| stage2 | stage3_shell | 2.09 | 1.31E-03 | V12B01_09116 | EAP94330.1 | ferredoxin | | K04755 | C | Ferredoxin | fdx | 53.6 | 0.66 | 2.06E-04 | 225.6 | 140.0 | 268.8 | 18.9 | 31.2 | 42.0 |
| stage3_free | stage3_shell | 3.05 | 1.64E-04 | V12B01_09131 | EAP94333.1 | nucleoside diphosphate kinase | Nucleoside and Nucleotide Biosynthesis | K00940 | F | Major role in the synthesis of nucleoside triphosphates other than ATP. The ATP gamma phosphate is transferred to the NDP beta phosphate via a ping-pong mechanism, using a phosphorylated active-site intermediate | ndk | 27.2 | 0.81 | 1.33E-05 | 225.0 | 167.9 | NA | 14.7 | 3.5 | 26.2 |
| **stage2** | **stage3_free** | **-2.18** | **1.36E-02** | **V12B01_09191** | **EAP94345.1** | **putative ureashort-chain amide or branched-chain amino acid uptake ABC transporter periplasmic solute-binding protein** | **Transport Proteins** | **K11959** | **E** | **COG0683 ABC-type branched-chain amino acid transport systems, periplasmic component** | **urtA** | **14.8** | **0.90** | **9.73E-04** | **5.5** | **5.0** | **6.1** | **55.1** | **19.0** | **NA** |
| stage3_free | stage3_shell | 2.23 | 5.55E-02 | V12B01_09196 | EAP94346.1 | putative ureashort-chain amide or branched-chain amino acid uptake ABC transporter permease protein, possibly fusion | Transport Proteins | K11960 | E | Belongs to the binding-protein-dependent transport system permease family | urtB | 5.6 | 1.64 | 1.72E-02 | 18.6 | 4.7 | NA | 0.0 | 1.9 | 0.0 |
| **stage2** | **stage3_free** | **-6.00** | **NA** | **V12B01_09226** | **EAP94352.1** | **urease, beta subunit** | **Amine and Polyamine Degradation** | **K01429** | **E** | **Belongs to the urease beta subunit family** | **ureB** | **2.8** | **1.92** | **5.77E-05** | **1.3** | **0.0** | **0.0** | **87.1** | **24.3** | **NA** |
| stage3_free | stage3_shell | 4.65 | 1.40E-02 | V12B01_09226 | EAP94352.1 | urease, beta subunit | Amine and Polyamine Degradation | K01429 | E | Belongs to the urease beta subunit family | ureB | 2.8 | 2.95 | 3.02E-03 | 87.1 | 24.3 | NA | 0.0 | 0.0 | 0.0 |
| **stage2** | **stage3_free** | **-3.81** | **1.24E-03** | **V12B01_09231** | **EAP94353.1** | **urease, alpha subunit** | **Amine and Polyamine Degradation** | **K01428** | **E** | **Belongs to the metallo-dependent hydrolases superfamily. Urease alpha subunit family** | **ureC** | **16.7** | **1.16** | **5.45E-05** | **4.2** | **1.2** | **1.3** | **82.7** | **14.4** | **NA** |
| stage3_free | stage3_shell | 7.59 | 4.78E-05 | V12B01_09231 | EAP94353.1 | urease, alpha subunit | Amine and Polyamine Degradation | K01428 | E | Belongs to the metallo-dependent hydrolases superfamily. Urease alpha subunit family | ureC | 16.7 | 2.67 | 3.17E-06 | 82.7 | 14.4 | NA | 0.0 | 0.0 | 0.0 |
| **stage2** | **stage3_free** | **-2.51** | **9.46E-04** | **V12B01_09261** | **EAP94359.1** | **inositol-5-monophosphate dehydrogenase** | **Nucleoside and Nucleotide Degradation** | **K00088** | **F** | **Catalyzes the conversion of inosine 5'-phosphate (IMP) to xanthosine 5'-phosphate (XMP), the first committed and rate- limiting step in the de novo synthesis of guanine nucleotides, and therefore plays an important role in the regulation of cell growth** | **guaB** | **340.5** | **0.72** | **3.75E-05** | **252.7** | **116.8** | **105.9** | **1219.0** | **1119.9** | **NA** |
| stage3_free | stage3_shell | 2.60 | 3.84E-04 | V12B01_09261 | EAP94359.1 | inositol-5-monophosphate dehydrogenase | Nucleoside and Nucleotide Degradation | K00088 | F | Catalyzes the conversion of inosine 5'-phosphate (IMP) to xanthosine 5'-phosphate (XMP), the first committed and rate- limiting step in the de novo synthesis of guanine nucleotides, and therefore plays an important role in the regulation of cell growth | guaB | 340.5 | 0.71 | 3.77E-05 | 1219.0 | 1119.9 | NA | 184.4 | 34.1 | 118.6 |
| stage3_free | stage3_shell | 3.17 | 9.10E-06 | V12B01_09266 | EAP94360.1 | bifunctional GMP synthase/glutamine amidotransferase protein | Amino Acid Degradation | K01951 | F | Catalyzes the synthesis of GMP from XMP | guaA | 274.5 | 0.69 | 4.21E-07 | 969.2 | 635.2 | NA | 23.6 | 27.3 | 102.6 |
| **stage2** | **stage3_free** | **-2.50** | **2.30E-05** | **V12B01_09276** | **EAP94362.1** | **hypothetical protein** | | | **C** | **C4-dicarboxylate anaerobic carrier** | **-** | **14.5** | **0.55** | **4.26E-07** | **4.9** | **4.1** | **4.8** | **42.2** | **18.6** | **NA** |
| stage3_free | stage3_shell | 2.50 | 7.88E-04 | V12B01_09276 | EAP94362.1 | hypothetical protein | | | C | C4-dicarboxylate anaerobic carrier | - | 14.5 | 0.73 | 8.61E-05 | 42.2 | 18.6 | NA | 4.1 | 2.0 | 3.7 |
| **stage2** | **stage3_shell** | **-2.39** | **9.79E-04** | **V12B01_09281** | **EAP94363.1** | **Predicted membrane protein** | **Membrane** | **K07034** | **S** | **GPR1/FUN34/yaaH family** | **yaaH** | **49.2** | **0.75** | **1.45E-04** | **3.0** | **9.5** | **6.3** | **37.7** | **30.6** | **24.0** |
| **stage3_free** | **stage3_shell** | **-2.61** | **2.07E-03** | **V12B01_09281** | **EAP94363.1** | **Predicted membrane protein** | **Membrane** | **K07034** | **S** | **GPR1/FUN34/yaaH family** | **yaaH** | **49.2** | **0.89** | **2.81E-04** | **3.1** | **7.3** | **NA** | **37.7** | **30.6** | **24.0** |
| stage2 | stage3_shell | 2.06 | 1.48E-05 | V12B01_09286 | EAP94364.1 | sodium/alanine symporter | Transport Proteins | K03310 | U | COG1115 Na alanine symporter | - | 269.5 | 0.46 | 1.21E-06 | 504.1 | 189.7 | 164.1 | 59.9 | 41.0 | 36.8 |
| stage3_free | stage3_shell | 2.88 | 7.30E-08 | V12B01_09286 | EAP94364.1 | sodium/alanine symporter | Transport Proteins | K03310 | U | COG1115 Na alanine symporter | - | 269.5 | 0.51 | 1.52E-09 | 608.0 | 423.0 | NA | 59.9 | 41.0 | 36.8 |
| stage3_free | stage3_shell | 2.52 | 9.13E-03 | V12B01_09311 | EAP94369.1 | hypothetical protein | | | S | Protein of unknown function (DUF429) | - | 13.1 | 0.96 | 1.71E-03 | 40.8 | 17.1 | NA | 0.0 | 4.2 | 3.9 |

| | | | | | | | | | | | | | | | | | | | | |
|---|---|---|---|---|---|---|---|---|---|---|---|---|---|---|---|---|---|---|---|---|
| stage2 | stage3_shell | -2.55 | 2.60E-03 | V12B01_09376 | EAP94382.1 | hypothetical protein | | | - | #N/A | #N/A | 11.3 | 0.91 | 4.68E-04 | 5.6 | 3.3 | 5.9 | 40.2 | 28.5 | 17.9 |
| stage2 | stage3_shell | -2.07 | 5.01E-02 | V12B01_09391 | EAP94385.1 | hypothetical protein | | | - | #N/A | #N/A | 2.1 | 2.69 | 1.69E-02 | 0.0 | 0.0 | 4.6 | 35.5 | 0.0 | 42.2 |
| stage3_free | stage3_shell | -2.20 | 3.05E-06 | V12B01_09401 | EAP94387.1 | signal transduction histidine kinase | Regulation | | T | His Kinase A (phospho-acceptor) domain | - | 581.8 | 0.45 | 1.11E-07 | 93.2 | 107.1 | NA | 423.8 | 564.1 | 150.1 |
| stage2 | stage3_free | -4.69 | 1.56E-11 | V12B01_09441 | EAP94395.1 | Sodium/glutamate symporter | Transport Proteins | | U | Belongs to the dicarboxylate amino acid cation symporter (DAACS) (TC 2.A.23) family | - | 75.2 | 0.66 | 6.54E-14 | 17.4 | 7.8 | 6.1 | 328.6 | 290.9 | NA |
| stage3_free | stage3_shell | 2.53 | 2.70E-04 | V12B01_09441 | EAP94395.1 | Sodium/glutamate symporter | Transport Proteins | | U | Belongs to the dicarboxylate amino acid cation symporter (DAACS) (TC 2.A.23) family | - | 75.2 | 0.67 | 2.44E-05 | 328.6 | 290.9 | NA | 67.7 | 7.1 | 22.3 |
| stage2 | stage3_shell | 2.10 | 1.58E-04 | V12B01_09451 | EAP94397.1 | molecular chaperone DnaK | Replication | K04043 | O | Heat shock 70 kDa protein | dnaK | 604.3 | 0.55 | 1.76E-05 | 697.0 | 126.2 | 804.6 | 61.8 | 63.2 | 108.6 |
| stage2 | stage3_free | -4.15 | 1.60E-13 | V12B01_09486 | EAP94404.1 | putative p-aminobenzoyl-glutamate transporter | Transport Proteins | K12942 | H | p-aminobenzoyl-glutamate transporter | abgT | 46.5 | 0.53 | 3.30E-16 | 6.0 | 5.9 | 4.4 | 91.3 | 121.4 | NA |
| stage3_free | stage3_shell | 2.28 | 1.01E-04 | V12B01_09486 | EAP94404.1 | putative p-aminobenzoyl-glutamate transporter | Transport Proteins | K12942 | H | p-aminobenzoyl-glutamate transporter | abgT | 46.5 | 0.57 | 7.49E-06 | 91.3 | 121.4 | NA | 10.2 | 13.6 | 18.2 |
| stage2 | stage3_shell | -4.42 | 5.60E-14 | V12B01_09506 | EAP93843.1 | hypothetical protein | | | - | - | - | 61.8 | 0.57 | 5.32E-16 | 32.9 | 65.6 | 55.5 | 603.6 | 885.3 | 1117.8 |
| stage3_free | stage3_shell | -2.91 | 6.43E-06 | V12B01_09506 | EAP93843.1 | hypothetical protein | | | - | - | - | 61.8 | 0.64 | 2.76E-07 | 176.2 | 103.2 | NA | 603.6 | 885.3 | 1117.8 |
| stage3_free | stage3_shell | 2.64 | 1.70E-04 | V12B01_09521 | EAP93846.1 | phosphoribosylformylglycinamidine synthase | Nucleoside and Nucleotide Biosynthesis | K01952 | F | Phosphoribosylformylglycinamidine synthase involved in the purines biosynthetic pathway. Catalyzes the ATP-dependent conversion of formylglycinamide ribonucleotide (FGAR) and glutamine to yield formylglycinamidine ribonucleotide (FGAM) and glutamate | purL | 654.5 | 0.68 | 1.39E-05 | 693.9 | 443.0 | NA | 31.6 | 57.2 | 70.8 |
| stage2 | stage3_shell | 2.14 | 1.00E-06 | V12B01_09541 | EAP93850.1 | 2,3,4,5-tetrahydropyridine-2-carboxylate N-succinyltransferase | | K00674 | H | Catalyzes the conversion of the cyclic tetrahydrodipicolinate (THDP) into the acyclic N-succinyl-L-2- amino-6-oxopimelate using succinyl-CoA | dapD | 80.2 | 0.42 | 5.43E-08 | 200.8 | 114.3 | 102.5 | 17.0 | 23.5 | 24.8 |
| stage3_free | stage3_shell | 2.10 | 1.27E-05 | V12B01_09541 | EAP93850.1 | 2,3,4,5-tetrahydropyridine-2-carboxylate N-succinyltransferase | | K00674 | H | Catalyzes the conversion of the cyclic tetrahydrodipicolinate (THDP) into the acyclic N-succinyl-L-2- amino-6-oxopimelate using succinyl-CoA | dapD | 80.2 | 0.45 | 6.23E-07 | 132.5 | 145.2 | NA | 17.0 | 23.5 | 24.8 |
| stage3_free | stage3_shell | 3.29 | 1.13E-08 | V12B01_09601 | EAP93862.1 | N-acetylglutamate synthase | Amino Acid Biosynthesis | K14682 | E | Amino-acid acetyltransferase | argA | 233.5 | 0.55 | 1.83E-10 | 571.2 | 348.6 | NA | 31.0 | 13.6 | 46.8 |
| stage2 | stage3_free | -2.34 | 1.60E-13 | V12B01_09636 | EAP93869.1 | outer membrane protein OmpK precursor | Outer membrane | K05517 | M | Nucleoside-specific channel-forming protein, Tsx | ompK | 76.4 | 0.30 | 2.84E-16 | 26.6 | 25.6 | 34.3 | 149.6 | 160.1 | NA |
| stage2 | stage3_shell | 2.60 | 7.57E-06 | V12B01_09656 | EAP93873.1 | lipoprotein, putative | | K07286 | M | lipoprotein | VV2594 | 58.2 | 0.56 | 5.47E-07 | 252.8 | 139.2 | 128.1 | 33.6 | 10.6 | 10.0 |
| stage3_free | stage3_shell | 2.80 | 9.91E-06 | V12B01_09656 | EAP93873.1 | lipoprotein, putative | | K07286 | M | lipoprotein | VV2594 | 58.2 | 0.60 | 4.69E-07 | 211.8 | 197.2 | NA | 33.6 | 10.6 | 10.0 |
| stage3_free | stage3_shell | -2.42 | 5.53E-08 | V12B01_09716 | EAP93885.1 | hypothetical protein | | | L | Belongs to the 'phage' integrase family | - | 714.3 | 0.42 | 1.07E-09 | 101.8 | 130.0 | NA | 617.2 | 697.9 | 192.6 |
| stage3_free | stage3_shell | -2.03 | 1.44E-03 | V12B01_09721 | EAP93886.1 | hypothetical protein | | | - | - | - | 344.6 | 0.64 | 1.77E-04 | 28.1 | 46.8 | NA | 137.0 | 252.8 | 31.0 |
| stage2 | stage3_shell | 2.64 | 7.91E-04 | V12B01_09876 | EAP91916.1 | DNA polymerase III subunit delta | Replication | K02340 | L | DNA polymerase III, delta subunit | holA | 24.2 | 0.80 | 1.12E-04 | 56.8 | 44.2 | 37.1 | 4.7 | 2.9 | 5.6 |
| stage3_free | stage3_shell | 2.08 | 1.07E-02 | V12B01_09876 | EAP91916.1 | DNA polymerase III subunit delta | | K02340 | L | DNA polymerase III, delta subunit | holA | 24.2 | 0.85 | 2.12E-03 | 47.7 | 20.1 | NA | 4.7 | 2.9 | 5.6 |
| stage2 | stage3_shell | -2.94 | 1.42E-12 | V12B01_09891 | EAP91919.1 | hypothetical protein | | | J | Zinc-ribbon containing domain | - | 923.2 | 0.40 | 1.76E-14 | 129.3 | 285.7 | 318.0 | 739.5 | 1715.2 | 1807.5 |
| stage3_free | stage3_shell | -2.58 | 6.05E-08 | V12B01_09891 | EAP91919.1 | hypothetical protein | | | J | Zinc-ribbon containing domain | - | 923.2 | 0.45 | 1.21E-09 | 290.3 | 335.5 | NA | 739.5 | 1715.2 | 1807.5 |
| stage2 | stage3_shell | 2.07 | 5.75E-05 | V12B01_09896 | EAP91920.1 | apolipoprotein N-acyltransferase | | K03820 | M | Transfers the fatty acyl group on membrane lipoproteins | Int | 36.3 | 0.49 | 5.57E-06 | 46.6 | 37.3 | 28.6 | 6.3 | 7.0 | 3.8 |
| stage3_free | stage3_shell | 2.47 | 2.43E-05 | V12B01_09916 | EAP91924.1 | tRNA 2-methylthioadenosine synthase | Translation | K06168 | J | Catalyzes the methylthiolation of N6-(dimethylallyl)adenosine (i(6)A), leading to the formation of 2- methylthio-N6-(dimethylallyl)adenosine (ms(2)i(6)A) at position 37 in tRNAs that read codons beginning with uridine | miaB | 81.7 | 0.55 | 1.36E-06 | 122.9 | 195.2 | NA | 31.4 | 17.0 | 6.0 |
| stage3_free | stage3_shell | -2.15 | 7.52E-06 | V12B01_09993 | EAP91868.1 | UDP-N-acetylmuramyl pentapeptide synthase | Cell Structure Biosynthesis | K01929 | M | Involved in cell wall formation. Catalyzes the final step in the synthesis of UDP-N-acetylmuramoyl-pentapeptide, the precursor of murein | murF | 163.3 | 0.46 | 3.40E-07 | 34.0 | 33.9 | NA | 65.1 | 202.6 | 95.3 |
| stage2 | stage3_shell | 2.22 | 2.40E-05 | V12B01_09998 | EAP91869.1 | UDP-N-acetylmuramoylalanyl-D-glutamate--2,6-diaminopimelate ligase | Cell Structure Biosynthesis | K01928 | M | Catalyzes the addition of meso-diaminopimelic acid to the nucleotide precursor UDP-N-acetylmuramoyl-L-alanyl-D-glutamate (UMAG) in the biosynthesis of bacterial cell-wall peptidoglycan | murE | 64.0 | 0.51 | 2.08E-06 | 76.7 | 76.3 | 85.3 | 4.3 | 19.4 | 9.6 |
| stage2 | stage3_shell | -6.45 | 3.79E-04 | V12B01_10018 | EAP91873.1 | hypothetical protein | | | - | - | - | 1733.7 | 2.17 | 4.75E-05 | 70.6 | 129.3 | 95.8 | 17042.7 | 27423.9 | 512.2 |
| stage3_free | stage3_shell | 2.08 | 1.28E-04 | V12B01_10033 | EAP91752.1 | hypothetical protein | | | - | #N/A | #N/A | 67.7 | 0.51 | 1.01E-05 | 1842.5 | 2751.8 | NA | 684.8 | 259.4 | 81.3 |
| stage2 | stage3_shell | 2.47 | 1.55E-09 | V12B01_10048 | EAP91755.1 | protease DO | | K04772 | M | Belongs to the peptidase S1C family | degQ | 173.9 | 0.39 | 3.47E-11 | 294.1 | 197.7 | 172.6 | 20.0 | 33.5 | 29.4 |
| stage3_free | stage3_shell | 2.11 | 7.30E-08 | V12B01_10123 | EAP92131.1 | pho4 family protein | | K03306 | P | COG0306 Phosphate sulphate permeases | pitA | 117.9 | 0.37 | 1.53E-09 | 213.3 | 219.8 | NA | 22.8 | 40.7 | 40.6 |
| stage2 | stage3_shell | 2.03 | 1.82E-05 | V12B01_10188 | EAP91727.1 | homocysteine synthase | | K01740 | E | COG2873 O-acetylhomoserine sulfhydrylase | metY | 102.9 | 0.46 | 1.52E-06 | 182.6 | 99.4 | 87.1 | 17.6 | 16.7 | 26.9 |
| stage2 | stage3_free | -5.25 | 2.89E-11 | V12B01_10193 | EAP91728.1 | D-fructose-6-phosphate amidotransferase | Carbohydrate Biosynthesis | K00820 | M | Catalyzes the first step in hexosamine metabolism, converting fructose-6P into glucosamine-6P using glutamine as a nitrogen source | glmS | 1045.7 | 0.74 | 1.27E-13 | 59.4 | 64.9 | 42.2 | 2232.9 | 2744.1 | NA |

| | | | | | | | | | | | | | | | | | | | | |
|---|---|---|---|---|---|---|---|---|---|---|---|---|---|---|---|---|---|---|---|---|
| stage2 | stage3_shell | -2.65 | 9.88E-05 | V12B01_10193 | EAP91728.1 | D-fructose-6-phosphate amidotransferase | Carbohydrate Biosynthesis | K00820 | M | Catalyzes the first step in hexosamine metabolism, converting fructose-6P into glucosamine-6P using glutamine as a nitrogen source | glmS | 1045.7 | 0.69 | 1.03E-05 | 59.4 | 64.9 | 42.2 | 142.1 | 122.2 | 574.7 |
| stage3_free | stage3_shell | 2.19 | 3.66E-03 | V12B01_10193 | EAP91728.1 | D-fructose-6-phosphate amidotransferase | Carbohydrate Biosynthesis | K00820 | M | Catalyzes the first step in hexosamine metabolism, converting fructose-6P into glucosamine-6P using glutamine as a nitrogen source | glmS | 1045.7 | 0.75 | 5.59E-04 | 2232.9 | 2744.1 | NA | 142.1 | 122.2 | 574.7 |
| stage2 | stage3_shell | -3.46 | 9.63E-05 | V12B01_10198 | EAP91729.1 | transcriptional regulator, DeoR family | Regulation | | K | COG1349 Transcriptional regulators of sugar metabolism | srlR | 375.9 | 0.91 | 9.99E-06 | 51.3 | 95.5 | 72.4 | 1742.8 | 157.0 | 443.1 |
| stage2 | stage3_shell | -4.53 | 1.54E-10 | V12B01_10215 | EAP92570.1 | hypothetical protein | | | - | - | - | 581.3 | 0.69 | 2.64E-12 | 64.0 | 501.8 | 236.4 | 6112.6 | 9237.7 | 1078.5 |
| stage3_free | stage3_shell | -5.87 | 5.42E-13 | V12B01_10215 | EAP92570.1 | hypothetical protein | | | - | - | - | 581.3 | 0.79 | 3.20E-15 | 72.8 | 138.3 | NA | 6112.6 | 9237.7 | 1078.5 |
| stage2 | stage3_shell | -3.46 | 1.82E-09 | V12B01_10377 | EAP92234.1 | malate dehydrogenase | TCA cycle | | - | - | - | 148.9 | 0.55 | 4.10E-11 | 19.4 | 91.4 | 37.1 | 418.5 | 720.4 | 226.0 |
| stage3_free | stage3_shell | -4.13 | 1.15E-09 | V12B01_10377 | EAP92234.1 | malate dehydrogenase | TCA cycle | | - | - | - | 148.9 | 0.66 | 1.55E-11 | 20.0 | 41.9 | NA | 418.5 | 720.4 | 226.0 |
| stage2 | stage3_shell | 2.16 | 4.69E-09 | V12B01_10382 | EAP92235.1 | malate dehydrogenase | | K00024 | C | Catalyzes the reversible oxidation of malate to oxaloacetate | mdh | 528.3 | 0.35 | 1.24E-10 | 1411.4 | 680.7 | 745.1 | 148.5 | 105.1 | 179.5 |
| stage3_free | stage3_shell | 2.37 | 1.10E-08 | V12B01_10382 | EAP92235.1 | malate dehydrogenase | TCA cycle | K00024 | C | Catalyzes the reversible oxidation of malate to oxaloacetate | mdh | 528.3 | 0.39 | 1.77E-10 | 1214.2 | 1021.3 | NA | 148.5 | 105.1 | 179.5 |
| stage2 | stage3_shell | -2.04 | 8.65E-09 | V12B01_10427 | EAP92244.1 | hypothetical protein | | K07175 | T | ATPase related to phosphate starvation-inducible protein PhoH | - | 207.6 | 0.33 | 2.48E-10 | 7.7 | 10.3 | 15.4 | 32.4 | 50.4 | 22.7 |
| stage2 | stage3_shell | 4.04 | 6.05E-03 | V12B01_10482 | EAP92147.1 | MSHA biogenesis protein MshP | Membrane | K12286 | NU | Pilus assembly protein PilX | mshP | 8.6 | 2.29 | 1.29E-03 | 24.7 | 37.0 | 15.3 | 0.0 | 0.0 | 0.0 |
| stage3_free | stage3_shell | 2.86 | 3.13E-02 | V12B01_10482 | EAP92147.1 | MSHA biogenesis protein MshP | Membrane | K12286 | NU | Pilus assembly protein PilX | mshP | 8.6 | 2.32 | 8.24E-03 | 18.3 | 14.9 | NA | 0.0 | 0.0 | 0.0 |
| stage2 | stage3_shell | 3.94 | 4.33E-03 | V12B01_10527 | EAP92156.1 | MSHA biogenesis protein MshN | Membrane | K12284 | S | MSHA biogenesis protein MshN | mshN | 5.2 | 2.12 | 8.57E-04 | 7.7 | 9.8 | 7.3 | 0.0 | 0.0 | 0.0 |
| stage3_free | stage3_shell | 2.22 | 5.05E-02 | V12B01_10527 | EAP92156.1 | MSHA biogenesis protein MshN | Membrane | K12284 | S | MSHA biogenesis protein MshN | mshN | 5.2 | 1.97 | 1.51E-02 | 3.6 | 4.6 | NA | 0.0 | 0.0 | 0.0 |
| stage3_free | stage3_shell | 2.20 | 2.95E-04 | V12B01_10567 | EAP92164.1 | single-strand DNA-binding protein | Regulation | K03111 | L | Plays an important role in DNA replication, recombination and repair. Binds to ssDNA and to an array of partner proteins to recruit them to their sites of action during DNA metabolism | ssb | 48.6 | 0.59 | 2.72E-05 | 234.9 | 325.7 | NA | 61.2 | 20.4 | 32.1 |
| stage3_free | stage3_shell | 2.33 | 2.38E-04 | V12B01_10597 | EAP91930.1 | aminotransferase, class V | | K00830 | E | COG0075 Serine-pyruvate aminotransferase archaeal aspartate aminotransferase | - | 110.7 | 0.62 | 2.10E-05 | 383.5 | 246.1 | NA | 63.1 | 10.6 | 40.0 |
| stage2 | stage3_shell | 2.00 | 1.27E-04 | V12B01_10602 | EAP91931.1 | aspartate kinase III | Amino Acid Biosynthesis | K00928 | E | Belongs to the aspartokinase family | lysC | 341.9 | 0.51 | 1.36E-05 | 380.6 | 152.0 | 169.5 | 28.0 | 27.6 | 56.2 |
| stage2 | stage3_shell | 2.00 | 4.90E-03 | V12B01_10630 | EAP93068.1 | phosphoadenosine phosphosulfate reductase | | K00390 | EH | Belongs to the PAPS reductase family. CysH subfamily | cysH | 229.4 | 0.75 | 9.90E-04 | 349.1 | 149.1 | 927.7 | 43.2 | 112.9 | 47.6 |
| stage2 | stage3_shell | -3.29 | 5.34E-12 | V12B01_10645 | EAP93071.1 | Thymidylate kinase | Nucleoside and Nucleotide Biosynthesis | | - | - | VV2968 | 294.9 | 0.46 | 7.44E-14 | 103.9 | 441.6 | 285.0 | 2068.1 | 3313.9 | 1232.6 |
| stage3_free | stage3_shell | -4.37 | 1.93E-15 | V12B01_10645 | EAP93071.1 | Thymidylate kinase | Nucleoside and Nucleotide Biosynthesis | | - | - | VV2968 | 294.9 | 0.53 | 5.70E-18 | 104.9 | 160.1 | NA | 2068.1 | 3313.9 | 1232.6 |
| stage2 | stage3_shell | -2.49 | 6.74E-04 | V12B01_10655 | EAP93073.1 | hypothetical protein | | | S | Phage shock protein G (Phageshock_PspG) | pspG | 7.2 | 0.76 | 9.23E-05 | 12.0 | 12.8 | 8.4 | 64.8 | 34.1 | 77.0 |
| stage2 | stage3_shell | -2.18 | 5.10E-08 | V12B01_10710 | EAP93084.1 | Outer membrane receptor protein | Outer membrane | | - | - | VPA0901 | 129.2 | 0.39 | 1.86E-09 | 28.9 | 36.9 | 27.9 | 86.2 | 81.6 | 160.5 |
| stage2 | stage3_shell | -2.66 | 1.06E-05 | V12B01_10730 | EAP93088.1 | hypothetical protein | | | - | - | - | 28.9 | 0.60 | 8.14E-07 | 9.1 | 7.2 | 10.4 | 20.7 | 44.8 | 79.1 |
| stage2 | stage3_shell | -2.91 | 7.86E-06 | V12B01_10735 | EAP93089.1 | hypothetical protein | | | - | - | - | 18.8 | 0.65 | 5.75E-07 | 2.8 | 6.7 | 11.8 | 30.1 | 38.1 | 80.6 |
| stage2 | stage3_shell | -2.62 | 2.60E-10 | V12B01_10740 | EAP93090.1 | putative replication initiation protein, phage-related | | | S | Bacteriophage replication gene A protein (GPA) | - | 131.4 | 0.40 | 4.84E-12 | 13.2 | 17.6 | 12.4 | 50.0 | 71.4 | 84.5 |
| stage2 | stage3_shell | -2.73 | 3.83E-04 | V12B01_10745 | EAP93091.1 | hypothetical protein | | | - | #N/A | #N/A | 7.3 | 0.78 | 4.81E-05 | 9.3 | 3.7 | 8.1 | 41.6 | 55.2 | 29.7 |
| stage2 | stage3_shell | -3.15 | 5.70E-05 | V12B01_10755 | EAP93093.1 | hypothetical protein | | | - | #N/A | #N/A | 11.8 | 0.79 | 5.49E-06 | 0.0 | 14.5 | 6.4 | 24.6 | 69.8 | 87.6 |
| stage2 | stage3_shell | -2.04 | 3.88E-10 | V12B01_10765 | EAP93095.1 | putative transcriptional regulator | Regulation | | K | Peptidase S24-like | - | 219.2 | 0.31 | 7.56E-12 | 137.8 | 102.5 | 136.4 | 296.7 | 351.7 | 512.0 |
| stage3_free | stage3_shell | -2.36 | 2.17E-10 | V12B01_10765 | EAP93095.1 | putative transcriptional regulator | Regulation | | K | Peptidase S24-like | - | 219.2 | 0.35 | 2.29E-12 | 104.3 | 97.6 | NA | 296.7 | 351.7 | 512.0 |
| stage2 | stage3_shell | 3.49 | 1.39E-02 | V12B01_10985 | EAP94417.1 | membrane transport protein | Transport Proteins | K07552 | EGP | Sugar (and other) transporter | - | 3.6 | 2.31 | 3.54E-03 | 10.3 | 6.4 | 4.8 | 0.0 | 0.0 | 0.0 |
| stage3_free | stage3_shell | 3.56 | 1.72E-02 | V12B01_10985 | EAP94417.1 | membrane transport protein | Transport Proteins | K07552 | EGP | Sugar (and other) transporter | - | 3.6 | 2.39 | 3.88E-03 | 6.3 | 9.5 | NA | 0.0 | 0.0 | 0.0 |
| stage3_free | stage3_shell | 2.19 | 5.87E-02 | V12B01_11005 | EAP94421.1 | Probable riboflavin biosynthesis protein RibD | Cofactor, Carrier, and Vitamin Biosynthesis | | H | MafB19-like deaminase | - | 4.1 | 2.31 | 1.84E-02 | 13.6 | 13.2 | NA | 0.0 | 0.0 | 0.0 |
| stage3_free | stage3_shell | -2.21 | 8.56E-04 | V12B01_11090 | EAP94438.1 | hypothetical protein | | | - | - | - | 109.7 | 0.67 | 9.55E-05 | 49.9 | 85.6 | NA | 455.5 | 317.2 | 83.6 |
| stage3_free | stage3_shell | -2.16 | 5.32E-02 | V12B01_11100 | EAP94440.1 | nucleoside permease | Transport Proteins | K03317 | U | Belongs to the concentrative nucleoside transporter (CNT) (TC 2.A.41) family | - | 5.8 | 1.60 | 1.63E-02 | 0.0 | 0.8 | NA | 2.6 | 6.3 | 0.0 |
| stage3_free | stage3_shell | -2.05 | 3.04E-02 | V12B01_11175 | EAP94455.1 | hypothetical protein | | | - | #N/A | #N/A | 7.4 | 1.24 | 7.93E-03 | 0.0 | 26.1 | NA | 49.5 | 117.3 | 44.1 |
| stage3_free | stage3_shell | -3.30 | 1.39E-04 | V12B01_11195 | EAP94459.1 | Multidrug resistance efflux pump | Transport Proteins | | V | multidrug resistance efflux pump | - | 13.4 | 0.86 | 1.11E-05 | 2.2 | 0.8 | NA | 18.3 | 9.9 | 14.0 |
| stage2 | stage3_shell | -2.61 | 5.77E-05 | V12B01_11215 | EAP94463.1 | hypothetical protein | | K07005 | S | Flavin-nucleotide-binding protein | - | 22.2 | 0.66 | 5.62E-06 | 4.0 | 8.6 | 5.2 | 9.7 | 27.5 | 60.4 |
| stage3_free | stage3_shell | -2.37 | 1.25E-03 | V12B01_11215 | EAP94463.1 | hypothetical protein | | K07005 | S | Flavin-nucleotide-binding protein | - | 22.2 | 0.76 | 1.50E-04 | 7.6 | 5.8 | NA | 9.7 | 27.5 | 60.4 |
| stage2 | stage3_free | -2.74 | 7.30E-04 | V12B01_11225 | EAP94465.1 | amino acid transporter | Transport Proteins | K16263 | E | COG0531 Amino acid transporters | - | 43.0 | 0.77 | 2.69E-05 | 8.4 | 8.4 | 9.8 | 80.4 | 75.0 | NA |
| stage3_free | stage3_shell | 2.17 | 7.75E-04 | V12B01_11265 | EAP94473.1 | glyceraldehyde-3-phosphate dehydrogenase | Glycolysis | K00134 | C | Belongs to the glyceraldehyde-3-phosphate dehydrogenase family | gapA-2 | 256.6 | 0.63 | 8.44E-05 | 678.2 | 705.3 | NA | 109.0 | 36.8 | 124.8 |
| stage2 | stage3_shell | -2.86 | 3.49E-17 | V12B01_11270 | EAP94474.1 | hypothetical protein | | | T | COG0642 Signal transduction histidine kinase | - | 356.4 | 0.33 | 1.77E-19 | 22.3 | 45.1 | 36.1 | 135.0 | 272.6 | 168.1 |
| stage3_free | stage3_shell | -3.09 | 1.77E-15 | V12B01_11270 | EAP94474.1 | hypothetical protein | | | T | COG0642 Signal transduction histidine kinase | - | 356.4 | 0.37 | 4.85E-18 | 27.3 | 31.9 | NA | 135.0 | 272.6 | 168.1 |
| stage2 | stage3_shell | -2.29 | 7.46E-08 | V12B01_11280 | EAP94476.1 | hypothetical protein | | | S | Peptidase M15 | - | 134.9 | 0.41 | 2.84E-09 | 36.2 | 33.3 | 29.4 | 214.1 | 115.8 | 54.5 |

| | | | | | | | | | | | | | | | | | | | | |
|---|---|---|---|---|---|---|---|---|---|---|---|---|---|---|---|---|---|---|---|---|
| stage2 | stage3_shell | 2.22 | 2.27E-03 | V12B01_11340 | EAP94488.1 | hypothetical protein | | | Q | COG0179 2-keto-4-pentenoate hydratase 2-oxohepta-3-ene-1,7-dioic acid hydratase (catechol pathway) | - | 15.9 | 0.70 | 3.99E-04 | 50.6 | 34.7 | 46.3 | 5.0 | 9.6 | 0.0 |
| stage3_free | stage3_shell | 2.07 | 6.64E-03 | V12B01_11340 | EAP94488.1 | hypothetical protein | | | Q | COG0179 2-keto-4-pentenoate hydratase 2-oxohepta-3-ene-1,7-dioic acid hydratase (catechol pathway) | - | 15.9 | 0.73 | 1.15E-03 | 42.6 | 39.5 | NA | 5.0 | 9.6 | 0.0 |
| **stage3_free** | **stage3_shell** | **-2.83** | **3.64E-07** | **V12B01_11350** | **EAP94490.1** | **Phosphomannomutase** | **Carbohydrate Biosynthesis** | **K01840** | **G** | **Phosphoglucomutase phosphomannomutase alpha beta subunit** | **-** | **1179.2** | **0.54** | **9.60E-09** | **26.0** | **33.4** | **NA** | **114.9** | **353.4** | **67.7** |
| **stage2** | **stage3_shell** | **-2.20** | **6.25E-04** | **V12B01_11355** | **EAP94491.1** | **cytochrome c oxidase, subunit II** | **Transport Proteins** | **K02275** | **C** | **Subunits I and II form the functional core of the enzyme complex. Electrons originating in cytochrome c are transferred via heme a and Cu(A) to the binuclear center formed by heme a3 and Cu(B)** | **coxB** | **508.8** | **0.65** | **8.48E-05** | **38.8** | **114.6** | **95.1** | **227.9** | **787.9** | **60.3** |
| **stage3_free** | **stage3_shell** | **-4.42** | **5.86E-09** | **V12B01_11355** | **EAP94491.1** | **cytochrome c oxidase, subunit II** | **Respiration** | **K02275** | **C** | **Subunits I and II form the functional core of the enzyme complex. Electrons originating in cytochrome c are transferred via heme a and Cu(A) to the binuclear center formed by heme a3 and Cu(B)** | **coxB** | **508.8** | **0.74** | **9.15E-11** | **16.0** | **21.1** | **NA** | **227.9** | **787.9** | **60.3** |
| stage2 | stage3_free | 2.86 | 1.08E-03 | V12B01_11360 | EAP94492.1 | cytochrome c oxidase, subunit I | Transport Proteins | K02274 | C | Cytochrome c oxidase is the component of the respiratory chain that catalyzes the reduction of oxygen to water. Subunits 1- 3 form the functional core of the enzyme complex. CO I is the catalytic subunit of the enzyme. Electrons originating in cytochrome c are transferred via the copper A center of subunit 2 and heme A of subunit 1 to the bimetallic center formed by heme A3 and copper B | coxA | 541.9 | 0.90 | 4.43E-05 | 29.4 | 87.6 | 54.6 | 3.9 | 7.8 | NA |
| **stage2** | **stage3_shell** | **-2.33** | **9.67E-04** | **V12B01_11360** | **EAP94492.1** | **cytochrome c oxidase, subunit I** | **Transport Proteins** | **K02274** | **C** | **Cytochrome c oxidase is the component of the respiratory chain that catalyzes the reduction of oxygen to water. Subunits 1- 3 form the functional core of the enzyme complex. CO I is the catalytic subunit of the enzyme. Electrons originating in cytochrome c are transferred via the copper A center of subunit 2 and heme A of subunit 1 to the bimetallic center formed by heme A3 and copper B** | **coxA** | **541.9** | **0.73** | **1.43E-04** | **29.4** | **87.6** | **54.6** | **128.8** | **682.7** | **37.7** |
| **stage3_free** | **stage3_shell** | **-5.74** | **2.05E-11** | **V12B01_11360** | **EAP94492.1** | **cytochrome c oxidase, subunit I** | **Respiration** | **K02274** | **C** | **Cytochrome c oxidase is the component of the respiratory chain that catalyzes the reduction of oxygen to water. Subunits 1- 3 form the functional core of the enzyme complex. CO I is the catalytic subunit of the enzyme. Electrons originating in cytochrome c are transferred via the copper A center of subunit 2 and heme A of subunit 1 to the bimetallic center formed by heme A3 and copper B** | **coxA** | **541.9** | **0.84** | **1.82E-13** | **3.9** | **7.8** | **NA** | **128.8** | **682.7** | **37.7** |
| stage2 | stage3_free | 2.36 | 4.51E-03 | V12B01_11365 | EAP94493.1 | cytochrome C oxidase assembly protein | Respiration | K02258 | O | COG3175 Cytochrome oxidase assembly factor | coxG | 254.6 | 0.88 | 2.48E-04 | 29.0 | 91.3 | 91.1 | 2.7 | 17.1 | NA |
| **stage2** | **stage3_shell** | **-2.74** | **5.14E-05** | **V12B01_11365** | **EAP94493.1** | **cytochrome C oxidase assembly protein** | **Respiration** | **K02258** | **O** | **COG3175 Cytochrome oxidase assembly factor** | **coxG** | **254.6** | **0.68** | **4.86E-06** | **29.0** | **91.3** | **91.1** | **201.2** | **977.2** | **122.3** |
| **stage3_free** | **stage3_shell** | **-5.63** | **6.54E-12** | **V12B01_11365** | **EAP94493.1** | **cytochrome C oxidase assembly protein** | **Respiration** | **K02258** | **O** | **COG3175 Cytochrome oxidase assembly factor** | **coxG** | **254.6** | **0.80** | **5.24E-14** | **2.7** | **17.1** | **NA** | **201.2** | **977.2** | **122.3** |
| **stage2** | **stage3_shell** | **-2.41** | **9.96E-05** | **V12B01_11370** | **EAP94494.1** | **cytochrome c oxidase, subunit III** | **Transport Proteins** | **K02276** | **C** | **COG1845 Heme copper-type cytochrome quinol oxidase, subunit 3** | **coxC** | **198.6** | **0.62** | **1.04E-05** | **21.6** | **59.2** | **40.2** | **102.9** | **417.1** | **61.1** |
| **stage3_free** | **stage3_shell** | **-4.53** | **1.31E-09** | **V12B01_11370** | **EAP94494.1** | **cytochrome c oxidase, subunit III** | **Respiration** | **K02276** | **C** | **COG1845 Heme copper-type cytochrome quinol oxidase, subunit 3** | **coxC** | **198.6** | **0.73** | **1.83E-11** | **8.3** | **10.9** | **NA** | **102.9** | **417.1** | **61.1** |
| **stage2** | **stage3_shell** | **-8.24** | **7.43E-06** | **V12B01_11375** | **EAP94495.1** | **hypothetical protein** | | | **S** | **Protein of unknown function (DUF2909)** | **-** | **6.3** | **2.98** | **5.36E-07** | **0.0** | **0.0** | **0.0** | **13.8** | **78.6** | **49.3** |
| **stage3_free** | **stage3_shell** | **-3.39** | **8.37E-03** | **V12B01_11375** | **EAP94495.1** | **hypothetical protein** | | | **S** | **Protein of unknown function (DUF2909)** | **-** | **6.3** | **1.59** | **1.54E-03** | **2.0** | **4.2** | **NA** | **13.8** | **78.6** | **49.3** |
| **stage2** | **stage3_shell** | **-2.98** | **2.12E-09** | **V12B01_11380** | **EAP94496.1** | **putative transmembrane cytochrome oxidase complex biogenesis factortransmembrane protein** | **Respiration** | **K14998** | **S** | **SURF1-like protein** | **VVA1112** | **247.9** | **0.48** | **4.82E-11** | **31.5** | **90.5** | **72.2** | **214.6** | **803.6** | **235.6** |
| **stage3_free** | **stage3_shell** | **-4.87** | **1.29E-16** | **V12B01_11380** | **EAP94496.1** | **putative transmembrane cytochrome oxidase complex biogenesis factortransmembrane protein** | **Respiration** | **K14998** | **S** | **SURF1-like protein** | **VVA1112** | **247.9** | **0.56** | **1.64E-19** | **15.9** | **19.6** | **NA** | **214.6** | **803.6** | **235.6** |
| stage2 | stage3_free | 2.97 | 3.67E-04 | V12B01_11385 | EAP94497.1 | hypothetical protein | Respiration | | S | signal sequence binding | VVA1110 | 104.0 | 0.81 | 1.15E-05 | 21.9 | 64.6 | 29.7 | 4.5 | 3.2 | NA |
| **stage2** | **stage3_shell** | **-2.86** | **9.54E-07** | **V12B01_11385** | **EAP94497.1** | **hypothetical protein** | | | **S** | **signal sequence binding** | **VVA1110** | **104.0** | **0.57** | **5.11E-08** | **21.9** | **64.6** | **29.7** | **196.8** | **397.5** | **126.2** |
| **stage3_free** | **stage3_shell** | **-6.21** | **1.51E-14** | **V12B01_11385** | **EAP94497.1** | **hypothetical protein** | | | **S** | **signal sequence binding** | **VVA1110** | **104.0** | **0.77** | **5.40E-17** | **4.5** | **3.2** | **NA** | **196.8** | **397.5** | **126.2** |

| | | | | | | | | | | | | | | | | | | | | |
|---|---|---|---|---|---|---|---|---|---|---|---|---|---|---|---|---|---|---|---|---|
| stage2 | stage3_shell | -2.24 | 1.45E-03 | V12B01_11390 | EAP94498.1 | putative cytochrome c oxidase assembly protein | Respiration | K02259 | O | protein required for cytochrome oxidase assembly | - | 155.1 | 0.73 | 2.33E-04 | 16.4 | 43.3 | 20.7 | 118.0 | 253.6 | 2.7 |
| stage3_free | stage3_shell | -4.03 | 2.43E-06 | V12B01_11390 | EAP94498.1 | putative cytochrome c oxidase assembly protein | Respiration | K02259 | O | protein required for cytochrome oxidase assembly | - | 155.1 | 0.85 | 8.56E-08 | 7.4 | 8.6 | NA | 118.0 | 253.6 | 2.7 |
| stage2 | stage3_shell | -2.66 | 6.38E-06 | V12B01_11395 | EAP94499.1 | protoheme IX farnesyltransferase | | K02257 | O | Converts heme B (protoheme IX) to heme O by substitution of the vinyl group on carbon 2 of heme B porphyrin ring with a hydroxyethyl farnesyl side group | cyoE | 230.5 | 0.58 | 4.36E-07 | 19.2 | 48.3 | 50.6 | 193.4 | 414.3 | 49.6 |
| stage3_free | stage3_shell | -3.44 | 4.41E-07 | V12B01_11395 | EAP94499.1 | protoheme IX farnesyltransferase | | K02257 | O | Converts heme B (protoheme IX) to heme O by substitution of the vinyl group on carbon 2 of heme B porphyrin ring with a hydroxyethyl farnesyl side group | cyoE | 230.5 | 0.66 | 1.20E-08 | 33.3 | 12.3 | NA | 193.4 | 414.3 | 49.6 |
| stage2 | stage3_shell | 2.06 | 2.36E-05 | V12B01_11425 | EAP94505.1 | 3-deoxy-7-phosphoheptulonate synthase | Other Biosynthesis | K01626 | E | Stereospecific condensation of phosphoenolpyruvate (PEP) and D-erythrose-4-phosphate (E4P) giving rise to 3-deoxy-D- arabino-heptulosonate-7-phosphate (DAHP) | aroG | 502.9 | 0.47 | 2.03E-06 | 858.5 | 499.8 | 466.5 | 127.5 | 48.8 | 108.2 |
| stage3_free | stage3_shell | -2.00 | 3.72E-08 | V12B01_11450 | EAP94510.1 | Cation transport protein | Transport Proteins | K03498 | P | COG0168 Trk-type K transport systems, membrane components | ktrB | 81.6 | 0.34 | 6.76E-10 | 25.1 | 26.0 | NA | 81.8 | 104.0 | 50.0 |
| stage2 | stage3_free | -5.47 | 8.34E-18 | V12B01_11520 | EAP92048.1 | TRAP dicarboxylate family transporter, DctP subunit | Transport Proteins | | Q | COG4663 TRAP-type mannitol chloroaromatic compound transport system, periplasmic component | - | 53.0 | 0.60 | 5.51E-21 | 4.4 | 3.5 | 2.5 | 136.3 | 197.4 | NA |
| stage2 | stage3_shell | -4.48 | 3.41E-14 | V12B01_11520 | EAP92048.1 | TRAP dicarboxylate family transporter, DctP subunit | Transport Proteins | | Q | COG4663 TRAP-type mannitol chloroaromatic compound transport system, periplasmic component | - | 53.0 | 0.57 | 3.17E-16 | 4.4 | 3.5 | 2.5 | 69.1 | 38.7 | 72.8 |
| stage2 | stage3_free | -3.30 | 1.05E-02 | V12B01_11525 | EAP92049.1 | TRAP dicarboxylate family transporter, DctQ subunit | Transport Proteins | | Q | COG4665 TRAP-type mannitol chloroaromatic compound transport system, small permease component | - | 6.9 | 1.36 | 6.90E-04 | 3.2 | 2.5 | 0.0 | 37.7 | 29.3 | NA |
| stage2 | stage3_free | -4.08 | 3.90E-06 | V12B01_11530 | EAP92050.1 | TRAP dicarboxylate transporter, DctM subunit | Transport Proteins | | Q | COG4664 TRAP-type mannitol chloroaromatic compound transport system, large permease component | - | 18.3 | 0.82 | 5.59E-08 | 1.7 | 1.1 | 1.9 | 23.9 | 41.8 | NA |
| stage2 | stage3_shell | -3.38 | 2.37E-05 | V12B01_11530 | EAP92050.1 | TRAP dicarboxylate transporter, DctM subunit | Transport Proteins | | Q | COG4664 TRAP-type mannitol chloroaromatic compound transport system, large permease component | - | 18.3 | 0.82 | 2.04E-06 | 1.7 | 1.1 | 1.9 | 3.6 | 12.6 | 28.0 |
| stage2 | stage3_shell | -2.07 | 6.60E-03 | V12B01_11535 | EAP92051.1 | gluconokinase | Carbohydrates and Carboxylates Degradation | K00851 | F | Gluconokinase | idnK | 9.0 | 0.84 | 1.42E-03 | 2.7 | 2.8 | 13.8 | 12.8 | 24.3 | 45.7 |
| stage2 | stage3_free | -3.29 | 3.20E-10 | V12B01_11555 | EAP93540.1 | di-/tripeptide transporter | Transport Proteins | K03305 | E | COG3104 Dipeptide tripeptide permease | - | 152.0 | 0.49 | 1.55E-12 | 2.4 | 2.9 | 5.4 | 42.2 | 36.4 | NA |
| stage2 | stage3_shell | -5.33 | 1.85E-30 | V12B01_11555 | EAP93540.1 | di-/tripeptide transporter | Transport Proteins | K03305 | E | COG3104 Dipeptide tripeptide permease | - | 152.0 | 0.45 | 7.80E-34 | 2.4 | 2.9 | 5.4 | 112.2 | 90.6 | 129.4 |
| stage3_free | stage3_shell | -2.28 | 1.93E-03 | V12B01_11645 | EAP93558.1 | chemotactic transducer-related protein | | | KT | COG0840 Methyl-accepting chemotaxis protein | - | 81.1 | 0.76 | 2.54E-04 | 2.5 | 6.3 | NA | 9.7 | 47.9 | 3.8 |
| stage2 | stage3_free | 2.00 | 6.58E-04 | V12B01_11685 | EAP93566.1 | hypothetical protein | | K02005 | M | Belongs to the membrane fusion protein (MFP) (TC 8.A.1) family | VVA1500 | 36.9 | 0.57 | 2.33E-05 | 99.2 | 38.4 | 46.5 | 11.3 | 14.8 | NA |
| stage2 | stage3_free | 2.69 | 8.55E-04 | V12B01_11705 | EAP93570.1 | ABC-type antimicrobial peptide transport system, ATPase component | Transport Proteins | K02003 | V | Part of the tripartite efflux system MacAB-TolC. MacB is a non-canonical ABC transporter that contains transmembrane domains (TMD), which form a pore in the inner membrane, and an ATP-binding domain (NBD), which is responsible for energy generation. Confers resistance against macrolides | macB | 26.0 | 0.81 | 3.24E-05 | 69.9 | 74.4 | 79.9 | 6.2 | 11.7 | NA |
| stage2 | stage3_shell | -3.94 | 6.34E-15 | V12B01_11710 | EAP93571.1 | hypothetical protein | | | K | Helix-turn-helix XRE-family like proteins | - | 115.1 | 0.49 | 4.87E-17 | 31.6 | 38.2 | 46.7 | 258.5 | 545.1 | 579.2 |
| stage3_free | stage3_shell | -3.25 | 1.37E-08 | V12B01_11710 | EAP93571.1 | hypothetical protein | | | K | Helix-turn-helix XRE-family like proteins | - | 115.1 | 0.55 | 2.28E-10 | 42.6 | 80.1 | NA | 258.5 | 545.1 | 579.2 |
| stage2 | stage3_shell | -2.90 | 8.61E-07 | V12B01_11735 | EAP93576.1 | hypothetical protein | | | - | - | - | 28.2 | 0.58 | 4.51E-08 | 4.6 | 8.2 | 6.9 | 22.3 | 35.1 | 66.1 |
| stage3_free | stage3_shell | -2.40 | 2.65E-04 | V12B01_11735 | EAP93576.1 | hypothetical protein | | | - | - | - | 28.2 | 0.66 | 2.38E-05 | 6.9 | 11.2 | NA | 22.3 | 35.1 | 66.1 |
| stage2 | stage3_free | -2.55 | 4.02E-05 | V12B01_11740 | EAP93577.1 | hypothetical protein | | | S | Protein of unknown function (DUF819) | - | 21.6 | 0.58 | 8.05E-07 | 10.5 | 10.4 | 9.7 | 62.8 | 78.9 | NA |
| stage3_free | stage3_shell | 2.18 | 1.18E-03 | V12B01_11740 | EAP93577.1 | hypothetical protein | | | S | Protein of unknown function (DUF819) | - | 21.6 | 0.65 | 1.39E-04 | 62.8 | 78.9 | NA | 15.7 | 7.4 | 4.7 |
| stage2 | stage3_shell | -4.01 | 1.36E-03 | V12B01_11770 | EAP93583.1 | alcohol dehydrogenase | | K13954 | C | alcohol dehydrogenase | adhB | 88.1 | 1.46 | 2.17E-04 | 3.9 | 7.7 | 8.4 | 32.0 | 71.1 | 265.1 |
| stage2 | stage3_shell | -3.50 | 1.00E-02 | V12B01_11775 | EAP93584.1 | hypothetical protein | | | - | - | - | 5.9 | 1.94 | 2.33E-03 | 0.0 | 6.9 | 2.0 | 0.0 | 29.7 | 148.9 |
| stage3_free | stage3_shell | -4.02 | 1.70E-02 | V12B01_11810 | EAP93591.1 | hypothetical protein | | | - | #N/A | #N/A | 1.8 | 3.16 | 3.84E-03 | 0.0 | 0.0 | NA | 51.2 | 9.7 | 0.0 |
| stage2 | stage3_free | -2.66 | 9.46E-04 | V12B01_11815 | EAP93592.1 | hypothetical protein | | | F | Protein of unknown function (DUF3029) | yjjI | 276.6 | 0.76 | 3.74E-05 | 1.1 | 6.4 | 5.2 | 28.8 | 41.3 | NA |
| stage2 | stage3_shell | -4.04 | 7.45E-09 | V12B01_11815 | EAP93592.1 | hypothetical protein | | | F | Protein of unknown function (DUF3029) | yjjI | 276.6 | 0.68 | 2.08E-10 | 1.1 | 6.4 | 5.2 | 18.5 | 86.5 | 69.5 |
| stage2 | stage3_shell | -2.86 | 1.19E-05 | V12B01_11820 | EAP93593.1 | pyruvate formate-lyase activating enzyme | | | C | COG1180 Pyruvate-formate lyase-activating enzyme | - | 33.3 | 0.65 | 9.36E-07 | 1.0 | 5.8 | 5.8 | 14.0 | 23.1 | 43.6 |
| stage3_free | stage3_shell | -2.95 | 4.94E-05 | V12B01_11840 | EAP93597.1 | hypothetical protein | | | - | - | - | 80.2 | 0.73 | 3.29E-06 | 108.2 | 51.7 | NA | 807.5 | 593.8 | 275.5 |
| stage3_free | stage3_shell | 2.38 | 3.01E-02 | V12B01_11920 | EAP95898.1 | probable two-component sensor | Regulation | | T | His Kinase A (phosphoacceptor) domain | - | 11.3 | 1.22 | 7.83E-03 | 17.3 | 6.7 | NA | 0.0 | 2.3 | 0.0 |
| stage2 | stage3_shell | 3.24 | 2.12E-02 | V12B01_11945 | EAP95903.1 | fructose-1-phosphate kinase | Carbohydrates and Carboxylates Degradation | K00882 | H | belongs to the carbohydrate kinase PfkB family | fruK | 10.0 | 2.49 | 5.94E-03 | 10.0 | 8.7 | 9.6 | 0.0 | 0.0 | 0.0 |

| | | | | | | | | | | | | | | | | | | | | |
|---|---|---|---|---|---|---|---|---|---|---|---|---|---|---|---|---|---|---|---|---|
| stage2 | stage3_shell | -4.35 | 2.03E-15 | V12B01_11995 | EAP95913.1 | Transposase | | K07486 | L | COG3547 Transposase and inactivated derivatives | - | 74.0 | 0.53 | 1.37E-17 | 3.5 | 12.4 | 8.2 | 87.2 | 126.8 | 169.3 |
| stage3_free | stage3_shell | -2.41 | 4.65E-05 | V12B01_11995 | EAP95913.1 | Transposase | | K07486 | L | COG3547 Transposase and inactivated derivatives | - | 74.0 | 0.59 | 3.06E-06 | 27.2 | 31.4 | NA | 87.2 | 126.8 | 169.3 |
| stage2 | stage3_shell | 2.95 | 1.82E-02 | V12B01_12030 | EAP95920.1 | putative transcriptional regulator, LysR family | Regulation | K03566 | K | Bacterial regulatory helix-turn-helix protein, lysR family | - | 5.9 | 2.02 | 4.93E-03 | 5.9 | 4.7 | 9.4 | 0.0 | 0.0 | 0.0 |
| stage3_free | stage3_shell | 3.87 | 6.49E-03 | V12B01_12030 | EAP95920.1 | putative transcriptional regulator, LysR family | Regulation | K03566 | K | Bacterial regulatory helix-turn-helix protein, lysR family | - | 5.9 | 2.15 | 1.13E-03 | 8.1 | 13.3 | NA | 0.0 | 0.0 | 0.0 |
| stage2 | stage3_shell | 3.63 | 1.10E-02 | V12B01_12160 | EAP95946.1 | Outer membrane receptor protein | Outer membrane | - | - | - | - | 6.3 | 2.28 | 2.62E-03 | 18.5 | 5.1 | 9.7 | 0.0 | 0.0 | 0.0 |
| stage3_free | stage3_shell | -2.34 | 6.65E-06 | V12B01_12265 | EAP95967.1 | hypothetical protein | | | K | negative regulation of transcription, DNA-templated | - | 107.7 | 0.50 | 2.86E-07 | 11.9 | 16.7 | NA | 42.3 | 102.7 | 33.0 |
| stage2 | stage3_shell | -4.47 | 1.00E-05 | V12B01_12280 | EAP95970.1 | hypothetical protein | | | - | - | - | 30.1 | 1.01 | 7.65E-07 | 0.0 | 7.7 | 22.2 | 296.9 | 388.6 | 0.0 |
| stage3_free | stage3_shell | -6.26 | 3.03E-05 | V12B01_12280 | EAP95970.1 | hypothetical protein | | | - | - | - | 30.1 | 1.45 | 1.79E-06 | 2.5 | 2.6 | NA | 296.9 | 388.6 | 0.0 |
| stage2 | stage3_shell | -2.06 | 1.08E-05 | V12B01_12300 | EAP95974.1 | Cytochrome c oxidase, cbb3-type, subunit I | Aerobic Respiration | K00404 | C | Cytochrome C and Quinol oxidase polypeptide I | - | 98.5 | 0.45 | 8.36E-07 | 10.6 | 10.7 | 10.7 | 30.3 | 63.7 | 14.0 |
| stage2 | stage3_shell | -2.92 | 1.18E-06 | V12B01_12305 | EAP95975.1 | cytochrome c554 | Respiration | | C | Cytochrome c | - | 57.7 | 0.58 | 6.47E-08 | 10.3 | 14.6 | 9.7 | 86.7 | 101.6 | 29.4 |
| stage2 | stage3_shell | -1.99 | 6.02E-05 | V12B01_12310 | EAP95976.1 | hypothetical protein | | | S | Protein conserved in bacteria | - | 72.9 | 0.48 | 5.93E-06 | 20.5 | 65.2 | 38.3 | 154.2 | 212.3 | 43.7 |
| stage3_free | stage3_shell | -2.38 | 3.35E-05 | V12B01_12310 | EAP95976.1 | hypothetical protein | | | S | Protein conserved in bacteria | - | 72.9 | 0.56 | 2.07E-06 | 32.1 | 30.5 | NA | 154.2 | 212.3 | 43.7 |
| stage2 | stage3_shell | -2.30 | 1.82E-05 | V12B01_12315 | EAP95977.1 | GGDEF family protein | Regulation | | T | COG2199 FOG GGDEF domain | cdgA | 137.9 | 0.52 | 1.53E-06 | 10.8 | 63.9 | 46.8 | 160.4 | 284.6 | 66.6 |
| stage3_free | stage3_shell | -3.08 | 6.98E-07 | V12B01_12315 | EAP95977.1 | GGDEF family protein | Regulation | | T | COG2199 FOG GGDEF domain | cdgA | 137.9 | 0.60 | 2.09E-08 | 24.5 | 22.5 | NA | 160.4 | 284.6 | 66.6 |
| stage2 | stage3_shell | 2.07 | 1.09E-02 | V12B01_12330 | EAP95980.1 | transporter, LysE family | Transport Proteins | | E | LysE type translocator | - | 14.9 | 0.82 | 2.59E-03 | 34.2 | 27.2 | 28.3 | 0.0 | 4.6 | 8.7 |
| stage3_free | stage3_shell | 3.20 | 3.50E-04 | V12B01_12330 | EAP95980.1 | transporter, LysE family | Transport Proteins | | E | LysE type translocator | - | 14.9 | 0.86 | 3.35E-05 | 79.4 | 46.5 | NA | 0.0 | 4.6 | 8.7 |
| stage2 | stage3_shell | 2.06 | 1.62E-04 | V12B01_12350 | EAP95984.1 | putative acetyltransferase | | K03830 | K | Acetyltransferase (GNAT) domain | - | 39.7 | 0.54 | 1.81E-05 | 173.2 | 92.2 | 120.3 | 13.3 | 15.8 | 35.6 |
| stage3_free | stage3_shell | -3.52 | 4.38E-05 | V12B01_12380 | EAP95990.1 | adenosine deaminase | Nucleoside and Nucleotide Degradation | K21053 | F | Catalyzes the hydrolytic deamination of adenine to hypoxanthine. Plays an important role in the purine salvage pathway and in nitrogen catabolism | - | 18.8 | 0.86 | 2.87E-06 | 0.9 | 3.4 | NA | 14.4 | 30.3 | 22.8 |
| stage3_free | stage3_shell | -2.92 | 9.67E-10 | V12B01_12385 | EAP95991.1 | hypothetical protein | | | K | Crp-like helix-turn-helix domain | - | 301.6 | 0.46 | 1.26E-11 | 70.2 | 95.8 | NA | 523.7 | 635.1 | 345.5 |
| stage3_free | stage3_shell | -3.13 | 1.04E-04 | V12B01_12390 | EAP95992.1 | hypothetical protein | | | - | - | - | 58.0 | 0.82 | 7.89E-06 | 54.7 | 87.6 | NA | 954.6 | 589.9 | 214.9 |
| stage2 | stage3_shell | -2.07 | 5.38E-03 | V12B01_12400 | EAP95994.1 | hypothetical protein | | | T | Histidine kinase | algC | 36.5 | 0.78 | 1.11E-03 | 9.3 | 19.8 | 10.2 | 11.2 | 137.5 | 13.3 |
| stage3_free | stage3_shell | -2.25 | 7.92E-03 | V12B01_12400 | EAP95994.1 | hypothetical protein | | | T | Histidine kinase | algC | 36.5 | 0.94 | 1.44E-03 | 13.9 | 7.8 | NA | 11.2 | 137.5 | 13.3 |
| stage2 | stage3_shell | -2.76 | 3.20E-05 | V12B01_12405 | EAP95995.1 | putative sensor histidine kinase | Regulation | | T | COG0642 Signal transduction histidine kinase | - | 147.7 | 0.66 | 2.86E-06 | 3.7 | 10.8 | 9.5 | 18.4 | 117.8 | 11.7 |
| stage3_free | stage3_shell | -2.43 | 9.89E-04 | V12B01_12405 | EAP95995.1 | putative sensor histidine kinase | Regulation | | T | COG0642 Signal transduction histidine kinase | - | 147.7 | 0.77 | 1.13E-04 | 8.9 | 10.1 | NA | 18.4 | 117.8 | 11.7 |
| stage2 | stage3_shell | -3.08 | 1.21E-07 | V12B01_12410 | EAP95996.1 | putative cytochrome c551 peroxidase | Respiration | K00428 | C | cytochrome c peroxidase | - | 117.2 | 0.57 | 4.94E-09 | 4.1 | 23.3 | 21.7 | 78.8 | 214.2 | 59.8 |
| stage3_free | stage3_shell | -3.18 | 1.98E-06 | V12B01_12410 | EAP95996.1 | putative cytochrome c551 peroxidase | Respiration | K00428 | C | cytochrome c peroxidase | - | 117.2 | 0.65 | 6.85E-08 | 13.4 | 16.7 | NA | 78.8 | 214.2 | 59.8 |
| stage2 | stage3_shell | -2.40 | 2.32E-06 | V12B01_12485 | EAP96011.1 | hypothetical protein | | | M | PFAM Sulphatase-modifying factor | - | 60.4 | 0.49 | 1.39E-07 | 11.3 | 28.3 | 18.6 | 50.6 | 147.6 | 54.1 |
| stage3_free | stage3_shell | -2.47 | 2.21E-05 | V12B01_12485 | EAP96011.1 | hypothetical protein | | | M | PFAM Sulphatase-modifying factor | - | 60.4 | 0.57 | 1.18E-06 | 17.4 | 18.9 | NA | 50.6 | 147.6 | 54.1 |
| stage2 | stage3_shell | -3.06 | 3.66E-08 | V12B01_12520 | EAP96018.1 | hypothetical protein | | | - | - | - | 31.9 | 0.54 | 1.22E-09 | 15.4 | 24.6 | 21.6 | 38.3 | 205.8 | 174.6 |
| stage2 | stage3_shell | -3.05 | 3.22E-06 | V12B01_12525 | EAP96019.1 | hypothetical protein | | K12541 | V | ABC transporter | - | 223.2 | 0.65 | 2.00E-07 | 5.8 | 20.9 | 18.5 | 29.1 | 178.2 | 109.2 |
| stage2 | stage3_free | -2.13 | 1.69E-03 | V12B01_12530 | EAP96020.1 | LapB-like protein | | K12541 | V | ABC transporter transmembrane region | - | 170.4 | 0.65 | 7.87E-05 | 8.7 | 19.4 | 21.8 | 110.1 | 73.6 | NA |
| stage2 | stage3_shell | -2.39 | 5.04E-05 | V12B01_12530 | EAP96020.1 | hypothetical protein | | K12541 | V | ABC transporter transmembrane region | - | 170.4 | 0.58 | 4.74E-06 | 8.7 | 19.4 | 21.8 | 25.2 | 143.0 | 53.8 |
| stage2 | stage3_free | -2.59 | 6.15E-05 | V12B01_12535 | EAP96021.1 | hypothetical ABC transporter outer membrane component | Outer membrane | K12340 | MU | Outer membrane efflux protein | - | 161.8 | 0.60 | 1.36E-06 | 7.0 | 27.2 | 22.4 | 146.5 | 125.6 | NA |
| stage2 | stage3_shell | -2.71 | 3.66E-08 | V12B01_12540 | EAP96022.1 | hypothetical protein | | K20972 | T | cheY-homologous receiver domain | - | 483.5 | 0.48 | 1.21E-09 | 14.1 | 50.6 | 41.9 | 142.5 | 236.6 | 177.3 |
| stage2 | stage3_shell | -4.28 | 7.41E-12 | V12B01_12545 | EAP96023.1 | hypothetical protein | | K12542 | M | HlyD family secretion protein | - | 435.8 | 0.61 | 1.05E-13 | 12.8 | 38.8 | 38.1 | 151.5 | 444.9 | 742.4 |
| stage3_free | stage3_shell | -3.09 | 1.06E-05 | V12B01_12545 | EAP96023.1 | hypothetical protein | | K12542 | M | HlyD family secretion protein | - | 435.8 | 0.70 | 5.06E-07 | 42.3 | 86.8 | NA | 151.5 | 444.9 | 742.4 |
| stage2 | stage3_shell | -3.43 | 5.11E-07 | V12B01_12550 | EAP96024.1 | hypothetical protein | | | - | - | - | 261.7 | 0.67 | 2.51E-08 | 22.0 | 142.5 | 117.8 | 259.7 | 1217.0 | 1043.2 |
| stage3_free | stage3_shell | -2.79 | 2.29E-04 | V12B01_12550 | EAP96024.1 | hypothetical protein | | | - | - | - | 261.7 | 0.78 | 2.00E-05 | 131.0 | 142.9 | NA | 259.7 | 1217.0 | 1043.2 |
| stage2 | stage3_free | -2.94 | 3.73E-05 | V12B01_12555 | EAP96025.1 | hypothetical protein | | K20276 | G | Hep Hag repeat protein | - | 35899.8 | 0.67 | 7.31E-07 | 44.4 | 170.3 | 120.0 | 946.4 | 1171.9 | NA |
| stage2 | stage3_shell | -5.26 | 5.88E-18 | V12B01_12555 | EAP96025.1 | hypothetical protein | | K20276 | G | Hep Hag repeat protein | - | 35899.8 | 0.59 | 2.73E-20 | 44.4 | 170.3 | 120.0 | 1256.3 | 2794.5 | 5487.6 |
| stage2 | stage3_free | -3.13 | 4.82E-05 | V12B01_12565 | EAP96027.1 | hypothetical protein | | | - | - | - | 97.4 | 0.72 | 1.03E-06 | 13.5 | 68.0 | 53.6 | 559.0 | 431.8 | NA |
| stage2 | stage3_shell | -6.08 | 4.21E-20 | V12B01_12565 | EAP96027.1 | hypothetical protein | | | - | #N/A | #N/A | 97.4 | 0.64 | 1.33E-22 | 13.5 | 68.0 | 53.6 | 758.5 | 1894.9 | 4227.7 |
| stage3_free | stage3_shell | -2.49 | 3.03E-04 | V12B01_12565 | EAP96027.1 | hypothetical protein | | | - | #N/A | #N/A | 97.4 | 0.70 | 2.82E-05 | 559.0 | 431.8 | NA | 758.5 | 1894.9 | 4227.7 |
| stage2 | stage3_shell | -4.20 | 5.16E-07 | V12B01_12570 | EAP96028.1 | Transposase | | K07486 | L | COG3547 Transposase and inactivated derivatives | - | 39.6 | 0.83 | 2.55E-08 | 1.5 | 6.8 | 4.6 | 26.9 | 86.6 | 92.7 |
| stage3_free | stage3_shell | -2.19 | 9.13E-03 | V12B01_12570 | EAP96028.1 | Transposase | | K07486 | L | COG3547 Transposase and inactivated derivatives | - | 39.6 | 0.97 | 1.72E-03 | 16.4 | 12.9 | NA | 26.9 | 86.6 | 92.7 |
| stage2 | stage3_shell | -3.61 | 3.69E-06 | V12B01_12600 | EAP96034.1 | hypothetical protein | | | - | #N/A | #N/A | 15.8 | 0.78 | 2.32E-07 | 6.2 | 44.3 | 26.0 | 111.0 | 189.1 | 553.6 |
| stage2 | stage3_shell | -4.24 | 2.70E-11 | V12B01_12605 | EAP96035.1 | hypothetical protein | | | - | #N/A | #N/A | 293.1 | 0.62 | 4.27E-13 | 49.4 | 375.4 | 304.1 | 1707.0 | 5094.6 | 4234.3 |
| stage3_free | stage3_shell | -2.12 | 2.02E-03 | V12B01_12605 | EAP96035.1 | hypothetical protein | | | - | #N/A | #N/A | 293.1 | 0.72 | 2.72E-04 | 969.4 | 927.9 | NA | 1707.0 | 5094.6 | 4234.3 |
| stage2 | stage3_shell | -3.51 | 5.09E-10 | V12B01_12625 | EAP96039.1 | hypothetical protein | | | - | - | - | 831.0 | 0.55 | 1.02E-11 | 379.1 | 819.6 | 685.1 | 1725.8 | 3830.7 | 10394.2 |
| stage3_free | stage3_shell | -3.13 | 8.92E-07 | V12B01_12625 | EAP96039.1 | hypothetical protein | | | - | - | - | 831.0 | 0.63 | 2.82E-08 | 750.5 | 831.4 | NA | 1725.8 | 3830.7 | 10394.2 |
| stage2 | stage3_shell | 3.26 | 1.48E-02 | V12B01_13015 | EAP96117.1 | hypothetical protein | | | - | #N/A | #N/A | 6.6 | 2.18 | 3.84E-03 | 35.9 | 103.7 | 88.3 | 0.0 | 0.0 | 0.0 |
| stage2 | stage3_shell | -2.52 | 4.05E-04 | V12B01_13170 | EAP96148.1 | hypothetical protein | | | - | #N/A | #N/A | 15.0 | 0.72 | 5.12E-05 | 35.6 | 66.1 | 70.7 | 703.0 | 121.0 | 75.9 |
| stage2 | stage3_shell | -2.36 | 2.88E-02 | V12B01_13420 | EAP96198.1 | hypothetical protein | | | - | #N/A | #N/A | 4.1 | 1.89 | 8.60E-03 | 2.7 | 4.4 | 0.0 | 0.0 | 9.3 | 70.3 |
| stage3_free | stage3_shell | -4.46 | 1.11E-02 | V12B01_13420 | EAP96198.1 | hypothetical protein | | | - | #N/A | #N/A | 4.1 | 3.22 | 2.27E-03 | 0.0 | 0.0 | NA | 0.0 | 9.3 | 70.3 |

| | | | | | | | | | | | | | | | | | | | | |
|---|---|---|---|---|---|---|---|---|---|---|---|---|---|---|---|---|---|---|---|---|
| stage3_free | stage3_shell | 2.01 | 2.24E-03 | V12B01_13705 | EAP96255.1 | glycine cleavage system P protein | Amino Acid Degradation | K00281 | E | The glycine cleavage system catalyzes the degradation of glycine. The P protein binds the alpha-amino group of glycine through its pyridoxal phosphate cofactor | gcvP | 330.5 | 0.65 | 3.09E-04 | 289.5 | 131.7 | NA | 12.9 | 27.1 | 49.1 |
| stage3_free | stage3_shell | 3.16 | 1.22E-13 | V12B01_13725 | EAP96259.1 | Glycine/serine hydroxymethyltransferase | Amino Acid Biosynthesis | K00600 | E | Catalyzes the reversible interconversion of serine and glycine with tetrahydrofolate (THF) serving as the one-carbon carrier. This reaction serves as the major source of one-carbon groups required for the biosynthesis of purines, thymidylate, methionine, and other important biomolecules. Also exhibits THF-independent aldolase activity toward beta-hydroxyamino acids, producing glycine and aldehydes, via a retro-aldol mechanism | glyA2 | 213.1 | 0.40 | 6.16E-16 | 808.4 | 400.3 | NA | 26.3 | 66.4 | 46.8 |
| stage2 | stage3_free | -3.57 | 5.56E-06 | V12B01_13740 | EAP96262.1 | hypothetical protein | | | - | - | - | 41.7 | 0.73 | 8.20E-08 | 174.7 | 50.6 | 66.9 | 1851.2 | 966.4 | NA |
| stage3_free | stage3_shell | 3.46 | 7.80E-05 | V12B01_13740 | EAP96262.1 | hypothetical protein | | | - | #N/A | #N/A | 41.7 | 0.84 | 5.51E-06 | 1851.2 | 966.4 | NA | 209.2 | 0.0 | 0.0 |
| stage2 | stage3_free | -2.88 | 4.92E-09 | V12B01_13750 | EAP96264.1 | hypothetical protein | | | - | - | - | 42.8 | 0.46 | 3.79E-11 | 52.9 | 75.9 | 104.0 | 733.2 | 549.0 | NA |
| stage3_free | stage3_shell | 3.34 | 2.14E-06 | V12B01_13750 | EAP96264.1 | hypothetical protein | | | - | #N/A | #N/A | 42.8 | 0.67 | 7.45E-08 | 733.2 | 549.0 | NA | 63.4 | 0.0 | 67.8 |
| stage2 | stage3_free | -3.07 | 1.64E-08 | V12B01_13775 | EAP92740.1 | glycine cleavage system T protein | Amino Acid Degradation | K00605 | H | COG0404 Glycine cleavage system T protein (aminomethyltransferase) | gcvT | 260.4 | 0.51 | 1.44E-10 | 116.5 | 50.9 | 55.6 | 885.7 | 527.4 | NA |
| stage3_free | stage3_shell | 2.44 | 9.52E-06 | V12B01_13775 | EAP92740.1 | glycine cleavage system T protein | Amino Acid Degradation | K00605 | H | COG0404 Glycine cleavage system T protein (aminomethyltransferase) | gcvT | 260.4 | 0.52 | 4.42E-07 | 885.7 | 527.4 | NA | 81.7 | 40.0 | 123.0 |
| stage2 | stage3_shell | -2.05 | 1.14E-05 | V12B01_13880 | EAP92761.1 | transporter, AcrB/D/F family | Transport Proteins | | V | Belongs to the resistance-nodulation-cell division (RND) (TC 2.A.6) family | - | 71.2 | 0.45 | 8.92E-07 | 5.8 | 12.5 | 8.9 | 26.1 | 30.2 | 33.5 |
| stage2 | stage3_shell | -2.89 | 8.53E-03 | V12B01_13910 | EAP92767.1 | CBS domain protein | | K06212 | P | COG2116 Formate nitrite family of transporters | VVA0737 | 22.4 | 1.43 | 1.93E-03 | 2.4 | 1.5 | 4.1 | 0.0 | 4.2 | 66.7 |
| stage2 | stage3_shell | -2.73 | 7.55E-10 | V12B01_13925 | EAP92770.1 | predicted ATP-dependent endonuclease | | K07459 | L | ATP-dependent endonuclease of the OLD family | ybjD | 173.0 | 0.43 | 1.59E-11 | 13.0 | 9.0 | 11.7 | 50.3 | 40.1 | 81.7 |
| stage2 | stage3_shell | 5.15 | 2.69E-04 | V12B01_13935 | EAP92772.1 | hypothetical protein | | | - | - | - | 11.9 | 2.16 | 3.24E-05 | 171.4 | 214.0 | 162.5 | 0.0 | 0.0 | 0.0 |
| stage3_free | stage3_shell | 2.92 | 1.98E-02 | V12B01_13935 | EAP92772.1 | hypothetical protein | | | - | - | - | 11.9 | 1.98 | 4.66E-03 | 61.9 | 65.0 | NA | 0.0 | 0.0 | 0.0 |
| stage2 | stage3_free | 2.62 | 8.54E-06 | V12B01_13990 | EAP93280.1 | Hydrolases of the alpha/beta superfamily | | | I | Bacterial virulence factor lipase N-terminal | - | 79.4 | 0.56 | 1.33E-07 | 173.6 | 58.7 | 48.4 | 12.9 | 14.1 | NA |
| stage2 | stage3_shell | 3.09 | 4.79E-08 | V12B01_13990 | EAP93280.1 | Hydrolases of the alpha/beta superfamily | | | I | Bacterial virulence factor lipase N-terminal | - | 79.4 | 0.54 | 1.71E-09 | 173.6 | 58.7 | 48.4 | 5.1 | 12.0 | 4.5 |
| stage2 | stage3_free | -2.25 | 5.36E-07 | V12B01_14035 | EAP93289.1 | hypothetical protein | | K08984 | S | membrane | - | 65.7 | 0.42 | 6.61E-09 | 13.9 | 13.4 | 16.5 | 74.7 | 79.3 | NA |
| stage2 | stage3_shell | -3.82 | 1.21E-21 | V12B01_14035 | EAP93289.1 | hypothetical protein | | K08984 | S | membrane | - | 65.7 | 0.38 | 3.05E-24 | 13.9 | 13.4 | 16.5 | 187.2 | 167.3 | 122.2 |
| stage2 | stage3_shell | -3.69 | NA | V12B01_14065 | EAP93295.1 | uncharacterized paraquat-inducible protein B | | K00209 | I | Involved in the final reduction of the elongation cycle of fatty acid synthesis (FAS II). Catalyzes the reduction of a carbon-carbon double bond in an enoyl moiety that is covalently linked to an acyl carrier protein (ACP) | fabV2 | 86.7 | 0.92 | NA | 12.2 | 7.7 | 6.8 | 50.6 | 55.5 | 196.9 |
| stage3_free | stage3_shell | 2.03 | 2.11E-04 | V12B01_14155 | EAP93313.1 | D-alanylalanine synthetase | Cell Structure Biosynthesis | K01921 | F | Belongs to the D-alanine--D-alanine ligase family | ddl | 55.9 | 0.52 | 1.80E-05 | 89.5 | 102.7 | NA | 28.9 | 9.1 | 5.7 |
| stage3_free | stage3_shell | -2.02 | 9.83E-06 | V12B01_14245 | EAP92335.1 | hypothetical protein | | | S | Late competence development protein ComFB | - | 248.5 | 0.44 | 4.60E-07 | 163.6 | 182.3 | NA | 493.6 | 824.1 | 370.4 |
| stage2 | stage3_shell | -2.07 | 5.30E-08 | V12B01_14265 | EAP92339.1 | putative regulatory component of sensory transduction system | Regulation | | T | COG3706 Response regulator containing a CheY-like receiver domain and a GGDEF domain | - | 513.3 | 0.36 | 1.97E-09 | 120.6 | 326.6 | 324.5 | 954.3 | 1102.4 | 506.0 |
| stage3_free | stage3_shell | -2.67 | 4.88E-10 | V12B01_14265 | EAP92339.1 | putative regulatory component of sensory transduction system | Regulation | | T | COG3706 Response regulator containing a CheY-like receiver domain and a GGDEF domain | - | 513.3 | 0.41 | 5.77E-12 | 164.5 | 172.7 | NA | 954.3 | 1102.4 | 506.0 |
| stage2 | stage3_shell | 2.36 | 8.86E-05 | V12B01_14270 | EAP92340.1 | hypothetical protein | | | S | Protein of unknown function (DUF3069) | VPA0928 | 67.3 | 0.58 | 9.13E-06 | 266.0 | 315.4 | 295.0 | 76.6 | 27.6 | 0.0 |
| stage3_free | stage3_shell | -2.46 | 4.65E-06 | V12B01_14280 | EAP92342.1 | molybdate transport permease protein | Transport Proteins | K02018 | P | COG4149 ABC-type molybdate transport system, permease component | modB | 51.8 | 0.51 | 1.83E-07 | 7.9 | 25.4 | NA | 50.3 | 123.4 | 48.9 |
| stage2 | stage3_shell | -2.42 | 1.97E-03 | V12B01_14345 | EAP92355.1 | anaerobic ribonucleoside triphosphate reductase | Nucleoside and Nucleotide Biosynthesis | K21636 | F | Ribonucleoside-triphosphate reductase | nrdD | 141.4 | 0.83 | 3.31E-04 | 8.4 | 18.4 | 18.7 | 37.7 | 48.5 | 132.9 |
| stage2 | stage3_free | 2.25 | 1.16E-04 | V12B01_14380 | EAP91778.1 | choline dehydrogenase | | | C | Involved in the biosynthesis of the osmoprotectant glycine betaine. Catalyzes the oxidation of choline to betaine aldehyde and betaine aldehyde to glycine betaine at the same rate | - | 80.6 | 0.56 | 2.84E-06 | 84.9 | 76.8 | 63.4 | 12.2 | 15.1 | NA |
| stage2 | stage3_free | -5.06 | 4.94E-12 | V12B01_14515 | EAP92987.1 | proton/glutamate symporter | Transport Proteins | | U | Belongs to the dicarboxylate amino acid cation symporter (DAACS) (TC 2.A.23) family | - | 252.1 | 0.69 | 1.74E-14 | 33.7 | 23.3 | 27.7 | 1197.2 | 972.5 | NA |
| stage3_free | stage3_shell | 3.18 | 1.38E-05 | V12B01_14515 | EAP92987.1 | proton/glutamate symporter | Transport Proteins | | U | Belongs to the dicarboxylate amino acid cation symporter (DAACS) (TC 2.A.23) family | - | 252.1 | 0.70 | 6.89E-07 | 1197.2 | 972.5 | NA | 83.3 | 11.6 | 113.5 |
| stage2 | stage3_shell | -2.24 | 9.78E-04 | V12B01_14535 | EAP92991.1 | hypothetical protein | | | S | Domain of unknown function (DUF4174) | VPA0983 | 61.7 | 0.70 | 1.45E-04 | 5.6 | 11.5 | 13.0 | 26.1 | 52.1 | 51.6 |
| stage3_free | stage3_shell | -2.84 | 4.77E-04 | V12B01_14535 | EAP92991.1 | hypothetical protein | | | S | Domain of unknown function (DUF4174) | VPA0983 | 61.7 | 0.84 | 4.82E-05 | 3.3 | 9.6 | NA | 26.1 | 52.1 | 51.6 |
| stage2 | stage3_free | 2.49 | 1.29E-07 | V12B01_14555 | EAP92995.1 | nitrite reductase [NAD(P)H] large subunit | Respiration | K00362 | C | Belongs to the nitrite and sulfite reductase 4Fe-4S domain family | nirB | 204.7 | 0.45 | 1.34E-09 | 48.8 | 36.1 | 38.5 | 5.7 | 7.7 | NA |

| | | | | | | | | | | | | | | | | | | | | |
|---|---|---|---|---|---|---|---|---|---|---|---|---|---|---|---|---|---|---|---|---|
| **stage2** | **stage3_shell** | **-2.13** | **2.69E-02** | **V12B01_14575** | **EAP92999.1** | **arsenite oxidase, small subunit** | | **K08355** | **C** | **Rieske [2Fe-2S] domain** | **-** | **22.9** | **1.39** | **7.89E-03** | **19.0** | **5.4** | **5.9** | **57.4** | **2.6** | **116.8** |
| **stage2** | **stage3_shell** | **-2.68** | **1.91E-03** | **V12B01_14580** | **EAP93000.1** | **probable cytochrome-c peroxidase** | | **K00428** | **C** | **COG1858 Cytochrome c peroxidase** | **-** | **31.1** | **0.95** | **3.18E-04** | **3.8** | **9.4** | **5.6** | **17.8** | **16.9** | **80.5** |
| **stage2** | **stage3_shell** | **-2.69** | **5.69E-03** | **V12B01_14585** | **EAP93001.1** | **SCO1/SenC family protein/methylamine utilization protein MauG, putative** | | **K00428** | **C** | **Di-haem cytochrome c peroxidase** | **-** | **24.3** | **1.16** | **1.19E-03** | **6.1** | **3.5** | **1.8** | **15.6** | **8.8** | **55.5** |
| **stage2** | **stage3_shell** | **-2.53** | **3.63E-03** | **V12B01_14615** | **EAP93007.1** | **lipase-related protein** | | **K01046** | **I** | **Lipase (class 3)** | **-** | **10.5** | **0.93** | **6.90E-04** | **0.6** | **5.4** | **3.2** | **28.4** | **26.8** | **0.0** |
| **stage3_free** | **stage3_shell** | **-2.11** | **2.06E-02** | **V12B01_14615** | **EAP93007.1** | **lipase-related protein** | | **K01046** | **I** | **Lipase (class 3)** | **-** | **10.5** | **1.12** | **4.89E-03** | **3.5** | **3.7** | **NA** | **28.4** | **26.8** | **0.0** |
| stage2 | stage3_free | 2.38 | 6.75E-06 | V12B01_14646 | EAP94513.1 | hypothetical protein | | | S | protein containing a von Willebrand factor type A (vWA) domain | yieM | 360.3 | 0.50 | 1.03E-07 | 379.7 | 420.9 | 504.3 | 93.3 | 55.9 | NA |
| stage2 | stage3_free | 4.53 | 9.67E-03 | V12B01_14671 | EAP94518.1 | putative orphan protein | | | - | - | - | 29.0 | 2.89 | 6.22E-04 | 28.8 | 323.6 | 266.2 | 0.0 | 5.7 | NA |
| stage2 | stage3_free | 3.72 | 6.14E-04 | V12B01_14676 | EAP94519.1 | putative orphan protein | | | - | - | - | 59.1 | 1.13 | 2.11E-05 | 103.6 | 576.8 | 507.6 | 12.9 | 28.9 | NA |
| stage2 | stage3_free | 4.01 | 2.91E-02 | V12B01_14681 | EAP94520.1 | putative orphan protein | | K07346 | M | Chaperone | - | 3.9 | 1.79 | 2.63E-03 | 8.6 | 16.6 | 12.9 | 0.0 | 0.7 | NA |
| **stage3_free** | **stage3_shell** | **-2.21** | **2.68E-04** | **V12B01_14766** | **EAP94537.1** | **transposase** | | | **L** | **User locus_tag** | **-** | **80.9** | **0.60** | **2.42E-05** | **32.0** | **33.6** | **NA** | **149.1** | **148.8** | **90.2** |
| stage2 | stage3_shell | 2.40 | 3.89E-02 | V12B01_14871 | EAP94558.1 | hypothetical protein | | K11897 | S | Gene 25-like lysozyme | - | 4.6 | 2.03 | 1.23E-02 | 11.6 | 10.9 | 13.3 | 0.0 | 0.0 | 0.0 |
| **stage2** | **stage3_free** | **-2.76** | **1.71E-06** | **V12B01_14926** | **EAP94569.1** | **putative acetyltransferase** | | | **K** | **COG0454 Histone acetyltransferase HPA2 and related acetyltransferases** | **-** | **147.0** | **0.54** | **2.26E-08** | **62.7** | **85.6** | **60.6** | **589.7** | **496.2** | **NA** |
| **stage2** | **stage3_shell** | **-2.98** | **1.78E-10** | **V12B01_14956** | **EAP94575.1** | **PTS family enzyme IIA (N-terminal); enzyme IIBC (C-terminal),induction of ompC** | **Transport Proteins** | **K11200** | **G** | **Phosphoenolpyruvate-dependent sugar phosphotransferase system, EIIA 2** | **-** | **75.1** | **0.45** | **3.19E-12** | **4.6** | **9.2** | **10.0** | **30.6** | **60.2** | **57.4** |
| **stage3_free** | **stage3_shell** | **-3.17** | **5.09E-09** | **V12B01_14956** | **EAP94575.1** | **PTS family enzyme IIA (N-terminal); enzyme IIBC (C-terminal),induction of ompC** | **Transport Proteins** | **K11200** | **G** | **Phosphoenolpyruvate-dependent sugar phosphotransferase system, EIIA 2** | **-** | **75.1** | **0.52** | **7.83E-11** | **5.2** | **8.7** | **NA** | **30.6** | **60.2** | **57.4** |
| **stage2** | **stage3_shell** | **-3.59** | **NA** | **V12B01_15011** | **EAP94586.1** | **C4-dicarboxylate transporter, anaerobic** | **Transport Proteins** | **K03326** | **C** | **C4-dicarboxylate transporter** | **dcuC** | **37.9** | **1.18** | **NA** | **4.5** | **3.9** | **1.9** | **10.8** | **12.8** | **99.0** |
| **stage3_free** | **stage3_shell** | **-2.19** | **3.03E-05** | **V12B01_15026** | **EAP94589.1** | **Signal transduction histidine kinase, nitrate/nitrite-specific** | **Regulation** | **K07674** | **T** | **COG3850 Signal transduction histidine kinase, nitrate nitrite-specific** | **narQ** | **162.2** | **0.51** | **1.78E-06** | **26.7** | **41.6** | **NA** | **105.2** | **151.2** | **123.4** |
| **stage3_free** | **stage3_shell** | **-2.51** | **2.32E-06** | **V12B01_15031** | **EAP94590.1** | **iron-sulfur cluster-binding protein NapF** | | **K02572** | **C** | **COG1145 Ferredoxin** | **napF** | **195.4** | **0.51** | **8.11E-08** | **94.3** | **93.3** | **NA** | **210.7** | **754.6** | **334.6** |
| stage3_free | stage3_shell | 2.50 | 5.15E-02 | V12B01_15111 | EAP94606.1 | polyhydroxyalkanoic acid synthase | Carbon storage | | I | Polyhydroxyalkanoic acid synthase | phaC | 3.5 | 2.04 | 1.55E-02 | 54.2 | 27.3 | NA | 0.0 | 3.7 | 0.0 |
| **stage2** | **stage3_free** | **-5.14** | **5.40E-15** | **V12B01_15121** | **EAP94608.1** | **acetyl-CoA acetyltransferase** | **Carbon storage** | **K00626** | **I** | **Belongs to the thiolase family** | **phbA** | **27.8** | **0.61** | **5.97E-18** | **1.7** | **4.9** | **4.9** | **198.2** | **108.8** | **NA** |
| stage3_free | stage3_shell | 2.87 | 2.35E-05 | V12B01_15121 | EAP94608.1 | acetyl-CoA acetyltransferase | Carbon storage | K00626 | I | Belongs to the thiolase family | phbA | 27.8 | 0.66 | 1.29E-06 | 198.2 | 108.8 | NA | 11.0 | 11.9 | 16.7 |
| **stage2** | **stage3_free** | **-4.70** | **2.62E-13** | **V12B01_15126** | **EAP91666.1** | **acetoacetyl-CoA reductase** | **Carbon storage** | **K00023** | **IQ** | **COG1028 Dehydrogenases with different specificities (related to short-chain alcohol dehydrogenases)** | **phbB** | **40.4** | **0.61** | **8.07E-16** | **7.8** | **13.9** | **7.2** | **304.3** | **260.0** | **NA** |
| **stage2** | **stage3_shell** | **-2.47** | **4.20E-05** | **V12B01_15126** | **EAP91666.1** | **acetoacetyl-CoA reductase** | **Carbon storage** | **K00023** | **IQ** | **COG1028 Dehydrogenases with different specificities (related to short-chain alcohol dehydrogenases)** | **phbB** | **40.4** | **0.59** | **3.85E-06** | **7.8** | **13.9** | **7.2** | **88.4** | **28.6** | **19.2** |
| **stage2** | **stage3_shell** | **-3.00** | **8.97E-03** | **V12B01_15136** | **EAP91668.1** | **hypothetical protein** | | | **-** | **-** | **-** | **4.1** | **1.37** | **2.04E-03** | **0.0** | **0.0** | **11.4** | **11.7** | **77.3** | **41.6** |
| **stage3_free** | **stage3_shell** | **-3.66** | **2.21E-02** | **V12B01_15136** | **EAP91668.1** | **hypothetical protein** | | | **-** | **-** | **-** | **4.1** | **1.94** | **5.36E-03** | **0.0** | **3.5** | **NA** | **11.7** | **77.3** | **41.6** |
| **stage2** | **stage3_shell** | **-2.60** | **3.09E-06** | **V12B01_15141** | **EAP92184.1** | **hypothetical protein** | | | **-** | **-** | **-** | **1867.9** | **0.55** | **1.90E-07** | **951.3** | **3305.7** | **2617.6** | **23629.2** | **8170.8** | **4228.8** |
| **stage3_free** | **stage3_shell** | **-2.12** | **6.80E-04** | **V12B01_15141** | **EAP92184.1** | **hypothetical protein** | | | **-** | **-** | **-** | **1867.9** | **0.63** | **7.22E-05** | **2518.3** | **3628.1** | **NA** | **23629.2** | **8170.8** | **4228.8** |
| stage2 | stage3_shell | 2.13 | 4.18E-03 | V12B01_15321 | EAP94091.1 | Putative translation initiation inhibitor, yjgF family | | | J | translation initiation inhibitor, yjgF family | - | 17.5 | 0.72 | 8.22E-04 | 87.0 | 62.4 | 67.1 | 0.0 | 25.0 | 0.0 |
| stage3_free | stage3_shell | 2.14 | 3.12E-04 | V12B01_15346 | EAP94096.1 | ribosomal-protein-alanine acetyltransferase | | K03790 | J | COG1670 Acetyltransferases, including N-acetylases of ribosomal proteins | - | 33.3 | 0.56 | 2.91E-05 | 108.4 | 99.0 | NA | 12.3 | 20.4 | 11.0 |
| stage2 | stage3_shell | 3.72 | 5.51E-05 | V12B01_15351 | EAP94097.1 | Acetyltransferase | | | S | maltose O-acetyltransferase activity | - | 28.6 | 0.86 | 5.29E-06 | 90.8 | 79.2 | 106.7 | 0.0 | 3.3 | 12.2 |
| stage3_free | stage3_shell | 3.93 | 4.21E-05 | V12B01_15351 | EAP94097.1 | Acetyltransferase | | | S | maltose O-acetyltransferase activity | - | 28.6 | 0.89 | 2.73E-06 | 104.3 | 111.6 | NA | 0.0 | 3.3 | 12.2 |
| stage2 | stage3_shell | 2.87 | 2.30E-02 | V12B01_15361 | EAP94099.1 | hypothetical protein | | | - | - | - | 4.6 | 2.11 | 6.54E-03 | 13.7 | 21.8 | 16.6 | 0.0 | 0.0 | 0.0 |
| **stage2** | **stage3_shell** | **-2.61** | **1.41E-08** | **V12B01_15406** | **EAP94108.1** | **hypothetical protein** | | | **O** | **ADP-ribosylglycohydrolase** | **-** | **123.3** | **0.45** | **4.33E-10** | **33.4** | **44.6** | **30.4** | **84.5** | **150.5** | **271.2** |
| **stage3_free** | **stage3_shell** | **-3.32** | **4.88E-10** | **V12B01_15406** | **EAP94108.1** | **hypothetical protein** | | | **O** | **ADP-ribosylglycohydrolase** | **-** | **123.3** | **0.51** | **5.70E-12** | **17.9** | **26.4** | **NA** | **84.5** | **150.5** | **271.2** |
| **stage3_free** | **stage3_shell** | **-2.00** | **6.78E-03** | **V12B01_15416** | **EAP94110.1** | **putative transcriptional regulator, AraC/XylS family** | **Regulation** | | **K** | **COG2207 AraC-type DNA-binding domain-containing proteins** | **-** | **29.7** | **0.79** | **1.20E-03** | **8.1** | **3.4** | **NA** | **29.7** | **10.7** | **25.2** |
| **stage2** | **stage3_shell** | **-3.65** | **2.75E-03** | **V12B01_15451** | **EAP94117.1** | **hypothetical protein** | | | **-** | **-** | **-** | **17.5** | **1.49** | **5.02E-04** | **8.6** | **13.7** | **0.0** | **24.8** | **52.8** | **264.9** |
| **stage3_free** | **stage3_shell** | **-2.54** | **2.05E-02** | **V12B01_15451** | **EAP94117.1** | **hypothetical protein** | | | **-** | **-** | **-** | **17.5** | **2.33** | **4.86E-03** | **7.1** | **13.0** | **NA** | **24.8** | **52.8** | **264.9** |
| **stage2** | **stage3_shell** | **-2.49** | **2.07E-07** | **V12B01_15456** | **EAP94118.1** | **hypothetical protein** | | | **-** | **-** | **-** | **212.7** | **0.46** | **9.05E-09** | **60.4** | **127.5** | **240.8** | **769.3** | **1005.5** | **228.5** |
| **stage3_free** | **stage3_shell** | **-2.81** | **2.97E-07** | **V12B01_15456** | **EAP94118.1** | **hypothetical protein** | | | **-** | **-** | **-** | **212.7** | **0.53** | **7.58E-09** | **106.3** | **120.6** | **NA** | **769.3** | **1005.5** | **228.5** |
| **stage3_free** | **stage3_shell** | **-2.02** | **3.64E-05** | **V12B01_15486** | **EAP94124.1** | **VCBS repeat protein** | **Membrane** | **K20276** | **Q** | **COG2931, RTX toxins and related Ca2 -binding proteins** | **-** | **1729.8** | **0.47** | **2.30E-06** | **28.7** | **37.8** | **NA** | **46.1** | **133.2** | **134.2** |
| **stage3_free** | **stage3_shell** | **-2.03** | **7.09E-06** | **V12B01_15751** | **EAP96587.1** | **response regulator/phosphatase** | **Regulation** | **K07814** | **T** | **COG3437 Response regulator containing a CheY-like receiver domain and an HD-GYP domain** | **-** | **164.3** | **0.43** | **3.12E-07** | **28.1** | **33.2** | **NA** | **56.1** | **176.9** | **66.6** |
| **stage3_free** | **stage3_shell** | **-2.22** | **2.81E-04** | **V12B01_15816** | **EAP96600.1** | **Sensory box/GGDEF family protein** | **Regulation** | | **T** | **PAS domain** | **-** | **25.1** | **0.60** | **2.56E-05** | **9.0** | **6.8** | **NA** | **27.1** | **45.6** | **21.4** |
| **stage3_free** | **stage3_shell** | **-2.02** | **6.15E-02** | **V12B01_15826** | **EAP96602.1** | **hypothetical protein** | | | **-** | **-** | **-** | **3.4** | **1.91** | **1.95E-02** | **2.0** | **0.0** | **NA** | **20.5** | **6.5** | **0.0** |
| **stage2** | **stage3_shell** | **-2.07** | **3.55E-07** | **V12B01_15831** | **EAP96603.1** | **hypothetical protein** | | **K07267** | **M** | **Carbohydrate-selective porin, OprB family** | **-** | **118.3** | **0.39** | **1.67E-08** | **21.6** | **33.6** | **27.3** | **52.7** | **129.8** | **89.3** |
| **stage3_free** | **stage3_shell** | **-2.32** | **6.48E-07** | **V12B01_15831** | **EAP96603.1** | **hypothetical protein** | | **K07267** | **M** | **Carbohydrate-selective porin, OprB family** | **-** | **118.3** | **0.45** | **1.89E-08** | **17.0** | **28.9** | **NA** | **52.7** | **129.8** | **89.3** |
| **stage3_free** | **stage3_shell** | **-2.58** | **7.95E-03** | **V12B01_15851** | **EAP96607.1** | **hypothetical membrane spanning protein** | **Membrane** | | **S** | **Chlorhexidine efflux transporter** | **-** | **7.0** | **1.07** | **1.44E-03** | **6.4** | **0.0** | **NA** | **26.1** | **24.7** | **13.3** |
| stage3_free | stage3_shell | 2.12 | 5.29E-05 | V12B01_15871 | EAP96611.1 | methyl-accepting chemotaxis protein | Chemotaxis | K03406 | NT | helical bimodular (HBM) domain | - | 89.7 | 0.50 | 3.55E-06 | 109.4 | 85.7 | NA | 8.2 | 21.0 | 14.6 |

| | | | | | | | | | | | | | | | | | | | |
|---|---|---|---|---|---|---|---|---|---|---|---|---|---|---|---|---|---|---|---|
| **stage3_free** | **stage3_shell** | **-2.11** | **5.99E-03** | **V12B01_15916** | **EAP96620.1** | **hypothetical protein** | | | **S** | **Domain of unknown function (DUF4382)** | **-** | **187.4** | **0.81** | **1.01E-03** | **1.3** | **8.9** | **NA** | **31.1** | **32.4** | **0.0** |
| stage2 | stage3_shell | 2.08 | 1.69E-02 | V12B01_15921 | EAP96621.1 | hypothetical protein | | | S | Protein of unknown function (DUF2999) | - | 8.9 | 0.90 | 4.51E-03 | 90.0 | 73.1 | 65.6 | 12.7 | 12.0 | 0.0 |
| **stage2** | **stage3_shell** | **-2.51** | **3.87E-06** | **V12B01_15936** | **EAP96624.1** | **hypothetical protein** | | | **M** | **Spondin_N** | **-** | **145.7** | **0.53** | **2.45E-07** | **68.6** | **113.6** | **93.0** | **479.3** | **747.0** | **118.6** |
| **stage3_free** | **stage3_shell** | **-2.35** | **1.16E-04** | **V12B01_15936** | **EAP96624.1** | **hypothetical protein** | | | **M** | **Spondin_N** | **-** | **145.7** | **0.60** | **8.97E-06** | **59.1** | **139.1** | **NA** | **479.3** | **747.0** | **118.6** |
| **stage2** | **stage3_shell** | **-2.21** | **1.28E-02** | **V12B01_15941** | **EAP96625.1** | **ISPsy11, transposase OrfA** | | **K07483** | **L** | **Evidence 2b Function of strongly homologous gene** | **-** | **11.3** | **1.03** | **3.16E-03** | **2.2** | **12.3** | **14.7** | **115.3** | **41.4** | **0.0** |
| **stage3_free** | **stage3_shell** | **-2.36** | **1.64E-02** | **V12B01_15941** | **EAP96625.1** | **ISPsy11, transposase OrfA** | | **K07483** | **L** | **Evidence 2b Function of strongly homologous gene** | **-** | **11.3** | **1.28** | **3.65E-03** | **1.1** | **14.3** | **NA** | **115.3** | **41.4** | **0.0** |
| **stage2** | **stage3_free** | **-3.30** | **2.67E-03** | **V12B01_15971** | **EAP96631.1** | **hypothetical protein** | | | **-** | **-** | **-** | **186.7** | **1.08** | **1.30E-04** | **47.0** | **167.0** | **204.2** | **2121.3** | **2069.9** | **NA** |
| **stage2** | **stage3_shell** | **-2.78** | **1.97E-03** | **V12B01_15971** | **EAP96631.1** | **hypothetical protein** | | | **-** | **-** | **-** | **186.7** | **0.98** | **3.31E-04** | **47.0** | **167.0** | **204.2** | **2760.3** | **73.8** | **266.2** |
| **stage2** | **stage3_shell** | **-2.08** | **2.60E-03** | **V12B01_15981** | **EAP96633.1** | **ISPsy11, transposase OrfA** | | **K07483** | **L** | **Evidence 2b Function of strongly homologous gene** | **-** | **11.6** | **0.71** | **4.69E-04** | **12.2** | **12.3** | **17.1** | **87.4** | **67.7** | **7.1** |
| **stage3_free** | **stage3_shell** | **-2.68** | **1.20E-03** | **V12B01_15981** | **EAP96633.1** | **ISPsy11, transposase OrfA** | | **K07483** | **L** | **Evidence 2b Function of strongly homologous gene** | **-** | **11.6** | **0.86** | **1.43E-04** | **6.8** | **10.8** | **NA** | **87.4** | **67.7** | **7.1** |
| **stage2** | **stage3_shell** | **-2.05** | **4.01E-03** | **V12B01_15986** | **EAP96634.1** | **Integrase, catalytic region** | | **K07497** | **L** | **Evidence 2b Function of strongly homologous gene** | **-** | **10.7** | **0.73** | **7.81E-04** | **1.1** | **2.6** | **6.2** | **11.9** | **22.5** | **3.5** |
| stage2 | stage3_shell | 2.63 | 2.98E-03 | V12B01_16001 | EAP96637.1 | hypothetical protein | | | CH | COG0654 2-polyprenyl-6-methoxyphenol hydroxylase and related FAD-dependent oxidoreductases | hpxO | 17.4 | 0.87 | 5.54E-04 | 30.9 | 15.2 | 25.6 | 2.8 | 2.7 | 0.0 |
| **stage2** | **stage3_shell** | **-2.38** | **6.52E-06** | **V12B01_16016** | **EAP96640.1** | **putative lactoylglutathione lyase** | | | **E** | **COG0346 Lactoylglutathione lyase and related lyases** | **-** | **232.2** | **0.51** | **4.51E-07** | **87.9** | **131.6** | **127.8** | **779.9** | **677.2** | **94.9** |
| **stage3_free** | **stage3_shell** | **-3.02** | **6.61E-07** | **V12B01_16016** | **EAP96640.1** | **putative lactoylglutathione lyase** | | | **E** | **COG0346 Lactoylglutathione lyase and related lyases** | **-** | **232.2** | **0.59** | **1.95E-08** | **56.6** | **91.5** | **NA** | **779.9** | **677.2** | **94.9** |
| stage2 | stage3_shell | 2.56 | 1.18E-02 | V12B01_16026 | EAP96642.1 | putative ferrichrome ABC transporter (permease) | Transport Proteins | K23182 | P | Belongs to the binding-protein-dependent transport system permease family. FecCD subfamily | - | 32.1 | 1.07 | 2.85E-03 | 43.5 | 17.1 | 8.0 | 4.9 | 0.0 | 0.0 |
| stage2 | stage3_shell | 2.66 | 1.08E-02 | V12B01_16041 | EAP96645.1 | ferric vibrioferrin receptor | Regulation | K16091 | P | Outer membrane protein beta-barrel family | - | 395.4 | 1.34 | 2.56E-03 | 329.8 | 28.6 | 14.1 | 3.0 | 7.1 | 13.4 |
| **stage3_free** | **stage3_shell** | **-2.32** | **1.09E-02** | **V12B01_16051** | **EAP96647.1** | **hypothetical protein** | | | **S** | **ATP-grasp domain** | **-** | **216.8** | **1.12** | **2.20E-03** | **7.5** | **17.8** | **NA** | **63.3** | **13.7** | **124.5** |
| stage2 | stage3_shell | 2.81 | 7.27E-03 | V12B01_16066 | EAP96650.1 | putative AcsD | | | Q | IucA / IucC family | iucA | 46.1 | 1.14 | 1.60E-03 | 46.9 | 5.5 | 2.7 | 1.7 | 1.6 | 0.0 |
| **stage2** | **stage3_shell** | **-2.06** | **3.78E-05** | **V12B01_16116** | **EAP96660.1** | **Cysteine synthase** | **Amino Acid Biosynthesis** | **K01738** | **E** | **Pyridoxal-phosphate dependent enzyme** | **-** | **249.8** | **0.48** | **3.45E-06** | **33.7** | **77.2** | **87.2** | **316.7** | **308.0** | **74.4** |
| **stage3_free** | **stage3_shell** | **-2.37** | **3.11E-05** | **V12B01_16116** | **EAP96660.1** | **Cysteine synthase** | **Amino Acid Biosynthesis** | **K01738** | **E** | **Pyridoxal-phosphate dependent enzyme** | **-** | **249.8** | **0.56** | **1.87E-06** | **43.3** | **61.9** | **NA** | **316.7** | **308.0** | **74.4** |
| **stage2** | **stage3_shell** | **-2.80** | **6.60E-08** | **V12B01_16121** | **EAP96661.1** | **hypothetical protein** | | | **-** | **-** | **-** | **172.1** | **0.50** | **2.49E-09** | **41.2** | **147.6** | **138.7** | **414.2** | **1155.7** | **316.4** |
| **stage3_free** | **stage3_shell** | **-2.87** | **1.15E-06** | **V12B01_16121** | **EAP96661.1** | **hypothetical protein** | | | **-** | **-** | **-** | **172.1** | **0.57** | **3.84E-08** | **116.5** | **86.7** | **NA** | **414.2** | **1155.7** | **316.4** |
| **stage2** | **stage3_shell** | **-2.86** | **3.85E-08** | **V12B01_16151** | **EAP96667.1** | **hypothetical protein** | | | **S** | **Protein of unknown function (DUF3024)** | **-** | **65.4** | **0.50** | **1.30E-09** | **24.5** | **46.8** | **42.1** | **96.8** | **350.1** | **219.6** |
| **stage3_free** | **stage3_shell** | **-2.33** | **6.21E-05** | **V12B01_16151** | **EAP96667.1** | **hypothetical protein** | | | **S** | **Protein of unknown function (DUF3024)** | **-** | **65.4** | **0.57** | **4.23E-06** | **50.4** | **55.6** | **NA** | **96.8** | **350.1** | **219.6** |
| stage2 | stage3_shell | 3.07 | 5.89E-04 | V12B01_16181 | EAP96673.1 | hypothetical protein | | | M | Mechanosensitive ion channel | - | 19.8 | 0.85 | 7.90E-05 | 58.6 | 30.7 | 35.5 | 3.8 | 3.6 | 0.0 |
| stage3_free | stage3_shell | 3.19 | 6.81E-04 | V12B01_16181 | EAP96673.1 | hypothetical protein | | | M | Mechanosensitive ion channel | - | 19.8 | 0.89 | 7.26E-05 | 50.5 | 42.7 | NA | 3.8 | 3.6 | 0.0 |
| **stage2** | **stage3_shell** | **-2.22** | **3.85E-06** | **V12B01_16196** | **EAP96676.1** | **hypothetical protein** | | | **-** | **-** | **-** | **263.2** | **0.46** | **2.43E-07** | **30.2** | **96.4** | **118.9** | **304.3** | **391.1** | **232.4** |
| **stage3_free** | **stage3_shell** | **-2.98** | **2.63E-07** | **V12B01_16196** | **EAP96676.1** | **hypothetical protein** | | | **-** | **-** | **-** | **263.2** | **0.56** | **6.54E-09** | **34.2** | **62.1** | **NA** | **304.3** | **391.1** | **232.4** |
| **stage3_free** | **stage3_shell** | **-2.24** | **3.41E-04** | **V12B01_16211** | **EAP96679.1** | **ferredoxin** | | **K05524** | **C** | **Ferredoxins are iron-sulfur proteins that transfer electrons in a wide variety of metabolic reactions** | **fdxA** | **41.3** | **0.62** | **3.22E-05** | **22.6** | **65.2** | **NA** | **221.9** | **289.5** | **35.2** |
| **stage3_free** | **stage3_shell** | **-6.61** | **8.11E-04** | **V12B01_16271** | **EAP96691.1** | **hypothetical protein** | | | **-** | **#N/A** | **#N/A** | **5.3** | **2.93** | **8.93E-05** | **0.0** | **0.0** | **NA** | **0.0** | **72.0** | **116.2** |
| **stage2** | **stage3_shell** | **-2.18** | **4.83E-07** | **V12B01_16376** | **EAP96712.1** | **multidrug resistance protein D** | | | **EGP** | **COG0477 Permeases of the major facilitator superfamily** | **-** | **42.3** | **0.41** | **2.35E-08** | **7.4** | **15.6** | **10.9** | **38.4** | **52.7** | **30.7** |
| **stage2** | **stage3_shell** | **-2.74** | **3.20E-09** | **V12B01_16386** | **EAP96714.1** | **GGDEF family protein** | **Regulation** | | **T** | **GGDEF family** | **-** | **1553.6** | **0.44** | **7.90E-11** | **32.9** | **80.9** | **72.5** | **304.1** | **554.5** | **159.5** |
| **stage3_free** | **stage3_shell** | **-2.41** | **4.50E-06** | **V12B01_16386** | **EAP96714.1** | **GGDEF family protein** | **Regulation** | | **T** | **GGDEF family** | **-** | **1553.6** | **0.51** | **1.74E-07** | **82.1** | **71.0** | **NA** | **304.1** | **554.5** | **159.5** |
| **stage2** | **stage3_free** | **-2.56** | **2.62E-13** | **V12B01_16416** | **EAP96720.1** | **hypothetical protein** | | | **S** | **protein conserved in bacteria** | **-** | **95.4** | **0.33** | **7.33E-16** | **13.6** | **20.2** | **16.2** | **91.7** | **118.1** | **NA** |
| **stage2** | **stage3_shell** | **-2.43** | **2.58E-13** | **V12B01_16416** | **EAP96720.1** | **hypothetical protein** | | | **S** | **protein conserved in bacteria** | **-** | **95.4** | **0.32** | **2.72E-15** | **13.6** | **20.2** | **16.2** | **104.9** | **64.0** | **37.4** |
| stage3_free | stage3_shell | 2.06 | 3.18E-05 | V12B01_16541 | EAP96745.1 | NAD(+) synthetase | Cofactor, Carrier, and Vitamin Biosynthesis | K01916 | F | Catalyzes the ATP-dependent amidation of deamido-NAD to form NAD. Uses ammonia as a nitrogen source | nadE | 85.7 | 0.47 | 1.91E-06 | 188.2 | 183.8 | NA | 42.3 | 18.2 | 27.4 |
| stage3_free | stage3_shell | 2.27 | 1.99E-04 | V12B01_16556 | EAP96748.1 | dihydroorotase | Nucleoside and Nucleotide Biosynthesis | K01465 | F | Catalyzes the reversible cyclization of carbamoyl aspartate to dihydroorotate | pyrC | 84.4 | 0.58 | 1.67E-05 | 209.8 | 261.4 | NA | 57.5 | 11.8 | 22.1 |
| **stage3_free** | **stage3_shell** | **-2.04** | **1.07E-06** | **V12B01_16561** | **EAP96749.1** | **hypothetical protein** | | **K09781** | **S** | **protein conserved in bacteria** | **yedI** | **173.3** | **0.40** | **3.51E-08** | **73.2** | **81.1** | **NA** | **335.1** | **335.6** | **94.3** |
| **stage2** | **stage3_free** | **-3.25** | **4.83E-05** | **V12B01_16571** | **EAP96751.1** | **putative transporter, BCCT family** | **Transport Proteins** | **K03451** | **P** | **Belongs to the BCCT transporter (TC 2.A.15) family** | **-** | **112.6** | **0.75** | **1.04E-06** | **35.4** | **23.1** | **16.1** | **259.1** | **334.6** | **NA** |
| stage2 | stage3_free | 2.78 | 5.87E-04 | V12B01_16596 | EAP96756.1 | TRAP-T family transporter, DctP subunit | Transport Proteins | K21395 | G | TRAP-type C4-dicarboxylate transport system, periplasmic component | - | 23.8 | 0.80 | 1.97E-05 | 44.8 | 49.9 | 57.3 | 6.5 | 5.0 | NA |
| **stage3_free** | **stage3_shell** | **-5.42** | **2.10E-03** | **V12B01_16616** | **EAP96760.1** | **hypothetical protein** | | | **S** | **FRG domain** | **-** | **12.9** | **2.63** | **2.87E-04** | **0.0** | **0.0** | **NA** | **7.8** | **7.4** | **0.0** |
| stage2 | stage3_shell | 2.47 | 4.08E-02 | V12B01_16626 | EAP96762.1 | putative MoxR-like ATPase | | K09882 | S | AAA domain (dynein-related subfamily) | cobS | 3.8 | 2.17 | 1.31E-02 | 3.2 | 4.7 | 8.6 | 0.0 | 0.0 | 0.0 |
| stage3_free | stage3_shell | 2.35 | 5.22E-02 | V12B01_16626 | EAP96762.1 | putative MoxR-like ATPase | | K09882 | S | AAA domain (dynein-related subfamily) | cobS | 3.8 | 2.26 | 1.58E-02 | 4.6 | 6.8 | NA | 0.0 | 0.0 | 0.0 |
| stage2 | stage3_shell | 2.43 | 3.37E-04 | V12B01_16656 | EAP96768.1 | malate synthase G | Other Energy | K01638 | C | Involved in the glycolate utilization. Catalyzes the condensation and subsequent hydrolysis of acetyl-coenzyme A (acetyl-CoA) and glyoxylate to form malate and CoA | glcB | 740.4 | 0.69 | 4.14E-05 | 463.0 | 58.4 | 207.2 | 11.7 | 34.6 | 33.9 |
| stage2 | stage3_free | 2.07 | 9.09E-04 | V12B01_16661 | EAP96769.1 | putative transcriptional regulator | Regulation | | K | Bacterial regulatory helix-turn-helix protein, lysR family | - | 41.3 | 0.61 | 3.46E-05 | 55.8 | 25.5 | 64.5 | 10.4 | 8.9 | NA |

| | | | | | | | | | | | | | | | | | | | |
|---|---|---|---|---|---|---|---|---|---|---|---|---|---|---|---|---|---|---|---|
| stage2 | stage3_shell | 2.76 | 2.08E-04 | V12B01_16661 | EAP96769.1 | putative transcriptional regulator | Regulation | | K | Bacterial regulatory helix-turn-helix protein, lysR family | - | 41.3 | 0.72 | 2.40E-05 | 55.8 | 25.5 | 64.5 | 6.6 | 3.1 | 2.9 |
| stage2 | stage3_shell | 4.59 | 1.61E-04 | V12B01_16666 | EAP96770.1 | isocitrate lyase | Other Energy | K01637 | C | Isocitrate lyase | aceA | 1180.9 | 1.32 | 1.79E-05 | 1798.4 | 84.8 | 317.7 | 0.0 | 12.3 | 28.5 |
| stage2 | stage3_shell | 3.41 | 5.10E-08 | V12B01_16711 | EAP96779.1 | hypothetical protein | | | G | Ribose-5-phosphate isomerase | - | 99.9 | 0.61 | 1.86E-09 | 545.6 | 565.4 | 379.9 | 12.5 | 42.6 | 35.6 |
| stage3_free | stage3_shell | 2.72 | 8.07E-05 | V12B01_16711 | EAP96779.1 | hypothetical protein | | | G | Ribose-5-phosphate isomerase | - | 99.9 | 0.67 | 5.72E-06 | 388.6 | 266.9 | NA | 12.5 | 42.6 | 35.6 |
| stage2 | stage3_shell | 2.32 | 1.02E-07 | V12B01_16716 | EAP96780.1 | 2-deoxy-D-gluconate 3-dehydrogenase | | K00065 | IQ | Enoyl-(Acyl carrier protein) reductase | kduD | 129.3 | 0.42 | 4.12E-09 | 508.3 | 633.5 | 456.6 | 67.1 | 79.4 | 71.0 |
| **stage2** | **stage3_shell** | **-3.04** | **2.17E-06** | **V12B01_16806** | **EAP96798.1** | **ABC transporter, ATP-binding protein** | | **K02003** | **V** | **ABC-type antimicrobial peptide transport system, ATPase component** | **-** | **88.5** | **0.64** | **1.29E-07** | **17.4** | **24.8** | **38.9** | **49.2** | **142.0** | **339.0** |
| **stage3_free** | **stage3_shell** | **-3.01** | **3.47E-05** | **V12B01_16806** | **EAP96798.1** | **ABC transporter, ATP-binding protein** | **Transport Proteins** | **K02003** | **V** | **ABC-type antimicrobial peptide transport system, ATPase component** | **-** | **88.5** | **0.73** | **2.18E-06** | **20.5** | **33.0** | **NA** | **49.2** | **142.0** | **339.0** |
| stage3_free | stage3_shell | 2.04 | 1.75E-04 | V12B01_16841 | EAP96805.1 | oligopeptide ABC transporter, periplasmic oligopeptide-binding protein | Transport Proteins | K15580 | E | Bacterial extracellular solute-binding proteins, family 5 Middle | - | 23.5 | 0.51 | 1.44E-05 | 46.1 | 38.4 | NA | 9.8 | 8.3 | 0.0 |
| stage2 | stage3_shell | 3.20 | 1.60E-03 | V12B01_16891 | EAP96815.1 | MoxR-related protein | | K03924 | S | COG0714 MoxR-like ATPases | - | 15.8 | 1.02 | 2.61E-04 | 86.9 | 22.2 | 17.0 | 0.0 | 6.3 | 0.0 |
| stage2 | stage3_shell | 2.28 | 1.82E-02 | V12B01_16896 | EAP96816.1 | hypothetical protein | | | S | conserved protein (some members contain a von Willebrand factor type A (vWA) domain) | - | 20.1 | 1.33 | 4.92E-03 | 93.1 | 23.3 | 21.2 | 0.0 | 0.0 | 12.3 |
| stage2 | stage3_free | 3.05 | 4.72E-06 | V12B01_16911 | EAP96819.1 | hypothetical protein | | K07114 | S | protein containing a von Willebrand factor type A (vWA) domain | - | 53.6 | 0.64 | 6.86E-08 | 81.4 | 39.0 | 55.1 | 5.1 | 7.1 | NA |
| stage2 | stage3_shell | 3.72 | 4.72E-07 | V12B01_16911 | EAP96819.1 | hypothetical protein | | K07114 | S | protein containing a von Willebrand factor type A (vWA) domain | - | 53.6 | 0.72 | 2.26E-08 | 81.4 | 39.0 | 55.1 | 2.4 | 3.1 | 2.9 |
| stage2 | stage3_free | 3.03 | 1.77E-04 | V12B01_16916 | EAP96820.1 | hypothetical protein | | | S | Oxygen tolerance | - | 42.1 | 0.80 | 4.81E-06 | 77.7 | 41.9 | 44.5 | 2.7 | 8.0 | NA |
| stage2 | stage3_shell | 2.95 | 1.43E-04 | V12B01_16916 | EAP96820.1 | hypothetical protein | | | S | Oxygen tolerance | - | 42.1 | 0.82 | 1.56E-05 | 77.7 | 41.9 | 44.5 | 1.9 | 0.0 | 11.8 |
| **stage3_free** | **stage3_shell** | **-3.17** | **6.04E-08** | **V12B01_16996** | **EAP96836.1** | **Murein lipoprotein** | **Membrane** | **K06078** | **M** | **Outer membrane lipoprotein** | **lpp** | **5912.0** | **0.57** | **1.20E-09** | **4200.1** | **6412.0** | **NA** | **35150.6** | **65541.0** | **19855.9** |
| **stage2** | **stage3_shell** | **-2.12** | **5.05E-07** | **V12B01_17016** | **EAP96840.1** | **Transposase** | | | **L** | **COG3677 Transposase and inactivated derivatives** | **-** | **390.0** | **0.40** | **2.47E-08** | **59.6** | **138.1** | **112.9** | **178.3** | **468.3** | **397.1** |
| **stage3_free** | **stage3_shell** | **-1.99** | **3.03E-05** | **V12B01_17016** | **EAP96840.1** | **Transposase** | | | **L** | **COG3677 Transposase and inactivated derivatives** | **-** | **390.0** | **0.46** | **1.79E-06** | **109.5** | **113.4** | **NA** | **178.3** | **468.3** | **397.1** |
| stage3_free | stage3_shell | 2.49 | 1.05E-06 | V12B01_17026 | EAP96842.1 | GGDEF family protein | Regulation | | T | COG2199 FOG GGDEF domain | - | 52.6 | 0.48 | 3.40E-08 | 68.3 | 48.0 | NA | 4.0 | 9.6 | 7.2 |
| stage2 | stage3_shell | 1.99 | 5.70E-05 | V12B01_17041 | EAP96845.1 | Multidrug resistance efflux pump | Transport Proteins | | V | multidrug resistance efflux pump | - | 101.7 | 0.48 | 5.48E-06 | 197.8 | 128.4 | 97.9 | 24.0 | 21.4 | 25.2 |
| stage2 | stage3_shell | 2.42 | 1.38E-02 | V12B01_17091 | EAP96855.1 | hypothetical protein | | | - | - | VPA1625 | 10.7 | 1.03 | 3.51E-03 | 32.3 | 34.5 | 34.0 | 0.0 | 0.0 | 12.9 |
| stage3_free | stage3_shell | 3.14 | 3.55E-03 | V12B01_17091 | EAP96855.1 | hypothetical protein | | | - | - | VPA1625 | 10.7 | 1.07 | 5.39E-04 | 56.0 | 51.2 | NA | 0.0 | 0.0 | 12.9 |
| stage2 | stage3_shell | 3.58 | 1.09E-02 | V12B01_17101 | EAP96857.1 | Uncharacterized protein conserved in bacteria | | K09796 | S | protein conserved in bacteria | - | 5.9 | 2.25 | 2.60E-03 | 26.2 | 19.3 | 13.6 | 0.0 | 0.0 | 0.0 |
| stage3_free | stage3_shell | 3.85 | 1.08E-02 | V12B01_17101 | EAP96857.1 | Uncharacterized protein conserved in bacteria | | K09796 | S | protein conserved in bacteria | - | 5.9 | 2.33 | 2.14E-03 | 18.9 | 28.3 | NA | 0.0 | 0.0 | 0.0 |
| **stage2** | **stage3_free** | **-3.20** | **1.40E-04** | **V12B01_17111** | **EAP96859.1** | **sodium/dicarboxylate symporter** | **Transport Proteins** | **K07862** | **E** | **Involved in the import of serine and threonine into the cell, with the concomitant import of sodium (symport system)** | **sstT** | **20.2** | **0.79** | **3.57E-06** | **8.8** | **2.3** | **2.6** | **47.8** | **59.3** | **NA** |
| **stage2** | **stage3_shell** | **-2.14** | **4.37E-05** | **V12B01_17146** | **EAP96866.1** | **putative cytochrome c** | | | **C** | **Cytochrome C oxidase, cbb3-type, subunit III** | **-** | **56.7** | **0.51** | **4.03E-06** | **18.4** | **39.6** | **13.7** | **101.1** | **117.8** | **41.6** |
| **stage2** | **stage3_shell** | **-2.12** | **4.01E-03** | **V12B01_17156** | **EAP96868.1** | **hypothetical protein** | | **K19713** | **C** | **Cytochrome C oxidase, cbb3-type, subunit III** | **-** | **19.4** | **0.76** | **7.79E-04** | **1.7** | **10.4** | **6.7** | **32.5** | **46.2** | **0.0** |
| **stage2** | **stage3_shell** | **-2.85** | **1.54E-04** | **V12B01_17161** | **EAP96869.1** | **hypothetical protein** | | | **C** | **Cytochrome C oxidase, cbb3-type, subunit III** | **-** | **18.1** | **0.76** | **1.69E-05** | **5.0** | **34.5** | **28.0** | **113.7** | **255.0** | **85.3** |
| **stage3_free** | **stage3_shell** | **-3.39** | **1.32E-04** | **V12B01_17161** | **EAP96869.1** | **hypothetical protein** | | | **C** | **Cytochrome C oxidase, cbb3-type, subunit III** | **-** | **18.1** | **0.91** | **1.04E-05** | **15.4** | **14.4** | **NA** | **113.7** | **255.0** | **85.3** |
| stage2 | stage3_shell | 2.18 | 4.68E-02 | V12B01_17236 | EAP95408.1 | GGDEF domain | Regulation | | T | GGDEF domain | - | 5.9 | 1.90 | 1.55E-02 | 3.2 | 2.1 | 2.6 | 0.0 | 0.0 | 0.0 |
| stage3_free | stage3_shell | 3.96 | 6.15E-03 | V12B01_17236 | EAP95408.1 | GGDEF domain | Regulation | | T | GGDEF domain | - | 5.9 | 2.18 | 1.05E-03 | 8.1 | 4.0 | NA | 0.0 | 0.0 | 0.0 |
| stage3_free | stage3_shell | 2.82 | 2.78E-02 | V12B01_17401 | EAP95441.1 | ACP:hemolysin acyltransferase | | K07389 | O | RTX toxin acyltransferase family | rtxC | 6.0 | 2.13 | 7.15E-03 | 12.9 | 14.6 | NA | 0.0 | 0.0 | 0.0 |
| **stage2** | **stage3_free** | **-2.70** | **8.27E-03** | **V12B01_17426** | **EAP95446.1** | **glycerol uptake facilitator protein GlpF** | **Transport Proteins** | **K02440** | **U** | **Belongs to the MIP aquaporin (TC 1.A.8) family** | **glpF** | **14.1** | **1.04** | **5.14E-04** | **14.0** | **7.5** | **2.2** | **93.1** | **68.0** | **NA** |
| **stage2** | **stage3_shell** | **-2.33** | **1.95E-06** | **V12B01_17431** | **EAP95447.1** | **glycerol kinase** | **Fatty Acid and Lipid Degradation** | **K00864** | **F** | **Key enzyme in the regulation of glycerol uptake and metabolism. Catalyzes the phosphorylation of glycerol to yield sn- glycerol 3-phosphate** | **glpK** | **219.4** | **0.48** | **1.12E-07** | **47.0** | **27.2** | **28.9** | **88.1** | **116.1** | **198.0** |
| **stage2** | **stage3_free** | **-2.08** | **2.28E-05** | **V12B01_17441** | **EAP95449.1** | **aerobic glycerol-3-phosphate dehydrogenase** | **Aerobic Respiration** | **K00111** | **C** | **Belongs to the FAD-dependent glycerol-3-phosphate dehydrogenase family** | **glpD** | **41.4** | **0.45** | **4.17E-07** | **10.3** | **12.6** | **16.6** | **57.6** | **68.7** | **NA** |
| **stage3_free** | **stage3_shell** | **-2.18** | **1.97E-02** | **V12B01_17461** | **EAP95453.1** | **hypothetical protein** | | | **S** | **Protein of unknown function (DUF1254)** | **-** | **6.4** | **1.20** | **4.60E-03** | **2.7** | **1.0** | **NA** | **6.4** | **3.0** | **22.7** |
| **stage2** | **stage3_shell** | **-3.39** | **1.94E-02** | **V12B01_17486** | **EAP95458.1** | **hypothetical protein** | | **K10536** | **E** | **agmatine deiminase** | **aguA** | **3.6** | **2.48** | **5.31E-03** | **0.0** | **0.0** | **0.6** | **0.0** | **0.0** | **15.6** |
| **stage2** | **stage3_shell** | **-2.12** | **8.61E-05** | **V12B01_17501** | **EAP95461.1** | **hypothetical transcriptional regulator** | | | **K** | **Bacterial regulatory proteins, tetR family** | **-** | **39.7** | **0.52** | **8.82E-06** | **12.1** | **13.6** | **29.4** | **119.8** | **62.7** | **18.2** |
| **stage2** | **stage3_shell** | **-2.30** | **1.96E-02** | **V12B01_17526** | **EAP95466.1** | **pyrimidine reductase** | | | **H** | **COG0262 Dihydrofolate reductase** | **-** | **4.1** | **1.25** | **5.41E-03** | **0.0** | **1.3** | **1.8** | **6.0** | **5.7** | **10.7** |
| **stage2** | **stage3_shell** | **-2.11** | **7.48E-04** | **V12B01_17536** | **EAP95468.1** | **Putative transcriptional regulator, LysR family** | **Regulation** | | **K** | **Bacterial regulatory helix-turn-helix protein, lysR family** | **-** | **28.5** | **0.63** | **1.05E-04** | **17.6** | **9.6** | **1.4** | **23.5** | **51.5** | **32.3** |
| **stage3_free** | **stage3_shell** | **-1.99** | **6.76E-07** | **V12B01_17596** | **EAP95480.1** | **hypothetical transcriptional regulator** | **Regulation** | | **K** | **Transcriptional regulator** | **-** | **104.2** | **0.38** | **2.01E-08** | **40.5** | **39.4** | **NA** | **109.7** | **197.7** | **66.2** |
| **stage2** | **stage3_shell** | **-3.73** | **1.07E-16** | **V12B01_17661** | **EAP95493.1** | **Transposase** | | **K07486** | **L** | **COG3547 Transposase and inactivated derivatives** | **-** | **54.1** | **0.43** | **6.53E-19** | **2.6** | **9.3** | **9.7** | **57.6** | **82.6** | **86.0** |
| stage2 | stage3_shell | 2.02 | 2.92E-02 | V12B01_17701 | EAP95501.1 | hypothetical protein | | | - | #N/A | #N/A | 9.8 | 0.97 | 8.71E-03 | 135.5 | 121.4 | 172.4 | 30.4 | 14.4 | 0.0 |
| **stage2** | **stage3_shell** | **-2.32** | **6.82E-06** | **V12B01_17806** | **EAP95522.1** | **PTS system, IIabc component** | **Transport Proteins** | **K02757** | **G** | **phosphotransferase system, EIIB** | **-** | **47.6** | **0.50** | **4.77E-07** | **5.6** | **11.4** | **13.3** | **55.7** | **60.1** | **9.9** |
| **stage2** | **stage3_shell** | **-2.39** | **7.75E-13** | **V12B01_17856** | **EAP95532.1** | **hypothetical protein** | | | **C** | **Disulfide bond formation protein** | **-** | **180.4** | **0.32** | **9.16E-15** | **65.3** | **78.9** | **75.8** | **188.1** | **285.0** | **397.2** |

| | | | | | | | | | | | | | | | | | | | | |
|---|---|---|---|---|---|---|---|---|---|---|---|---|---|---|---|---|---|---|---|---|
| **stage2** | **stage3_shell** | **-3.41** | **1.41E-07** | **V12B01_17861** | **EAP95533.1** | **hypothetical protein** | | | **-** | **-** | **-** | **56.4** | **0.64** | **5.86E-09** | **94.1** | **61.3** | **42.0** | **338.0** | **523.7** | **839.6** |
| **stage3_free** | **stage3_shell** | **-3.91** | **2.89E-07** | **V12B01_17861** | **EAP95533.1** | **hypothetical protein** | | | **-** | **-** | **-** | **56.4** | **0.75** | **7.25E-09** | **20.5** | **70.8** | **NA** | **338.0** | **523.7** | **839.6** |
| **stage2** | **stage3_shell** | **-2.23** | **9.90E-08** | **V12B01_17866** | **EAP95534.1** | **hypothetical protein** | | | **-** | **-** | **-** | **78.7** | **0.40** | **3.95E-09** | **4.4** | **11.7** | **11.5** | **38.4** | **44.5** | **20.3** |
| **stage3_free** | **stage3_shell** | **-2.15** | **7.09E-06** | **V12B01_17866** | **EAP95534.1** | **hypothetical protein** | | | **-** | **-** | **-** | **78.7** | **0.46** | **3.13E-07** | **11.0** | **8.3** | **NA** | **38.4** | **44.5** | **20.3** |
| stage2 | stage3_shell | 2.67 | 1.24E-02 | V12B01_17946 | EAP95725.1 | transcriptional regulator, MarR family | Regulation | | K | COG1846 Transcriptional regulators | - | 11.0 | 1.11 | 3.04E-03 | 37.7 | 28.6 | 38.1 | 0.0 | 6.3 | 0.0 |
| stage3_free | stage3_shell | 2.36 | 2.77E-02 | V12B01_17946 | EAP95725.1 | transcriptional regulator, MarR family | Regulation | | K | COG1846 Transcriptional regulators | - | 11.0 | 1.16 | 7.11E-03 | 29.4 | 31.8 | NA | 0.0 | 6.3 | 0.0 |
| stage2 | stage3_shell | 2.96 | 2.77E-02 | V12B01_17986 | EAP95733.1 | C4-dicarboxylate transporter family protein, DctM subunit | Transport Proteins | K11690 | G | Tripartite ATP-independent periplasmic transporter, DctM component | - | 4.0 | 2.45 | 8.14E-03 | 5.3 | 5.5 | 6.7 | 0.0 | 0.0 | 0.0 |
| stage2 | stage3_free | 4.52 | 1.19E-03 | V12B01_17996 | EAP95735.1 | putative C4-dicarboxylate-binding periplasmic protein DctP | Transport Proteins | K21395 | G | Bacterial extracellular solute-binding protein, family 7 | - | 18.7 | 1.49 | 5.15E-05 | 51.5 | 50.9 | 52.6 | 2.6 | 0.0 | NA |
| stage2 | stage3_free | 2.43 | 3.14E-02 | V12B01_18001 | EAP95736.1 | amidohydrolase family protein | | | S | COG1473 Metal-dependent amidase aminoacylase carboxypeptidase | - | 4.8 | 1.39 | 2.92E-03 | 9.1 | 8.1 | 7.4 | 0.0 | 1.6 | NA |
| stage2 | stage3_free | 2.95 | 9.86E-03 | V12B01_18006 | EAP95737.1 | Asp/Glu/hydantoin racemase | | K01799 | Q | Catalyzes the conversion of maleate to fumarate | - | 7.4 | 1.17 | 6.37E-04 | 19.0 | 12.0 | 13.3 | 2.3 | 0.0 | NA |
| stage2 | stage3_free | 4.59 | 1.63E-07 | V12B01_18011 | EAP95738.1 | hypothetical transcriptional regulator | | K03719 | K | helix_turn_helix ASNC type | - | 23.0 | 0.82 | 1.86E-09 | 107.6 | 68.4 | 114.8 | 4.6 | 2.0 | NA |
| **stage3_free** | **stage3_shell** | **-2.24** | **1.02E-02** | **V12B01_18011** | **EAP95738.1** | **hypothetical transcriptional regulator** | **Regulation** | **K03719** | **K** | **helix_turn_helix ASNC type** | **-** | **23.0** | **0.95** | **1.98E-03** | **4.6** | **2.0** | **NA** | **19.5** | **18.4** | **11.6** |
| stage2 | stage3_free | 3.10 | 3.65E-11 | V12B01_18041 | EAP95744.1 | Zinc metalloprotease | | K08604 | E | COG3227 Zinc metalloprotease (elastase) | - | 169.1 | 0.45 | 1.69E-13 | 340.3 | 294.0 | 258.7 | 34.4 | 30.8 | NA |
| stage2 | stage3_shell | 4.03 | 1.66E-18 | V12B01_18041 | EAP95744.1 | Zinc metalloprotease | | K08604 | E | COG3227 Zinc metalloprotease (elastase) | - | 169.1 | 0.45 | 5.95E-21 | 340.3 | 294.0 | 258.7 | 4.1 | 14.1 | 20.6 |
| stage2 | stage3_free | 2.15 | 1.57E-02 | V12B01_18046 | EAP95745.1 | aminopeptidase | | K05994 | G | Bacterial pre-peptidase C-terminal domain | - | 10.6 | 1.01 | 1.17E-03 | 46.8 | 19.6 | 12.2 | 2.9 | 4.9 | NA |
| stage2 | stage3_shell | 4.07 | 9.27E-04 | V12B01_18046 | EAP95745.1 | aminopeptidase | | K05994 | G | Bacterial pre-peptidase C-terminal domain | - | 10.6 | 1.22 | 1.35E-04 | 46.8 | 19.6 | 12.2 | 0.0 | 1.9 | 0.0 |
| stage2 | stage3_shell | 2.40 | 3.78E-02 | V12B01_18056 | EAP95747.1 | hypothetical protein | | | - | - | - | 6.7 | 2.01 | 1.19E-02 | 1.7 | 1.6 | 2.5 | 0.0 | 0.0 | 0.0 |
| stage3_free | stage3_shell | 3.00 | 2.34E-02 | V12B01_18056 | EAP95747.1 | hypothetical protein | | | - | - | - | 6.7 | 2.17 | 5.78E-03 | 2.8 | 2.6 | NA | 0.0 | 0.0 | 0.0 |
| stage2 | stage3_shell | 1.99 | 1.96E-03 | V12B01_18076 | EAP95751.1 | hypothetical protein | | | S | Tripartite ATP-independent periplasmic transporter, DctM component | - | 49.0 | 0.65 | 3.28E-04 | 110.9 | 50.6 | 18.9 | 16.1 | 3.5 | 7.8 |
| stage3_free | stage3_shell | 2.38 | 5.00E-02 | V12B01_18101 | EAP95756.1 | putative 2-keto-3-deoxy-6-phosphogluconatealdolase | | K01625 | G | COG0800 2-keto-3-deoxy-6-phosphogluconate aldolase | - | 3.3 | 2.30 | 1.49E-02 | 10.1 | 9.1 | NA | 0.0 | 0.0 | 0.0 |
| **stage2** | **stage3_shell** | **-2.09** | **8.53E-06** | **V12B01_18151** | **EAP95766.1** | **predicted membrane-associated, metal-dependent hydrolase** | | **K03760** | **S** | **membrane-associated, metal-dependent hydrolase** | **eptA** | **53.3** | **0.45** | **6.33E-07** | **4.7** | **10.9** | **9.6** | **49.2** | **18.6** | **19.3** |
| **stage3_free** | **stage3_shell** | **-2.30** | **2.46E-05** | **V12B01_18151** | **EAP95766.1** | **predicted membrane-associated, metal-dependent hydrolase** | **Membrane** | **K03760** | **S** | **membrane-associated, metal-dependent hydrolase** | **eptA** | **53.3** | **0.53** | **1.38E-06** | **6.8** | **7.7** | **NA** | **49.2** | **18.6** | **19.3** |
| **stage2** | **stage3_shell** | **-2.06** | **1.13E-03** | **V12B01_18166** | **EAP95769.1** | **acetyltransferase, GNAT family** | | | **J** | **Acetyltransferase (GNAT) domain** | **-** | **12.0** | **0.65** | **1.71E-04** | **3.2** | **3.3** | **5.9** | **7.6** | **14.3** | **26.9** |
| **stage2** | **stage3_shell** | **-2.22** | **1.40E-05** | **V12B01_18211** | **EAP95778.1** | **hypothetical protein** | | | **S** | **Protein conserved in bacteria** | **-** | **21.1** | **0.50** | **1.14E-06** | **12.0** | **9.5** | **10.6** | **24.3** | **58.4** | **40.0** |
| **stage3_free** | **stage3_shell** | **-3.23** | **4.98E-07** | **V12B01_18211** | **EAP95778.1** | **hypothetical protein** | | | **S** | **Protein conserved in bacteria** | **-** | **21.1** | **0.62** | **1.40E-08** | **3.2** | **7.3** | **NA** | **24.3** | **58.4** | **40.0** |
| **stage2** | **stage3_shell** | **-2.12** | **1.36E-04** | **V12B01_18226** | **EAP95781.1** | **hypothetical protein** | | | **C** | **COG4230 Delta 1-pyrroline-5-carboxylate dehydrogenase** | **-** | **85.5** | **0.55** | **1.47E-05** | **39.7** | **23.6** | **16.4** | **34.0** | **90.1** | **157.5** |
| stage3_free | stage3_shell | 2.10 | 6.81E-04 | V12B01_18246 | EAP95785.1 | hypothetical protein | | | S | Sulfatase-modifying factor enzyme 1 | - | 90.2 | 0.60 | 7.27E-05 | 100.7 | 104.7 | NA | 7.0 | 15.7 | 21.8 |
| **stage2** | **stage3_shell** | **-3.46** | **8.16E-17** | **V12B01_18316** | **EAP95799.1** | **hypothetical protein** | | | **-** | **-** | **-** | **94.6** | **0.40** | **4.52E-19** | **23.6** | **24.6** | **25.1** | **102.4** | **249.4** | **265.9** |
| **stage3_free** | **stage3_shell** | **-4.15** | **3.35E-16** | **V12B01_18316** | **EAP95799.1** | **hypothetical protein** | | | **-** | **-** | **-** | **94.6** | **0.49** | **6.36E-19** | **10.1** | **20.2** | **NA** | **102.4** | **249.4** | **265.9** |
| **stage3_free** | **stage3_shell** | **-2.26** | **7.13E-07** | **V12B01_18346** | **EAP95805.1** | **hypothetical two-component sensor** | **Regulation** | | **T** | **Evidence 3 Function proposed based on presence of conserved amino acid motif, structural feature or limited homology** | **-** | **506.2** | **0.43** | **2.15E-08** | **41.0** | **60.7** | **NA** | **172.3** | **340.6** | **84.4** |
| **stage2** | **stage3_shell** | **-5.26** | **2.00E-17** | **V12B01_18386** | **EAP95813.1** | **4-hydroxyphenylpyruvate dioxygenase** | | **K00457** | **C** | **COG3185 4-hydroxyphenylpyruvate dioxygenase and related hemolysins** | **hppD** | **641.8** | **0.60** | **9.73E-20** | **5.8** | **5.6** | **8.4** | **305.0** | **321.2** | **18.6** |
| **stage3_free** | **stage3_shell** | **-4.50** | **1.19E-10** | **V12B01_18386** | **EAP95813.1** | **4-hydroxyphenylpyruvate dioxygenase** | | **K00457** | **C** | **COG3185 4-hydroxyphenylpyruvate dioxygenase and related hemolysins** | **hppD** | **641.8** | **0.67** | **1.21E-12** | **10.2** | **11.6** | **NA** | **305.0** | **321.2** | **18.6** |
| **stage2** | **stage3_shell** | **-5.08** | **9.79E-16** | **V12B01_18391** | **EAP95814.1** | **putative oxidoreductase** | | **K00451** | **Q** | **COG3508 Homogentisate 1,2-dioxygenase** | **-** | **318.7** | **0.61** | **6.40E-18** | **4.6** | **4.0** | **4.8** | **156.9** | **200.7** | **24.5** |
| **stage3_free** | **stage3_shell** | **-4.45** | **5.03E-10** | **V12B01_18391** | **EAP95814.1** | **putative oxidoreductase** | | **K00451** | **Q** | **COG3508 Homogentisate 1,2-dioxygenase** | **-** | **318.7** | **0.69** | **6.05E-12** | **4.3** | **9.1** | **NA** | **156.9** | **200.7** | **24.5** |
| **stage2** | **stage3_shell** | **-5.73** | **4.23E-12** | **V12B01_18396** | **EAP95815.1** | **hypothetical protein** | | **K16171** | **Q** | **COG0179 2-keto-4-pentenoate hydratase 2-oxohepta-3-ene-1,7-dioic acid hydratase (catechol pathway)** | **-** | **288.0** | **0.80** | **5.56E-14** | **5.1** | **0.7** | **3.0** | **197.0** | **215.7** | **5.5** |
| **stage3_free** | **stage3_shell** | **-4.87** | **1.42E-07** | **V12B01_18396** | **EAP95815.1** | **hypothetical protein** | | **K16171** | **Q** | **COG0179 2-keto-4-pentenoate hydratase 2-oxohepta-3-ene-1,7-dioic acid hydratase (catechol pathway)** | **-** | **288.0** | **0.91** | **3.23E-09** | **5.3** | **4.6** | **NA** | **197.0** | **215.7** | **5.5** |
| **stage2** | **stage3_shell** | **-3.72** | **7.41E-12** | **V12B01_18401** | **EAP95816.1** | **putative glutathione S-transferase** | | | **O** | **maleylacetoacetate isomerase** | **maiA** | **95.9** | **0.52** | **1.06E-13** | **3.2** | **8.0** | **4.9** | **63.5** | **83.7** | **24.2** |
| **stage3_free** | **stage3_shell** | **-2.82** | **2.46E-06** | **V12B01_18401** | **EAP95816.1** | **putative glutathione S-transferase** | | | **O** | **maleylacetoacetate isomerase** | **maiA** | **95.9** | **0.57** | **8.73E-08** | **14.9** | **4.8** | **NA** | **63.5** | **83.7** | **24.2** |
| **stage3_free** | **stage3_shell** | **-3.13** | **8.58E-06** | **V12B01_18411** | **EAP95818.1** | **acetyl-CoA synthase** | **Carbohydrates and Carboxylates Degradation** | **K01907** | **I** | **COG0365 Acyl-coenzyme A synthetases AMP-(fatty) acid ligases** | **acs-2** | **110.3** | **0.69** | **3.95E-07** | **2.1** | **6.5** | **NA** | **31.4** | **62.9** | **4.3** |
| **stage3_free** | **stage3_shell** | **-2.11** | **1.47E-02** | **V12B01_18416** | **EAP95819.1** | **acetyl-CoA synthase** | **Carbohydrates and Carboxylates Degradation** | **K01907** | **I** | **COG0365 Acyl-coenzyme A synthetases AMP-(fatty) acid ligases** | **acs-2** | **83.9** | **1.02** | **3.22E-03** | **4.2** | **10.7** | **NA** | **37.7** | **71.5** | **0.0** |
| **stage2** | **stage3_shell** | **-3.10** | **5.04E-10** | **V12B01_18421** | **EAP95820.1** | **phenylalanine-4-hydroxylase** | **Amino Acid Degradation** | **K00500** | **E** | **Phenylalanine-4-hydroxylase** | **phhA** | **624.9** | **0.48** | **9.99E-12** | **6.2** | **15.6** | **17.3** | **100.5** | **121.8** | **50.1** |

| | | | | | | | | | | | | | | | | | | | |
|---|---|---|---|---|---|---|---|---|---|---|---|---|---|---|---|---|---|---|---|
| stage3_free | stage3_shell | -3.17 | 4.76E-08 | V12B01_18421 | EAP95820.1 | phenylalanine-4-hydroxylase | Amino Acid Degradation | K00500 | E | Phenylalanine-4-hydroxylase | phhA | 624.9 | 0.56 | 8.95E-10 | 13.8 | 10.9 | NA | 100.5 | 121.8 | 50.1 |
| stage2 | stage3_shell | -3.26 | 4.79E-08 | V12B01_18426 | EAP95821.1 | pterin-4-alpha-carbinolamine dehydratase | Amino Acid Degradation | K01724 | H | pterin-4-alpha-carbinolamine dehydratase | phhB | 161.7 | 0.58 | 1.70E-09 | 5.2 | 15.7 | 2.8 | 56.6 | 93.7 | 33.6 |
| stage3_free | stage3_shell | -2.39 | 1.96E-04 | V12B01_18426 | EAP95821.1 | pterin-4-alpha-carbinolamine dehydratase | Amino Acid Degradation | K01724 | H | pterin-4-alpha-carbinolamine dehydratase | phhB | 161.7 | 0.63 | 1.64E-05 | 16.2 | 11.3 | NA | 56.6 | 93.7 | 33.6 |
| stage2 | stage3_shell | -2.74 | 1.96E-14 | V12B01_18466 | EAP95829.1 | Fructose-bisphosphate aldolase I | Glycolysis | K01624 | H | fructose-bisphosphate aldolase, class II, Calvin cycle subtype | cbbA | 97.0 | 0.34 | 1.69E-16 | 16.9 | 28.6 | 25.5 | 78.9 | 171.0 | 116.3 |
| stage3_free | stage3_shell | -3.14 | 7.39E-14 | V12B01_18466 | EAP95829.1 | Fructose-bisphosphate aldolase I | Glycolysis | K01624 | H | fructose-bisphosphate aldolase, class II, Calvin cycle subtype | cbbA | 97.0 | 0.40 | 3.31E-16 | 15.6 | 20.5 | NA | 78.9 | 171.0 | 116.3 |
| stage2 | stage3_shell | -2.11 | 1.21E-05 | V12B01_18471 | EAP95830.1 | putative sugar kinase | | | H | Catalyzes the phosphorylation of ribose at O-5 in a reaction requiring ATP and magnesium. The resulting D-ribose-5-phosphate can then be used either for sythesis of nucleotides, histidine, and tryptophan, or as a component of the pentose phosphate pathway | rbsK | 62.9 | 0.47 | 9.56E-07 | 9.9 | 28.6 | 35.8 | 80.4 | 106.2 | 73.3 |
| stage3_free | stage3_shell | -2.82 | 6.32E-07 | V12B01_18471 | EAP95830.1 | putative sugar kinase | | | H | Catalyzes the phosphorylation of ribose at O-5 in a reaction requiring ATP and magnesium. The resulting D-ribose-5-phosphate can then be used either for sythesis of nucleotides, histidine, and tryptophan, or as a component of the pentose phosphate pathway | rbsK | 62.9 | 0.55 | 1.83E-08 | 16.0 | 14.0 | NA | 80.4 | 106.2 | 73.3 |
| stage3_free | stage3_shell | -2.10 | 3.16E-04 | V12B01_18476 | EAP95831.1 | hypothetical protein | | | F | COG1072 Panthothenate kinase | frcK | 32.6 | 0.57 | 2.96E-05 | 11.1 | 8.8 | NA | 34.0 | 32.2 | 43.3 |
| stage2 | stage3_free | -3.73 | 7.26E-07 | V12B01_18591 | EAP95854.1 | N-ethylmaleimide reductase | | K10680 | C | COG1902 NADH flavin oxidoreductases, Old Yellow Enzyme family | nemA | 936.6 | 0.70 | 9.12E-09 | 4.9 | 7.8 | 5.2 | 164.2 | 24.7 | NA |
| stage2 | stage3_shell | -2.37 | 4.42E-04 | V12B01_18591 | EAP95854.1 | N-ethylmaleimide reductase | | K10680 | C | COG1902 NADH flavin oxidoreductases, Old Yellow Enzyme family | nemA | 936.6 | 0.67 | 5.66E-05 | 4.9 | 7.8 | 5.2 | 29.3 | 45.8 | 7.8 |
| stage2 | stage3_free | -4.59 | 2.04E-12 | V12B01_18596 | EAP95855.1 | hypothetical MFS transporter | Transport Proteins | K19577 | EGP | Major Facilitator Superfamily | - | 577.5 | 0.62 | 6.74E-15 | 2.9 | 2.9 | 0.8 | 88.2 | 26.8 | NA |
| stage2 | stage3_shell | -3.39 | 1.01E-07 | V12B01_18596 | EAP95855.1 | hypothetical MFS transporter | Transport Proteins | K19577 | EGP | Major Facilitator Superfamily | - | 577.5 | 0.62 | 4.03E-09 | 2.9 | 2.9 | 0.8 | 19.3 | 24.4 | 11.5 |
| stage2 | stage3_free | -3.78 | 1.47E-07 | V12B01_18601 | EAP95856.1 | putative dehydrogenase | | | IQ | COG1028 Dehydrogenases with different specificities (related to short-chain alcohol dehydrogenases) | - | 475.9 | 0.67 | 1.56E-09 | 1.3 | 10.4 | 4.1 | 115.7 | 57.6 | NA |
| stage2 | stage3_shell | -3.25 | 9.68E-07 | V12B01_18601 | EAP95856.1 | putative dehydrogenase | | | IQ | COG1028 Dehydrogenases with different specificities (related to short-chain alcohol dehydrogenases) | - | 475.9 | 0.65 | 5.21E-08 | 1.3 | 10.4 | 4.1 | 58.4 | 55.3 | 16.7 |
| stage2 | stage3_shell | 2.46 | 2.82E-03 | V12B01_18611 | EAP95858.1 | hypothetical protein | | | - | - | - | 40.0 | 0.86 | 5.19E-04 | 184.5 | 28.0 | 67.8 | 0.0 | 14.9 | 14.1 |
| stage3_free | stage3_shell | 2.23 | 1.01E-02 | V12B01_18611 | EAP95858.1 | hypothetical protein | | | - | - | - | 40.0 | 0.92 | 1.96E-03 | 99.4 | 71.2 | NA | 0.0 | 14.9 | 14.1 |
| stage2 | stage3_shell | 3.13 | 1.90E-04 | V12B01_18616 | EAP95859.1 | hypothetical protein | | | Q | Flavin containing amine oxidoreductase | - | 90.5 | 0.84 | 2.16E-05 | 134.0 | 22.0 | 21.2 | 8.2 | 2.9 | 0.0 |
| stage3_free | stage3_shell | 2.98 | 1.06E-03 | V12B01_18616 | EAP95859.1 | hypothetical protein | | | Q | Flavin containing amine oxidoreductase | - | 90.5 | 0.90 | 1.23E-04 | 70.2 | 43.6 | NA | 8.2 | 2.9 | 0.0 |
| stage3_free | stage3_shell | 2.22 | 9.86E-05 | V12B01_18626 | EAP95861.1 | hypothetical protein | | | S | Glutathione-dependent formaldehyde-activating enzyme | - | 39.6 | 0.54 | 7.24E-06 | 180.4 | 182.1 | NA | 32.5 | 15.4 | 29.0 |
| stage3_free | stage3_shell | -2.14 | 6.89E-03 | V12B01_18666 | EAP95869.1 | Uncharacterized conserved secreted protein | | | S | Protein of unknown function (DUF3833) | PP2730 | 138.3 | 0.86 | 1.22E-03 | 11.8 | 13.3 | NA | 41.2 | 103.1 | 21.0 |
| stage2 | stage3_shell | -2.40 | 5.98E-04 | V12B01_18671 | EAP95870.1 | hypothetical protein | | | H | Chalcone isomerase-like | - | 208.8 | 0.70 | 8.07E-05 | 3.8 | 13.3 | 9.0 | 13.6 | 92.6 | 19.4 |
| stage3_free | stage3_shell | -2.98 | 3.60E-04 | V12B01_18671 | EAP95870.1 | hypothetical protein | | | H | Chalcone isomerase-like | - | 208.8 | 0.85 | 3.48E-05 | 3.9 | 7.4 | NA | 13.6 | 92.6 | 19.4 |
| stage3_free | stage3_shell | -2.05 | 9.46E-03 | V12B01_18676 | EAP95871.1 | hypothetical protein | | | S | Protein of unknown function (DUF2878) | - | 45.4 | 0.83 | 1.81E-03 | 7.9 | 3.7 | NA | 18.4 | 49.3 | 0.0 |
| stage2 | stage3_shell | -2.38 | 1.44E-05 | V12B01_18696 | EAP95875.1 | oxidoreductase, short-chain dehydrogenase/reductase family | | | S | Enoyl-(Acyl carrier protein) reductase | - | 49.6 | 0.54 | 1.17E-06 | 24.9 | 16.9 | 11.9 | 43.7 | 88.9 | 97.2 |
| stage3_free | stage3_shell | -2.29 | 2.36E-04 | V12B01_18696 | EAP95875.1 | oxidoreductase, short-chain dehydrogenase/reductase family | | | S | Enoyl-(Acyl carrier protein) reductase | - | 49.6 | 0.62 | 2.07E-05 | 23.1 | 13.8 | NA | 43.7 | 88.9 | 97.2 |
| stage2 | stage3_shell | -2.04 | 6.67E-05 | V12B01_18711 | EAP95878.1 | hypothetical protein | | | S | Protein of unknown function (DUF1722) | - | 745.2 | 0.50 | 6.71E-06 | 61.4 | 82.6 | 52.0 | 210.7 | 405.3 | 72.1 |
| stage2 | stage3_shell | -2.57 | 8.86E-07 | V12B01_18716 | EAP95879.1 | putative transcriptional activator | Regulation | K07167 | T | ChrR Cupin-like domain | - | 971.0 | 0.51 | 4.69E-08 | 39.4 | 60.3 | 40.4 | 155.1 | 419.2 | 120.1 |
| stage3_free | stage3_shell | -2.18 | 2.00E-04 | V12B01_18716 | EAP95879.1 | putative transcriptional activator | Transcription | K07167 | T | ChrR Cupin-like domain | - | 971.0 | 0.58 | 1.68E-05 | 52.7 | 65.5 | NA | 155.1 | 419.2 | 120.1 |
| stage2 | stage3_shell | -2.11 | 8.61E-05 | V12B01_18721 | EAP95880.1 | RNA polymerase sigma-70 factor | Sigma Factors | | K | Belongs to the sigma-70 factor family. ECF subfamily | - | 813.4 | 0.52 | 8.81E-06 | 15.1 | 53.6 | 25.5 | 91.2 | 170.4 | 79.1 |
| stage3_free | stage3_shell | -2.26 | 2.01E-04 | V12B01_18721 | EAP95880.1 | RNA polymerase sigma-70 factor | Sigma Factors | | K | Belongs to the sigma-70 factor family. ECF subfamily | - | 813.4 | 0.61 | 1.70E-05 | 28.9 | 25.9 | NA | 91.2 | 170.4 | 79.1 |
| stage2 | stage3_shell | -2.67 | 7.70E-03 | V12B01_18726 | EAP95881.1 | hypothetical protein | | K07157 | S | ATP-dependent protease La (LON) substrate-binding domain | - | 54.3 | 1.19 | 1.71E-03 | 2.9 | 0.6 | 0.0 | 15.4 | 0.0 | 9.2 |
| stage2 | stage3_shell | -3.37 | 1.96E-07 | V12B01_18801 | EAP92535.1 | hypothetical protein | | | - | - | - | 4824.4 | 0.64 | 8.39E-09 | 998.3 | 5080.0 | 4318.3 | 42638.1 | 49850.8 | 4201.0 |
| stage3_free | stage3_shell | -3.15 | 1.65E-05 | V12B01_18801 | EAP92535.1 | hypothetical protein | | | - | - | - | 4824.4 | 0.73 | 8.47E-07 | 3338.2 | 4445.7 | NA | 42638.1 | 49850.8 | 4201.0 |
| stage2 | stage3_shell | -3.06 | 5.64E-03 | V12B01_18831 | EAP92541.1 | hypothetical protein | | | S | Nucleoside recognition | yjiH | 22.1 | 1.37 | 1.17E-03 | 1.8 | 8.2 | 8.5 | 17.3 | 14.3 | 146.6 |
| stage2 | stage3_shell | -3.50 | 7.63E-03 | V12B01_18836 | EAP92542.1 | hypothetical protein | | | S | Nucleoside recognition | yjiG | 11.3 | 1.82 | 1.69E-03 | 0.0 | 4.5 | 2.7 | 6.8 | 19.4 | 97.3 |
| stage3_free | stage3_shell | -2.90 | 3.18E-09 | V12B01_18846 | EAP92544.1 | hypothetical protein | | | - | - | - | 71.1 | 0.47 | 4.69E-11 | 31.0 | 30.8 | NA | 99.3 | 273.5 | 171.6 |
| stage2 | stage3_shell | -2.21 | 3.44E-04 | V12B01_18851 | EAP92545.1 | hypothetical protein | | | S | START domain | - | 47.2 | 0.62 | 4.26E-05 | 23.8 | 15.0 | 29.9 | 20.1 | 73.7 | 170.1 |
| stage3_free | stage3_shell | -2.87 | 7.24E-05 | V12B01_18851 | EAP92545.1 | hypothetical protein | | | S | START domain | - | 47.2 | 0.73 | 5.02E-06 | 17.3 | 11.3 | NA | 20.1 | 73.7 | 170.1 |

| | | | | | | | | | | | | | | | | | | | | |
|---|---|---|---|---|---|---|---|---|---|---|---|---|---|---|---|---|---|---|---|---|
| stage2 | stage3_shell | 2.75 | 3.10E-09 | V12B01_18901 | EAP92555.1 | hypothetical protein | | K09913 | S | Catalyzes the phosphorolysis of diverse nucleosides, yielding D-ribose 1-phosphate and the respective free bases. Can use uridine, adenosine, guanosine, cytidine, thymidine, inosine and xanthosine as substrates. Also catalyzes the reverse reactions | ppnP | 151.5 | 0.45 | 7.53E-11 | 777.4 | 895.5 | 794.5 | 68.0 | 75.1 | 111.1 |
| **stage2** | stage3_shell | **-2.44** | **5.75E-05** | **V12B01_18906** | **EAP92556.1** | **pirin-related protein** | | **K06911** | **S** | **Belongs to the pirin family** | **-** | **69.3** | **0.60** | **5.59E-06** | **8.7** | **48.3** | **26.1** | **129.9** | **227.0** | **39.1** |
| **stage3_free** | stage3_shell | **-3.03** | **1.49E-05** | **V12B01_18906** | **EAP92556.1** | **pirin-related protein** | | **K06911** | **S** | **Belongs to the pirin family** | **-** | **69.3** | **0.69** | **7.47E-07** | **14.1** | **22.0** | **NA** | **129.9** | **227.0** | **39.1** |
| **stage2** | stage3_shell | **-5.62** | **2.37E-18** | **V12B01_18911** | **EAP92557.1** | **hypothetical protein** | | | **-** | **-** | **-** | **922.9** | **0.62** | **1.05E-20** | **63.1** | **246.2** | **282.2** | **1841.4** | **6003.2** | **13610.1** |
| **stage3_free** | stage3_shell | **-5.85** | **1.64E-15** | **V12B01_18911** | **EAP92557.1** | **hypothetical protein** | | | **-** | **-** | **-** | **922.9** | **0.71** | **4.14E-18** | **157.3** | **175.5** | **NA** | **1841.4** | **6003.2** | **13610.1** |
| stage3_free | stage3_shell | 2.89 | 3.18E-02 | V12B01_18946 | EAP92564.1 | regulatory protein UhpC | | K07783 | G | COG2271 Sugar phosphate permease | uhpC | 4.7 | 2.35 | 8.41E-03 | 6.1 | 5.0 | NA | 0.0 | 0.0 | 0.0 |
| **stage2** | stage3_shell | **-3.48** | **7.36E-13** | **V12B01_18991** | **EAP95136.1** | **ABC transporter ATP-binding protein** | **Transport Proteins** | | **EP** | **ATPases associated with a variety of cellular activities** | **-** | **62.2** | **0.47** | **8.54E-15** | **5.9** | **9.3** | **16.0** | **71.6** | **123.6** | **82.5** |
| **stage3_free** | stage3_shell | **-4.64** | **3.04E-13** | **V12B01_18991** | **EAP95136.1** | **ABC transporter ATP-binding protein** | **Transport Proteins** | | **EP** | **ATPases associated with a variety of cellular activities** | **-** | **62.2** | **0.61** | **1.60E-15** | **5.4** | **3.8** | **NA** | **71.6** | **123.6** | **82.5** |
| **stage2** | stage3_shell | **-2.23** | **4.80E-05** | **V12B01_18996** | **EAP95137.1** | **oligopeptide porter** | **Transport Proteins** | | **E** | **ATPases associated with a variety of cellular activities** | **-** | **28.0** | **0.55** | **4.51E-06** | **5.7** | **8.7** | **4.0** | **9.4** | **28.4** | **36.8** |
| **stage3_free** | stage3_shell | **-3.08** | **1.19E-05** | **V12B01_18996** | **EAP95137.1** | **oligopeptide porter** | **Transport Proteins** | | **E** | **ATPases associated with a variety of cellular activities** | **-** | **28.0** | **0.69** | **5.75E-07** | **3.2** | **3.4** | **NA** | **9.4** | **28.4** | **36.8** |
| **stage2** | stage3_shell | **-3.01** | **2.20E-09** | **V12B01_19056** | **EAP95149.1** | **transcriptional regulator** | **Regulation** | **K02529** | **K** | **transcriptional regulators** | **-** | **75.0** | **0.49** | **5.09E-11** | **11.0** | **25.6** | **31.9** | **88.8** | **127.6** | **211.8** |
| **stage3_free** | stage3_shell | **-2.40** | **2.16E-05** | **V12B01_19056** | **EAP95149.1** | **transcriptional regulator** | **Regulation** | **K02529** | **K** | **transcriptional regulators** | **-** | **75.0** | **0.55** | **1.14E-06** | **28.6** | **39.1** | **NA** | **88.8** | **127.6** | **211.8** |
| stage2 | stage3_free | 2.41 | 1.58E-02 | V12B01_19131 | EAP95164.1 | hypothetical protein | | | - | - | - | 66.2 | 1.09 | 1.19E-03 | 62.9 | 14.3 | 6.3 | 3.7 | 2.9 | NA |
| **stage3_free** | stage3_shell | **-2.69** | **5.47E-03** | **V12B01_19131** | **EAP95164.1** | **hypothetical protein** | | | **-** | **-** | **-** | **66.2** | **1.09** | **9.04E-04** | **3.7** | **2.9** | **NA** | **25.8** | **12.2** | **34.5** |
| **stage2** | stage3_shell | **-2.56** | **3.93E-04** | **V12B01_19261** | **EAP95190.1** | **transcriptional regulator, AraC family** | **Regulation** | **K18954** | **K** | **helix_turn_helix, arabinose operon control protein** | **-** | **13.6** | **0.75** | **4.96E-05** | **4.3** | **6.5** | **4.6** | **15.5** | **18.4** | **48.5** |
| **stage2** | stage3_shell | **-3.27** | **5.36E-10** | **V12B01_19301** | **EAP95198.1** | **putative transmembrane protein** | **Membrane** | **K15269** | **EG** | **of the drug metabolite transporter (DMT) superfamily** | **-** | **39.8** | **0.51** | **1.09E-11** | **8.0** | **13.5** | **9.4** | **98.3** | **57.5** | **82.8** |
| **stage3_free** | stage3_shell | **-2.89** | **1.43E-06** | **V12B01_19301** | **EAP95198.1** | **putative transmembrane protein** | **Membrane** | **K15269** | **EG** | **of the drug metabolite transporter (DMT) superfamily** | **-** | **39.8** | **0.58** | **4.79E-08** | **10.2** | **16.1** | **NA** | **98.3** | **57.5** | **82.8** |
| **stage3_free** | stage3_shell | **-2.00** | **6.83E-05** | **V12B01_19491** | **EAP95236.1** | **putative xanthine dehydrogenase, XdhB subunit** | **Nucleoside and Nucleotide Degradation** | **K13482** | **F** | **COG4631 Xanthine dehydrogenase, molybdopterin-binding subunit B** | **xdhB** | **283.1** | **0.49** | **4.69E-06** | **7.8** | **8.8** | **NA** | **14.6** | **28.9** | **36.7** |
| **stage2** | stage3_shell | **-2.04** | **4.58E-04** | **V12B01_19496** | **EAP95237.1** | **putative xanthine dehydrogenase, XdhA subunit** | **Nucleoside and Nucleotide Degradation** | **K13481** | **F** | **COG4630 Xanthine dehydrogenase, iron-sulfur cluster and FAD-binding subunit A** | **xdhA** | **101.2** | **0.58** | **5.89E-05** | **19.5** | **13.8** | **12.8** | **41.4** | **70.8** | **49.2** |
| **stage3_free** | stage3_shell | **-3.63** | **2.92E-07** | **V12B01_19496** | **EAP95237.1** | **putative xanthine dehydrogenase, XdhA subunit** | **Nucleoside and Nucleotide Degradation** | **K13481** | **F** | **COG4630 Xanthine dehydrogenase, iron-sulfur cluster and FAD-binding subunit A** | **xdhA** | **101.2** | **0.69** | **7.39E-09** | **9.9** | **0.3** | **NA** | **41.4** | **70.8** | **49.2** |
| stage2 | stage3_shell | 3.24 | 1.61E-02 | V12B01_19506 | EAP95239.1 | DNA-3-methyladenine glycosylase | | | L | 3-methyladenine DNA glycosylase | - | 4.1 | 2.21 | 4.27E-03 | 14.2 | 5.2 | 13.6 | 0.0 | 0.0 | 0.0 |
| stage3_free | stage3_shell | 2.80 | 3.29E-02 | V12B01_19506 | EAP95239.1 | DNA-3-methyladenine glycosylase | | | L | 3-methyladenine DNA glycosylase | - | 4.1 | 2.26 | 8.72E-03 | 12.6 | 7.0 | NA | 0.0 | 0.0 | 0.0 |
| stage2 | stage3_shell | 2.44 | 4.17E-02 | V12B01_19511 | EAP95240.1 | putative acetyltransferase | | | J | COG1670 Acetyltransferases, including N-acetylases of ribosomal proteins | - | 3.4 | 2.24 | 1.35E-02 | 16.5 | 11.2 | 4.6 | 0.0 | 0.0 | 0.0 |
| **stage2** | stage3_shell | **-4.04** | **NA** | **V12B01_19691** | **EAP93771.1** | **mannose-6-phosphate isomerase** | **Carbohydrates and Carboxylates Degradation** | **K01809** | **G** | **Belongs to the mannose-6-phosphate isomerase type 1 family** | **manA** | **1.5** | **3.66** | **6.39E-03** | **0.0** | **0.0** | **0.0** | **5.2** | **0.0** | **0.0** |
| stage2 | stage3_shell | 2.10 | 1.58E-06 | V12B01_19751 | EAP93783.1 | methyl-accepting chemotaxis protein | Chemotaxis | | NT | Nitrate and nitrite sensing | - | 64.2 | 0.42 | 8.99E-08 | 67.4 | 63.0 | 69.3 | 6.5 | 13.0 | 12.9 |
| stage3_free | stage3_shell | 2.11 | 7.52E-06 | V12B01_19836 | EAP93800.1 | methyl-accepting chemotaxis protein | Chemotaxis | K03406 | T | methyl-accepting chemotaxis protein | - | 65.7 | 0.45 | 3.39E-07 | 125.2 | 98.0 | NA | 13.5 | 18.2 | 20.6 |
| **stage2** | stage3_shell | **-2.01** | **4.43E-05** | **V12B01_19861** | **EAP93805.1** | **hypothetical protein** | | | **S** | **Domain of unknown function (DUF1611_N) Rossmann-like domain** | **-** | **58.5** | **0.47** | **4.09E-06** | **10.6** | **26.0** | **19.2** | **72.9** | **96.1** | **16.9** |
| **stage3_free** | stage3_shell | **-2.87** | **8.76E-07** | **V12B01_19861** | **EAP93805.1** | **hypothetical protein** | | | **S** | **Domain of unknown function (DUF1611_N) Rossmann-like domain** | **-** | **58.5** | **0.56** | **2.76E-08** | **10.0** | **10.5** | **NA** | **72.9** | **96.1** | **16.9** |
| **stage2** | stage3_shell | **-2.04** | **2.93E-04** | **V12B01_19866** | **EAP93806.1** | **putative muconate cycloisomerase I** | | **K19802** | **M** | **Belongs to the mandelate racemase muconate lactonizing enzyme family** | **-** | **70.9** | **0.55** | **3.56E-05** | **10.4** | **21.5** | **25.3** | **53.4** | **113.4** | **34.6** |
| **stage3_free** | stage3_shell | **-2.54** | **9.56E-05** | **V12B01_19866** | **EAP93806.1** | **putative muconate cycloisomerase I** | | **K19802** | **M** | **Belongs to the mandelate racemase muconate lactonizing enzyme family** | **-** | **70.9** | **0.64** | **7.00E-06** | **13.4** | **13.1** | **NA** | **53.4** | **113.4** | **34.6** |
| **stage3_free** | stage3_shell | **-3.51** | **2.36E-05** | **V12B01_19901** | **EAP93813.1** | **phosphonate ABC transporter, permease protein** | **Transport Proteins** | **K02042** | **P** | **COG3639 ABC-type phosphate phosphonate transport system, permease component** | **-** | **32.9** | **0.84** | **1.30E-06** | **1.1** | **13.7** | **NA** | **63.6** | **112.9** | **56.7** |
| **stage2** | **stage3_free** | **-4.05** | **1.92E-03** | **V12B01_19931** | **EAP93819.1** | **glucose-1-phosphate adenylyltransferase** | | **K00975** | **H** | **Catalyzes the synthesis of ADP-glucose, a sugar donor used in elongation reactions on alpha-glucans** | **glgC2** | **10.8** | **1.29** | **9.17E-05** | **1.5** | **1.2** | **2.0** | **67.0** | **17.3** | **NA** |
| stage3_free | stage3_shell | 3.51 | 8.53E-03 | V12B01_19931 | EAP93819.1 | glucose-1-phosphate adenylyltransferase | | K00975 | H | Catalyzes the synthesis of ADP-glucose, a sugar donor used in elongation reactions on alpha-glucans | glgC2 | 10.8 | 1.65 | 1.57E-03 | 67.0 | 17.3 | NA | 0.0 | 0.0 | 4.7 |
| **stage2** | **stage3_free** | **-5.43** | **6.12E-04** | **V12B01_19936** | **EAP93820.1** | **outer membrane protein OmpW** | **Outer membrane** | **K07275** | **M** | **Outer membrane protein W** | **ompW** | **186.8** | **1.52** | **2.09E-05** | **11.1** | **11.0** | **10.7** | **849.2** | **794.6** | **NA** |
| **stage2** | stage3_shell | **-4.65** | **3.45E-04** | **V12B01_19936** | **EAP93820.1** | **outer membrane protein OmpW** | **Outer membrane** | **K07275** | **M** | **Outer membrane protein W** | **ompW** | **186.8** | **1.44** | **4.29E-05** | **11.1** | **11.0** | **10.7** | **47.3** | **115.5** | **709.6** |
| **stage2** | **stage3_free** | **-4.11** | **2.66E-06** | **V12B01_19941** | **EAP93821.1** | **putative alkaline phosphatase** | **Cofactor, Carrier, and Vitamin Biosynthesis** | | **S** | **alkaline phosphatase** | **-** | **25.2** | **0.82** | **3.75E-08** | **1.5** | **1.4** | **2.8** | **31.3** | **49.4** | **NA** |
| **stage2** | stage3_shell | **-3.18** | **4.09E-05** | **V12B01_19941** | **EAP93821.1** | **putative alkaline phosphatase** | **Cofactor, Carrier, and Vitamin Biosynthesis** | | **S** | **alkaline phosphatase** | **-** | **25.2** | **0.79** | **3.74E-06** | **1.5** | **1.4** | **2.8** | **7.1** | **8.4** | **29.9** |

| | | | | | | | | | | | | | | | | | | | | |
|---|---|---|---|---|---|---|---|---|---|---|---|---|---|---|---|---|---|---|---|---|
| stage3_free | stage3_shell | -3.30 | 6.25E-06 | V12B01_19961 | EAP93825.1 | oxidoreductase, short-chain dehydrogenase/reductase family | | | IQ | KR domain | - | 20.4 | 0.70 | 2.64E-07 | 1.2 | 5.0 | NA | 29.1 | 39.4 | 7.4 |
| stage3_free | stage3_shell | 2.45 | 5.15E-02 | V12B01_19981 | EAP93829.1 | hypothetical protein | | | - | - | - | 3.5 | 2.53 | 1.55E-02 | 11.6 | 4.4 | NA | 0.0 | 0.0 | 0.0 |
| stage3_free | stage3_shell | -2.13 | 1.33E-02 | V12B01_20043 | EAP92403.1 | ISPsy11, transposase OrfA | | K07483 | L | Evidence 2b Function of strongly homologous gene | - | 11.8 | 0.97 | 2.83E-03 | 4.6 | 10.8 | NA | 83.5 | 22.6 | 0.0 |
| stage2 | stage3_shell | -2.06 | 1.88E-05 | V12B01_20053 | EAP92405.1 | predicted PE--lipooligosaccharide phosphorylethanolaminetransferase | | | - | #N/A | #N/A | 41.1 | 0.46 | 1.58E-06 | 6.6 | 12.9 | 10.9 | 55.9 | 39.7 | 7.7 |
| stage3_free | stage3_shell | -2.15 | 9.42E-05 | V12B01_20053 | EAP92405.1 | predicted PE--lipooligosaccharide phosphorylethanolaminetransferase | | | - | #N/A | #N/A | 41.1 | 0.53 | 6.87E-06 | 7.4 | 11.3 | NA | 55.9 | 39.7 | 7.7 |
| stage2 | stage3_shell | -2.12 | 1.49E-03 | V12B01_20058 | EAP92406.1 | hypothetical protein | | | C | glycerophosphoryl diester phosphodiesterase | - | 45.2 | 0.68 | 2.41E-04 | 7.3 | 18.6 | 23.4 | 123.3 | 77.9 | 0.0 |
| stage3_free | stage3_shell | -2.62 | 6.99E-04 | V12B01_20058 | EAP92406.1 | hypothetical protein | | | C | glycerophosphoryl diester phosphodiesterase | - | 45.2 | 0.80 | 7.49E-05 | 8.0 | 14.6 | NA | 123.3 | 77.9 | 0.0 |
| stage3_free | stage3_shell | 2.08 | 4.05E-07 | V12B01_20068 | EAP92408.1 | phosphoenolpyruvate synthase | Glycolysis | K01007 | H | Catalyzes the phosphorylation of pyruvate to phosphoenolpyruvate | ppsA | 255.7 | 0.38 | 1.08E-08 | 255.2 | 178.7 | NA | 37.0 | 31.9 | 34.8 |
| stage3_free | stage3_shell | -2.27 | 1.75E-03 | V12B01_20153 | EAP92425.1 | Uncharacterized protein conserved in archaea | | | O | protein conserved in archaea | - | 51.6 | 0.76 | 2.25E-04 | 23.5 | 29.1 | NA | 59.6 | 95.2 | 185.8 |
| stage3_free | stage3_shell | -2.18 | 7.93E-08 | V12B01_20211 | EAP91959.1 | putative low molecular weight phosphotyrosine protein phosphatase | | K01104 | T | Belongs to the low molecular weight phosphotyrosine protein phosphatase family | - | 86.1 | 0.38 | 1.67E-09 | 61.2 | 84.6 | NA | 303.8 | 326.7 | 140.4 |
| stage3_free | stage3_shell | 2.06 | 2.21E-04 | V12B01_20236 | EAP91964.1 | methionyl-tRNA synthetase | Translation | K01874 | J | Is required not only for elongation of protein synthesis but also for the initiation of all mRNA translation through initiator tRNA(fMet) aminoacylation | metG | 155.7 | 0.53 | 1.90E-05 | 192.8 | 229.3 | NA | 49.6 | 14.7 | 31.8 |
| stage2 | stage3_shell | 2.74 | 8.26E-04 | V12B01_20261 | EAP92862.1 | disulfide bond formation protein B | Translation | K03611 | C | Required for disulfide bond formation in some periplasmic proteins. Acts by oxidizing the DsbA protein | dsbB | 21.3 | 0.81 | 1.18E-04 | 120.3 | 72.9 | 40.4 | 18.3 | 0.0 | 0.0 |
| stage3_free | stage3_shell | 2.40 | 2.83E-04 | V12B01_20326 | EAP92875.1 | oligopeptide ABC transporter, permease protein | Transport Proteins | K15581 | P | COG0601 ABC-type dipeptide oligopeptide nickel transport systems, permease components | oppB | 85.0 | 0.65 | 2.58E-05 | 331.7 | 227.9 | NA | 10.4 | 36.1 | 49.5 |
| stage3_free | stage3_shell | 2.25 | 1.51E-06 | V12B01_20331 | EAP92876.1 | oligopeptide ABC transporter, periplasmic oligopeptide-binding protein | Transport Proteins | K15580 | E | COG4166 ABC-type oligopeptide transport system, periplasmic component | oppA | 917.2 | 0.44 | 5.08E-08 | 1898.1 | 1593.3 | NA | 240.8 | 203.0 | 272.2 |
| stage3_free | stage3_shell | -2.42 | 1.26E-05 | V12B01_20406 | EAP92891.1 | putative transcriptional regulator ToxR | | | K | COG3710 DNA-binding winged-HTH domains | - | 39.5 | 0.53 | 6.17E-07 | 11.8 | 10.6 | NA | 78.6 | 33.5 | 35.0 |
| stage2 | stage3_shell | 3.72 | 7.84E-03 | V12B01_20461 | EAP92902.1 | ribonuclease T | | | L | Trims short 3' overhangs of a variety of RNA species, leaving a one or two nucleotide 3' overhang. Responsible for the end-turnover of tRNA specifically removes the terminal AMP residue from uncharged tRNA (tRNA-C-C-A). Also appears to be involved in tRNA biosynthesis | rnt | 7.0 | 2.19 | 1.75E-03 | 18.6 | 49.4 | 61.0 | 0.0 | 0.0 | 0.0 |
| stage3_free | stage3_shell | 4.07 | 6.78E-03 | V12B01_20461 | EAP92902.1 | ribonuclease T | | | L | Trims short 3' overhangs of a variety of RNA species, leaving a one or two nucleotide 3' overhang. Responsible for the end-turnover of tRNA specifically removes the terminal AMP residue from uncharged tRNA (tRNA-C-C-A). Also appears to be involved in tRNA biosynthesis | rnt | 7.0 | 2.27 | 1.20E-03 | 51.1 | 55.9 | NA | 0.0 | 0.0 | 0.0 |
| stage2 | stage3_shell | 2.67 | 3.71E-07 | V12B01_20468 | EAP91706.1 | superoxide dismutase | | K04564 | C | Destroys radicals which are normally produced within the cells and which are toxic to biological systems | sodB | 248.9 | 0.51 | 1.75E-08 | 882.5 | 397.8 | 463.5 | 95.6 | 46.6 | 38.9 |
| stage2 | stage3_free | -2.13 | 2.03E-04 | V12B01_20513 | EAP92637.1 | Na+/H+ antiporter | Transport Proteins | K03315 | C | Na H antiporter | nhaC | 10.5 | 0.53 | 5.77E-06 | 3.7 | 3.0 | 4.3 | 23.2 | 14.7 | NA |
| stage3_free | stage3_shell | -2.12 | 1.02E-02 | V12B01_20553 | EAP92645.1 | hypothetical protein | | K13771 | K | Nitric oxide-sensitive repressor of genes involved in protecting the cell against nitrosative stress. May require iron for activity | nsrR | 13.8 | 0.89 | 1.99E-03 | 2.1 | 7.6 | NA | 36.0 | 13.6 | 12.8 |
| stage2 | stage3_shell | -3.82 | 7.99E-03 | V12B01_20827 | EAP92930.1 | putative phage gene | | | S | P2 phage tail completion protein R (GpR) | - | 2.7 | 1.87 | 1.78E-03 | 0.0 | 0.7 | 0.0 | 0.0 | 9.2 | 11.6 |
| stage3_free | stage3_shell | -2.41 | 2.39E-04 | V12B01_20862 | EAP92937.1 | putative phage gene | | | S | Ogr/Delta-like zinc finger | - | 44.5 | 0.65 | 2.11E-05 | 16.7 | 31.2 | NA | 162.4 | 98.4 | 69.4 |
| stage3_free | stage3_shell | -2.42 | 1.97E-07 | V12B01_20972 | EAP92959.1 | hypothetical protein | | K09160 | S | Belongs to the UPF0260 family | VV2402 | 56.9 | 0.44 | 4.62E-09 | 34.2 | 63.2 | NA | 275.3 | 209.3 | 135.6 |
| stage2 | stage3_shell | -2.17 | 8.09E-07 | V12B01_21022 | EAP91992.1 | Response regulator | Regulation | K02485 | T | COG0784 FOG CheY-like receiver | rssB | 1145.3 | 0.42 | 4.20E-08 | 145.0 | 232.9 | 239.1 | 1037.7 | 1042.8 | 221.8 |
| stage3_free | stage3_shell | -2.60 | 2.06E-07 | V12B01_21022 | EAP91992.1 | Response regulator | Regulation | K02485 | T | COG0784 FOG CheY-like receiver | rssB | 1145.3 | 0.48 | 4.92E-09 | 154.9 | 149.2 | NA | 1037.7 | 1042.8 | 221.8 |
| stage3_free | stage3_shell | 2.28 | 5.12E-13 | V12B01_21032 | EAP91994.1 | amidophosphoribosyltransferase | Nucleoside and Nucleotide Biosynthesis | K00764 | F | Catalyzes the formation of phosphoribosylamine from phosphoribosylpyrophosphate (PRPP) and glutamine | purF | 183.3 | 0.30 | 2.91E-15 | 410.3 | 306.6 | NA | 59.1 | 46.9 | 48.9 |

| | | | | | | | | | | | | | | | | | | | | |
|---|---|---|---|---|---|---|---|---|---|---|---|---|---|---|---|---|---|---|---|---|
| stage3_free | stage3_shell | 2.13 | 5.47E-05 | V12B01_21117 | EAP91902.1 | methylenetetrahydrofolate dehydrogenase/methenyltetrahydrofolatecyclohydrolase | Cofactor, Carrier, and Vitamin Biosynthesis | K01491 | F | Catalyzes the oxidation of 5,10-methylenetetrahydrofolate to 5,10-methenyltetrahydrofolate and then the hydrolysis of 5,10-methenyltetrahydrofolate to 10-formyltetrahydrofolate | folD | 68.3 | 0.50 | 3.70E-06 | 219.1 | 229.0 | NA | 42.8 | 17.6 | 39.8 |
| stage2 | stage3_shell | -2.25 | 1.36E-06 | V12B01_21194 | EAP92781.1 | hypothetical protein | | | - | - | - | 128.2 | 0.45 | 7.64E-08 | 115.0 | 329.5 | 257.7 | 651.8 | 1400.0 | 651.6 |
| stage2 | stage3_shell | 2.28 | 2.85E-07 | V12B01_21199 | EAP92782.1 | formyltetrahydrofolate deformylase | Cofactor, Carrier, and Vitamin Biosynthesis | K01433 | F | Catalyzes the hydrolysis of 10-formyltetrahydrofolate (formyl-FH4) to formate and tetrahydrofolate (FH4) | purU | 116.4 | 0.43 | 1.30E-08 | 194.8 | 134.5 | 139.7 | 11.4 | 23.4 | 33.9 |
| stage3_free | stage3_shell | 2.28 | 2.59E-06 | V12B01_21199 | EAP92782.1 | formyltetrahydrofolate deformylase | Cofactor, Carrier, and Vitamin Biosynthesis | K01433 | F | Catalyzes the hydrolysis of 10-formyltetrahydrofolate (formyl-FH4) to formate and tetrahydrofolate (FH4) | purU | 116.4 | 0.46 | 9.29E-08 | 177.5 | 143.5 | NA | 11.4 | 23.4 | 33.9 |
| stage3_free | stage3_shell | 2.06 | 1.53E-05 | V12B01_21209 | EAP92784.1 | arginyl-tRNA synthetase | Other Biosynthesis | K01887 | J | Arginyl-tRNA synthetase | argS | 195.0 | 0.45 | 7.81E-07 | 249.5 | 359.6 | NA | 82.0 | 36.6 | 26.3 |
| stage2 | stage3_shell | -3.69 | 2.92E-07 | V12B01_21219 | EAP92786.1 | hypothetical protein | | | S | FeoC like transcriptional regulator | VV1042 | 50.8 | 0.71 | 1.34E-08 | 19.2 | 32.2 | 36.5 | 332.0 | 58.9 | 542.3 |
| stage2 | stage3_shell | -4.39 | 3.23E-10 | V12B01_21224 | EAP92787.1 | Fe2+ transport system protein B | Transport Proteins | K04759 | P | transporter of a GTP-driven Fe(2 ) uptake system | feoB | 378.5 | 0.68 | 6.20E-12 | 12.5 | 17.3 | 15.6 | 219.2 | 65.2 | 453.3 |
| stage3_free | stage3_shell | -2.69 | 4.27E-04 | V12B01_21224 | EAP92787.1 | Fe2+ transport system protein B | Transport Proteins | K04759 | P | transporter of a GTP-driven Fe(2 ) uptake system | feoB | 378.5 | 0.80 | 4.26E-05 | 30.2 | 59.4 | NA | 219.2 | 65.2 | 453.3 |
| stage2 | stage3_shell | -4.77 | 1.12E-10 | V12B01_21229 | EAP92788.1 | Fe2+ transport system protein A | Transport Proteins | K04758 | P | Fe2 transport system protein A | feoA | 105.2 | 0.72 | 1.89E-12 | 24.0 | 25.5 | 54.8 | 475.0 | 374.8 | 1385.1 |
| stage3_free | stage3_shell | -4.10 | 8.03E-07 | V12B01_21229 | EAP92788.1 | Fe2+ transport system protein A | Transport Proteins | K04758 | P | Fe2 transport system protein A | feoA | 105.2 | 0.83 | 2.49E-08 | 61.8 | 43.3 | NA | 475.0 | 374.8 | 1385.1 |
| stage3_free | stage3_shell | -2.97 | 2.24E-02 | V12B01_21234 | EAP92789.1 | hypothetical protein | | | - | #N/A | #N/A | 4.4 | 1.86 | 5.45E-03 | 2.6 | 2.7 | NA | 27.1 | 25.6 | 64.3 |
| stage2 | stage3_shell | 2.33 | 2.61E-06 | V12B01_21259 | EAP92794.1 | succinyl-CoA synthetase alpha subunit | TCA cycle | K01902 | C | Succinyl-CoA synthetase functions in the citric acid cycle (TCA), coupling the hydrolysis of succinyl-CoA to the synthesis of either ATP or GTP and thus represents the only step of substrate-level phosphorylation in the TCA. The alpha subunit of the enzyme binds the substrates coenzyme A and phosphate, while succinate binding and nucleotide specificity is provided by the beta subunit | sucD | 563.1 | 0.48 | 1.58E-07 | 2242.7 | 1011.0 | 1181.0 | 87.8 | 190.6 | 283.7 |
| stage2 | stage3_shell | 2.13 | 1.25E-03 | V12B01_21264 | EAP92795.1 | succinyl-CoA synthetase subunit beta | TCA cycle | K01903 | F | Succinyl-CoA synthetase functions in the citric acid cycle (TCA), coupling the hydrolysis of succinyl-CoA to the synthesis of either ATP or GTP and thus represents the only step of substrate-level phosphorylation in the TCA. The beta subunit provides nucleotide specificity of the enzyme and binds the substrate succinate, while the binding sites for coenzyme A and phosphate are found in the alpha subunit | sucC | 573.0 | 0.67 | 1.94E-04 | 2144.5 | 730.8 | 780.5 | 27.4 | 142.6 | 295.2 |
| stage2 | stage3_shell | 2.39 | 1.98E-04 | V12B01_21274 | EAP92797.1 | 2-oxoglutarate dehydrogenase complex, dehydrogenase (E1) component | | K00164 | C | COG0567 2-oxoglutarate dehydrogenase complex, dehydrogenase (E1) component, and related enzymes | sucA | 1151.0 | 0.65 | 2.27E-05 | 1546.9 | 540.7 | 567.1 | 24.4 | 81.5 | 181.7 |
| stage3_free | stage3_shell | 2.99 | 3.32E-05 | V12B01_21299 | EAP92802.1 | citrate synthase | TCA cycle | K01647 | C | Belongs to the citrate synthase family | gltA | 855.2 | 0.70 | 2.04E-06 | 2408.7 | 1807.1 | NA | 126.3 | 42.2 | 278.1 |
| stage2 | stage3_shell | 2.39 | 4.88E-04 | V12B01_21349 | EAP92812.1 | hypothetical protein | | | - | #N/A | #N/A | 24.0 | 0.66 | 6.36E-05 | 439.6 | 382.7 | 441.2 | 18.4 | 69.6 | 65.5 |
| stage2 | stage3_shell | 2.45 | 3.43E-05 | V12B01_21384 | EAP92476.1 | asparagine synthetase B | Amino Acid Degradation | K01953 | E | Asparagine synthetase B | asnB | 307.2 | 0.58 | 3.08E-06 | 367.5 | 210.0 | 218.3 | 15.3 | 37.1 | 39.2 |
| stage3_free | stage3_shell | 3.47 | 2.37E-07 | V12B01_21384 | EAP92476.1 | asparagine synthetase B | Amino Acid Degradation | K01953 | E | Asparagine synthetase B | asnB | 307.2 | 0.64 | 5.75E-09 | 857.5 | 229.4 | NA | 15.3 | 37.1 | 39.2 |
| stage2 | stage3_shell | -3.83 | 4.41E-09 | V12B01_21394 | EAP92478.1 | dipeptide/tripeptide permease | Transport Proteins | K03305 | U | COG3104 Dipeptide tripeptide permease | - | 53.5 | 0.63 | 1.13E-10 | 2.6 | 6.1 | 0.5 | 55.3 | 37.1 | 16.4 |
| stage2 | stage3_free | -2.32 | 1.77E-02 | V12B01_21404 | EAP92480.1 | adenylate kinase | Nucleoside and Nucleotide Biosynthesis | K00939 | F | Catalyzes the reversible transfer of the terminal phosphate group between ATP and AMP. Plays an important role in cellular energy homeostasis and in adenine nucleotide metabolism | adk | 273.6 | 1.04 | 1.36E-03 | 319.2 | 197.1 | 203.7 | 2025.1 | 1878.3 | NA |
| stage3_free | stage3_shell | 2.01 | 2.33E-02 | V12B01_21404 | EAP92480.1 | adenylate kinase | Nucleoside and Nucleotide Biosynthesis | K00939 | F | Catalyzes the reversible transfer of the terminal phosphate group between ATP and AMP. Plays an important role in cellular energy homeostasis and in adenine nucleotide metabolism | adk | 273.6 | 0.98 | 5.75E-03 | 2025.1 | 1878.3 | NA | 649.0 | 14.1 | 61.8 |
| stage2 | stage3_shell | 2.22 | 2.93E-04 | V12B01_21409 | EAP92481.1 | heat shock protein 90 | | K04079 | O | Molecular chaperone. Has ATPase activity | htpG | 115.2 | 0.61 | 3.56E-05 | 121.4 | 24.5 | 199.4 | 15.1 | 12.7 | 17.9 |
| stage2 | stage3_shell | -2.00 | 3.90E-03 | V12B01_21424 | EAP92484.1 | selenoprotein W-related protein | | K07401 | O | selenoprotein W-related protein | - | 11.8 | 0.72 | 7.52E-04 | 11.1 | 19.9 | 15.6 | 89.6 | 56.6 | 26.6 |
| stage2 | stage3_shell | -2.06 | 2.32E-06 | V12B01_21429 | EAP92485.1 | predicted phosphoesterase | | K07095 | S | Phosphoesterase | yfcE | 171.2 | 0.42 | 1.40E-07 | 104.2 | 157.0 | 138.9 | 324.8 | 274.3 | 667.5 |
| stage2 | stage3_shell | -3.16 | 5.26E-08 | V12B01_21449 | EAP92489.1 | Permease of the major facilitator superfamily | Transport Proteins | | S | Protein of unknown function (DUF1538) | - | 385.0 | 0.57 | 1.94E-09 | 92.6 | 150.8 | 163.0 | 384.8 | 874.8 | 1545.6 |
| stage3_free | stage3_shell | -2.56 | 8.23E-05 | V12B01_21449 | EAP92489.1 | Permease of the major facilitator superfamily | Transport Proteins | | S | Protein of unknown function (DUF1538) | - | 385.0 | 0.65 | 5.85E-06 | 172.4 | 219.6 | NA | 384.8 | 874.8 | 1545.6 |
| stage2 | stage3_shell | -2.20 | 7.75E-14 | V12B01_21454 | EAP92490.1 | putative sugar nucleotide epimerase | | K07071 | S | nucleoside-diphosphate sugar epimerase | yfcH | 362.7 | 0.28 | 7.68E-16 | 67.6 | 100.3 | 125.5 | 268.9 | 444.8 | 304.9 |

| | | | | | | | | | | | | | | | | | | | |
|---|---|---|---|---|---|---|---|---|---|---|---|---|---|---|---|---|---|---|---|
| stage3_free | stage3_shell | -2.04 | 1.05E-09 | V12B01_21454 | EAP92490.1 | putative sugar nucleotide epimerase | | K07071 | S | nucleoside-diphosphate sugar epimerase | yfcH | 362.7 | 0.32 | 1.39E-11 | 93.5 | 123.9 | NA | 268.9 | 444.8 | 304.9 |
| stage2 | stage3_shell | -2.68 | 1.36E-04 | V12B01_21484 | EAP92496.1 | hypothetical protein | | | - | - | - | 114.1 | 0.71 | 1.48E-05 | 35.2 | 298.1 | 160.4 | 1606.8 | 1256.5 | 128.7 |
| stage3_free | stage3_shell | -3.37 | 3.20E-05 | V12B01_21484 | EAP92496.1 | hypothetical protein | | | - | - | - | 114.1 | 0.82 | 1.95E-06 | 87.9 | 114.0 | NA | 1606.8 | 1256.5 | 128.7 |
| stage2 | stage3_shell | 3.10 | 6.04E-06 | V12B01_21519 | EAP92503.1 | cysteine synthase A | Amino Acid Biosynthesis | K01738 | E | Belongs to the cysteine synthase cystathionine beta- synthase family | cysK | 124.1 | 0.67 | 4.03E-07 | 1010.8 | 580.6 | 1440.9 | 69.8 | 115.7 | 31.1 |
| stage3_free | stage3_shell | 2.49 | 8.40E-04 | V12B01_21519 | EAP92503.1 | cysteine synthase A | Amino Acid Biosynthesis | K01738 | E | Belongs to the cysteine synthase cystathionine beta- synthase family | cysK | 124.1 | 0.73 | 9.30E-05 | 817.2 | 601.1 | NA | 69.8 | 115.7 | 31.1 |
| stage2 | stage3_shell | 2.02 | 4.70E-04 | V12B01_21546 | EAP93683.1 | flagellin | Motility | | - | #N/A | #N/A | 36.3 | 0.56 | 6.09E-05 | 592.9 | 651.2 | 446.3 | 85.9 | 113.9 | 61.2 |
| stage2 | stage3_shell | -5.80 | NA | V12B01_21581 | EAP93690.1 | hypothetical protein | | | S | protein conserved in bacteria | - | 843.0 | 1.48 | NA | 27.8 | 157.4 | 92.1 | 1642.1 | 1560.2 | 12575.2 |
| stage2 | stage3_shell | 2.02 | 2.51E-03 | V12B01_21586 | EAP93691.1 | putative arsenate reductase | | | P | Belongs to the ArsC family | yffB | 23.0 | 0.65 | 4.45E-04 | 117.6 | 90.6 | 62.8 | 27.5 | 8.7 | 0.0 |
| stage3_free | stage3_shell | 2.60 | 9.08E-04 | V12B01_21616 | EAP93697.1 | putative glycine cleavage system transcriptional repressor | Regulation | K03567 | E | glycine cleavage system | gcvR | 28.7 | 0.76 | 1.02E-04 | 114.5 | 103.5 | NA | 17.7 | 11.1 | 0.0 |
| stage3_free | stage3_shell | -2.06 | 3.84E-05 | V12B01_21636 | EAP93701.1 | hypothetical protein | | | - | - | - | 179.7 | 0.48 | 2.45E-06 | 26.2 | 31.3 | NA | 42.1 | 168.9 | 79.5 |
| stage2 | stage3_shell | -5.14 | NA | V12B01_21671 | EAP93708.1 | hypothetical protein | | | - | #N/A | #N/A | 1.3 | 3.29 | 2.18E-03 | 0.0 | 0.0 | 0.0 | 25.4 | 48.0 | 0.0 |
| stage2 | stage3_shell | 2.15 | 2.04E-05 | V12B01_21686 | EAP93711.1 | probable cysteine synthase A | Amino Acid Biosynthesis | K01738 | E | Belongs to the cysteine synthase cystathionine beta- synthase family | - | 97.3 | 0.50 | 1.74E-06 | 286.2 | 187.2 | 139.6 | 21.0 | 29.9 | 40.6 |
| stage3_free | stage3_shell | 2.32 | 3.64E-05 | V12B01_21686 | EAP93711.1 | probable cysteine synthase A | Amino Acid Biosynthesis | K01738 | E | Belongs to the cysteine synthase cystathionine beta- synthase family | - | 97.3 | 0.54 | 2.30E-06 | 326.5 | 145.9 | NA | 21.0 | 29.9 | 40.6 |
| stage2 | stage3_shell | -4.18 | 5.99E-17 | V12B01_21706 | EAP93715.1 | Transposase | | K07486 | L | COG3547 Transposase and inactivated derivatives | - | 82.1 | 0.48 | 3.16E-19 | 3.9 | 11.4 | 12.8 | 77.9 | 125.3 | 188.7 |
| stage3_free | stage3_shell | -2.52 | 4.65E-06 | V12B01_21706 | EAP93715.1 | Transposase | | K07486 | L | COG3547 Transposase and inactivated derivatives | - | 82.1 | 0.53 | 1.82E-07 | 29.4 | 28.1 | NA | 77.9 | 125.3 | 188.7 |
| stage2 | stage3_shell | -2.23 | 2.03E-06 | V12B01_21746 | EAP93723.1 | bacterioferritin comigratory protein | | | O | COG1225 Peroxiredoxin | bcp | 170.3 | 0.45 | 1.17E-07 | 151.4 | 341.3 | 212.6 | 1308.3 | 997.6 | 412.6 |
| stage3_free | stage3_shell | -3.10 | 1.66E-08 | V12B01_21746 | EAP93723.1 | bacterioferritin comigratory protein | | | O | COG1225 Peroxiredoxin | bcp | 170.3 | 0.53 | 2.83E-10 | 96.1 | 163.6 | NA | 1308.3 | 997.6 | 412.6 |
| stage2 | stage3_shell | -2.00 | 2.22E-03 | V12B01_21761 | EAP93726.1 | Predicted redox protein | | K04085 | O | Belongs to the sulfur carrier protein TusA family | - | 15.4 | 0.66 | 3.85E-04 | 11.1 | 38.4 | 53.3 | 99.9 | 189.1 | 83.0 |
| stage3_free | stage3_shell | -2.78 | 3.77E-04 | V12B01_21761 | EAP93726.1 | Predicted redox protein | | K04085 | O | Belongs to the sulfur carrier protein TusA family | - | 15.4 | 0.80 | 3.70E-05 | 19.1 | 20.0 | NA | 99.9 | 189.1 | 83.0 |
| stage2 | stage3_shell | 2.05 | 1.26E-02 | V12B01_21771 | EAP93728.1 | arsenate reductase | | K00537 | C | COG1393 Arsenate reductase and related proteins, glutaredoxin family | arsC | 13.7 | 0.87 | 3.11E-03 | 114.0 | 44.4 | 47.1 | 18.2 | 0.0 | 8.1 |
| stage2 | stage3_shell | 3.13 | 8.81E-04 | V12B01_21776 | EAP93729.1 | Trp repressor binding protein WrbA | | K03809 | S | COG0655 Multimeric flavodoxin WrbA | wrbA | 18.6 | 0.92 | 1.28E-04 | 71.1 | 40.5 | 44.1 | 0.0 | 2.5 | 9.6 |
| stage3_free | stage3_shell | 2.19 | 1.76E-02 | V12B01_21776 | EAP93729.1 | Trp repressor binding protein WrbA | | K03809 | S | COG0655 Multimeric flavodoxin WrbA | wrbA | 18.6 | 0.95 | 4.03E-03 | 37.0 | 23.5 | NA | 0.0 | 2.5 | 9.6 |
| stage2 | stage3_shell | 2.44 | 2.16E-02 | V12B01_21781 | EAP93730.1 | Predicted membrane protein | Membrane | | S | Predicted membrane protein (DUF2069) | - | 6.0 | 1.16 | 6.07E-03 | 48.7 | 27.5 | 27.1 | 0.0 | 6.9 | 0.0 |
| stage3_free | stage3_shell | 2.43 | 1.77E-03 | V12B01_21791 | EAP93732.1 | uracil permease | Transport Proteins | K02824 | F | COG2233 Xanthine uracil permeases | uraA | 31.9 | 0.79 | 2.27E-04 | 73.8 | 129.1 | NA | 10.3 | 3.7 | 18.4 |
| stage2 | stage3_shell | 2.42 | 4.50E-08 | V12B01_21801 | EAP93734.1 | phosphoribosylaminoimidazole synthetase | Nucleoside and Nucleotide Biosynthesis | K01933 | F | Phosphoribosylformylglycinamidine cyclo-ligase | purM | 231.2 | 0.42 | 1.55E-09 | 375.0 | 287.5 | 258.1 | 58.3 | 23.3 | 35.6 |
| stage3_free | stage3_shell | 3.91 | 1.15E-15 | V12B01_21801 | EAP93734.1 | phosphoribosylaminoimidazole synthetase | Nucleoside and Nucleotide Biosynthesis | K01933 | F | Phosphoribosylformylglycinamidine cyclo-ligase | purM | 231.2 | 0.46 | 2.44E-18 | 871.2 | 855.1 | NA | 58.3 | 23.3 | 35.6 |
| stage2 | stage3_shell | 2.02 | 3.38E-02 | V12B01_21806 | EAP93735.1 | phosphoribosylglycinamide formyltransferase | Nucleoside and Nucleotide Biosynthesis | K11175 | F | Catalyzes the transfer of a formyl group from 10- formyltetrahydrofolate to 5-phospho-ribosyl-glycinamide (GAR), producing 5-phospho-ribosyl-N-formylglycinamide (FGAR) and tetrahydrofolate | purN | 9.3 | 1.20 | 1.04E-02 | 46.8 | 32.5 | 11.6 | 2.4 | 0.0 | 8.4 |
| stage3_free | stage3_shell | 2.24 | 3.07E-02 | V12B01_21806 | EAP93735.1 | phosphoribosylglycinamide formyltransferase | Nucleoside and Nucleotide Biosynthesis | K11175 | F | Catalyzes the transfer of a formyl group from 10- formyltetrahydrofolate to 5-phospho-ribosyl-glycinamide (GAR), producing 5-phospho-ribosyl-N-formylglycinamide (FGAR) and tetrahydrofolate | purN | 9.3 | 1.28 | 8.06E-03 | 29.8 | 44.8 | NA | 2.4 | 0.0 | 8.4 |
| stage2 | stage3_shell | 2.63 | 9.36E-08 | V12B01_21951 | EAP93764.1 | ribosome releasing factor | Translation | K02838 | J | Responsible for the release of ribosomes from messenger RNA at the termination of protein biosynthesis. May increase the efficiency of translation by recycling ribosomes from one round of translation to another | frr | 77.2 | 0.47 | 3.67E-09 | 316.4 | 178.4 | 186.8 | 31.5 | 21.7 | 20.4 |
| stage3_free | stage3_shell | 3.21 | 1.75E-09 | V12B01_21951 | EAP93764.1 | ribosome releasing factor | Translation | K02838 | J | Responsible for the release of ribosomes from messenger RNA at the termination of protein biosynthesis. May increase the efficiency of translation by recycling ribosomes from one round of translation to another | frr | 77.2 | 0.50 | 2.48E-11 | 390.1 | 295.3 | NA | 31.5 | 21.7 | 20.4 |
| stage3_free | stage3_shell | 2.38 | 6.31E-06 | V12B01_21956 | EAP91876.1 | elongation factor Ts | Translation | K02357 | J | Associates with the EF-Tu.GDP complex and induces the exchange of GDP to GTP. It remains bound to the aminoacyl-tRNA.EF- Tu.GTP complex up to the GTP hydrolysis stage on the ribosome | tsf | 244.7 | 0.50 | 2.67E-07 | 1798.9 | 1564.3 | NA | 151.6 | 209.5 | 262.8 |
| stage3_free | stage3_shell | 2.22 | 2.23E-04 | V12B01_21961 | EAP91877.1 | 30S ribosomal protein S2 | Translation | K02967 | J | Belongs to the universal ribosomal protein uS2 family | rpsB | 1028.7 | 0.58 | 1.93E-05 | 3946.3 | 3629.8 | NA | 547.8 | 393.1 | 561.7 |
| stage2 | stage3_shell | 2.71 | 3.12E-02 | V12B01_21996 | EAP91884.1 | hypothetical protein | | | S | Protein of unknown function (DUF3301) | - | 3.6 | 2.22 | 9.45E-03 | 24.5 | 10.3 | 25.2 | 0.0 | 0.0 | 0.0 |

| | | | | | | | | | | | | | | | | | | | | |
|---|---|---|---|---|---|---|---|---|---|---|---|---|---|---|---|---|---|---|---|---|
| stage2 | stage3_free | -2.61 | 2.10E-04 | V12B01_22001 | EAP91885.1 | acetyltransferase-related protein | | K03790 | J | COG1670 Acetyltransferases, including N-acetylases of ribosomal proteins | - | 236.0 | 0.66 | 6.02E-06 | 133.7 | 150.8 | 203.2 | 1246.9 | 1197.7 | NA |
| stage2 | stage3_free | -2.58 | 2.20E-04 | V12B01_22021 | EAP91889.1 | putative efflux pump component MtrF | Transport Proteins | K12942 | H | p-aminobenzoyl-glutamate transporter | - | 22.6 | 0.66 | 6.41E-06 | 6.7 | 5.4 | 8.7 | 68.0 | 33.9 | NA |
| stage2 | stage3_shell | -2.31 | 4.17E-09 | V12B01_22061 | EAP91852.1 | PTS system, trehalose-specific IIBC component | Transport Proteins | K02819 | G | pts system | treB | 83.2 | 0.38 | 1.07E-10 | 16.9 | 9.2 | 11.4 | 35.9 | 70.1 | 38.0 |
| stage3_free | stage3_shell | 2.78 | 6.34E-06 | V12B01_22261 | EAP94709.1 | hypothetical protein | | K06901 | S | COG2252 Permeases | yieG | 115.0 | 0.58 | 2.70E-07 | 266.3 | 325.5 | NA | 57.0 | 14.1 | 11.0 |
| stage2 | stage3_free | -2.15 | 2.34E-04 | V12B01_22271 | EAP94711.1 | aminoacyl-histidine dipeptidase | | K01270 | E | aminoacyl-histidine dipeptidase | pepD | 152.9 | 0.54 | 6.89E-06 | 116.8 | 38.7 | 44.3 | 382.8 | 314.7 | NA |
| stage3_free | stage3_shell | 2.28 | 7.62E-05 | V12B01_22271 | EAP94711.1 | aminoacyl-histidine dipeptidase | | K01270 | E | aminoacyl-histidine dipeptidase | pepD | 152.9 | 0.56 | 5.35E-06 | 382.8 | 314.7 | NA | 26.0 | 32.9 | 73.5 |
| stage2 | stage3_shell | 6.05 | 1.45E-04 | V12B01_22276 | EAP94712.1 | hypothetical protein | | | S | Domain of unknown function (DUF3332) | - | 13.5 | 2.38 | 1.58E-05 | 47.8 | 59.7 | 68.2 | 0.0 | 0.0 | 0.0 |
| stage3_free | stage3_shell | 2.19 | 5.45E-02 | V12B01_22276 | EAP94712.1 | hypothetical protein | | | S | Domain of unknown function (DUF3332) | - | 13.5 | 2.18 | 1.67E-02 | 11.9 | 8.9 | NA | 0.0 | 0.0 | 0.0 |
| stage2 | stage3_shell | -2.35 | 4.39E-04 | V12B01_22291 | EAP94715.1 | adenosine deaminase | Nucleoside and Nucleotide Degradation | K21053 | F | Catalyzes the hydrolytic deamination of adenine to hypoxanthine. Plays an important role in the purine salvage pathway and in nitrogen catabolism | - | 46.3 | 0.68 | 5.60E-05 | 6.2 | 4.2 | 5.2 | 12.6 | 29.9 | 28.2 |
| stage3_free | stage3_shell | 2.37 | 9.22E-04 | V12B01_22311 | EAP94719.1 | transcriptional regulator, LacI family | Regulation | K02529 | K | transcriptional regulators | galR | 21.2 | 0.68 | 1.04E-04 | 41.7 | 38.9 | NA | 6.4 | 6.1 | 0.0 |
| stage2 | stage3_shell | -2.44 | 1.68E-08 | V12B01_22316 | EAP94720.1 | Galactose-1-epimerase | Carbohydrates and Carboxylates Degradation | K01785 | G | Converts alpha-aldose to the beta-anomer | galM | 623.6 | 0.42 | 5.24E-10 | 153.2 | 346.6 | 310.9 | 1130.3 | 1697.8 | 717.1 |
| stage3_free | stage3_shell | -2.54 | 2.37E-07 | V12B01_22316 | EAP94720.1 | Galactose-1-epimerase | Carbohydrates and Carboxylates Degradation | K01785 | G | Converts alpha-aldose to the beta-anomer | galM | 623.6 | 0.47 | 5.72E-09 | 258.3 | 241.1 | NA | 1130.3 | 1697.8 | 717.1 |
| stage2 | stage3_shell | -2.02 | 4.44E-04 | V12B01_22391 | EAP94735.1 | putative sugar ABC transporter ATP-binding protein | Transport Proteins | K10111 | P | Belongs to the ABC transporter superfamily | - | 79.8 | 0.57 | 5.69E-05 | 29.6 | 27.5 | 21.9 | 56.1 | 125.7 | 89.4 |
| stage2 | stage3_shell | -2.00 | 4.00E-02 | V12B01_22401 | EAP94737.1 | hypothetical protein | | | S | Domain of unknown function (DUF1330) | - | 15.1 | 1.29 | 1.27E-02 | 0.0 | 2.3 | 4.1 | 26.4 | 10.0 | 0.0 |
| stage3_free | stage3_shell | -2.55 | 1.51E-04 | V12B01_22406 | EAP94738.1 | putative sodium/myo-inositol cotransporter | Transport Proteins | K03307 | S | Belongs to the sodium solute symporter (SSF) (TC 2.A.21) family | - | 32.9 | 0.67 | 1.21E-05 | 4.6 | 7.9 | NA | 43.3 | 47.2 | 6.7 |
| stage2 | stage3_shell | 3.91 | 6.64E-03 | V12B01_22426 | EAP94742.1 | Chromate transport protein ChrA | Transport Proteins | K07240 | P | COG2059 Chromate transport protein ChrA | - | 5.5 | 2.25 | 1.43E-03 | 11.6 | 8.3 | 5.4 | 0.0 | 0.0 | 0.0 |
| stage3_free | stage3_shell | 4.26 | 5.81E-03 | V12B01_22426 | EAP94742.1 | Chromate transport protein ChrA | Transport Proteins | K07240 | P | COG2059 Chromate transport protein ChrA | - | 5.5 | 2.33 | 9.75E-04 | 8.8 | 12.1 | NA | 0.0 | 0.0 | 0.0 |
| stage2 | stage3_shell | -3.11 | 1.91E-08 | V12B01_22436 | EAP94744.1 | hypothetical protein | | | S | oligosaccharyl transferase activity | - | 132.5 | 0.54 | 6.01E-10 | 8.8 | 26.9 | 19.7 | 200.6 | 180.6 | 28.9 |
| stage3_free | stage3_shell | -3.33 | 1.47E-07 | V12B01_22436 | EAP94744.1 | hypothetical protein | | | S | oligosaccharyl transferase activity | - | 132.5 | 0.61 | 3.38E-09 | 11.8 | 19.5 | NA | 200.6 | 180.6 | 28.9 |
| stage2 | stage3_shell | -2.90 | 1.36E-08 | V12B01_22441 | EAP94745.1 | TPR repeat protein | | | S | COG0457 FOG TPR repeat | VP2409 | 424.3 | 0.49 | 4.08E-10 | 23.5 | 54.3 | 54.6 | 284.6 | 492.2 | 59.5 |
| stage3_free | stage3_shell | -2.84 | 9.82E-07 | V12B01_22441 | EAP94745.1 | TPR repeat protein | | | S | COG0457 FOG TPR repeat | VP2409 | 424.3 | 0.56 | 3.17E-08 | 46.0 | 44.5 | NA | 284.6 | 492.2 | 59.5 |
| stage2 | stage3_shell | -3.37 | 4.00E-16 | V12B01_22446 | EAP94746.1 | Flp pilus assembly protein TadC | Membrane | K12511 | NU | COG2064 Flp pilus assembly protein TadC | tadC | 498.1 | 0.40 | 2.53E-18 | 16.0 | 39.0 | 33.7 | 207.1 | 404.7 | 117.2 |
| stage3_free | stage3_shell | -2.28 | 1.07E-06 | V12B01_22446 | EAP94746.1 | Flp pilus assembly protein TadC | Membrane | K12511 | NU | COG2064 Flp pilus assembly protein TadC | tadC | 498.1 | 0.45 | 3.51E-08 | 62.6 | 60.8 | NA | 207.1 | 404.7 | 117.2 |
| stage2 | stage3_shell | -3.24 | 7.73E-15 | V12B01_22451 | EAP94747.1 | Flp pilus assembly protein TadB | Membrane | K12510 | U | COG4965 Flp pilus assembly protein TadB | tadB | 560.1 | 0.40 | 6.36E-17 | 24.4 | 29.3 | 37.1 | 175.3 | 419.7 | 91.4 |
| stage3_free | stage3_shell | -2.15 | 5.00E-06 | V12B01_22451 | EAP94747.1 | Flp pilus assembly protein TadB | Membrane | K12510 | U | COG4965 Flp pilus assembly protein TadB | tadA | 560.1 | 0.45 | 1.99E-07 | 59.7 | 66.1 | NA | 175.3 | 419.7 | 91.4 |
| stage2 | stage3_shell | -3.90 | 4.27E-36 | V12B01_22456 | EAP94748.1 | Flp pilus assembly protein TadA | Membrane | K02283 | U | COG4962 Flp pilus assembly protein, ATPase CpaF | tadA | 1428.7 | 0.30 | 9.01E-40 | 44.5 | 99.2 | 89.2 | 943.9 | 1175.6 | 567.9 |
| stage3_free | stage3_shell | -3.59 | 5.77E-24 | V12B01_22456 | EAP94748.1 | Flp pilus assembly protein TadA | Membrane | K02283 | U | COG4962 Flp pilus assembly protein, ATPase CpaF | tadA | 1428.7 | 0.34 | 1.22E-27 | 79.9 | 113.0 | NA | 943.9 | 1175.6 | 567.9 |
| stage2 | stage3_shell | -3.63 | 7.87E-22 | V12B01_22461 | EAP94749.1 | Flp pilus assembly protein | Membrane | K02282 | U | COG4963 Flp pilus assembly protein, ATPase CpaE | cpaE | 1985.9 | 0.36 | 1.83E-24 | 82.7 | 201.0 | 230.5 | 1979.9 | 2229.0 | 813.8 |
| stage3_free | stage3_shell | -3.97 | 3.24E-20 | V12B01_22461 | EAP94749.1 | Flp pilus assembly protein | Membrane | K02282 | U | COG4963 Flp pilus assembly protein, ATPase CpaE | cpaE | 1985.9 | 0.41 | 2.05E-23 | 117.2 | 152.6 | NA | 1979.9 | 2229.0 | 813.8 |
| stage2 | stage3_shell | -3.97 | 4.53E-20 | V12B01_22466 | EAP94750.1 | hypothetical protein | | | U | TadE-like protein | VP2414 | 578.5 | 0.42 | 1.53E-22 | 22.3 | 58.3 | 90.9 | 805.3 | 960.0 | 365.9 |
| stage3_free | stage3_shell | -4.17 | 1.28E-16 | V12B01_22466 | EAP94750.1 | hypothetical protein | | | U | TadE-like protein | VP2414 | 578.5 | 0.48 | 1.35E-19 | 36.7 | 62.7 | NA | 805.3 | 960.0 | 365.9 |
| stage2 | stage3_shell | -4.74 | 7.12E-30 | V12B01_22471 | EAP94751.1 | hypothetical protein | | | U | TadE-like protein | - | 412.3 | 0.40 | 4.51E-33 | 17.6 | 47.9 | 24.7 | 696.5 | 906.3 | 298.7 |
| stage3_free | stage3_shell | -4.62 | 2.10E-21 | V12B01_22471 | EAP94751.1 | hypothetical protein | | | U | TadE-like protein | - | 412.3 | 0.46 | 8.85E-25 | 38.4 | 26.9 | NA | 696.5 | 906.3 | 298.7 |
| stage2 | stage3_shell | -4.99 | 1.76E-24 | V12B01_22476 | EAP94752.1 | hypothetical protein | | | S | Putative Flp pilus-assembly TadE/G-like | - | 1586.3 | 0.47 | 2.60E-27 | 13.0 | 35.5 | 36.2 | 598.2 | 1371.9 | 236.1 |
| stage3_free | stage3_shell | -4.57 | 2.89E-16 | V12B01_22476 | EAP94752.1 | hypothetical protein | | | S | Putative Flp pilus-assembly TadE/G-like | - | 1586.3 | 0.53 | 4.88E-19 | 38.3 | 36.8 | NA | 598.2 | 1371.9 | 236.1 |
| stage2 | stage3_shell | -5.56 | 1.73E-20 | V12B01_22481 | EAP94753.1 | hypothetical protein | | | - | - | - | 325.5 | 0.58 | 4.74E-23 | 9.9 | 28.9 | 45.1 | 650.9 | 2370.2 | 316.4 |
| stage3_free | stage3_shell | -5.13 | 6.72E-14 | V12B01_22481 | EAP94753.1 | hypothetical protein | | | - | - | - | 325.5 | 0.66 | 2.83E-16 | 32.2 | 42.7 | NA | 650.9 | 2370.2 | 316.4 |
| stage2 | stage3_shell | -4.51 | 4.69E-09 | V12B01_22486 | EAP94754.1 | Flp pilus assembly protein | Membrane | K02280 | U | Belongs to the GSP D family | - | 1543.6 | 0.75 | 1.25E-10 | 6.7 | 29.5 | 31.7 | 319.6 | 1031.5 | 68.5 |
| stage3_free | stage3_shell | -2.80 | 7.89E-04 | V12B01_22486 | EAP94754.1 | Flp pilus assembly protein | Membrane | K02280 | U | Belongs to the GSP D family | - | 1543.6 | 0.90 | 8.64E-05 | 46.2 | 84.5 | NA | 319.6 | 1031.5 | 68.5 |
| stage2 | stage3_shell | -4.24 | 2.65E-04 | V12B01_22491 | EAP94755.1 | Flp pilus assembly protein CpaB | Membrane | K02279 | U | COG3745 Flp pilus assembly protein CpaB | VV2661 | 507.5 | 1.26 | 3.17E-05 | 1.8 | 20.8 | 15.4 | 245.0 | 564.8 | 45.6 |
| stage3_free | stage3_shell | -2.65 | 1.47E-02 | V12B01_22491 | EAP94755.1 | Flp pilus assembly protein CpaB | Membrane | K02279 | U | COG3745 Flp pilus assembly protein CpaB | VV2661 | 507.5 | 1.82 | 3.21E-03 | 12.8 | 39.1 | NA | 245.0 | 564.8 | 45.6 |
| stage2 | stage3_shell | -4.53 | 5.29E-09 | V12B01_22496 | EAP94756.1 | predicted ATPase with chaperone activity | | | O | ATPase with chaperone activity | - | 1242.5 | 0.76 | 1.43E-10 | 5.6 | 18.1 | 14.6 | 288.8 | 514.4 | 12.7 |
| stage3_free | stage3_shell | -4.83 | 5.64E-08 | V12B01_22496 | EAP94756.1 | predicted ATPase with chaperone activity | Membrane | | O | ATPase with chaperone activity | - | 1242.5 | 0.88 | 1.11E-09 | 9.8 | 10.3 | NA | 288.8 | 514.4 | 12.7 |
| stage2 | stage3_shell | -3.77 | 8.35E-04 | V12B01_22501 | EAP94757.1 | hypothetical protein | | | OU | COG4960 Flp pilus assembly protein, protease CpaA | - | 709.5 | 1.26 | 1.20E-04 | 4.9 | 40.0 | 32.9 | 582.9 | 720.0 | 15.8 |

| | | | | | | | | | | | | | | | | | | | | |
|---|---|---|---|---|---|---|---|---|---|---|---|---|---|---|---|---|---|---|---|---|
| **stage3_free** | **stage3_shell** | **-5.12** | **1.24E-04** | **V12B01_22501** | **EAP94757.1** | **hypothetical protein** | | | **OU** | **COG4960 Flp pilus assembly protein, protease CpaA** | **-** | **709.5** | **1.49** | **9.70E-06** | **5.9** | **14.2** | **NA** | **582.9** | **720.0** | **15.8** |
| **stage2** | **stage3_shell** | **-3.99** | **1.83E-07** | **V12B01_22506** | **EAP94758.1** | **hypothetical protein** | | | **-** | **#N/A** | **#N/A** | **709.1** | **0.75** | **7.81E-09** | **5.6** | **82.4** | **98.1** | **723.5** | **1855.4** | **161.2** |
| **stage3_free** | **stage3_shell** | **-3.90** | **7.52E-06** | **V12B01_22506** | **EAP94758.1** | **hypothetical protein** | | | **-** | **#N/A** | **#N/A** | **709.1** | **0.88** | **3.35E-07** | **69.0** | **57.4** | **NA** | **723.5** | **1855.4** | **161.2** |
| **stage2** | **stage3_shell** | **-4.46** | **8.94E-09** | **V12B01_22511** | **EAP94759.1** | **hypothetical protein** | | **K02651** | **-** | **-** | **-** | **1380.7** | **0.76** | **2.62E-10** | **15.6** | **167.8** | **224.5** | **2616.2** | **5317.2** | **316.4** |
| **stage3_free** | **stage3_shell** | **-4.45** | **4.37E-07** | **V12B01_22511** | **EAP94759.1** | **hypothetical protein** | | **K02651** | **-** | **-** | **-** | **1380.7** | **0.88** | **1.18E-08** | **104.3** | **157.3** | **NA** | **2616.2** | **5317.2** | **316.4** |
| **stage2** | **stage3_shell** | **-3.01** | **7.75E-04** | **V12B01_22516** | **EAP94760.1** | **Response regulator** | | | **K** | **helix_turn_helix, arabinose operon control protein** | **-** | **206.4** | **0.96** | **1.09E-04** | **15.5** | **112.8** | **93.5** | **1462.5** | **455.5** | **7.7** |
| **stage3_free** | **stage3_shell** | **-4.30** | **3.72E-05** | **V12B01_22516** | **EAP94760.1** | **Response regulator** | **Regulation** | | **K** | **helix_turn_helix, arabinose operon control protein** | **-** | **206.4** | **1.10** | **2.36E-06** | **27.7** | **32.9** | **NA** | **1462.5** | **455.5** | **7.7** |
| stage2 | stage3_shell | 2.80 | 2.71E-10 | V12B01_22636 | EAP94784.1 | c-di-GMP phosphodiesterase A-related protein | Regulation | | T | signal transduction protein containing a membrane domain, an EAL and a GGDEF domain | VV2690 | 85.8 | 0.43 | 5.09E-12 | 81.3 | 79.3 | 63.0 | 6.3 | 5.9 | 11.2 |
| stage3_free | stage3_shell | 2.45 | 3.44E-07 | V12B01_22636 | EAP94784.1 | c-di-GMP phosphodiesterase A-related protein | Regulation | | T | signal transduction protein containing a membrane domain, an EAL and a GGDEF domain | VV2690 | 85.8 | 0.46 | 9.00E-09 | 64.8 | 55.6 | NA | 6.3 | 5.9 | 11.2 |
| **stage2** | **stage3_shell** | **-3.04** | **NA** | **V12B01_22641** | **EAP94785.1** | **putative protease** | | **K08303** | **O** | **COG0826 Collagenase and related proteases** | **yhbU** | **27.0** | **1.19** | **NA** | **2.2** | **6.3** | **3.7** | **12.6** | **11.9** | **81.4** |
| **stage2** | **stage3_shell** | **-2.37** | **NA** | **V12B01_22646** | **EAP94786.1** | **putative protease** | | | **O** | **COG0826 Collagenase and related proteases** | **yhbV** | **19.9** | **1.49** | **NA** | **3.0** | **4.4** | **1.4** | **12.7** | **13.8** | **38.9** |
| stage3_free | stage3_shell | 2.19 | 1.15E-05 | V12B01_22666 | EAP94790.1 | polyribonucleotide nucleotidyltransferase | | K00962 | J | Involved in mRNA degradation. Catalyzes the phosphorolysis of single-stranded polyribonucleotides processively in the 3'- to 5'-direction | pnp | 696.9 | 0.47 | 5.52E-07 | 896.4 | 765.0 | NA | 134.1 | 82.7 | 135.6 |
| stage2 | stage3_shell | 2.28 | 8.02E-03 | V12B01_22760 | EAP91762.1 | hypothetical protein | | | P | Mg Co Ni transporter MgtE (Contains CBS domain) | VP2469 | 22.5 | 0.85 | 1.79E-03 | 90.5 | 46.0 | 87.6 | 5.6 | 15.9 | 0.0 |
| stage3_free | stage3_shell | 2.59 | 4.90E-03 | V12B01_22760 | EAP91762.1 | hypothetical protein | | | P | Mg Co Ni transporter MgtE (Contains CBS domain) | VP2469 | 22.5 | 0.89 | 7.88E-04 | 118.8 | 69.1 | NA | 5.6 | 15.9 | 0.0 |
| **stage2** | **stage3_shell** | **-2.01** | **9.66E-04** | **V12B01_22770** | **EAP91764.1** | **putative integrase protein** | | **K07497** | **L** | **Evidence 2b Function of strongly homologous gene** | **-** | **21.0** | **0.62** | **1.43E-04** | **9.1** | **13.5** | **8.2** | **28.2** | **26.7** | **54.4** |
| **stage3_free** | **stage3_shell** | **-2.34** | **9.65E-04** | **V12B01_22770** | **EAP91764.1** | **putative integrase protein** | | **K07497** | **L** | **Evidence 2b Function of strongly homologous gene** | **-** | **21.0** | **0.73** | **1.10E-04** | **8.7** | **7.1** | **NA** | **28.2** | **26.7** | **54.4** |
| stage3_free | stage3_shell | 3.36 | 2.29E-12 | V12B01_22810 | EAP92452.1 | glutamate-1-semialdehyde aminotransferase | Cofactor, Carrier, and Vitamin Biosynthesis | K01845 | H | Glutamate-1-semialdehyde aminotransferase | hemL | 79.5 | 0.46 | 1.64E-14 | 253.4 | 211.3 | NA | 21.0 | 8.2 | 17.6 |
| **stage3_free** | **stage3_shell** | **-4.44** | **1.16E-02** | **V12B01_22865** | **EAP92463.1** | **Predicted N-acetylglucosamine kinase** | **Amine and Polyamine Degradation** | **K18676** | **G** | **N-acetylglucosamine kinase** | **gspK** | **3.3** | **3.10** | **2.40E-03** | **0.0** | **0.0** | **NA** | **0.0** | **6.8** | **6.4** |
| **stage2** | **stage3_shell** | **-2.25** | **1.29E-05** | **V12B01_22880** | **EAP92466.1** | **phosphoglucomutase** | **Carbohydrates and Carboxylates Degradation** | | **G** | **Phosphoglucomutase/phosphomannomutase, C-terminal domain** | **-** | **32.1** | **0.50** | **1.04E-06** | **5.3** | **3.3** | **6.2** | **15.8** | **25.7** | **16.1** |
| **stage3_free** | **stage3_shell** | **-2.10** | **3.44E-04** | **V12B01_22880** | **EAP92466.1** | **phosphoglucomutase** | **Carbohydrates and Carboxylates Degradation** | | **G** | **Phosphoglucomutase/phosphomannomutase, C-terminal domain** | **-** | **32.1** | **0.57** | **3.26E-05** | **5.2** | **5.4** | **NA** | **15.8** | **25.7** | **16.1** |
| stage3_free | stage3_shell | 2.51 | 3.40E-02 | V12B01_22910 | EAP95019.1 | hypothetical protein | | | - | - | VV10822 | 5.8 | 1.40 | 9.09E-03 | 24.1 | 17.2 | NA | 3.0 | 0.0 | 0.0 |
| stage3_free | stage3_shell | 2.17 | 7.93E-07 | V12B01_23060 | EAP95049.1 | dTDP-D-glucose 4,6-dehydratase | Carbohydrate Biosynthesis | K01710 | M | Belongs to the NAD(P)-dependent epimerase dehydratase family. dTDP-glucose dehydratase subfamily | rffG | 90.0 | 0.41 | 2.44E-08 | 164.3 | 157.5 | NA | 37.2 | 23.0 | 10.2 |
| stage2 | stage3_free | 2.80 | 2.56E-04 | V12B01_23080 | EAP95053.1 | hypothetical protein | | | - | - | - | 44.6 | 0.76 | 7.62E-06 | 36.4 | 167.3 | 167.3 | 8.3 | 20.1 | NA |
| stage2 | stage3_free | 3.25 | 5.22E-04 | V12B01_23090 | EAP95055.1 | putative integral membrane protein | Membrane | | - | - | - | 43.8 | 0.95 | 1.74E-05 | 30.4 | 143.2 | 123.4 | 4.1 | 11.2 | NA |
| stage2 | stage3_free | 2.37 | 9.79E-04 | V12B01_23095 | EAP95056.1 | putative rhamnosyl transferase | | | - | - | - | 68.9 | 0.72 | 3.93E-05 | 77.8 | 214.9 | 235.7 | 19.7 | 34.6 | NA |
| stage2 | stage3_free | 2.05 | 1.19E-03 | V12B01_23100 | EAP95057.1 | hypothetical protein | | | - | - | - | 46.9 | 0.62 | 5.06E-05 | 64.9 | 77.5 | 64.6 | 9.3 | 18.3 | NA |
| stage2 | stage3_shell | 2.12 | 7.20E-04 | V12B01_23145 | EAP95066.1 | hypothetical protein | | | S | Capsule biosynthesis GfcC | - | 18.9 | 0.60 | 9.96E-05 | 49.0 | 38.3 | 28.3 | 6.5 | 4.6 | 5.8 |
| stage3_free | stage3_shell | 2.07 | 1.77E-03 | V12B01_23145 | EAP95066.1 | hypothetical protein | | | S | Capsule biosynthesis GfcC | - | 18.9 | 0.64 | 2.28E-04 | 36.8 | 40.2 | NA | 6.5 | 4.6 | 5.8 |
| stage3_free | stage3_shell | 2.15 | 6.69E-06 | V12B01_23165 | EAP95070.1 | putative tyrosine-protein kinase Wzc | | K16692 | D | protein involved in exopolysaccharide biosynthesis | wzc | 89.0 | 0.45 | 2.92E-07 | 115.2 | 66.1 | NA | 8.1 | 19.5 | 13.1 |
| stage2 | stage3_free | 2.42 | 3.42E-03 | V12B01_23225 | EAP95082.1 | hypothetical protein | | | - | - | - | 14.9 | 0.85 | 1.78E-04 | 11.8 | 50.1 | 26.8 | 3.0 | 5.2 | NA |
| stage2 | stage3_shell | 2.83 | 4.52E-03 | V12B01_23300 | EAP95097.1 | 10 kDa chaperonin GROES | Translation | K04078 | O | Binds to Cpn60 in the presence of Mg-ATP and suppresses the ATPase activity of the latter | groS | 18.8 | 1.07 | 8.99E-04 | 331.6 | 11.0 | 153.3 | 11.0 | 20.8 | 0.0 |
| stage2 | stage3_shell | 2.96 | 8.23E-05 | V12B01_23305 | EAP95098.1 | chaperonin GroEL | Translation | K04077 | O | Prevents misfolding and promotes the refolding and proper assembly of unfolded polypeptides generated under stress conditions | groL | 211.0 | 0.77 | 8.37E-06 | 641.2 | 42.6 | 245.6 | 13.6 | 16.5 | 38.0 |
| **stage2** | **stage3_shell** | **-2.17** | **8.94E-04** | **V12B01_23325** | **EAP95102.1** | **Mg(2+) transport ATPase protein C** | **Transport Proteins** | **K07507** | **S** | **MgtC family** | **-** | **35.7** | **0.65** | **1.30E-04** | **9.1** | **11.6** | **9.6** | **39.2** | **68.1** | **11.6** |
| **stage3_free** | **stage3_shell** | **-3.69** | **3.32E-05** | **V12B01_23325** | **EAP95102.1** | **Mg(2+) transport ATPase protein C** | **Transport Proteins** | **K07507** | **S** | **MgtC family** | **-** | **35.7** | **0.87** | **2.03E-06** | **0.9** | **5.9** | **NA** | **39.2** | **68.1** | **11.6** |
| **stage2** | **stage3_free** | **-4.31** | **4.90E-03** | **V12B01_23345** | **EAP95106.1** | **fumarate reductase subunit D** | **Fermentation** | **K00247** | **C** | **Seems to be involved in the anchoring of the catalytic components of the fumarate reductase complex to the cytoplasmic membrane** | **frdD** | **207.0** | **1.57** | **2.73E-04** | **72.5** | **30.6** | **24.7** | **1632.9** | **1643.9** | **NA** |
| **stage2** | **stage3_shell** | **-3.51** | **3.65E-03** | **V12B01_23345** | **EAP95106.1** | **fumarate reductase subunit D** | **Fermentation** | **K00247** | **C** | **Seems to be involved in the anchoring of the catalytic components of the fumarate reductase complex to the cytoplasmic membrane** | **frdD** | **207.0** | **1.50** | **6.97E-04** | **72.5** | **30.6** | **24.7** | **209.1** | **82.2** | **1420.1** |

| | | | | | | | | | | | | | | | | | | | | |
|---|---|---|---|---|---|---|---|---|---|---|---|---|---|---|---|---|---|---|---|---|
| stage2 | stage3_free | -4.65 | 3.01E-03 | V12B01_23350 | EAP95107.1 | fumarate reductase subunit C | Fermentation | K00246 | C | Seems to be involved in the anchoring of the catalytic components of the fumarate reductase complex to the cytoplasmic membrane | frdC | 137.9 | 1.58 | 1.52E-04 | 24.3 | 17.5 | 23.6 | 1014.8 | 1053.3 | NA |
| stage2 | stage3_shell | -4.35 | 7.32E-04 | V12B01_23350 | EAP95107.1 | fumarate reductase subunit C | Fermentation | K00246 | C | Seems to be involved in the anchoring of the catalytic components of the fumarate reductase complex to the cytoplasmic membrane | frdC | 137.9 | 1.46 | 1.02E-04 | 24.3 | 17.5 | 23.6 | 45.8 | 197.0 | 1208.6 |
| stage2 | stage3_free | -4.07 | 1.16E-02 | V12B01_23355 | EAP95108.1 | succinate dehydrogenase | Aerobic Respiration | K00245 | C | Belongs to the succinate dehydrogenase fumarate reductase iron-sulfur protein family | frdB | 376.3 | 1.78 | 7.87E-04 | 52.3 | 23.1 | 17.1 | 1057.0 | 1251.5 | NA |
| stage2 | stage3_shell | -4.49 | 8.59E-04 | V12B01_23355 | EAP95108.1 | succinate dehydrogenase | Aerobic Respiration | K00245 | C | Belongs to the succinate dehydrogenase fumarate reductase iron-sulfur protein family | frdB | 376.3 | 1.56 | 1.24E-04 | 52.3 | 23.1 | 17.1 | 112.8 | 243.2 | 2021.0 |
| stage2 | stage3_free | -3.20 | 2.88E-02 | V12B01_23360 | EAP95109.1 | fumarate reductase | Fermentation | K00244 | C | fumarate reductase, flavoprotein subunit | frdA | 755.7 | 1.90 | 2.58E-03 | 54.6 | 24.3 | 24.0 | 640.9 | 944.4 | NA |
| stage2 | stage3_shell | -4.09 | 1.46E-03 | V12B01_23360 | EAP95109.1 | fumarate reductase | Fermentation | K00244 | C | fumarate reductase, flavoprotein subunit | frdA | 755.7 | 1.52 | 2.36E-04 | 54.6 | 24.3 | 24.0 | 147.2 | 276.2 | 1601.6 |
| stage2 | stage3_shell | -2.40 | 5.80E-03 | V12B01_23380 | EAP95113.1 | putative transporter | | K14445 | P | COG0471 Di- and tricarboxylate transporters | - | 26.6 | 1.00 | 1.22E-03 | 7.9 | 6.0 | 4.2 | 12.4 | 23.5 | 60.3 |
| stage3_free | stage3_shell | 2.07 | 5.85E-04 | V12B01_23405 | EAP95118.1 | diaminobutyrate-pyruvate transaminase & L-2,4-diaminobutyratedecarboxylase | | K00836 | E | COG0160 4-aminobutyrate aminotransferase and related aminotransferases | - | 137.0 | 0.58 | 6.05E-05 | 136.8 | 112.8 | NA | 9.2 | 12.8 | 31.8 |
| stage2 | stage3_free | -3.26 | 6.97E-05 | V12B01_23435 | EAP95124.1 | putative anaerobic dimethyl sulfoxide reductase subunit C | Respiration | K07308 | S | DMSO reductase anchor subunit (DmsC) | - | 54.9 | 0.77 | 1.58E-06 | 10.0 | 17.2 | 10.3 | 184.9 | 118.1 | NA |
| stage2 | stage3_shell | -3.01 | 1.90E-05 | V12B01_23435 | EAP95124.1 | putative anaerobic dimethyl sulfoxide reductase subunit C | Respiration | K07308 | S | DMSO reductase anchor subunit (DmsC) | - | 54.9 | 0.71 | 1.61E-06 | 10.0 | 17.2 | 10.3 | 22.7 | 69.7 | 158.2 |
| stage2 | stage3_free | -4.94 | 9.24E-04 | V12B01_23440 | EAP95125.1 | anaerobic dimethyl sulfoxide reductase chain B | Respiration | K07307 | C | 4Fe-4S dicluster domain | - | 22.8 | 1.44 | 3.62E-05 | 1.4 | 1.1 | 4.0 | 111.2 | 123.0 | NA |
| stage2 | stage3_shell | -4.41 | 4.10E-04 | V12B01_23440 | EAP95125.1 | anaerobic dimethyl sulfoxide reductase chain B | Respiration | K07307 | C | 4Fe-4S dicluster domain | - | 22.8 | 1.38 | 5.21E-05 | 1.4 | 1.1 | 4.0 | 5.1 | 12.2 | 133.0 |
| stage2 | stage3_shell | -4.88 | NA | V12B01_23445 | EAP95126.1 | putative anaerobic dimethyl sulfoxide reductase, subunit A | Respiration | K07306 | C | Molybdopterin oxidoreductase Fe4S4 domain | dmsA | 48.6 | 1.98 | NA | 0.5 | 0.9 | 0.3 | 3.9 | 1.2 | 63.0 |
| stage2 | stage3_shell | -5.30 | 2.21E-06 | V12B01_23455 | EAP95128.1 | hypothetical ferredoxin-type protein NapF | Respiration | | C | 4Fe-4S binding domain | - | 17.5 | 1.12 | 1.33E-07 | 0.0 | 1.4 | 5.5 | 37.4 | 59.0 | 161.0 |
| stage3_free | stage3_shell | -2.24 | 2.02E-02 | V12B01_23455 | EAP95128.1 | hypothetical ferredoxin-type protein NapF | | | C | 4Fe-4S binding domain | - | 17.5 | 1.41 | 4.77E-03 | 4.5 | 23.4 | NA | 37.4 | 59.0 | 161.0 |
| stage2 | stage3_shell | -2.67 | 5.20E-03 | V12B01_23499 | EAP93130.1 | putative YfrE protein | | K04018 | O | COG4235 Cytochrome c biogenesis factor | nrfG | 16.1 | 1.13 | 1.07E-03 | 7.1 | 7.3 | 6.4 | 10.1 | 33.6 | 99.4 |
| stage2 | stage3_free | -3.72 | 6.71E-08 | V12B01_23509 | EAP93132.1 | nitrite reductase periplasmic cytochrome c552 | Respiration | K03385 | C | Catalyzes the reduction of nitrite to ammonia, consuming six electrons in the process | nrfA | 297.2 | 0.64 | 6.51E-10 | 7.5 | 26.0 | 13.7 | 238.7 | 248.1 | NA |
| stage2 | stage3_shell | -4.50 | 5.65E-14 | V12B01_23509 | EAP93132.1 | nitrite reductase periplasmic cytochrome c552 | Respiration | K03385 | C | Catalyzes the reduction of nitrite to ammonia, consuming six electrons in the process | nrfA | 297.2 | 0.58 | 5.48E-16 | 7.5 | 26.0 | 13.7 | 125.1 | 572.0 | 189.0 |
| stage2 | stage3_free | -3.26 | 3.90E-05 | V12B01_23514 | EAP93133.1 | cytochrome c-type protein NrfB precursor | Respiration | K04013 | C | Cytochrome c554 and c-prime | nrfB | 151.2 | 0.74 | 7.73E-07 | 6.8 | 25.4 | 20.3 | 158.7 | 261.1 | NA |
| stage2 | stage3_shell | -4.93 | 4.47E-13 | V12B01_23514 | EAP93133.1 | cytochrome c-type protein NrfB precursor | Respiration | K04013 | C | Cytochrome c554 and c-prime | nrfB | 151.2 | 0.66 | 4.90E-15 | 6.8 | 25.4 | 20.3 | 103.8 | 871.5 | 355.3 |
| stage2 | stage3_free | -3.98 | 7.75E-06 | V12B01_23519 | EAP93134.1 | Fe-S-cluster-containing hydrogenase component 1 | Respiration | K04014 | C | COG0437 Fe-S-cluster-containing hydrogenase components 1 | nrfC | 84.5 | 0.83 | 1.19E-07 | 7.1 | 16.0 | 5.9 | 190.4 | 197.2 | NA |
| stage2 | stage3_shell | -4.38 | 1.39E-08 | V12B01_23519 | EAP93134.1 | Fe-S-cluster-containing hydrogenase component 1 | | K04014 | C | COG0437 Fe-S-cluster-containing hydrogenase components 1 | nrfC | 84.5 | 0.76 | 4.21E-10 | 7.1 | 16.0 | 5.9 | 48.8 | 310.5 | 161.6 |
| stage2 | stage3_free | -4.83 | 4.52E-09 | V12B01_23524 | EAP93135.1 | NrfD protein | Respiration | K04015 | P | Formate-dependent nitrite reductase, membrane component | nrfD | 124.8 | 0.77 | 3.39E-11 | 6.4 | 5.1 | 6.7 | 183.4 | 226.6 | NA |
| stage2 | stage3_shell | -5.18 | 8.94E-13 | V12B01_23524 | EAP93135.1 | NrfD protein | Respiration | K04015 | P | Formate-dependent nitrite reductase, membrane component | nrfD | 124.8 | 0.71 | 1.08E-14 | 6.4 | 5.1 | 6.7 | 98.3 | 187.8 | 242.4 |
| stage2 | stage3_free | -3.22 | 1.08E-05 | V12B01_23529 | EAP93136.1 | heme chaperone--apocytochrome heme-lyase | | K02198 | O | COG1138 Cytochrome c biogenesis factor | nrfE | 62.0 | 0.68 | 1.76E-07 | 2.5 | 6.6 | 2.4 | 46.1 | 41.7 | NA |
| stage2 | stage3_shell | -3.80 | 3.25E-09 | V12B01_23529 | EAP93136.1 | heme chaperone--apocytochrome heme-lyase | | K02198 | O | COG1138 Cytochrome c biogenesis factor | nrfE | 62.0 | 0.63 | 8.09E-11 | 2.5 | 6.6 | 2.4 | 34.9 | 26.0 | 68.1 |
| stage2 | stage3_free | -2.14 | 4.63E-03 | V12B01_23534 | EAP93137.1 | thiol:disulfide interchange protein DsbE | | K02199 | CO | Thiol disulfide interchange protein dsbE | - | 15.4 | 0.73 | 2.57E-04 | 8.0 | 8.9 | 10.7 | 51.9 | 56.3 | NA |
| stage2 | stage3_shell | -3.07 | 2.35E-09 | V12B01_23574 | EAP93145.1 | Uncharacterized protein conserved in bacteria | | K21470 | S | protein conserved in bacteria | VP1916 | 533.9 | 0.50 | 5.55E-11 | 22.5 | 85.6 | 68.7 | 580.3 | 530.8 | 140.6 |
| stage3_free | stage3_shell | -4.03 | 5.98E-12 | V12B01_23574 | EAP93145.1 | Uncharacterized protein conserved in bacteria | | K21470 | S | protein conserved in bacteria | VP1916 | 533.9 | 0.57 | 4.67E-14 | 21.4 | 39.6 | NA | 580.3 | 530.8 | 140.6 |
| stage2 | stage3_shell | -3.31 | 1.35E-09 | V12B01_23589 | EAP93148.1 | hypothetical protein | | | S | Protein of unknown function (DUF2982) | VV12961 | 82.9 | 0.52 | 2.98E-11 | 7.6 | 11.3 | 9.2 | 43.1 | 150.2 | 34.9 |
| stage3_free | stage3_shell | -2.97 | 1.64E-06 | V12B01_23589 | EAP93148.1 | hypothetical protein | | | S | Protein of unknown function (DUF2982) | VV12961 | 82.9 | 0.60 | 5.60E-08 | 11.2 | 11.8 | NA | 43.1 | 150.2 | 34.9 |
| stage3_free | stage3_shell | 2.18 | 1.65E-03 | V12B01_23634 | EAP93157.1 | methyl-accepting chemotaxis protein | | K03406 | NT | methyl-accepting chemotaxis protein | - | 128.8 | 0.68 | 2.09E-04 | 196.1 | 277.4 | NA | 61.1 | 4.0 | 27.2 |
| stage3_free | stage3_shell | 2.22 | 2.63E-04 | V12B01_23654 | EAP93161.1 | thioredoxin reductase | Cofactor, Carrier, and Vitamin Biosynthesis | K00384 | C | Belongs to the class-II pyridine nucleotide-disulfide oxidoreductase family | trxB | 94.5 | 0.59 | 2.36E-05 | 120.1 | 129.1 | NA | 23.3 | 9.5 | 17.8 |
| stage2 | stage3_shell | -3.63 | 2.17E-06 | V12B01_23679 | EAP93166.1 | hypothetical protein | | | S | protein conserved in bacteria | - | 837.9 | 0.76 | 1.29E-07 | 12.2 | 32.9 | 39.4 | 328.3 | 635.5 | 17.3 |
| stage2 | stage3_shell | -3.65 | 4.73E-07 | V12B01_23684 | EAP93167.1 | NifS-related protein | Amino Acid Biosynthesis | | E | COG0520 Selenocysteine lyase | - | 913.2 | 0.71 | 2.27E-08 | 11.5 | 34.7 | 34.6 | 246.9 | 647.5 | 41.5 |

| | | | | | | | | | | | | | | | | | | | | |
|---|---|---|---|---|---|---|---|---|---|---|---|---|---|---|---|---|---|---|---|---|
| stage3_free | stage3_shell | -2.47 | 1.70E-03 | V12B01_23684 | EAP93167.1 | NifS-related protein | Amino Acid Biosynthesis | | E | COG0520 Selenocysteine lyase | - | 913.2 | 0.84 | 2.17E-04 | 64.0 | 46.2 | NA | 246.9 | 647.5 | 41.5 |
| stage2 | stage3_free | -8.87 | 5.50E-16 | V12B01_23714 | EAP93390.1 | transporter, NadC family | Transport Proteins | K14445 | P | COG0471 Di- and tricarboxylate transporters | - | 146.9 | 1.02 | 4.85E-19 | 1.3 | 1.8 | 0.7 | 765.9 | 671.7 | NA |
| stage2 | stage3_shell | -5.37 | 4.56E-08 | V12B01_23714 | EAP93390.1 | transporter, NadC family | Transport Proteins | K14445 | P | COG0471 Di- and tricarboxylate transporters | - | 146.9 | 0.97 | 1.61E-09 | 1.3 | 1.8 | 0.7 | 83.9 | 4.4 | 51.9 |
| stage3_free | stage3_shell | 2.96 | 2.45E-03 | V12B01_23714 | EAP93390.1 | transporter, NadC family | | K14445 | P | COG0471 Di- and tricarboxylate transporters | - | 146.9 | 1.00 | 3.44E-04 | 765.9 | 671.7 | NA | 83.9 | 4.4 | 51.9 |
| stage2 | stage3_free | -3.50 | 2.50E-06 | V12B01_23734 | EAP93394.1 | hypothetical protein | | | - | - | - | 64.5 | 0.69 | 3.47E-08 | 28.2 | 59.0 | 49.5 | 655.8 | 592.2 | NA |
| stage3_free | stage3_shell | 3.05 | 4.98E-07 | V12B01_23739 | EAP93395.1 | phosphoribosylaminoimidazole-succinocarboxamide synthase | Nucleoside and Nucleotide Biosynthesis | K01923 | F | Belongs to the SAICAR synthetase family | purC | 108.6 | 0.58 | 1.40E-08 | 360.0 | 368.1 | NA | 46.3 | 12.3 | 25.8 |
| stage2 | stage3_shell | -3.04 | 2.21E-04 | V12B01_23819 | EAP93411.1 | hypothetical protein | | | S | Protein of unknown function (DUF3187) | - | 9.3 | 0.83 | 2.58E-05 | 1.9 | 16.4 | 4.0 | 47.2 | 102.1 | 24.0 |
| stage3_free | stage3_shell | -4.20 | 3.35E-04 | V12B01_23819 | EAP93411.1 | hypothetical protein | | | S | Protein of unknown function (DUF3187) | - | 9.3 | 1.16 | 3.15E-05 | 0.0 | 6.1 | NA | 47.2 | 102.1 | 24.0 |
| stage3_free | stage3_shell | -3.15 | 1.40E-03 | V12B01_23824 | EAP93412.1 | hypothetical protein | | | S | Protein of unknown function (DUF3187) | - | 12.2 | 1.00 | 1.72E-04 | 0.0 | 4.9 | NA | 26.0 | 36.9 | 0.0 |
| stage2 | stage3_free | 2.20 | 7.90E-05 | V12B01_23844 | EAP93416.1 | hypothetical protein | | | L | Protein of unknown function (DUF3732) | - | 223.2 | 0.53 | 1.86E-06 | 168.7 | 216.5 | 343.6 | 41.5 | 50.7 | NA |
| stage2 | stage3_shell | -3.86 | 9.70E-03 | V12B01_23899 | EAP93427.1 | hypothetical protein | | | - | #N/A | #N/A | 2.3 | 2.11 | 2.24E-03 | 2.5 | 0.0 | 0.0 | 53.3 | 0.0 | 15.8 |
| stage3_free | stage3_shell | -2.24 | 6.12E-03 | V12B01_24034 | EAP93454.1 | Phage tail sheath protein FI-like | | K06907 | S | tail sheath protein | - | 9.5 | 0.88 | 1.04E-03 | 2.0 | 2.1 | NA | 4.1 | 12.9 | 12.2 |
| stage2 | stage3_free | -5.47 | 1.26E-07 | V12B01_24084 | EAP93464.1 | hypothetical protein | | | - | - | - | 5388.5 | 0.96 | 1.27E-09 | 50.9 | 301.3 | 229.8 | 7608.9 | 14324.8 | NA |
| stage2 | stage3_shell | -9.15 | 1.19E-24 | V12B01_24084 | EAP93464.1 | hypothetical protein | | | - | #N/A | #N/A | 5388.5 | 0.86 | 1.25E-27 | 50.9 | 301.3 | 229.8 | 38783.3 | 29038.2 | 177278.3 |
| stage3_free | stage3_shell | -2.93 | 1.73E-03 | V12B01_24084 | EAP93464.1 | hypothetical protein | | | - | #N/A | #N/A | 5388.5 | 1.08 | 2.21E-04 | 7608.9 | 14324.8 | NA | 38783.3 | 29038.2 | 177278.3 |
| stage2 | stage3_shell | -3.28 | 2.46E-10 | V12B01_24089 | EAP93465.1 | hypothetical protein | | | - | #N/A | #N/A | 541.7 | 0.50 | 4.51E-12 | 200.3 | 139.7 | 149.4 | 792.1 | 580.9 | 2147.4 |
| stage3_free | stage3_shell | -2.12 | 2.48E-04 | V12B01_24089 | EAP93465.1 | hypothetical protein | | | - | #N/A | #N/A | 541.7 | 0.57 | 2.21E-05 | 321.8 | 374.0 | NA | 792.1 | 580.9 | 2147.4 |
| stage2 | stage3_shell | 2.05 | 2.08E-07 | V12B01_24139 | EAP93047.1 | 50S ribosomal protein L20 | Translation | K02887 | J | Binds directly to 23S ribosomal RNA and is necessary for the in vitro assembly process of the 50S ribosomal subunit. It is not involved in the protein synthesizing functions of that subunit | rplT | 1204.9 | 0.38 | 9.15E-09 | 4165.3 | 7703.9 | 7350.9 | 1182.5 | 1162.4 | 836.5 |
| stage3_free | stage3_shell | 3.25 | 1.50E-03 | V12B01_24169 | EAP93053.1 | coniferyl aldehyde dehydrogenase | | K00154 | C | Belongs to the aldehyde dehydrogenase family | calB | 19.3 | 0.99 | 1.85E-04 | 29.0 | 30.1 | NA | 0.0 | 4.2 | 0.0 |
| stage2 | stage3_shell | 3.36 | 2.51E-03 | V12B01_24179 | EAP93055.1 | agmatinase | Amino Acid Degradation | K01480 | E | Belongs to the arginase family | speB | 55.5 | 1.21 | 4.47E-04 | 7.7 | 36.2 | 58.0 | 0.0 | 0.0 | 6.1 |
| stage2 | stage3_shell | -2.52 | 2.37E-15 | V12B01_24184 | EAP93056.1 | hypothetical protein | | | Q | COG2931, RTX toxins and related Ca2 -binding proteins | - | 8453.3 | 0.31 | 1.65E-17 | 91.1 | 125.9 | 128.9 | 307.6 | 567.3 | 593.8 |
| stage3_free | stage3_shell | -2.71 | 5.64E-14 | V12B01_24184 | EAP93056.1 | hypothetical protein | | | Q | COG2931, RTX toxins and related Ca2 -binding proteins | - | 8453.3 | 0.34 | 2.26E-16 | 93.0 | 109.3 | NA | 307.6 | 567.3 | 593.8 |
| stage2 | stage3_shell | 2.09 | 2.78E-04 | V12B01_24189 | EAP93057.1 | cyclohexadienyl dehydratase precursor | | K01713 | ET | belongs to the bacterial solute-binding protein 3 family | - | 34.1 | 0.56 | 3.35E-05 | 73.8 | 68.1 | 95.0 | 16.7 | 7.9 | 11.2 |
| stage2 | stage3_shell | 2.27 | 6.57E-07 | V12B01_24194 | EAP93058.1 | putative transporter | Transport Proteins | K03307 | E | Belongs to the sodium solute symporter (SSF) (TC 2.A.21) family | toaA | 209.7 | 0.44 | 3.32E-08 | 436.1 | 347.6 | 219.0 | 28.0 | 32.5 | 75.7 |
| stage2 | stage3_shell | 3.12 | 1.46E-11 | V12B01_24199 | EAP93059.1 | keto-hydroxyglutarate-aldolase/keto-deoxy-phosphogluconate aldolase | | K01625 | G | COG0800 2-keto-3-deoxy-6-phosphogluconate aldolase | eda | 360.1 | 0.44 | 2.22E-13 | 2083.6 | 2576.4 | 1455.8 | 135.1 | 241.3 | 109.0 |
| stage2 | stage3_shell | 3.17 | 4.49E-13 | V12B01_24204 | EAP93060.1 | putative 2-dehydro-3-deoxygluconokinase | Carbohydrates and Carboxylates Degradation | K00874 | G | COG0524 Sugar kinases, ribokinase family | kdgK | 291.9 | 0.42 | 5.02E-15 | 1517.0 | 925.1 | 608.1 | 49.7 | 113.5 | 70.2 |
| stage3_free | stage3_shell | 2.08 | 2.60E-05 | V12B01_24204 | EAP93060.1 | putative 2-dehydro-3-deoxygluconokinase | Carbohydrates and Carboxylates Degradation | K00874 | G | COG0524 Sugar kinases, ribokinase family | kdgK | 291.9 | 0.47 | 1.47E-06 | 576.0 | 426.8 | NA | 49.7 | 113.5 | 70.2 |
| stage2 | stage3_shell | 3.07 | 1.08E-08 | V12B01_24209 | EAP93061.1 | pectin degradation protein | Carbohydrate degradation | | S | Cupin domain | - | 129.9 | 0.52 | 3.19E-10 | 1745.6 | 1253.3 | 883.2 | 85.6 | 103.6 | 118.6 |
| stage2 | stage3_shell | 2.54 | 5.84E-07 | V12B01_24214 | EAP93062.1 | putative chondroitin AC/alginate lyase | Carbohydrates and Carboxylates Degradation | K20525 | S | Alginate lyase | oalB | 347.8 | 0.49 | 2.91E-08 | 754.8 | 431.8 | 270.2 | 23.2 | 71.3 | 68.3 |
| stage2 | stage3_shell | -2.60 | 5.49E-06 | V12B01_24224 | EAP93064.1 | hypothetical protein | | | Q | Domain of unknown function (DUF2437) | - | 108.1 | 0.56 | 3.62E-07 | 14.6 | 47.7 | 48.8 | 300.2 | 249.4 | 38.0 |
| stage3_free | stage3_shell | -2.85 | 1.31E-05 | V12B01_24224 | EAP93064.1 | hypothetical protein | | | Q | Domain of unknown function (DUF2437) | - | 108.1 | 0.64 | 6.46E-07 | 32.7 | 28.5 | NA | 300.2 | 249.4 | 38.0 |
| stage2 | stage3_shell | 2.89 | 6.91E-07 | V12B01_24234 | EAP93066.1 | putative transporter | Transport Proteins | K03307 | E | Belongs to the sodium solute symporter (SSF) (TC 2.A.21) family | toaB | 792.1 | 0.57 | 3.54E-08 | 2277.1 | 1308.0 | 663.9 | 43.1 | 176.0 | 144.0 |
| stage3_free | stage3_shell | 2.12 | 1.11E-03 | V12B01_24234 | EAP93066.1 | putative transporter | Transport Proteins | K03307 | E | Belongs to the sodium solute symporter (SSF) (TC 2.A.21) family | toaB | 792.1 | 0.64 | 1.30E-04 | 1162.0 | 617.8 | NA | 43.1 | 176.0 | 144.0 |
| stage2 | stage3_shell | 2.83 | 8.67E-04 | V12B01_24244 | EAP94919.1 | putative tartronate semialdehyde reductase (TSAR) | Carbohydrates and Carboxylates Degradation | | I | NAD-binding of NADP-dependent 3-hydroxyisobutyrate dehydrogenase | dehR | 350.3 | 0.90 | 1.25E-04 | 2361.5 | 884.6 | 482.9 | 7.2 | 63.5 | 193.7 |
| stage3_free | stage3_shell | 2.02 | 2.21E-02 | V12B01_24244 | EAP94919.1 | putative tartronate semialdehyde reductase (TSAR) | Carbohydrates and Carboxylates Degradation | | I | NAD-binding of NADP-dependent 3-hydroxyisobutyrate dehydrogenase | dehR | 350.3 | 0.99 | 5.36E-03 | 1043.3 | 605.1 | NA | 7.2 | 63.5 | 193.7 |
| stage2 | stage3_shell | 2.53 | 4.55E-05 | V12B01_24249 | EAP94920.1 | methyl-accepting chemotaxis protein | Chemotaxis | K03406 | NT | methyl-accepting chemotaxis protein | - | 110.2 | 0.62 | 4.21E-06 | 269.1 | 193.7 | 136.0 | 6.0 | 21.0 | 36.5 |
| stage2 | stage3_shell | 3.92 | 3.88E-10 | V12B01_24254 | EAP94921.1 | lyase, putative | Carbohydrate degradation | | G | Alginate lyase | alyA | 119.5 | 0.61 | 7.62E-12 | 261.7 | 268.0 | 246.0 | 3.7 | 13.9 | 16.3 |
| stage3_free | stage3_shell | 2.26 | 8.86E-04 | V12B01_24254 | EAP94921.1 | lyase, putative | Carbohydrates and Carboxylates Degradation | | G | Alginate lyase | alyA | 119.5 | 0.67 | 9.93E-05 | 105.5 | 75.2 | NA | 3.7 | 13.9 | 16.3 |
| stage2 | stage3_shell | 3.67 | 1.35E-07 | V12B01_24259 | EAP94922.1 | putative alginate lyase | Carbohydrates and Carboxylates Degradation | | G | F5/8 type C domain | alyB | 544.9 | 0.68 | 5.56E-09 | 2086.5 | 858.9 | 633.1 | 12.2 | 55.0 | 101.6 |

| | | | | | | | | | | | | | | | | | | | | |
|---|---|---|---|---|---|---|---|---|---|---|---|---|---|---|---|---|---|---|---|---|
| stage3_free | stage3_shell | 2.57 | 8.05E-04 | V12B01_24259 | EAP94922.1 | putative alginate lyase | Carbohydrates and Carboxylates Degradation | | G | F5/8 type C domain | alyB | 544.9 | 0.76 | 8.83E-05 | 744.2 | 491.4 | NA | 12.2 | 55.0 | 101.6 |
| stage2 | stage3_shell | 3.32 | 1.31E-06 | V12B01_24264 | EAP94923.1 | hypothetical protein | | | - | #N/A | alyC | 1041.5 | 0.68 | 7.29E-08 | 4585.1 | 2498.0 | 1956.8 | 28.6 | 157.4 | 338.0 |
| stage3_free | stage3_shell | 2.21 | 3.52E-03 | V12B01_24264 | EAP94923.1 | hypothetical protein | | | - | #N/A | alyC | 1041.5 | 0.75 | 5.29E-04 | 1960.3 | 1100.1 | NA | 28.6 | 157.4 | 338.0 |
| stage2 | stage3_shell | 3.30 | 1.56E-06 | V12B01_24269 | EAP94924.1 | putative oligogalacturonate specific porin | Transport Proteins | K22110 | M | Oligogalacturonate-specific porin protein (KdgM) | kdgM | 837.2 | 0.68 | 8.84E-08 | 5444.5 | 2754.0 | 2132.6 | 89.1 | 196.8 | 340.6 |
| stage3_free | stage3_shell | 2.14 | 4.34E-03 | V12B01_24269 | EAP94924.1 | putative oligogalacturonate specific porin | | K22110 | M | Oligogalacturonate-specific porin protein (KdgM) | kdgM | 837.2 | 0.75 | 6.82E-04 | 2098.2 | 1337.9 | NA | 89.1 | 196.8 | 340.6 |
| stage2 | stage3_shell | 3.56 | 8.79E-11 | V12B01_24274 | EAP94925.1 | putative alginate lyase | Carbohydrates and Carboxylates Degradation | | M | Alginate lyase | alyD | 369.4 | 0.53 | 1.45E-12 | 1423.1 | 1536.8 | 977.4 | 21.5 | 88.9 | 107.0 |
| stage3_free | stage3_shell | 2.01 | 9.30E-04 | V12B01_24274 | EAP94925.1 | putative alginate lyase | Carbohydrates and Carboxylates Degradation | | M | Alginate lyase | alyD | 369.4 | 0.59 | 1.06E-04 | 581.6 | 394.7 | NA | 21.5 | 88.9 | 107.0 |
| stage2 | stage3_shell | 2.92 | 1.25E-05 | V12B01_24284 | EAP94927.1 | GntR-famly transcriptional regulator | Regulation | | K | Evidence 3 Function proposed based on presence of conserved amino acid motif, structural feature or limited homology | - | 20.6 | 0.64 | 9.85E-07 | 93.5 | 86.7 | 61.3 | 4.4 | 8.4 | 7.9 |
| stage3_free | stage3_shell | 2.40 | 6.99E-04 | V12B01_24284 | EAP94927.1 | GntR-famly transcriptional regulator | Regulation | | K | Evidence 3 Function proposed based on presence of conserved amino acid motif, structural feature or limited homology | - | 20.6 | 0.68 | 7.48E-05 | 57.6 | 59.8 | NA | 4.4 | 8.4 | 7.9 |
| stage2 | stage3_shell | 2.43 | 2.14E-08 | V12B01_24289 | EAP94928.1 | 3-ketoacyl-CoA reductase PhaB | | K00059 | IQ | COG1028 Dehydrogenases with different specificities (related to short-chain alcohol dehydrogenases) | - | 125.3 | 0.42 | 6.91E-10 | 364.5 | 599.0 | 368.3 | 35.0 | 85.7 | 47.6 |
| stage2 | stage3_shell | 3.81 | 8.23E-10 | V12B01_24309 | EAP94932.1 | hypothetical protein | | K22110 | M | Oligogalacturonate-specific porin protein (KdgM) | kdgN | 69.2 | 0.61 | 1.75E-11 | 211.9 | 534.7 | 330.1 | 13.2 | 8.3 | 31.4 |
| stage2 | stage3_shell | 3.26 | 3.76E-05 | V12B01_24324 | EAP94935.1 | putative transporter | Transport Proteins | K03307 | E | Belongs to the sodium solute symporter (SSF) (TC 2.A.21) family | toaC | 139.4 | 0.80 | 3.42E-06 | 444.9 | 242.3 | 148.4 | 3.9 | 21.0 | 25.7 |
| stage3_free | stage3_shell | 2.30 | 6.65E-03 | V12B01_24324 | EAP94935.1 | putative transporter | Transport Proteins | K03307 | E | Belongs to the sodium solute symporter (SSF) (TC 2.A.21) family | toaC | 139.4 | 0.89 | 1.16E-03 | 221.4 | 103.7 | NA | 3.9 | 21.0 | 25.7 |
| **stage2** | **stage3_free** | **-3.90** | **5.28E-10** | **V12B01_24344** | **EAP94939.1** | **putative tryptophanase** | **Amino Acid Degradation** | **K01667** | **H** | **Belongs to the beta-eliminating lyase family** | **tnaA** | **449.9** | **0.59** | **2.79E-12** | **6.7** | **6.9** | **13.1** | **61.0** | **240.5** | **NA** |
| **stage2** | **stage3_shell** | **-3.86** | **4.23E-12** | **V12B01_24344** | **EAP94939.1** | **putative tryptophanase** | | **K01667** | **H** | **Belongs to the beta-eliminating lyase family** | **tnaA** | **449.9** | **0.54** | **5.62E-14** | **6.7** | **6.9** | **13.1** | **178.6** | **85.0** | **54.0** |
| **stage2** | **stage3_free** | **-2.89** | **1.38E-08** | **V12B01_24369** | **EAP94944.1** | **Zinc metalloprotease** | | **K08604** | **E** | **COG3227 Zinc metalloprotease (elastase)** | **hap** | **96.8** | **0.48** | **1.18E-10** | **8.3** | **5.8** | **3.9** | **35.6** | **63.2** | **NA** |
| **stage2** | **stage3_shell** | **-4.59** | **1.55E-24** | **V12B01_24369** | **EAP94944.1** | **Zinc metalloprotease** | | **K08604** | **E** | **COG3227 Zinc metalloprotease (elastase)** | **hap** | **96.8** | **0.43** | **1.97E-27** | **8.3** | **5.8** | **3.9** | **102.5** | **148.5** | **84.3** |
| stage2 | stage3_free | 2.34 | 1.77E-02 | V12B01_24454 | EAP94961.1 | 5-methyltetrahydropteroyltriglutamate--homocysteine methyltransferase | Amino Acid Biosynthesis | K00549 | E | COG0620 Methionine synthase II (cobalamin-independent) | metE | 15.2 | 1.21 | 1.37E-03 | 42.8 | 7.9 | 103.1 | 1.8 | 10.3 | NA |
| stage2 | stage3_shell | 2.95 | 3.65E-03 | V12B01_24454 | EAP94961.1 | 5-methyltetrahydropteroyltriglutamate--homocysteine methyltransferase | Amino Acid Biosynthesis | K00549 | E | COG0620 Methionine synthase II (cobalamin-independent) | metE | 15.2 | 1.08 | 6.97E-04 | 42.8 | 7.9 | 103.1 | 6.2 | 2.9 | 0.0 |
| stage2 | stage3_free | 3.17 | 1.52E-02 | V12B01_24459 | EAP94962.1 | hypothetical protein | | | S | Domain of unknown function (DUF1852) | - | 7.5 | 1.63 | 1.12E-03 | 40.0 | 5.4 | 67.3 | 1.9 | 2.5 | NA |
| stage2 | stage3_shell | 6.34 | 5.59E-04 | V12B01_24459 | EAP94962.1 | hypothetical protein | | | S | Domain of unknown function (DUF1852) | - | 7.5 | 2.75 | 7.44E-05 | 40.0 | 5.4 | 67.3 | 0.0 | 0.0 | 0.0 |
| **stage2** | **stage3_free** | **-3.62** | **1.49E-05** | **V12B01_24499** | **EAP94970.1** | **Na+/alanine symporter** | **Transport Proteins** | **K03310** | **U** | **COG1115 Na alanine symporter** | **-** | **39.8** | **0.78** | **2.55E-07** | **12.6** | **7.5** | **1.6** | **133.1** | **89.7** | **NA** |
| stage3_free | stage3_shell | 2.13 | 3.31E-03 | V12B01_24539 | EAP94978.1 | hypothetical protein | | K01993 | V | COG0845 Membrane-fusion protein | - | 20.8 | 0.70 | 4.90E-04 | 50.2 | 45.3 | NA | 6.6 | 10.9 | 0.0 |
| **stage3_free** | **stage3_shell** | **-2.31** | **5.75E-03** | **V12B01_24554** | **EAP94981.1** | **ISPsy11, transposase OrfA** | | **K07483** | **L** | **Evidence 2b Function of strongly homologous gene** | **-** | **14.2** | **0.91** | **9.59E-04** | **11.4** | **9.6** | **NA** | **99.4** | **45.2** | **14.2** |
| **stage2** | **stage3_free** | **-2.39** | **1.60E-03** | **V12B01_24559** | **EAP94982.1** | **Cation transport ATPase** | **Transport Proteins** | | **-** | **-** | **-** | **131.8** | **0.72** | **7.32E-05** | **19.5** | **69.9** | **60.2** | **316.9** | **362.4** | **NA** |
| **stage2** | **stage3_shell** | **-5.09** | **5.49E-15** | **V12B01_24559** | **EAP94982.1** | **Cation transport ATPase** | **Transport Proteins** | | **-** | **-** | **-** | **131.8** | **0.63** | **4.06E-17** | **19.5** | **69.9** | **60.2** | **441.5** | **703.4** | **2634.9** |
| **stage3_free** | **stage3_shell** | **-2.16** | **1.88E-03** | **V12B01_24559** | **EAP94982.1** | **Cation transport ATPase** | **Transport Proteins** | | **-** | **-** | **-** | **131.8** | **0.73** | **2.46E-04** | **316.9** | **362.4** | **NA** | **441.5** | **703.4** | **2634.9** |
| **stage2** | **stage3_free** | **-2.01** | **4.82E-05** | **V12B01_24589** | **EAP94988.1** | **Response regulator** | **Regulation** | | **KT** | **HD domain** | **-** | **49.7** | **0.46** | **1.02E-06** | **16.3** | **11.1** | **16.8** | **76.8** | **58.5** | **NA** |
| stage2 | stage3_shell | 3.72 | 9.83E-03 | V12B01_24604 | EAP94991.1 | hypothetical protein | | K07090 | S | membrane transporter protein | - | 4.8 | 2.29 | 2.28E-03 | 19.9 | 8.4 | 7.8 | 0.0 | 0.0 | 0.0 |
| stage3_free | stage3_shell | 3.97 | 9.89E-03 | V12B01_24604 | EAP94991.1 | hypothetical protein | | K07090 | S | membrane transporter protein | - | 4.8 | 2.37 | 1.91E-03 | 12.1 | 16.5 | NA | 0.0 | 0.0 | 0.0 |
| **stage2** | **stage3_free** | **-2.54** | **NA** | **V12B01_24644** | **EAP94999.1** | **Transcriptional regulator** | **Regulation** | | **K** | **COG1802 Transcriptional regulators** | **-** | **55.1** | **0.93** | **NA** | **9.9** | **15.2** | **23.4** | **142.3** | **133.4** | **NA** |
| **stage2** | **stage3_shell** | **-2.56** | **NA** | **V12B01_24644** | **EAP94999.1** | **Transcriptional regulator** | **Regulation** | | **K** | **COG1802 Transcriptional regulators** | **-** | **55.1** | **0.83** | **NA** | **9.9** | **15.2** | **23.4** | **255.2** | **12.6** | **15.8** |
| **stage2** | **stage3_free** | **-3.14** | **4.37E-07** | **V12B01_24649** | **EAP95000.1** | **carboxyphosphonoenolpyruvate phosphonomutase** | | **K03417** | **H** | **Catalyzes the thermodynamically favored C-C bond cleavage of (2R,3S)-2-methylisocitrate to yield pyruvate and succinate** | **prpB** | **91.4** | **0.58** | **5.29E-09** | **21.8** | **11.5** | **19.8** | **234.5** | **126.4** | **NA** |
| **stage2** | **stage3_free** | **-2.18** | **2.92E-03** | **V12B01_24654** | **EAP95001.1** | **citrate synthase** | **TCA cycle** | **K01659** | **C** | **Belongs to the citrate synthase family** | **prpC** | **55.9** | **0.70** | **1.45E-04** | **19.6** | **8.5** | **5.4** | **89.3** | **41.5** | **NA** |
| stage3_free | stage3_shell | 2.33 | 2.65E-03 | V12B01_24654 | EAP95001.1 | citrate synthase | TCA cycle | K01659 | C | Belongs to the citrate synthase family | prpC | 55.9 | 0.78 | 3.77E-04 | 89.3 | 41.5 | NA | 9.3 | 7.5 | 4.7 |
| **stage3_free** | **stage3_shell** | **-2.47** | **2.34E-05** | **V12B01_24684** | **EAP95007.1** | **acetyltransferase, putative** | | | **K** | **COG0454 Histone acetyltransferase HPA2 and related acetyltransferases** | **-** | **50.1** | **0.57** | **1.27E-06** | **23.8** | **22.0** | **NA** | **79.9** | **110.3** | **124.6** |
| **stage3_free** | **stage3_shell** | **-2.19** | **3.11E-05** | **V12B01_24709** | **EAP95012.1** | **putative GGDEF family protein** | **Regulation** | | **T** | **Tetratricopeptide repeat** | **-** | **26.9** | **0.50** | **1.86E-06** | **4.1** | **2.7** | **NA** | **13.7** | **15.2** | **8.6** |
| **stage2** | **stage3_shell** | **-2.76** | **4.22E-07** | **V12B01_24759** | **EAP91822.1** | **ISPsy11, transposase OrfA** | | **K07483** | **L** | **Evidence 2b Function of strongly homologous gene** | **-** | **12.0** | **0.53** | **2.01E-08** | **6.6** | **10.6** | **11.6** | **71.5** | **60.2** | **28.3** |
| **stage3_free** | **stage3_shell** | **-2.39** | **1.05E-04** | **V12B01_24759** | **EAP91822.1** | **ISPsy11, transposase OrfA** | | **K07483** | **L** | **Evidence 2b Function of strongly homologous gene** | **-** | **12.0** | **0.60** | **8.02E-06** | **13.7** | **10.8** | **NA** | **71.5** | **60.2** | **28.3** |
| **stage2** | **stage3_shell** | **-4.19** | **2.53E-13** | **V12B01_24874** | **EAP92078.1** | **WD40 repeat protein** | | | **-** | **-** | **VP1605** | **254.2** | **0.55** | **2.61E-15** | **11.7** | **39.4** | **20.5** | **429.8** | **464.5** | **183.2** |
| **stage3_free** | **stage3_shell** | **-4.75** | **9.31E-13** | **V12B01_24874** | **EAP92078.1** | **WD40 repeat protein** | | | **-** | **-** | **VP1605** | **254.2** | **0.64** | **6.09E-15** | **18.7** | **13.3** | **NA** | **429.8** | **464.5** | **183.2** |

| | | | | | | | | | | | | | | | | | | | |
|---|---|---|---|---|---|---|---|---|---|---|---|---|---|---|---|---|---|---|---|
| stage3_free | stage3_shell | 2.48 | 3.59E-06 | V12B01_24894 | EAP92082.1 | dihydroorotate dehydrogenase | Nucleoside and Nucleotide Biosynthesis | K00254 | F | Catalyzes the conversion of dihydroorotate to orotate with quinone as electron acceptor | pyrD | 75.5 | 0.50 | 1.33E-07 | 195.0 | 318.3 | NA | 45.8 | 26.9 | 16.9 |
| stage2 | stage3_shell | 2.39 | 2.05E-02 | V12B01_24959 | EAP96276.1 | hypothetical protein | | | S | Outer membrane protein beta-barrel domain | - | 7.6 | 1.09 | 5.69E-03 | 25.9 | 20.1 | 25.0 | 5.3 | 0.0 | 0.0 |
| stage3_free | stage3_shell | 2.10 | 6.64E-04 | V12B01_25034 | EAP96291.1 | Cbb3-type cytochrome oxidase, subunit 1 | Respiration | K00404 | C | Belongs to the heme-copper respiratory oxidase family | ccoN | 426.4 | 0.59 | 7.01E-05 | 942.1 | 778.5 | NA | 90.2 | 90.7 | 176.7 |
| stage3_free | stage3_shell | 2.04 | 4.19E-03 | V12B01_25039 | EAP96292.1 | Cbb3-type cytochrome oxidase, cytochrome c subunit | Respiration | K00405 | C | Cbb3-type cytochrome oxidase, cytochrome c subunit | ccoO | 186.3 | 0.72 | 6.54E-04 | 820.4 | 615.0 | NA | 33.4 | 104.8 | 151.3 |
| stage2 | stage3_free | -4.20 | 4.11E-08 | V12B01_25134 | EAP96311.1 | NAD-dependent deacetylase | | K12410 | K | NAD-dependent lysine deacetylase and desuccinylase that specifically removes acetyl and succinyl groups on target proteins. Modulates the activities of several proteins which are inactive in their acylated form | cobB | 426.9 | 0.72 | 3.89E-10 | 52.0 | 80.0 | 133.7 | 1877.9 | 2014.3 | NA |
| stage2 | stage3_shell | -2.77 | 2.50E-05 | V12B01_25134 | EAP96311.1 | NAD-dependent deacetylase | | K12410 | K | NAD-dependent lysine deacetylase and desuccinylase that specifically removes acetyl and succinyl groups on target proteins. Modulates the activities of several proteins which are inactive in their acylated form | cobB | 426.9 | 0.66 | 2.18E-06 | 52.0 | 80.0 | 133.7 | 1171.7 | 92.6 | 333.1 |
| stage2 | stage3_free | -6.86 | 1.60E-13 | V12B01_25139 | EAP96312.1 | Ammonia permease | Transport Proteins | K03320 | P | ammonium transporter | - | 71.7 | 0.87 | 3.87E-16 | 1.5 | 5.2 | 2.0 | 569.0 | 242.3 | NA |
| stage3_free | stage3_shell | 5.20 | 1.37E-07 | V12B01_25139 | EAP96312.1 | Ammonia permease | Transport Proteins | K03320 | P | ammonium transporter | - | 71.7 | 0.92 | 3.07E-09 | 569.0 | 242.3 | NA | 9.2 | 9.9 | 0.0 |
| stage2 | stage3_shell | -2.32 | 4.80E-05 | V12B01_25144 | EAP96313.1 | hypothetical protein | | | - | - | VV1697 | 172.1 | 0.56 | 4.51E-06 | 106.6 | 251.4 | 175.9 | 452.1 | 1567.3 | 307.3 |
| stage2 | stage3_shell | -2.71 | 2.75E-07 | V12B01_25149 | EAP96314.1 | formate dehydrogenase, cytochrome b556 subunit | Respiration | K00127 | C | COG2864 Cytochrome b subunit of formate dehydrogenase | fdnI | 468.5 | 0.51 | 1.25E-08 | 106.0 | 195.6 | 156.3 | 325.8 | 1746.7 | 450.7 |
| stage2 | stage3_shell | -3.34 | 4.56E-08 | V12B01_25154 | EAP96315.1 | formate dehydrogenase, iron-sulfur subunit | Respiration | K00124 | C | formate dehydrogenase iron-sulfur subunit | fdhB | 371.2 | 0.60 | 1.58E-09 | 115.4 | 221.0 | 118.9 | 280.7 | 2695.4 | 935.0 |
| stage2 | stage3_shell | -2.78 | 1.13E-06 | V12B01_25159 | EAP96316.1 | putative formate dehydrogenase large subunit | Respiration | K00123 | C | Belongs to the prokaryotic molybdopterin-containing oxidoreductase family | fdhA | 1858.9 | 0.56 | 6.18E-08 | 164.6 | 245.4 | 120.2 | 250.1 | 2192.5 | 657.0 |
| stage2 | stage3_shell | -2.51 | 8.90E-06 | V12B01_25164 | EAP96317.1 | hypothetical protein | | | S | TIGRFAM formate dehydrogenase region TAT target | - | 186.9 | 0.56 | 6.67E-07 | 154.9 | 442.3 | 340.0 | 413.3 | 2980.6 | 1133.2 |
| stage2 | stage3_free | -2.03 | 3.49E-04 | V12B01_25169 | EAP96318.1 | putative formate dehydrogenase-specific chaperone | | | S | component of anaerobic dehydrogenases | - | 193.0 | 0.53 | 1.06E-05 | 62.5 | 168.9 | 114.3 | 553.1 | 557.2 | NA |
| stage2 | stage3_shell | -2.26 | 2.95E-06 | V12B01_25169 | EAP96318.1 | putative formate dehydrogenase-specific chaperone | Respiration | | S | component of anaerobic dehydrogenases | - | 193.0 | 0.47 | 1.81E-07 | 62.5 | 168.9 | 114.3 | 246.0 | 870.6 | 263.9 |
| stage2 | stage3_free | -2.37 | 1.48E-05 | V12B01_25174 | EAP96319.1 | iron-sulfur cluster-binding protein | | | C | 4Fe-4S dicluster domain | - | 408.7 | 0.50 | 2.48E-07 | 61.0 | 129.8 | 83.6 | 528.6 | 555.2 | NA |
| stage2 | stage3_shell | -2.10 | 7.20E-06 | V12B01_25174 | EAP96319.1 | iron-sulfur cluster-binding protein | | | C | 4Fe-4S dicluster domain | - | 408.7 | 0.45 | 5.12E-07 | 61.0 | 129.8 | 83.6 | 317.2 | 581.7 | 94.2 |
| stage2 | stage3_shell | -2.09 | 5.75E-05 | V12B01_25179 | EAP96320.1 | hypothetical protein | | | S | Protein of unknown function (DUF3306) | - | 115.6 | 0.51 | 5.59E-06 | 29.6 | 98.6 | 66.0 | 215.3 | 414.4 | 78.4 |
| stage2 | stage3_shell | -2.38 | 8.94E-09 | V12B01_25184 | EAP96321.1 | hypothetical protein | | | S | Protein of unknown function (DUF3305) | - | 147.9 | 0.40 | 2.61E-10 | 55.5 | 169.9 | 124.2 | 405.5 | 709.3 | 330.6 |
| stage3_free | stage3_shell | -2.11 | 4.86E-03 | V12B01_25234 | EAP96331.1 | Methyl-accepting chemotaxis protein | Chemotaxis | K03406 | NT | methyl-accepting chemotaxis protein | - | 153.5 | 0.80 | 7.77E-04 | 15.0 | 22.1 | NA | 46.5 | 191.3 | 6.0 |
| stage2 | stage3_free | -2.26 | 2.55E-03 | V12B01_25254 | EAP96335.1 | Methyl-accepting chemotaxis protein | Chemotaxis | K03406 | T | methyl-accepting chemotaxis protein | - | 156.6 | 0.72 | 1.24E-04 | 35.7 | 29.7 | 23.7 | 211.2 | 161.0 | NA |
| stage2 | stage3_shell | -2.32 | 2.67E-04 | V12B01_25254 | EAP96335.1 | Methyl-accepting chemotaxis protein | Chemotaxis | K03406 | T | methyl-accepting chemotaxis protein | - | 156.6 | 0.64 | 3.20E-05 | 35.7 | 29.7 | 23.7 | 94.7 | 85.9 | 188.1 |
| stage2 | stage3_free | -2.30 | 1.07E-08 | V12B01_25269 | EAP96338.1 | hypothetical protein | | K05772 | H | COG2998 ABC-type tungstate transport system, permease component | tupA | 114.5 | 0.38 | 8.93E-11 | 88.7 | 154.0 | 83.1 | 587.7 | 582.1 | NA |
| stage2 | stage3_free | -2.42 | 1.05E-05 | V12B01_25274 | EAP96339.1 | ABC transporter, permease protein | Transport Proteins | K05773 | P | COG4662 ABC-type tungstate transport system, periplasmic component | - | 32.3 | 0.51 | 1.66E-07 | 24.2 | 24.3 | 15.2 | 123.0 | 138.1 | NA |
| stage2 | stage3_shell | -2.91 | 1.01E-22 | V12B01_25329 | EAP96350.1 | hypothetical protein | | | G | COG3001 Fructosamine-3-kinase | - | 599.2 | 0.29 | 1.93E-25 | 146.7 | 160.1 | 208.0 | 702.1 | 1275.6 | 922.8 |
| stage3_free | stage3_shell | -2.52 | 7.39E-14 | V12B01_25329 | EAP96350.1 | hypothetical protein | | | G | COG3001 Fructosamine-3-kinase | - | 599.2 | 0.32 | 3.53E-16 | 212.6 | 235.5 | NA | 702.1 | 1275.6 | 922.8 |
| stage2 | stage3_shell | -3.04 | 1.10E-08 | V12B01_25334 | EAP96351.1 | hypothetical protein | | | S | Cysteine-rich CPXCG | - | 315.0 | 0.52 | 3.29E-10 | 130.8 | 229.2 | 293.7 | 446.4 | 1490.6 | 2191.2 |
| stage3_free | stage3_shell | -2.52 | 2.60E-05 | V12B01_25334 | EAP96351.1 | hypothetical protein | | | S | Cysteine-rich CPXCG | - | 315.0 | 0.59 | 1.47E-06 | 251.1 | 353.2 | NA | 446.4 | 1490.6 | 2191.2 |
| stage2 | stage3_shell | -2.69 | 2.38E-04 | V12B01_25354 | EAP96355.1 | Anti-anti-sigma regulatory factor | Regulation | K04749 | T | COG1366 Anti-anti-sigma regulatory factor (antagonist of anti-sigma factor) | spoIIAA | 9.7 | 0.74 | 2.81E-05 | 6.7 | 6.4 | 0.9 | 19.4 | 41.3 | 17.3 |
| stage3_free | stage3_shell | -2.01 | 1.00E-04 | V12B01_25364 | EAP96357.1 | periplasmic protein involved in polysaccharide export | Transport Proteins | | M | Periplasmic protein involved in polysaccharide export | - | 100.5 | 0.50 | 7.41E-06 | 10.5 | 10.5 | NA | 12.2 | 59.8 | 31.2 |
| stage2 | stage3_shell | -2.58 | 5.09E-05 | V12B01_25369 | EAP96358.1 | ATPase involved in chromosome partitioning | Replication | | D | COG0489 ATPases involved in chromosome partitioning | - | 24.9 | 0.64 | 4.80E-06 | 6.4 | 7.2 | 2.3 | 18.5 | 30.7 | 33.0 |
| stage2 | stage3_shell | -2.51 | 1.77E-10 | V12B01_25549 | EAP96394.1 | hypothetical protein | | | - | #N/A | #N/A | 143.6 | 0.38 | 3.13E-12 | 43.8 | 59.4 | 77.9 | 294.8 | 412.3 | 113.4 |
| stage3_free | stage3_shell | -3.13 | 8.21E-12 | V12B01_25549 | EAP96394.1 | hypothetical protein | | | - | #N/A | #N/A | 143.6 | 0.44 | 6.76E-14 | 33.6 | 45.3 | NA | 294.8 | 412.3 | 113.4 |
| stage2 | stage3_shell | 2.36 | 1.98E-03 | V12B01_25559 | EAP96396.1 | hypothetical protein | | | - | - | - | 40.0 | 0.81 | 3.33E-04 | 89.9 | 121.3 | 194.5 | 12.6 | 9.9 | 22.4 |
| stage2 | stage3_shell | -4.26 | 6.34E-15 | V12B01_25564 | EAP96397.1 | Transposase | | K07486 | L | COG3547 Transposase and inactivated derivatives | - | 64.7 | 0.53 | 5.08E-17 | 3.9 | 12.1 | 8.2 | 63.8 | 103.2 | 188.7 |
| stage3_free | stage3_shell | -2.60 | 1.34E-05 | V12B01_25564 | EAP96397.1 | Transposase | | K07486 | L | COG3547 Transposase and inactivated derivatives | - | 64.7 | 0.59 | 6.66E-07 | 27.2 | 21.5 | NA | 63.8 | 103.2 | 188.7 |
| stage2 | stage3_shell | 2.33 | 3.10E-03 | V12B01_25569 | EAP96398.1 | hypothetical protein | | | - | #N/A | #N/A | 19.4 | 0.78 | 5.78E-04 | 61.8 | 54.1 | 67.5 | 9.5 | 9.0 | 0.0 |
| stage2 | stage3_shell | -3.57 | 8.67E-09 | V12B01_25574 | EAP96399.1 | Transposase | | K07486 | L | COG3547 Transposase and inactivated derivatives | - | 56.4 | 0.60 | 2.51E-10 | 2.2 | 13.5 | 5.8 | 28.0 | 88.5 | 91.6 |
| stage2 | stage3_free | -4.15 | 1.30E-04 | V12B01_25584 | EAP96401.1 | arginine ABC transporter, permease protein | Transport Proteins | K09998 | P | COG4160 ABC-type arginine histidine transport system, permease component | artM | 27.3 | 1.02 | 3.23E-06 | 8.6 | 3.2 | 7.9 | 225.6 | 99.1 | NA |

| | | | | | | | | | | | | | | | | | | | |
|---|---|---|---|---|---|---|---|---|---|---|---|---|---|---|---|---|---|---|---|
| stage2 | stage3_free | -3.24 | 2.13E-06 | V12B01_25589 | EAP96402.1 | arginine ABC transporter, permease protein | Transport Proteins | K09999 | P | COG4215 ABC-type arginine transport system, permease component | artQ | 108.0 | 0.64 | 2.90E-08 | 14.2 | 55.7 | 51.8 | 604.5 | 305.6 | NA |
| stage2 | stage3_free | -4.64 | 3.52E-19 | V12B01_25594 | EAP96403.1 | ABC-type amino acid transport/signal transduction system | Transport Proteins | K09997 | ET | Belongs to the bacterial solute-binding protein 3 family | artI | 100.0 | 0.49 | 1.55E-22 | 23.1 | 22.3 | 25.2 | 765.4 | 510.5 | NA |
| stage3_free | stage3_shell | 3.02 | 2.75E-08 | V12B01_25594 | EAP96403.1 | ABC-type amino acid transport/signal transduction system | Transport Proteins | K09997 | ET | Belongs to the bacterial solute-binding protein 3 family | artI | 100.0 | 0.52 | 4.82E-10 | 765.4 | 510.5 | NA | 45.8 | 20.7 | 89.5 |
| stage2 | stage3_free | -3.61 | 2.01E-04 | V12B01_25599 | EAP96404.1 | arginine ABC transporter, ATP-binding protein | Transport Proteins | K10000 | E | COG4161 ABC-type arginine transport system, ATPase component | artP | 98.8 | 0.92 | 5.63E-06 | 22.0 | 44.7 | 24.8 | 427.9 | 589.1 | NA |
| stage2 | stage3_shell | 2.54 | 6.86E-06 | V12B01_25609 | EAP96406.1 | hypothetical protein | | K07107 | S | Thioesterase-like superfamily | - | 32.0 | 0.54 | 4.85E-07 | 239.1 | 169.9 | 140.9 | 27.5 | 29.3 | 0.0 |
| stage3_free | stage3_shell | -2.30 | 7.41E-07 | V12B01_25694 | EAP96423.1 | hypothetical protein | | | S | Domain of unknown function DUF302 | - | 585.8 | 0.44 | 2.25E-08 | 434.4 | 574.6 | NA | 1907.6 | 2524.8 | 1476.3 |
| stage3_free | stage3_shell | 2.21 | 1.64E-03 | V12B01_25709 | EAP96426.1 | hypothetical protein | | | - | - | - | 23.2 | 0.67 | 2.07E-04 | 98.9 | 90.4 | NA | 0.0 | 23.1 | 14.5 |
| stage2 | stage3_shell | 2.00 | 1.61E-03 | V12B01_25799 | EAP96444.1 | hypothetical protein | | K09962 | S | Nucleotidyltransferase | - | 23.5 | 0.61 | 2.62E-04 | 78.9 | 52.0 | 69.3 | 14.3 | 13.6 | 0.0 |
| stage3_free | stage3_shell | 2.70 | 2.38E-04 | V12B01_25834 | EAP96451.1 | Glutathione-dependent formaldehyde-activating, GFA | | | S | Evidence 3 Function proposed based on presence of conserved amino acid motif, structural feature or limited homology | - | 17.1 | 0.70 | 2.10E-05 | 112.7 | 109.1 | NA | 0.0 | 18.3 | 13.8 |
| stage2 | stage3_shell | -3.83 | 1.57E-10 | V12B01_25869 | EAP96458.1 | Transposase | | K07486 | L | COG3547 Transposase and inactivated derivatives | - | 56.4 | 0.58 | 2.71E-12 | 3.0 | 7.6 | 8.5 | 34.3 | 69.3 | 113.8 |
| stage3_free | stage3_shell | 2.11 | 2.12E-02 | V12B01_25879 | EAP96460.1 | acetyltransferase | | K00633 | S | COG0110 Acetyltransferase (isoleucine patch superfamily) | - | 7.7 | 0.95 | 5.09E-03 | 34.0 | 29.8 | NA | 5.0 | 4.7 | 0.0 |
| stage3_free | stage3_shell | 2.60 | 4.65E-02 | V12B01_25894 | EAP96463.1 | hypothetical protein | | | - | - | - | 2.5 | 2.47 | 1.35E-02 | 42.3 | 41.5 | NA | 0.0 | 0.0 | 0.0 |
| stage3_free | stage3_shell | -3.93 | 1.46E-02 | V12B01_25974 | EAP96479.1 | deoxyribodipyrimidine photolyase | | K01669 | L | Belongs to the DNA photolyase family | cry | 29.4 | 1.70 | 3.18E-03 | 0.3 | 0.0 | NA | 0.0 | 8.7 | 4.1 |
| stage2 | stage3_free | 3.37 | 4.03E-02 | V12B01_25994 | EAP96483.1 | hypothetical protein | | | S | protein conserved in bacteria | - | 8.5 | 1.40 | 4.04E-03 | 21.6 | 41.8 | 45.5 | 3.2 | 0.0 | NA |
| stage3_free | stage3_shell | -3.77 | 1.46E-02 | V12B01_25994 | EAP96483.1 | hypothetical protein | | | S | protein conserved in bacteria | - | 8.5 | 1.53 | 3.18E-03 | 3.2 | 0.0 | NA | 11.1 | 52.5 | 39.5 |
| stage2 | stage3_free | -2.85 | 6.23E-05 | V12B01_26009 | EAP96486.1 | hypothetical protein | | | M | OmpA-like transmembrane domain | - | 18.4 | 0.67 | 1.39E-06 | 8.4 | 9.2 | 16.2 | 97.2 | 102.9 | NA |
| stage3_free | stage3_shell | -2.21 | 4.67E-02 | V12B01_26044 | EAP96493.1 | hypothetical protein | | | Q | Methyltransferase domain | - | 4.0 | 1.55 | 1.36E-02 | 1.2 | 0.0 | NA | 4.2 | 4.0 | 7.5 |
| stage2 | stage3_shell | 2.35 | 8.29E-12 | V12B01_26059 | EAP96496.1 | chitinase A | Carbohydrates and Carboxylates Degradation | K01183 | G | Domain of unknown function (DUF5011) | - | 197.5 | 0.33 | 1.21E-13 | 249.1 | 184.4 | 191.1 | 19.0 | 35.9 | 31.2 |
| stage2 | stage3_shell | 3.27 | 1.79E-04 | V12B01_26064 | EAP96497.1 | acetyltransferase | | | L | single-stranded DNA 5'-3' exodeoxyribonuclease activity | - | 79.3 | 0.87 | 2.02E-05 | 62.7 | 37.7 | 174.2 | 3.2 | 11.9 | 0.0 |
| stage3_free | stage3_shell | 2.14 | 1.60E-02 | V12B01_26064 | EAP96497.1 | acetyltransferase | | | L | single-stranded DNA 5'-3' exodeoxyribonuclease activity | - | 79.3 | 0.95 | 3.55E-03 | 49.2 | 46.4 | NA | 3.2 | 11.9 | 0.0 |
| stage2 | stage3_shell | 2.50 | 7.09E-03 | V12B01_26069 | EAP96498.1 | PAS factor | | | S | Pas factor saposin fold | - | 8.3 | 1.00 | 1.54E-03 | 137.1 | 155.0 | 66.3 | 0.0 | 12.6 | 23.7 |
| stage3_free | stage3_shell | 2.24 | 7.87E-03 | V12B01_26089 | EAP96502.1 | transcriptional regulator, LysR family | Regulation | | K | Transcriptional regulator | VVA1234 | 18.0 | 0.84 | 1.43E-03 | 38.4 | 30.9 | NA | 0.0 | 10.4 | 0.0 |
| stage2 | stage3_shell | -3.43 | 7.46E-07 | V12B01_26094 | EAP96503.1 | adenylosuccinate synthetase | Nucleoside and Nucleotide Biosynthesis | K01939 | F | Plays an important role in the de novo pathway of purine nucleotide biosynthesis. Catalyzes the first committed step in the biosynthesis of AMP from IMP | VVA1235 | 34.6 | 0.69 | 3.85E-08 | 2.8 | 3.7 | 1.5 | 5.1 | 31.2 | 36.2 |
| stage2 | stage3_free | 3.86 | 6.93E-03 | V12B01_26149 | EAP96514.1 | hypothetical protein | | K02282 | D | COG4963 Flp pilus assembly protein, ATPase CpaE | - | 5.4 | 1.36 | 4.09E-04 | 5.7 | 8.6 | 14.6 | 0.0 | 0.8 | NA |
| stage2 | stage3_free | 3.66 | 7.21E-03 | V12B01_26174 | EAP96519.1 | hypothetical protein | | K02651 | U | Flp/Fap pilin component | - | 12.5 | 1.61 | 4.36E-04 | 64.4 | 407.2 | 262.3 | 4.4 | 16.2 | NA |
| stage2 | stage3_shell | 2.91 | 1.34E-02 | V12B01_26174 | EAP96519.1 | hypothetical protein | | K02651 | U | Flp/Fap pilin component | - | 12.5 | 1.42 | 3.38E-03 | 64.4 | 407.2 | 262.3 | 7.7 | 29.2 | 0.0 |
| stage2 | stage3_free | 2.04 | 1.84E-03 | V12B01_26359 | EAP91655.1 | putative outer membrane protein | Outer membrane | K22110 | M | Oligogalacturonate-specific porin protein (KdgM) | kdgM | 18.5 | 0.65 | 8.62E-05 | 121.1 | 226.1 | 157.0 | 40.9 | 26.2 | NA |
| stage2 | stage3_shell | 5.17 | 7.27E-06 | V12B01_26359 | EAP91655.1 | putative outer membrane protein | Outer membrane | K22110 | M | Oligogalacturonate-specific porin protein (KdgM) | - | 18.5 | 1.08 | 5.19E-07 | 121.1 | 226.1 | 157.0 | 0.0 | 6.6 | 0.0 |
| stage3_free | stage3_shell | 2.48 | 2.09E-02 | V12B01_26359 | EAP91655.1 | putative outer membrane protein | Outer membrane | K22110 | M | Oligogalacturonate-specific porin protein (KdgM) | - | 18.5 | 1.12 | 4.99E-03 | 40.9 | 26.2 | NA | 0.0 | 6.6 | 0.0 |
